# SNP-to-gene linking strategies reveal contributions of enhancer-related and candidate master-regulator genes to autoimmune disease

**DOI:** 10.1101/2020.09.02.279059

**Authors:** Kushal K. Dey, Steven Gazal, Bryce van de Geijn, Samuel Sungil Kim, Joseph Nasser, Jesse M. Engreitz, Alkes L. Price

## Abstract

Gene regulation is known to play a fundamental role in human disease, but mechanisms of regulation vary greatly across genes. Here, we explore the contributions to disease of two types of genes: genes whose regulation is driven by enhancer regions as opposed to promoter regions (enhancer-related) and genes that regulate other genes in trans (candidate master-regulator). We link these genes to SNPs using a comprehensive set of SNP-to-gene (S2G) strategies and apply stratified LD score regression to the resulting SNP annotations to draw three main conclusions about 11 autoimmune diseases and blood cell traits (average *N*_case_=13K across 6 autoimmune diseases, average *N* =443K across 5 blood cell traits). First, several characterizations of enhancer-related genes defined in blood using functional genomics data (e.g. ATAC-seq, RNA-seq, PC-HiC) are conditionally informative for autoimmune disease heritability, after conditioning on a broad set of regulatory annotations from the baseline-LD model. Second, candidate master-regulator genes defined using trans-eQTL in blood are also conditionally informative for autoimmune disease heritability. Third, integrating enhancer-related and candidate master-regulator gene sets with protein-protein interaction (PPI) network information magnified their disease signal. The resulting PPI-enhancer gene score produced *>*2x stronger conditional signal (maximum standardized SNP annotation effect size (*τ^*^*) = 2.0 (s.e. 0.3) vs. 0.91 (s.e. 0.21)), and *>*2x stronger gene-level enrichment for approved autoimmune disease drug targets (5.3x vs. 2.1x), as compared to the recently proposed Enhancer Domain Score (EDS). In each case, using functionally informed S2G strategies to link genes to SNPs that may regulate them produced much stronger disease signals (4.1x-13x larger *τ^*^* values) than conventional window-based S2G strategies. We conclude that our characterizations of enhancer-related and candidate master-regulator genes identify gene sets that are important for autoimmune disease, and that combining those gene sets with functionally informed S2G strategies enables us to identify SNP annotations in which disease heritability is concentrated.

## 1 Introduction

Disease risk variants associated with complex traits and diseases predominantly lie in non-coding regulatory regions of the genes, motivating the need to assess the relative importance of genes for disease through the lens of gene regulation^1–6^. Several recent studies have performed disease-specific gene-level prioritization by integrating GWAS summary statistics data with functional genomics data, including gene expression and gene networks^7–14^. Here, we investigate the contribution to autoimmune disease of gene sets reflecting two specific aspects of gene regulation in blood—genes with strong evidence of enhancer-related regulation and candidate master-regulator genes that may potentially regulate many other genes; previous studies suggested that both of these characterizations are important for understanding human disease^9, 15–24^. For example, several common non-coding variants associated with Hirschsprung disease have been identified in intronic enhancer elements of *RET* gene and have been shown to synergistically regulate its expression^25, 26^ and *NLRC5* acts as a master regulator of MHC class genes in immune response^27^. Our two main goals are to characterize which types of genes are important for autoimmune disease, and to construct SNP annotations derived from those genes that are conditionally informative for disease heritability, conditional on all other annotations.

A major challenge in gene-level analyses of disease is to link genes to SNPs that may regulate them, a prerequisite to integrative analyses of GWAS summary statistics. Previous studies have often employed window-based strategies such as *±*100kb^8, 9, 11^, linking each gene to all SNPs within 100kb; however, this approach lacks specificity. Here, we incorporated functionally informed SNP-to-gene (S2G) linking strategies that capture both distal and proximal components of gene regulation. We evaluated the resulting SNP annotations by applying stratified LD score regression^28^ (S-LDSC) conditional on a broad set of coding, conserved, regulatory and LD-related annotations from the baseline-LD model^29, 30^, meta-analyzing the results across 11 autoimmune diseases and blood cell traits; we focused on autoimmune diseases and blood cell traits because the functional data underlying the gene scores and S2G strategies that we analyze is primarily measured in blood. We also assessed gene-level enrichment for disease-related gene sets, including approved drug targets for autoimmune disease^10^.

### Results

#### Overview of methods

We define an *annotation* as an assignment of a numeric value to each SNP with minor allele count *≥*5 in a 1000 Genomes Project European reference panel^31^, as in our previous work^28^; we primarily focus on annotations with values between 0 and 1. We define a *gene score* as an assignment of a numeric value between 0 and 1 to each gene; gene scores predict the relevance of each gene to disease. We primarily focus on binary gene sets defined by the top 10% of genes; we made this choice to be consistent with ref.^9^, and to ensure that all resulting SNP annotations (gene scores x S2G strategies; see below) were of reasonable size (0.2% of SNPs or larger). We consider 11 gene scores prioritizing enhancer-related genes, candidate master-regulator genes, and genes with high network connectivity to enhancer-related or candidate master-regulator genes (Table 1, Supplementary Figure S1); these gene scores were only mildly correlated (average *r*= 0.08, Supplementary Figure S2). We considered enhancer-related and candidate master-regulator genes because previous studies suggested that both of these characterizations are important for understanding human disease^9, 15–24^.

**Table 1.** List of 11 gene scores. : For each gene score, including (A) 7 enhancer-related genes scores (red font), (B) 2 candidate master-regulator gene scores (blue font) and (C) 2 PPI-network informed gene scores (corresponding to enhancer-related and candidate master-regulator gene scores), we provide a brief description and report its size (average gene score across 22,020 genes; equal to % of genes for binary gene scores). Gene scores are listed alphabetically within each category. All gene scores are binary except ATAC-distal and eQTL-CTS, which are probabilistic. Density plots of the distribution of the metric underlying each gene score are provided in Figure S1. Further details are provided in the Methods section.

| Gene score | Description | Size (%) |
| --- | --- | --- |
| (A) Enhancer-related gene scores |  |  |
| **ABC-G** | Genes in top 10% genes of number of genic and intergenic enhancer-gene connections in blood, assessed using Activity-By-Contact<sup>32</sup> | 10% |
| **ATAC-distal** | Proportion of mouse gene expression variability across immune cell types explained by distal ATAC-seq peaks<sup>48</sup> | 29% |
| **EDS-binary** | Genes in top 10% of blood-specific Enhancer Domain Score (EDS), reflecting the number of bases in enhancers linked to a gene<sup>24</sup> | 10% |
| **eQTL-CTS** | Proportion of eQTLs<sup>99</sup> (FDR < 0.05) that are specific to a single cell type (union across blood cell types). | 32% |
| **Expecto-MVP** | Genes in top 10% of magnitude of variation potential (MVP), based on Expecto $\Delta$ predictions of regions surrounding the TSS<sup>7</sup>. | 10% |
| **PC-HiC-distal** | Genes in top 10% genes of number of distal Promoter Capture HiC (PC-HiC) connections in blood cell types<sup>108</sup>. | 10% |
| **SEG-GTEx** | Specifically expressed genes<sup>9</sup>(SEG) in GTEx whole blood<sup>112</sup> | 10% |
| (B) candidate master-regulator gene scores |  |  |
| **Trans-master** | Genes that significantly trans-regulate $\geq 3$ genes by any significant cis-eQTL of the focal gene | 10% |
| **TF** | Curated list of human Transcription Factor genes<sup>57</sup> | 7.4% |
| (C) PPI network-based gene score |  |  |
| **PPI-enhancer** | Genes with high network connectivity to enhancer-related genes in STRING<sup>53</sup> PPI network | 10% |
| **PPI-master** | Genes with high network connectivity to candidate master-regulator genes in STRING<sup>53</sup> PPI network | 10% |

We define a *SNP-to-gene (S2G) linking strategy* as an assignment of 0, 1 or more linked genes to each SNP. We consider 10 S2G strategies capturing both distal and proximal gene regulation (see Methods, Figure 1A and Table 2); these S2G strategies aim to link SNPs to genes that they regulate. For each gene score X and S2G strategy Y, we define a corresponding *combined* annotation X *×* Y by assigning to each SNP the maximum gene score among genes linked to that SNP (or 0 for SNPs with no linked genes); this generalizes the standard approach of constructing annotations from gene scores using window-based strategies^8, 9^. For example, EDS-binary *×* ABC annotates SNPs linked by Activity-By-Contact enhancer-gene links^32, 33^ to any gene from the EDS-binary gene set, whereas EDS-binary *×* 100kb annotates all SNPs within 100kb of any gene from the EDS-binary gene set. We considered combined annotations based on S2G strategies related to gene regulation because SNPs that regulate functionally important genes may be important for disease. For each S2G strategy, we also define a corresponding binary S2G annotation defined by SNPs linked to the set of all genes. We have publicly released all gene scores, S2G links, and annotations analyzed in this study (see URLs). We have also included annotations for 93 million Haplotype Reference Consortium (HRC) SNPs^34^ (MAF *>* 0.1% in imputed UK Biobank data^35^) and 170 million TOPMed SNPs^36^ (Freeze 3A).

**Figure 1.**
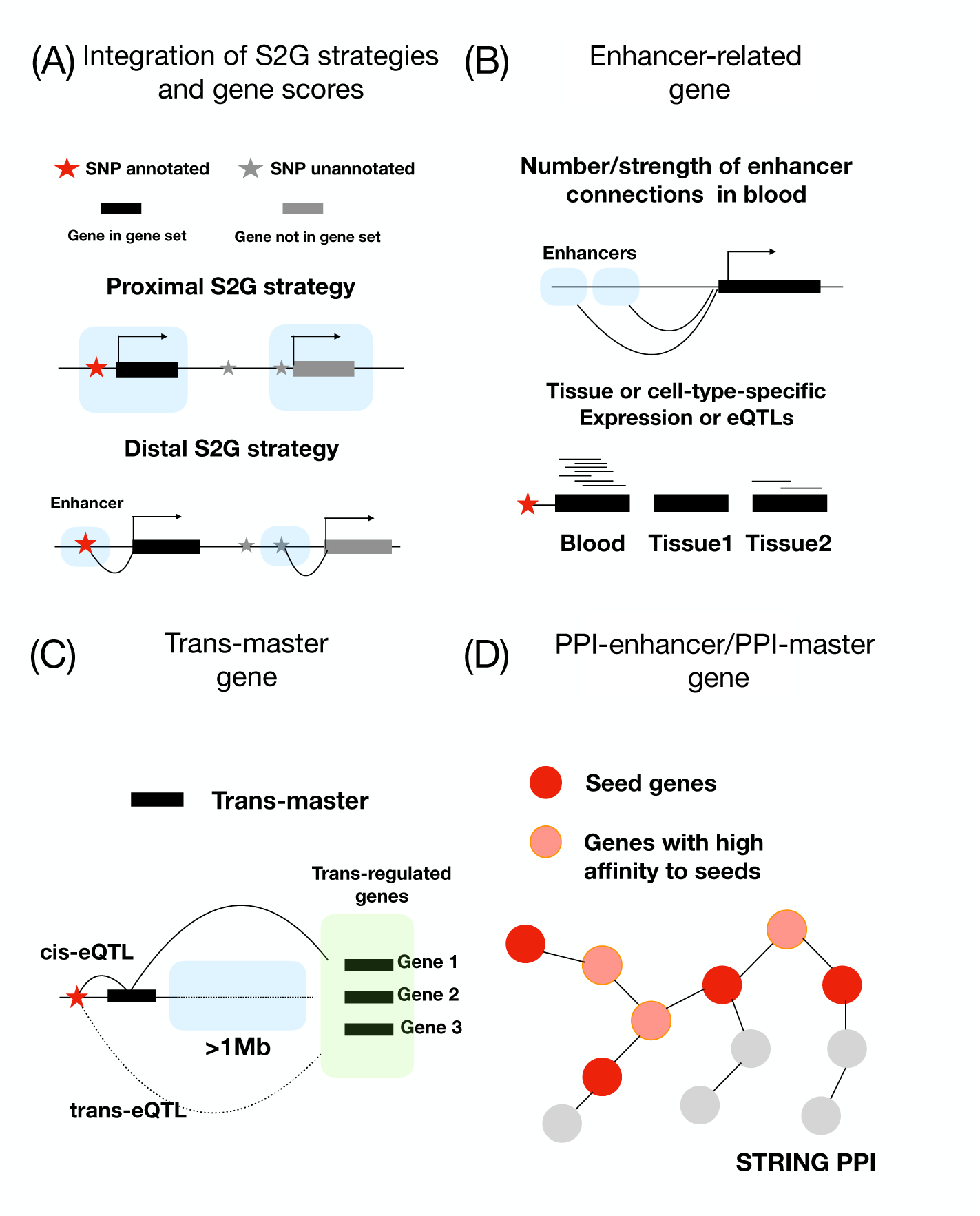
Illustration of S2G strategies and gene scores: (A) SNP annotations defined by integration of genes in gene set with proximal (close to gene body) and distal S2G strategies. (B) Examples of approaches used to define enhancer-related genes. (C) A Trans-master gene regulates multiple distal genes, via a cis-eQTL that is a trans-eQTL of the distal genes. (D) PPI-Enhancer genes have high connectivity to enhancer-related genes in a PPI network.

**Table 2.** List of 10 S2G strategies: For each S2G strategy, we provide a brief description, indicate whether the S2G strategy prioritizes distal or proximal SNPs relative to the gene, and report its size (% of SNPs linked to genes). S2G strategies are listed in order of increasing size. Further details are provided in the Methods section.

| S2G strategy | Description | Distal/<br>Proximal | Size (%) |
| --- | --- | --- | --- |
| ABC | Inter-genic SNPs with distal enhancer-gene connections, assessed by Activity-By-Contact<sup>32,33</sup> across blood cell types. | Distal | 1.4 |
| TSS | SNPs in predicted Transcription start sites<sup>103,104</sup> overlapping Ensembl gene±5kb window. | Proximal | 1.6 |
| Coding | SNPs in coding regions | Proximal | 1.6 |
| ATAC | SNPs in ATAC-seq peaks >50% correlated to mouse expression across blood cell-types<sup>48</sup> (mapped to human). | Distal | 1.6 |
| eQTL | SNPs with fine-mapped causal posterior probability<sup>37</sup> (CPP) >0.001 in GTEx whole blood. | Distal<br>+Proximal | 2.4 |
| Roadmap | SNPs in predicted enhancer-gene links, assessed using Roadmap Epigenomics data<sup>98,107</sup>. | Distal | 3.2 |
| Promoter | SNPs in promoter regions. | Proximal | 4.3 |
| PC-HiC | Distal SNPs with Promoter Capture HiC (PC-HiC)<sup>108</sup> connections to promoter regions in blood cell-types. | Distal | 27 |
| 5kb | SNPs in ±5kb window around gene body. | Proximal | 53 |
| 100kb | SNPs in ±100kb window around gene body<sup>8,9,11</sup>. | Distal | 81 |

We assessed the informativeness of the resulting annotations for disease heritability by applying stratified LD score regression (S-LDSC)^28^ to 11 independent blood-related traits (6 autoimmune diseases and 5 blood cell traits; average *N_case_*=13K for autoimmune diseases and *N* =443K for blood cell traits, Supplementary Table S1) and meta-analyzing S-LDSC results across traits; we also assessed results meta-analyzed across autoimmune diseases or blood cell traits only, as well as results for individual diseases/traits. We conditioned on 86 coding, conserved, regulatory and LD-related annotations from the baseline-LD model (v2.1)^29, 30^ (see URLs). S-LDSC uses two metrics to evaluate informativeness for disease heritability: enrichment and standardized effect size (*τ\**). Enrichment is defined as the proportion of heritability explained by SNPs in an annotation divided by the proportion of SNPs in the annotation^28^, and generalizes to annotations with values between 0 and 1^37^. Standardized effect size (*τ\**) is defined as the proportionate change in per-SNP heritability associated with a 1 standard deviation increase in the value of the annotation, conditional on other annotations included in the model^29^; unlike enrichment, *τ^*^* quantifies effects that are *conditionally informative*, i.e. unique to the focal annotation conditional on other annotations included in the model. In our “marginal” analyses, we estimated *τ^*^* for each focal annotation conditional on the baseline-LD annotations. In our “joint” analyses, we merged baseline-LD annotations with focal annotations that were marginally significant after Bonferroni correction and performed forward stepwise elimination to iteratively remove focal annotations that had conditionally non-significant *τ^*^* values after Bonferroni correction, as in ref.^11, 29, 37–42^. We did not consider other feature selection methods, as previous research determined that a LASSO-based feature selection method is computationally expensive and did not perform better in predicting off-chromosome *χ*^2^ association statistics (R. Cui and H. Finucane, personal correspondence). The difference between marginal *τ^*^* and joint *τ^*^* is that marginal *τ^*^* assesses informativeness for disease conditional only on baseline-LD model annotations, whereas joint *τ^*^* assesses informativeness for disease conditional on baseline-LD model annotations as well as other annotations in the joint model.

As a preliminary assessment of the potential of the 10 S2G strategies, we considered the 10 S2G annotations defined by SNPs linked to the set of all genes. The S2G annotations were only weakly positively correlated (average *r* = 0.09; Supplementary Figure S3). We analyzed the 10 S2G annotations via a marginal analysis, running S-LDSC^28^ conditional on the baseline-LD model and meta-analyzing the results across the 11 blood-related traits. In the marginal analysis, all 10 S2G annotations were significantly enriched for disease heritability, with larger enrichments for smaller annotations (Figure 2A and Supplementary Table S2); values of standardized enrichment (defined as enrichment scaled by the standard deviation of the annotation^11^) were more similar across annotations (Supplementary Figure S4 and Supplementary Table S3). 7 S2G annotations attained conditionally significant *τ^*^* values after Bonferroni correction (*p <* 0.05*/*10) (Figure 2B and Supplementary Table S2). In the joint analysis, 3 of these 7 S2G annotations were jointly significant: TSS (joint *τ^*^* = 0.97), Roadmap (joint *τ^*^* = 0.84) and Activity-by-Contact (ABC) (joint *τ^*^* = 0.44) (Figure 2B and Supplementary Table S4). This suggests that these 3 S2G annotations are highly informative for disease. Subsequent analyses were conditioned on the baseline-LD+ model defined by 86 baseline-LD model annotations plus all S2G annotations (except Coding, TSS and Promoter, which were already part of the baseline-LD model), to ensure that conditionally significant *τ^*^* values for (gene scores x S2G strategies) annotations are specific to the gene scores and cannot be explained by (all genes x S2G strategies) annotations. Accordingly, we confirmed that (random genes x S2G strategies) annotations did not produce conditionally significant *τ^*^* values for any S2G strategy (Supplementary Table S5).

**Figure 2.**
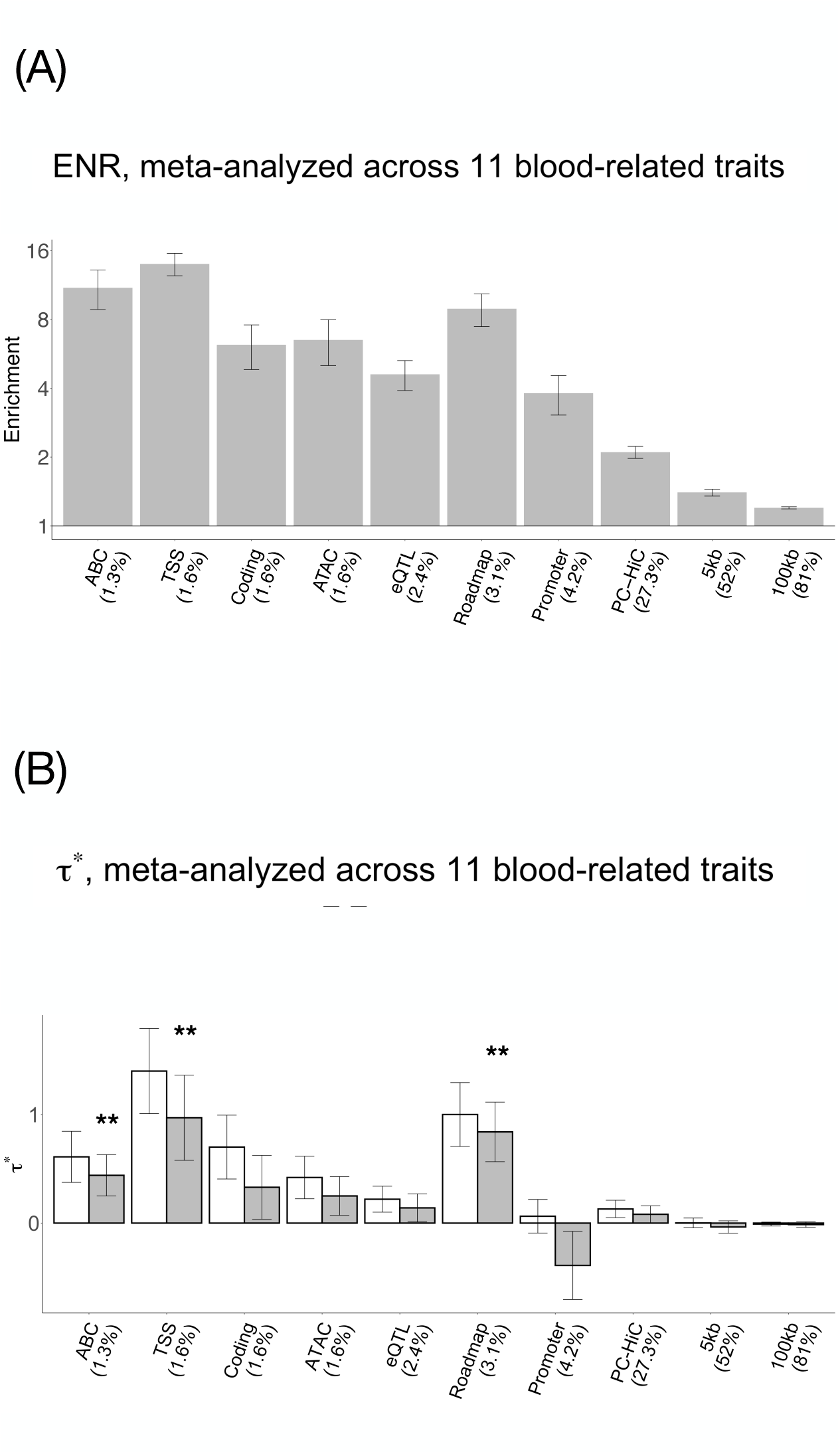
Disease informativeness of S2G annotations: We evaluated 10 S2G annotations, defined from the corresponding S2G strategies by SNPs linked to the set of all genes. (A) Heritabilty enrichment (log scale), conditional on the baseline-LD model. Horizontal line denotes no enrichment. (B) Standardized effect size (*τ^*^*), conditional on either the baseline-LD model (marginal analyses: left column, white) or the baseline-LD+ model, which includes all 10 S2G annotations (right column, dark shading). Results are meta-analyzed across 11 blood-related traits. ** denotes *P <* 0.05*/*10. Error bars denote 95% confidence intervals. Numerical results are reported in Supplementary Tables S2 and S4.

We validated the gene scores implicated in our study by investigating whether they were enriched in 5 “gold-standard” disease-related gene sets: 195 approved drug target genes for autoimmune diseases^10, 43^; 550 Mendelian genes related to immune dysregulation^44^, 390 Mendelian genes related to blood disorders^45^, 146 “Bone Marrow/Immune” genes defined by the Developmental Disorders Database/Genotype-Phenotype Database (DDD/G2P)^46^, and 2200 (top 10%) high-pLI genes^47^ (Figure 3C and Supplementary Table S6). (We note that the high-pLI genes should not be viewed as a strict gold standard as not all of these genes are disease-related, but *≈*30% of these genes have an established human disease phenotype^47^.)

**Figure 3.**
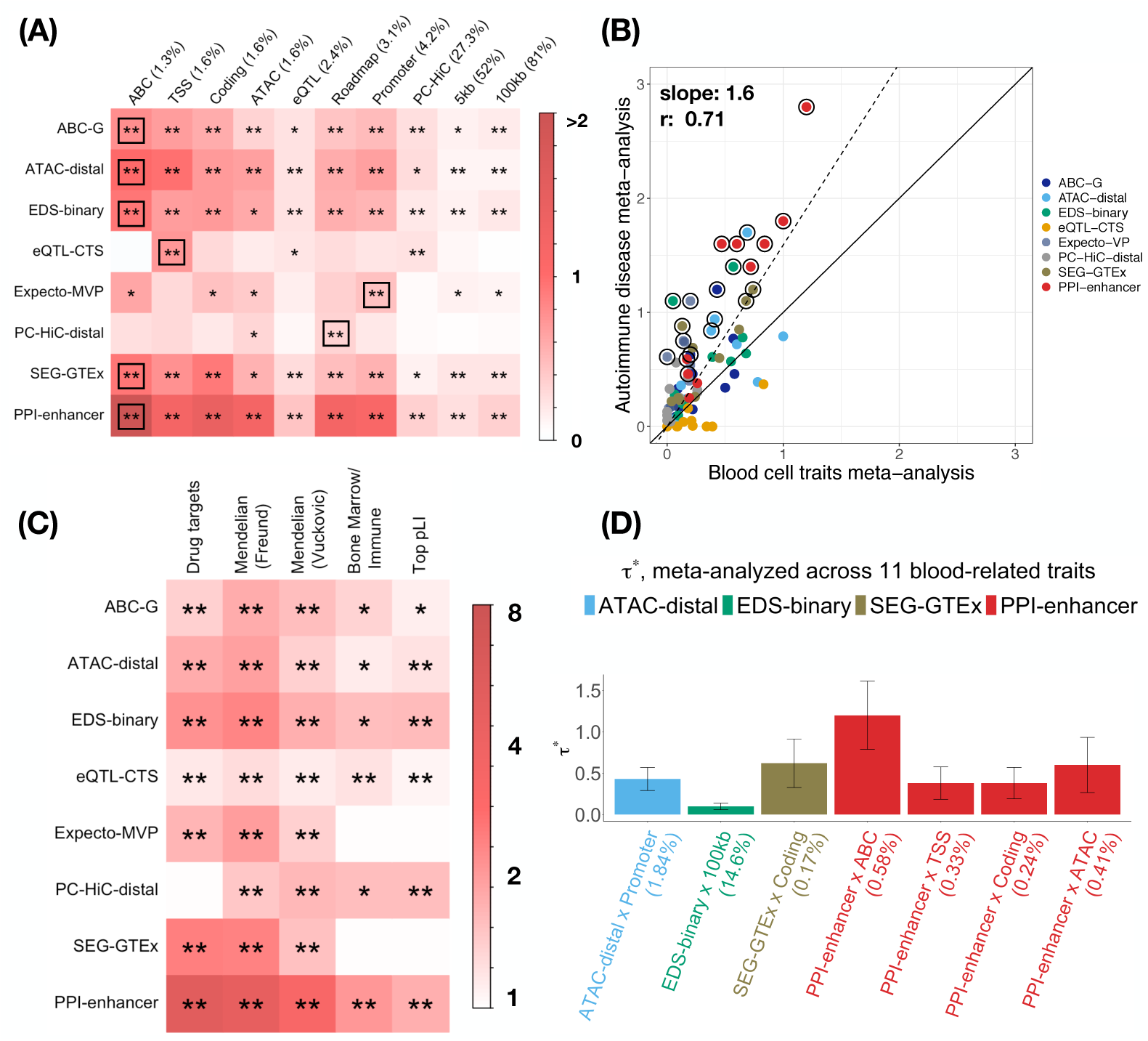
Disease informativeness of enhancer-related and PPI-enhancer annotations: We evaluated 80 annotations constructed by combining 7 enhancer-related + 1 PPI-enhancer gene scores with 10 S2G strategies. (A) Standardized effect size (*τ^*^*), conditional on the baselineLD+ model. (B) Comparison of meta-analyzed standardized effect size (*τ^*^*) across 6 autoimmune diseases vs. 5 blood cell traits. (C) Enrichment of enhancer-related and PPI-enhancer genes in 5 “gold standard” disease-related gene sets. (D) Standardized effect size (*τ^*^*), conditional on the baseline-LD+ model plus 7 jointly significant enhancer-related + PPI-enhancer annotations. In panels A and D, results are meta-analyzed across 11 blood-related traits. In panels A and C, ** denotes Bonferroni-significant (*P <* 0.05*/*110 in Panel A and *P <* 0.05*/*55 in Panel C) and * denotes FDR *<* 0.05. In panel A, the black box in each row denotes the S2G strategy with highest *τ^*^*. In Panel B, circled dots denote annotations with significant (FDR*<*5%) difference in effect size between the two meta-analyses, the solid line denotes y=x, and the dashed line denotes the regression slope. We report the slope of the regression and the Pearson correlation for enhancer-related and PPI-enhancer annotations (slope=1.3, *r*=0.57 for enhancer-related annotations only). Numerical results are reported in Supplementary Tables S8, S10, S11, S6, S23 and S26.

Subsequent subsections are organized in the following order: description of gene scores for that subsection; marginal analyses using S-LDSC; joint analyses using S-LDSC; and validation of the genes scores implicated in our study using “gold-standard” disease-related gene sets.

#### Enhancer-related genes are conditionally informative for autoimmune disease heritability

We assessed the disease informativeness of 7 gene scores prioritizing enhancer-related genes in blood. We defined these gene scores based on distal enhancer-gene connections, tissue-specific expression, or tissue-specific eQTL, all of which can characterize enhancer-related regulation (Figure 1B, Table 1 and Methods). Some of these gene scores were derived from the same functional data that we used to define S2G strategies (e.g. ABC^32, 33^ and ATAC-seq^48^; see URLs). We included two published gene scores, (binarized) blood-specific enhancer domain score (EDS)^24^ and specifically expressed genes in GTEx whole blood^9^ (SEG-GTEx). We use the term “enhancer-related” to broadly describe gene scores with high predicted functionality under a diverse set of metrics, notwithstanding the fact that all genes require the activation of enhancers and their promoters. 4 of our enhancer-related gene scores (ABC-G, ATAC-distal, EDS-binary, PC-HiC) were explicitly defined based on distal enhancer-gene connections. Using the established EDS-binary (derived from the published Enhancer Domain Score (EDS)^24^) as a point of reference, we determined that the other 3 gene scores (ABC-G, ATAC-distal, PC-HiC) had an average excess overlap of 1.7x with the EDS-binary score (P-values per gene score: 2e-08 to 6e-06; Supplementary Table S7), confirming that they prioritize enhancer-related genes. 3 of our enhancer-related scores (eQTL-CTS, Expecto-MVP, SEG-GTEx) were not explicitly defined based on distal enhancer-gene connections. We determined that these 3 gene scores also had an average excess overlap of 1.5x with the EDS-binary score (P-values per gene score: 4e-07 to 1e-04; Supplementary Table S7)) confirming that they prioritize enhancer-related genes; notably, the excess overlap of 1.5x was almost as large as the excess of overlap of 1.7x for gene scores defined based on distal enhancer-gene connections.

We combined the 7 enhancer-related gene scores with the 10 S2G strategies (Table 2) to define 70 annotations. In our marginal analysis using S-LDSC conditional on the baseline-LD+ model (meta-analyzing S-LDSC results across 11 autoimmune diseases and blood cell traits), all 70 enhancer-related annotations were significantly enriched for disease heritability, with larger enrichments for smaller annotations (Supplementary Figure S5 and Supplementary Table S8); values of standardized enrichment were more similar across annotations (Supplementary Figure S6 and Supplementary Table S9). 37 of the 70 enhancer-related annotations attained conditionally significant *τ^*^* values after Bonferroni correction (*p <* 0.05*/*110) (Figure 3A and Supplementary Table S8). We observed the strongest conditional signal for ATAC-distal *×* ABC (*τ^*^* = 1.0*±*0.2). ATAC-distal is defined by the proportion of mouse gene expression variability across blood cell types that is explained by distal ATAC-seq peaks in mouse^48^; the mouse genes are mapped to orthologous human genes. 4 of the 7 gene scores (ABC-G, ATAC-distal, EDS-binary and SEG-GTEx) produced strong conditional signals across many S2G strategies; however none of them attained Bonferroni-significant *τ^*^* for all 10 S2G strategies (Figure 3A). Among the S2G strategies, average conditional signals were strongest for the ABC strategy (average *τ^*^* = 0.59) and TSS strategy (average *τ\** = 0.52), which greatly outperformed the window-based S2G strategies (average *τ^*^* = 0.04-0.07), emphasizing the high added value of S2G strategies incorporating functional data (especially the ABC and TSS strategies).

We compared meta-analyses of S-LDSC results across 6 autoimmune diseases vs. 5 blood cell traits(Figure 3B, Supplementary Figure S7, Supplementary Table S1, Supplementary Table S10, and Supplementary Table S11). Results were broadly concordant (*r* = 0.57 between *τ^*^* estimates), with slightly stronger signals for autoimmune diseases (slope=1.3). We also compared meta-analyses of results across 2 granulocyte-related blood cell traits (white blood cell count, eosinophil count) vs. 3 red blood cell or platelet-related blood cell traits (red blood cell count, red blood cell distribution width, platelet count) (Supplementary Figure S8, Supplementary Table S12, and Supplementary Table S13). Results were broadly concordant (*r* = 0.65, slope = 1.1). We also examined S-LDSC results for individual disease/traits and applied a test for heterogeneity^49^ (Supplementary Figure S9, Supplementary Figure S10, Supplementary Table S14, Supplementary Table S15). Results were generally underpowered (FDR*<*5% for 16 of 770 annotation-trait pairs), with limited evidence of heterogeneity across diseases/traits (FDR*<*5% for 11 of 70 annotations).

We jointly analyzed the 37 enhancer-related annotations that were Bonferroni-significant in our marginal analysis (Figure 3A and Supplementary Table S8) by performing forward stepwise elimination to iteratively remove annotations that had conditionally non-significant *τ^*^* values after Bonferroni correction. Of these, 6 annotations were jointly significant in the resulting enhancer-related joint model (Supplementary Figure S11 and Supplementary Table S16), corresponding to 4 enhancer-related gene scores: ABC-G, ATAC-distal, EDS-binary and SEG-GTEx.

We assessed the enrichment of the 7 enhancer-related gene scores (Table 1) in 5 “gold-standard” disease-related gene sets (drug target genes^10, 43^, Mendelian genes (Freund)^44^, Mendelian genes (Vuckovic)^45^, immune genes^46^, and high-pLI genes^47^) (Figure 3C and Supplementary Table S6). 6 of the 7 gene scores were significantly enriched (after Bonferroni correction; *p <* 0.05*/*55) in the drug target genes, all 7 were significantly enriched in both Mendelian gene sets, 3 of 7 were significantly enriched in the immune genes, and 5 of 7 were significantly enriched in the high-pLI genes. The largest enrichment was observed for SEG-GTEx genes in the drug target genes (2.4x, s.e. 0.1) and Mendelian genes (Freund) (2.4x, s.e. 0.1). These findings validate the high importance to disease of enhancer-related genes..

We performed 5 secondary analyses. First, for each of the 6 annotations from the enhancer-related joint model (Supplementary Figure S11), we assessed their functional enrichment for fine-mapped SNPs for blood-related traits from two previous studies^50, 51^. We observed large and significant enrichments for all 6 annotations (Supplementary Table S17), consistent with the S-LDSC results. Second, for each of the 7 enhancer-related gene scores, we performed pathway enrichment analyses to assess their enrichment in pathways from the ConsensusPathDB database^52^; all 7 gene scores were significantly enriched in immune-related and signaling pathways (Supplementary Table S18). Third, we explored other approaches to combining information across genes that are linked to a SNP using S2G strategies, by using either the mean across genes or the sum across genes of the gene scores linked to a SNP, instead of the maximum across genes. We determined that results for either the mean or the sum were very similar to the results for the maximum, with no significant difference in standardized effect sizes of the resulting SNP annotations (Supplementary Table S8, Supplementary Table S19, Supplementary Table S20). Fourth, we repeated our analyses of the 5 enhancer-related gene scores for which the top 10% (of genes) threshold was applied, using top 5% or top 20% thresholds instead (Supplementary Table S21 and Supplementary Table S22). We observed very similar results, with largely non-significant differences in standardized effect sizes. Fifth, we confirmed that our forward stepwise elimination procedure produced identical results when applied to all 70 enhancer-related annotations, instead of just the 37 enhancer-related annotations that were Bonferroni-significant in our marginal analysis.

We conclude that 4 of the 7 characterizations of enhancer-related genes are conditionally informative for autoimmune diseases and blood-related traits when using functionally informed S2G strategies.

#### Genes with high network connectivity to enhancer-related genes are even more informative

We assessed the disease informativeness of a gene score prioritizing genes with high connectivity to enhancer-related genes in a protein-protein interaction (PPI) network (PPI-enhancer).We hypothesized that (i) genes that are connected to enhancer-related genes in biological networks are likely to be important, and that (ii) combining potentially noisy metrics defining enhancer-related genes would increase statistical signal. We used the STRING PPI network^53^ to quantify the network connectivity of each gene with respect to each of the 4 jointly informative enhancer-related gene scores from Supplementary Figure S11 (ABC-G, ATAC-distal, EDS-binary and SEG-GTEx) (Figure 1D). Network connectivity scores were computed using a random walk with restart algorithm^10, 54^ (see Methods). We defined the PPI-enhancer gene score based on genes in the top 10% of average network connectivity across the 4 enhancer-related gene scores (Table 1). The PPI-enhancer gene score was only moderately positively correlated with the 4 underlying enhancer-related gene scores (average *r*=0.28; Supplementary Figure S2).

We combined the PPI-enhancer gene score with the 10 S2G strategies (Table 2) to define 10 annotations. In our marginal analysis using S-LDSC (meta-analyzing S-LDSC results across 11 autoimmune diseases and blood cell traits), all 10 PPI-enhancer annotations were significantly enriched for disease heritability, with larger enrichments for smaller annotations (Supplementary Figure S5 and Supplementary Table S23); values of standardized enrichment were more similar across annotations (Supplementary Figure S6 and Supplementary Table S24). All 10 PPI-enhancer annotations attained conditionally significant *τ^*^* values after Bonferroni correction (*p <* 0.05*/*110) (Figure 3A and Supplementary Table S23). Notably, the maximum *τ^*^* (2.0 (s.e. 0.3) for PPI-enhancer x ABC) was *>*2x larger than the maximum *τ^*^* for the recently proposed Enhancer Domain Score^24^ (EDS) (0.91 (s.e. 0.21) for EDS-binary x ABC). All 10 PPI-enhancer annotations remained significant when conditioned on the enhancer-related joint model from Supplementary Figure S11 (Supplementary Table S25). In a comparison of meta-analyses of S-LDSC results across 5 blood cell traits vs. 6 autoimmune diseases, results were broadly concordant (*r* = 0.93 between *τ^*^* estimates), but with much stronger signals for autoimmune diseases (slope=2.2) (Figure 3B, Supplementary Figure S7, Supplementary Table S10, and Supplementary Table S11). In a comparison of meta-analyses across 2 granulocyte-related blood cell traits vs. 3 red blood cell or platelet-related blood cell traits, results were broadly concordant (*r* = 0.83), but with much stronger signals for granulocyte-related blood cell traits (slope = 2.1); providing a further validation that the PPI-enhancer gene score is related to immune response (Supplementary Figure S8 and Supplementary Tables S12, S13). In analyses of individual traits, 62 of 110 PPI-enhancer annotation-trait pairs were significant (FDR*<*5%) (Supplementary Figure S9, Supplementary Figure S10, Supplementary Table S14), 8 of them with evidence of heterogeneity across diseases/traits (FDR*<*5% for 8 of 10 PPI-enhancer annotations) (Supplementary Table S15).

We jointly analyzed the 6 enhancer-related annotations from the enhancer-related joint model (Supplementary Figure S11) and the 10 marginally significant PPI-enhancer annotations conditional on the enhancer-related joint model in Supplementary Table S25. Of these, 3 enhancer-related and 4 PPI-enhancer annotations were jointly significant in the resulting PPI-enhancer-related joint model (Figure 3D and Supplementary Table S26). The joint signal was strongest for PPI-enhancer *×* ABC (*τ^*^* = 1.2*±*0.21), highlighting the informativeness of the ABC S2G strategy. 3 of the 7 annotations attained *τ^*^ >* 0.5; annotations with *τ^*^ >* 0.5 are unusual, and considered to be important^39^.

We assessed the enrichment of the PPI-enhancer gene score in the 5 “gold-standard” disease-related gene sets (drug target genes^10, 43^, Mendelian genes (Freund)^44^, Mendelian genes (Vuckovic)^45^, immune genes^46^, and high-pLI genes^47^) (Figure 3C and Supplementary Table S6). The PPI-enhancer gene score showed significant enrichment in all 5 gene sets, with higher magnitude of enrichment compared to any of the 7 enhancer-related gene scores. In particular, the PPI-enhancer gene score was 5.3x (s.e. 0.1) enriched in drug target genes and 4.6x (s.e. 0.1) enriched in Mendelian genes (Freund), a *≥*2x stronger enrichment in each case than the EDS-binary gene score^24^ (2.1x (s.e. 0.1) and 2.3x (s.e. 0.1)).

We sought to assess whether the PPI-enhancer disease signal derives from (i) the information in the PPI network or (ii) the improved signal-to-noise of combining different enhancer-related gene scores (see above). To assess this, we constructed an optimally weighted linear combination of the 4 enhancer-related scores from Figure 3D, without using PPI network information (Weighted-enhancer; see Methods). We repeated the above analyses using Weighted-enhancer instead of PPI-enhancer. We determined that marginal *τ^*^* values were considerably lower for Weighted-enhancer vs. PPI-enhancer annotations (0.65x, Bootstrap p: 3.4e-09 for 1,000 re-samples per annotation, Supplementary Table S27 vs. Supplementary Table S23). (In addition, none of the Weighted-enhancer annotations were significant conditional on the PPI-enhancer-related joint model from Figure 3D; see Supplementary Table S28). This confirms that the additional PPI-enhancer signal derives from the information in the PPI network. To verify that the PPI-enhancer disease signal is driven not just by the PPI network but also by the input gene scores, we defined a new gene score analogous to PPI-enhancer but using 4 randomly generated binary gene sets of size 10% as input (PPI-control). We determined that marginal *τ^*^* values were much lower for PPI-control vs. PPI-enhancer annotations (0.52x, Bootstrap p-value: 4.4e-16 over 1,000 re-samples per annotation, Supplementary Table S29 vs. Supplementary Table S23). (In addition, none of the PPI-control annotations were significant conditional on the PPI-enhancer-related joint model from Figure 3D; see Supplementary Table S30). This confirms that the PPI-enhancer disease signal is driven not just by the PPI network but also by the input gene scores.

We performed 4 secondary analyses. First, we defined a new gene score (RegNet-Enhancer) using the regulatory network from ref.^55^ instead of the STRING PPI network, and repeated the above analyses. We determined that the STRING PPI network and the RegNet regulatory network are similarly informative (Supplementary Table S31 and Supplementary Table S32); we elected to use the STRING PPI network in our main analyses because the RegNet regulatory network uses GTEx expression data, which is also used by the SEG-GTEx gene score, complicating interpretation of the results. Second, for each of the 4 jointly significant PPI-enhancer annotations from Figure 3D, we assessed their functional enrichment for fine-mapped SNPs for blood-related traits from two previous studies^50, 51^. We observed large and significant enrichments for all 4 annotations (Supplementary Table S17), consistent with the S-LDSC results (and with the similar analysis of enhancer-related annotations described above). Third, we performed a pathway enrichment analysis to assess the enrichment of the PPI-enhancer gene score in pathways from the ConsensusPathDB database^52^; this gene score was enriched in immune-related pathways (Supplementary Table S18). Fourth, we confirmed that our forward stepwise elimination procedure produced identical results when applied to all 80 enhancer-related and PPI-enhancer annotations, instead of just the 6 enhancer-related annotations from the enhancer-related joint model (Supplementary Figure S11) and the 10 PPI-enhancer annotations.

We conclude that genes with high network connectivity to enhancer-related genes are conditionally informative for autoimmune diseases and blood-related traits when using functionally informed S2G strategies.

#### Candidate master-regulator genes are conditionally informative for autoimmune disease heritability

We assessed the disease informativeness of two gene scores prioritizing candidate master-regulator genes in blood. We defined these gene scores using whole blood eQTL data from the eQTLGen consortium^56^ (Trans-master) and a published list of known transcription factors in humans^57^ (TF) (Figure 1C, Table 1 and Methods). We note that TF genes do not necessarily act as master regulators and only a small number of TFs regulate many downstream genes, but TF genes can still be viewed as candidate master regulators. Using 97 known master-regulator genes from 18 master-regulator families^58–62^ as a point of reference, we determined that Trans-master and TF genes had 3.5x and 5.4x excess overlaps with the 97 candidate master-regulator genes (P-values: 5.6e-72 and 2.2e-160; Supplementary Table S33 and Supplementary Table S34), confirming that they prioritize candidate master-regulator genes.

In detail, Trans-master is a binary gene score defined by genes that significantly regulate 3 or more other genes in trans via SNPs that are significant cis-eQTLs of the focal gene (10% of genes); the median value of the number of genes trans-regulated by a Trans-master gene is 14. Notably, trans-eQTL data from the eQTLGen consortium^56^ was only available for 10,317 previously disease-associated SNPs. It is possible that genes with significant cis-eQTL that are disease-associated SNPs may be enriched for disease heritability irrespective of trans signals. To account for this gene-level bias, we conditioned all analyses of Trans-master annotations on both (i) 10 annotations based on a gene score defined by genes with at least 1 disease-associated cis-eQTL, combined with each of the 10 S2G strategies, and (ii) 10 annotations based on a gene score defined by genes with at least 3 unlinked disease-associated cis-eQTL, combined with each of the 10 S2G strategies; we chose the number 3 to maximize the correlation between this gene score and the Trans-master gene score (*r* = 0.32). Thus, our primary analyses were conditioned on 93 baseline-LD+ and 20 additional annotations (113 baseline-LD+cis model annotations); additional secondary analyses are described below. We did not consider a SNP annotation defined by trans-eQTLs, because the trans-eQTLs in eQTLGen data were restricted to disease-associated SNPs, which would bias our results.

We combined the Trans-master gene score with the 10 S2G strategies (Table 2) to define 10 annotations. In our marginal analysis using S-LDSC conditional on the baseline-LD+cis model, all 10 Trans-master annotations were strongly and significantly enriched for disease heritability, with larger enrichments for smaller annotations (Supplementary Figure S5 and Supplementary Table S35); values of standardized enrichment were more similar across annotations (Supplementary Figure S6 and Supplementary Table S36). All 10 Trans-master annotations attained conditionally significant *τ^*^* values after Bonferroni correction (*p <* 0.05*/*110) (Figure 4A and Supplementary Table S35). We observed the strongest conditional signals for Trans-master *×* TSS (*τ^*^* = 1.6, vs. *τ^*^* = 0.37-0.39 for candidate master-regulator *×* window-based S2G strategies). We observed similar (slightly more significant) results when conditioning on baseline-LD+ annotations only (Supplementary Table S37).

**Figure 4.**
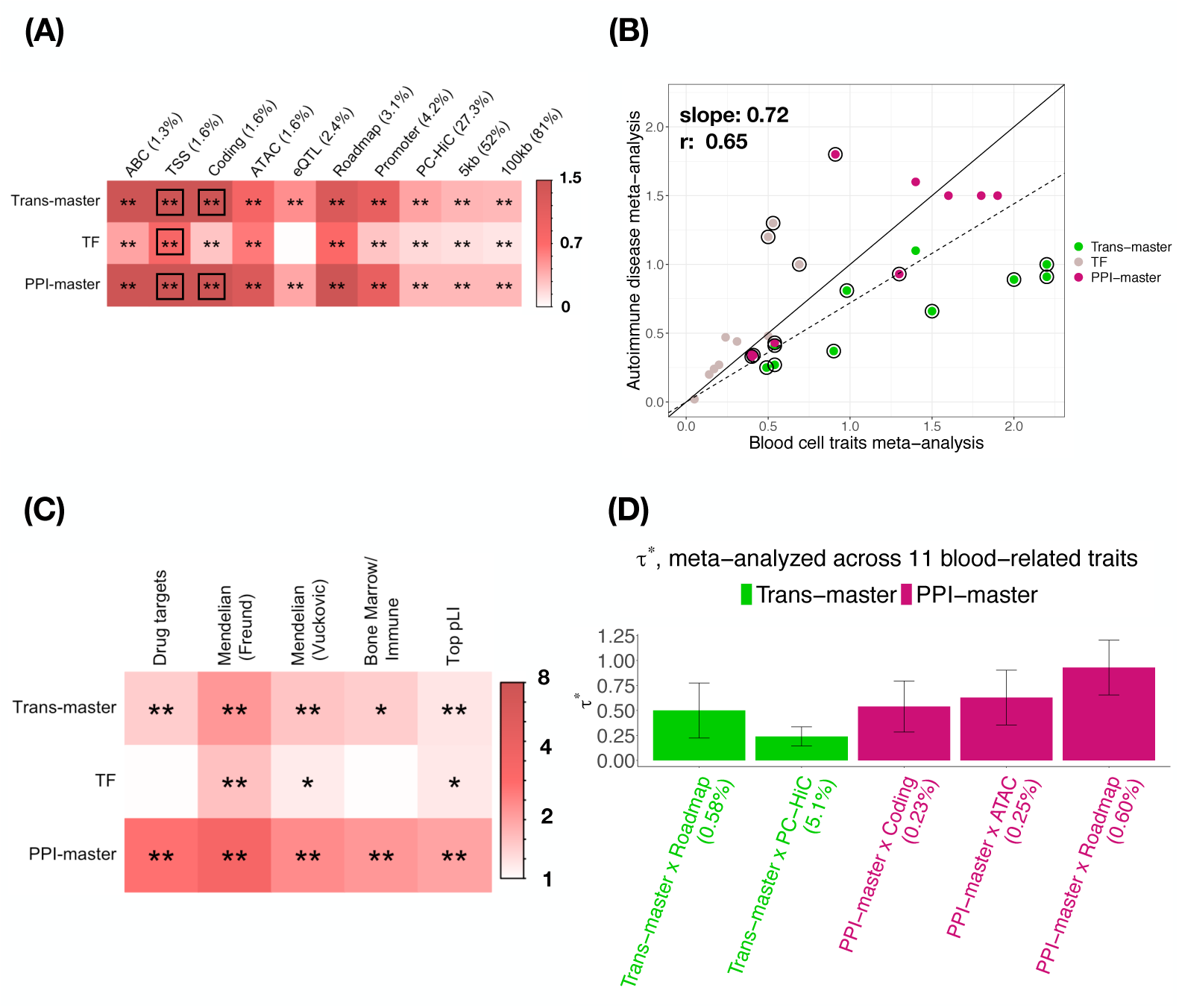
Disease informativeness of Master-regulator and PPI-master annotations: We evaluated 30 annotations constructed by combining 2 Master-regulator + 1 PPI-master gene scores with 10 S2G strategies. (A) Standardized effect size (*τ^*^*), conditional on the 113 baselineLD+cis model annotations. (B) Comparison of meta-analyzed standardized effect size (*τ^*^*) across 6 autoimmune diseases vs. 5 blood cell traits. (C) Enrichment of Master-regulator and PPI-master genes in 5 “gold standard” disease-related gene sets.(D) Standardized effect size (*τ^*^*), conditional on the baseline-LD+cis model plus 5 jointly significant Master-regulator + PPI-master annotations. In panels A and D, results are meta-analyzed across 11 blood-related traits. In panels A and C, ** denotes Bonferroni-significant (*P <* 0.05*/*110 in Panel A and *P <* 0.05*/*55 in Panel C) and * denotes FDR *<* 0.05. In panel A, the black box in each row denotes the S2G strategy with highest *τ^*^*. In Panel B, circled dots denote annotations with significant (FDR*<*5%) difference in effect size between the two meta-analyses, the solid line denotes y=x, and the dashed line denotes the regression slope. We report the slope of the regression and the Pearson correlation for master-regulator and PPI-master annotations (slope=0.57, *r*=0.56 for master-regulator annotations only). Numerical results are reported in Supplementary Tables S35, Table S44, S45, S6, S53 and S57.

As noted above, trans-eQTL data from the eQTLGen consortium^56^ was only available for 10,317 previously disease-associated SNPs, and we thus defined and conditioned on baseline-LD+cis model annotations to account for gene-level bias. We verified that conditioning on annotations derived from gene scores defined by other minimum numbers of cis-eQTL and/or unlinked cis-eQTL produced similar results (Supplementary Table S38, Supplementary Table S39, Supplementary Table S40, Supplementary Table S41, Supplementary Table S42). To verify that our results were not impacted by SNP-level bias, we adjusted each of the 10 Trans-master annotations by removing all disease-associated trans-eQTL SNPs in the eQTLGen data from the annotation, as well as any linked SNPs (Methods). We verified that these adjusted annotations produced similar results (Supplementary Table S43).

TF is a binary gene score defined by a published list of 1,639 known transcription factors in humans^57^. We combined TF with the 10 S2G strategies (Table 2) to define 10 annotations. In our marginal analysis conditional on the baseline-LD+cis model, all 10 TF annotations were significantly enriched for heritability, but with smaller enrichments than the Trans-master annotations (Supplementary Table S35); see Supplementary Table S36 for standardized enrichments. 9 TF annotations attained significant *τ^*^* values after Bonferroni correction (Figure 4A and Supplementary Table S35) (the same 9 annotations were also significant conditional on the baseline-LD+ model; Supplementary Table S37). Across all S2G strategies, *τ^*^* values of Trans-master annotations were larger than those of TF annotations (Supplementary Table S35).

We compared meta-analyses of S-LDSC results across 6 autoimmune diseases vs. 5 blood cell traits(Figure 4B, Supplementary Figure S7, Supplementary Table S1, Supplementary Table S44, Supplementary Table S45). Results were broadly concordant (*r* = 0.56 between *τ^*^* estimates), with slightly stronger signals for blood cell traits (slope=0.57). We also compared meta-analyses of results across 2 granulocyte-related blood cell traits vs. 3 red blood cell or platelet-related blood cell traits (Supplementary Figure S8, Supplementary Table S46, and Supplementary Table S47). Results were broadly concordant (*r* = 0.94, slope = 1.12). We also examined S-LDSC results for individual disease/traits and applied a test for heterogeneity^49^ (Supplementary Figure S12, Supplementary Figure S13, Supplementary Table S14, Supplementary Table S15). We observed several annotation-trait pairs with disease signal (FDR*<*5% for 96 of 220 annotation-trait pairs), with evidence of heterogeneity across diseases/traits (FDR*<*5% for 10 of 20 annotations).

We jointly analyzed the 10 Trans-master and 9 TF annotations that were Bonferroni-significant in our marginal analysis (Figure 4A and Supplementary Table S35) by performing forward stepwise elimination to iteratively remove annotations that had conditionally non-significant *τ^*^* values after Bonferroni correction. Of these, 3 Trans-master annotations and 2 TF annotations were jointly significant in the resulting candidate master-regulator joint model (Supplementary Figure S14 and Supplementary Table S48). The joint signal was strongest for Trans-master *×* Roadmap (*τ^*^* = 0.81, s.e. = 0.13), emphasizing the high added value of the Roadmap S2G strategy.

We assessed the enrichment of the Trans-master and TF gene scores in the 5 “gold standard” disease-related gene sets (drug target genes^10, 43^, Mendelian genes (Freund)^44^, Mendelian genes (Vuckovic)^45^, immune genes^46^, and high-pLI genes^47^) (Figure 4C and Supplementary Table S6). The Trans-master gene score showed higher enrichment in all 5 gene sets compared to the TF gene score. The enrichments for candidate master-regulator genes were lower (1.4x, s.e. 0.07) for drug target genes in comparison to some enhancer-related genes and the PPI-enhancer gene score (Figure 3C); this can be attributed to the fact that candidate master-regulator genes may tend to disrupt genes across several pathways, rendering them unsuitable as drug targets.

We performed 7 secondary analyses. First, for comparison purposes, we defined a binary gene score (Trans-regulated) based on genes with at least one significant trans-eQTL. We combined Trans-regulated genes with the 10 S2G strategies to define 10 annotations. In our marginal analysis using S-LDSC conditional on the baseline-LD+cis model, none of the Trans-regulated annotations attained conditionally significant *τ^*^* values after Bonferroni correction (*p <* 0.05*/*110) (Supplementary Table S49). (In contrast, 3 of the annotations were significant when conditioning only on the baseline-LD+ model (Supplementary Table S50).) Second, a potential complexity is that trans-eQTL in whole blood may be inherently enriched for blood cell trait-associated SNPs (since SNPs that regulate the abundance of a specific blood cell type would result in trans-eQTL effects on genes that are specifically expressed in that cell type^56^), potentially limiting the generalizability of our results to non-blood cell traits. To ensure that our results were robust to this complexity, we verified that analyses restricted to the 5 autoimmune diseases (Supplementary Table S1) produced similar results (Supplementary Table S51). Third, for each of the 5 annotations from the candidate master-regulator joint model (Supplementary Figure S14), we assessed their functional enrichment for fine-mapped SNPs for blood-related traits from two previous studies^50, 51^. We observed large and significant enrichments for all 5 annotations (Supplementary Table S17), consistent with the S-LDSC results (and with similar analyses described above). Fourth, we performed pathway enrichment analyses to assess the enrichment of the Trans-master and TF gene scores in pathways from the ConsensusPathDB database^52^. The Trans-master gene score was significantly enriched in immune-related pathways (Supplementary Table S18). Fifth, we explored other approaches to combining information across genes that are linked to a SNP using S2G strategies, by using either the mean across genes or the sum across genes of the gene scores linked to a SNP, instead of the maximum across genes. We determined that results for either the mean or the sum were very similar to the results for the maximum, with no significant difference in standardized effect sizes of the resulting SNP annotations (Supplementary Table S35, Supplementary Table S19 and Supplementary Table S20). Sixth, we repeated our analyses of the Trans-master gene score, defined in our primary analyses based on 2,215 genes that trans-regulate *≥* 3 genes, using either 3,717 genes that trans-regulate *≥* 1 gene (most of which trans-regulate multiple genes) or 1,170 genes that trans-regulate *≥* 10 genes (Supplementary Table S52). We observed very similar results, with largely non-significant differences in standardized effect sizes. Seventh, we confirmed that our forward stepwise elimination procedure produced identical results when applied to all 20 candidate master-regulator annotations, instead of just the 19 candidate master-regulator annotations that were Bonferroni-significant in our marginal analysis.

We conclude that candidate master-regulator genes are conditionally informative for autoimmune diseases and blood-related traits when using functionally informed S2G strategies.

#### Genes with high network connectivity to candidate master-regulator genes are even more informative

We assessed the disease informativeness of a gene score prioritizing genes with high connectivity to candidate master-regulator genes in the STRING PPI network^53^ (PPI-master, analogous to PPI-enhancer; see Methods and Table 1). The PPI-master gene score was positively correlated with the 2 underlying candidate master-regulator gene scores (average *r* = 0.43) and modestly correlated with PPI-enhancer (r=0.22) (Supplementary Figure S2). In addition, it had an excess overlap of 7.2x with the 97 known Master regulator genes^58–62^ (P = 2e-214; Supplementary Table S33 and Supplementary Table S34).

We combined the PPI-master gene score with the 10 S2G strategies (Table 2) to define 10 annotations. In our marginal analysis using S-LDSC conditional on the baseline-LD+cis model, all 10 PPI-master annotations were significantly enriched for disease heritability, with larger enrichments for smaller annotations (Figure 4A and Supplementary Table S53); values of standardized enrichment were more similar across annotations (Supplementary Figure S6 and Supplementary Table S54). All 10 PPI-master annotations attained conditionally significant *τ^∗^* values after Bonferroni correction (*p <* 0.05*/*110) (Figure 4B and Supplementary Table S53) (as expected, results were similar when conditioning only on the baseline-LD+ model; Supplementary Table S55). We observed the strongest conditional signals for PPI-master combined with TSS (*τ\**=1.7, s.e. 0.16), Coding (*τ\**=1.7, s.e. 0.14) and ABC (*τ\**=1.6, s.e. 0.17) S2G strategies, again emphasizing the high added value of S2G strategies incorporating functional data (Supplementary Table S53). 9 of the 10 PPI-master annotations remained significant when conditioning on the candidate master-regulator joint model from Supplementary Figure S14 (Supplementary Table S56). In a comparison of meta-analyses of S-LDSC results across 5 blood cell traits vs. 6 autoimmune diseases, results were broadly concordant (*r* = 0.81 between *τ^*^* estimates, slope = 0.93) (Figure 4B, Supplementary Figure S7, Supplementary Table S44, and Supplementary Table S45). In a comparison of meta-analyses across 2 granulocyte-related blood cell traits vs. 3 red blood cell or platelet-related blood cell traits, results were broadly concordant, but with slightly stronger signals for granulocyte-related traits (*r* = 0.92, slope = 1.3), providing a further validation that the PPI-master gene score is related to immune response (Supplementary Figure S8 and Supplementary Tables S46, S47). In the analyses of individual traits, 101 of 110 PPI-enhancer annotation-trait pairs were significant (FDR*<*5%) (Supplementary Figure S12, Supplementary Figure S13, Supplementary Table S14), with evidence of heterogeneity across diseases/traits (FDR*<*5% for 6 of 10 PPI-master annotations)(Supplementary Table S15).

We jointly analyzed the 5 candidate master-regulator annotations from the candidate master-regulator joint model (Supplementary Figure S14 and Supplementary Table S48) and the 9 PPI-master annotations significant conditional on the candidate master-regulator joint model in Supplementary Table S56. Of these, 2 Trans-master and 3 PPI-master annotations were jointly significant in the resulting PPI-master-regulator joint model (Figure 4D and Supplementary Table S57). The joint signal was strongest for PPI-master *×* Roadmap (*τ^*^* = 0.94*±*0.14),and 4 of the 5 annotations attained *τ^*^ >* 0.5.

We assessed the enrichment of the PPI-master gene score in the 5 “gold standard” disease-related gene sets (drug target genes^10, 43^, Mendelian genes (Freund)^44^, Mendelian genes (Vuckovic)^45^, immune genes^46^, and high-pLI genes^47^) (Figure 4C and Supplementary Table S6). The PPI-master gene score showed significant enrichment in all 5 gene sets, with higher magnitude of enrichment compared to either of the candidate master-regulator gene scores. In particular, the PPI-master gene score was 2.7x (s.e. 0.1) enriched in drug target genes and 3.4x (s.e. 0.1) enriched in Mendelian genes (Freund).

We performed 3 secondary analyses. First, for each of the 3 jointly significant PPI-master annotations from Figure 4D, we assessed their functional enrichment for fine-mapped SNPs for blood-related traits from two previous studies^50, 51^. We observed large and significant enrichments for all 3 annotations (Supplementary Table S17), consistent with the S-LDSC results (and with similar analyses described above). Second, we performed a pathway enrichment analysis to assess the enrichment of the PPI-master gene score in pathways from the ConsensusPathDB database^52^ and report the top enriched pathways (Supplementary Table S18). Third, we confirmed that our forward stepwise elimination procedure produced identical results when applied to all 30 candidate master-regulatorand PPI-master annotations, instead of just the 5 candidate master-regulator annotations from the candidate master-regulator joint model (Supplementary Figure S14) and the 9 PPI-master annotations that were Bonferroni-significant in our marginal analysis.

We conclude that genes with high network connectivity to candidate master-regulator genes are conditionally informative for autoimmune diseases and blood-related traits when using functionally informed S2G strategies.

#### Combined joint model

We constructed a combined joint model containing annotations from the above analyses that were jointly significant, contributing information conditional on all other annotations. We merged the baseline-LD+cis model with annotations from the PPI-enhancer (Figure 3D) and PPI-master (Figure 4D) joint models, and performed forward stepwise elimination to iteratively remove annotations that had conditionally non-significant *τ^*^* values after Bonferroni correction (*p <* 0.05*/*110). The combined joint model contained 8 new annotations, including 2 enhancer-related, 2 PPI-enhancer, 2 Trans-master and 2 PPI-master annotations (Figure 5 and Supplementary Table S58). The joint signals were strongest for PPI-enhancer *×* ABC (*τ^*^* = 0.99, s.e. 0.23) and PPI-master*×*Roadmap (*τ^*^* = 0.91, s.e. 0.12) highlighting the importance of two distal S2G strategies, ABC and Roadmap; 5 of the 8 new annotations attained *τ^*^ >* 0.5. We defined a new metric quantifying the conditional informativeness of a heritability model (combined *τ\**, generalizing the combined *τ^*^* metric of ref.^63^ to more than two annotations; see Methods). As expected, the combined joint model attained a larger combined *τ^*^* (2.5, s.e. 0.24) than the PPI-enhancer (1.5, s.e. 0.15) or PPI-master (1.9, s.e. 0.14) joint models (Supplementary Figure S15, Supplementary Table S59, Supplementary Table S60, Supplementary Table S61).

**Figure 5.**
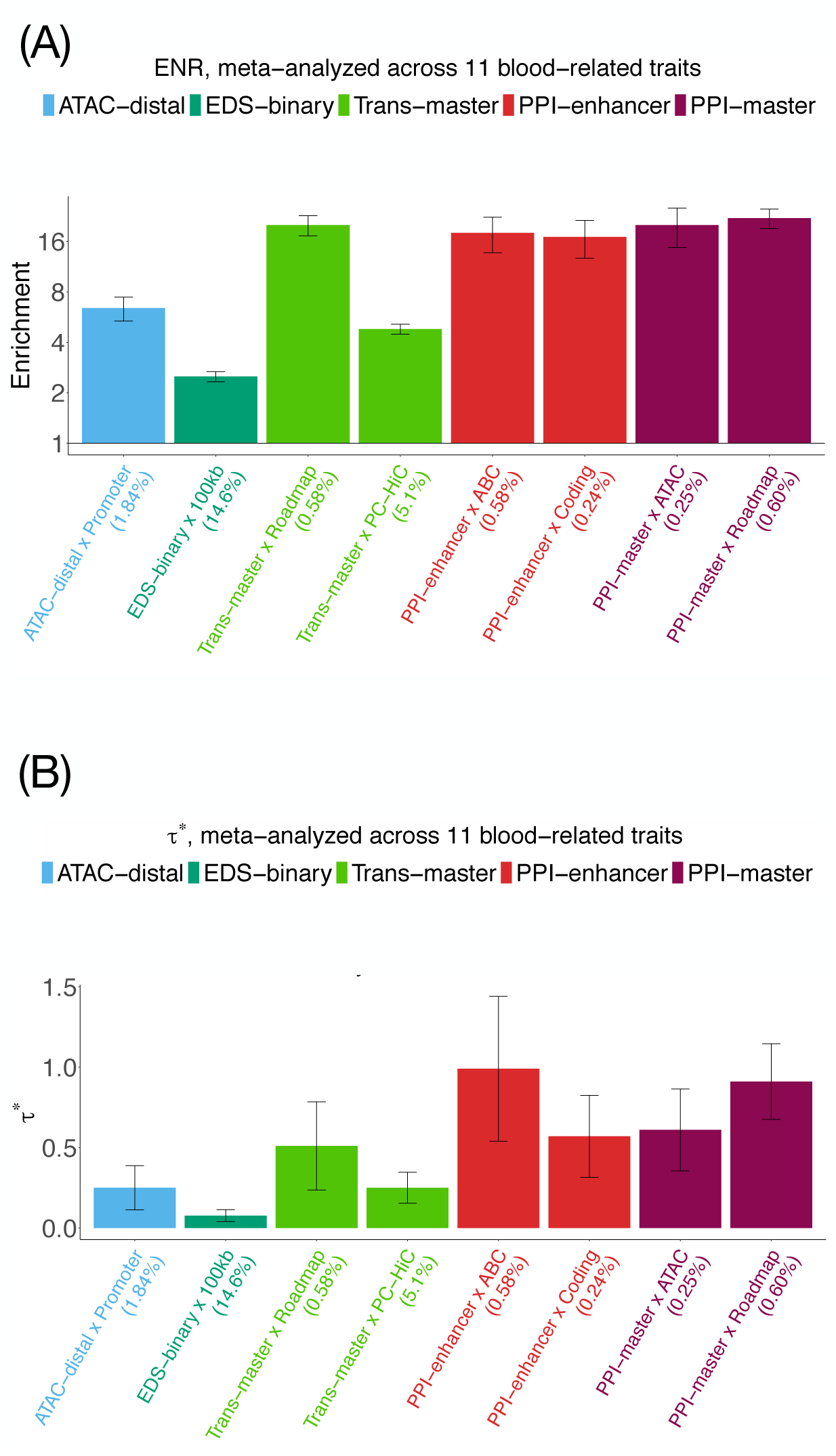
Combined joint model: (A) Heritability enrichment (log scale) of the 8 jointly significant enhancer-related, Master-regulator, PPI-enhancer-related and PPI-master-regulator annotations, conditional on the baseline-LD+cis model. Horizontal line denotes no enrichment. (B) Standardized effect size (*τ^*^*) conditional on the baseline-LD+cis model plus the 8 jointly significant annotations. Significance is corrected for multiple testing by Bonferroni correction (*P <* 0.05*/*110). Errors bars denote 95% confidence intervals. Numerical results are reported in Supplementary Table S58.

We evaluated the combined joint model of Figure 5 (and other models) by computing log*l*_SS_^64^ (an approximate likelihood metric) relative to a model with no functional annotations (Δlog*l*_SS_), averaged across a subset of 6 blood-related traits (1 autoimmune disease and 5 blood cell traits) from the UK Biobank (Supplementary Table S1). The combined joint model attained a +12.3% larger Δlog*l*_SS_ than the baseline-LD model (Supplementary Table S62); most of the improvement derived from the 7 S2G annotations (Figure 2) and the 8 enhancer-related and candidate master-regulator annotations (Figure 5). The combined joint model also attained a 27.2% larger Δlog*l*_SS_ than the baseline-LD model in a separate analysis of 24 non-blood-related traits from the UK Biobank (Supplementary Table S62; traits listed in Supplementary Table S63), implying that the value of the annotations introduced in this paper is not restricted to autoimmune diseases and blood-related traits. However, the non-blood-related traits had considerably lower absolute Δlog*l*_SS_ compared to the blood-related traits. Accordingly, in a broader analysis of 36 non-blood-related traits from UK Biobank and non-UK Biobank sources (Supplementary Table S64), meta-analyzed *τ^∗^* values were considerably lower for non-blood-related traits than for blood-related traits (Supplementary Figure S16, Supplementary Table S65 and Supplementary Table S66).

We investigated the biology of individual loci by examining 1,198 SNPs that were previously confidently fine-mapped (posterior inclusion probability (PIP) *>* 0.90) for 1 autoimmune disease and 5 blood cell traits from the UK Biobank. As noted above, fine-mapped SNPs from ref.^51^ were highly enriched for all 8 annotations from the combined joint model (Supplementary Table S17); accordingly, focusing on the 4 highly enriched regulatory annotations from Figure 5 (enrichment*≥*18; 1.5% of SNPs in total), 194 of the 1,198 SNPs belonged to one or more of these 4 annotations. A list of these 194 SNPs is provided in Supplementary Table S67. We highlight 3 notable examples. First, rs231779, a fine-mapped SNP (PIP=0.91) for “All Auto Immune Traits” (Supplementary Table S1), was linked by the ABC S2G strategy to *CTLA4*, a high-scoring gene for the PPI-enhancer gene score (ranked 109) (Supplementary Figure S17A). *CTLA4* acts as an immune checkpoint for activation of T cells and is a key target gene for cancer immunotherapy^65–67^. Second, rs6908626, a fine-mapped SNP (PIP=0.99) for ”All Auto Immune Traits” (Supplementary Table S1), was linked by the Roadmap S2G strategy to *BACH2*, a high-scoring gene for the Trans-master gene score (ranked 311) (Supplementary Figure S17B). *BACH2* is a known master-regulator TF that functions in innate and adaptive lineages to control immune responses^68, 69^, has been shown to control autoimmunity in mice knockout studies^70^, and has been implicated in several autoimmune and allergic diseases including lupus, type 1 diabetes and asthma^71–73^. Third, rs113473633, a fine-mapped SNP (PIP = 0.99 and PIP = 0.99) for white blood cell (WBC) count and eosinophil count (Supplementary Table S1), was linked by the Roadmap S2G strategy to *NFKB1*, a high-scoring gene for the Trans-master and PPI-master gene scores (ranked 409 and 111) (Supplementary Figure S17C for WBC count, Supplementary Figure S17D for eosinophil count). *NFKB1* is a major transcription factor involved in immune response^74^ and is critical for development and proliferation of lymphocytes^75, 76^, and has previously been implicated in blood cell traits^45^. In each of these examples, we we nominate both the causal gene and the SNP-gene link.

We performed 4 secondary analyses. First, we investigated whether the 8 annotations of the combined joint model still contributed unique information after including the pLI gene score^47^, which has previously been shown to be conditionally informative for disease heritability^37, 41, 77^. We confirmed that all 8 annotations from Figure 5 remained jointly significant (Supplementary Figure S18 and Supplementary Table S68). Second, we considered integrating PPI network information via a single gene score (PPI-all) instead of two separate gene scores (PPI-enhancer and PPI-master). We determined that the combined joint model derived from PPI-all attained a similar combined *τ^*^* (2.5, s.e. 0.22; Supplementary Table S59; see Supplementary Table S69 for individual *τ\** values) as the combined joint model derived from PPI-enhancer and PPI-master (2.5, 0.24; Supplementary Table S59), and we believe it is less interpretable. Third, we constructed a less restrictive combined joint model by conditioning on the baseline-LD+ model instead of the baseline-LD+cis model. The less restrictive combined joint model included 1 additional annotation, SEG-GTEx *×* Coding (Supplementary Table S70). This implies that the combined joint model is largely invariant to conditioning on the baseline-LD+ or baseline-LD+cis model. Fourth, we analyzed binarized versions of all 11 gene scores (Table 1) using MAGMA^78^, an alternative gene set analysis method. 9 of the 11 gene scores produced significant signals (Supplementary Table S71), 11 marginally significant gene scores (Figure 3 and Figure 4) and 5 gene scores included in the combined joint model of Figure 5 in the S-LDSC analysis. However, MAGMA does not allow for conditioning on the baseline-LD model, does not allow for joint analysis of multiple gene scores to assess joint significance, and does not allow for incorporation of S2G strategies. Fifth, we confirmed that our forward stepwise elimination procedure produced identical results when applied to all 110 enhancer-related, candidate master-regulator, PPI-enhancer and PPI-master annotations, instead of just the 12 annotations from the PPI-enhancer (Figure 3D) and PPI-master (Figure 4D) joint models. Sixth, we assessed the model fit of the final joint model by correlating the residuals from stratified LD score regression with the independent variables in the regression (annotation-specific LD scores) for each of the 11 blood-related traits (Supplementary Figure S19). We observed an average squared correlation of 0.02 across annotation-specific LD scores and traits, suggesting good model fit.

We conclude that both enhancer-related genes and candidate master-regulator genes, as well as genes with high network connectivity to those genes, are jointly informative for autoimmune diseases and blood-related traits when using functionally informed S2G strategies.

### Discussion

#### Summary of findings

We have assessed the contribution to autoimmune disease of enhancer-related genes and candidate master-regulator genes, incorporating PPI network information and 10 functionally informed S2G strategies. We determined that our characterizations of enhancer-related and candidate master-regulator genes, informed by PPI networks, identify gene sets that are important for autoimmune disease, and that combining those gene sets with functionally informed S2G strategies enables us to identify SNP annotations in which disease heritability is concentrated. Our primary results were meta-analyzed across 11 autoimmune diseases and blood-related traits; we determined that results of meta-analyses across 6 autoimmune diseases and meta-analyses across 5 blood cell traits were quite similar, and that analyses of individual diseases/traits were generally underpowered. Our primary analyses used S-LDSC to assess functional enrichment, but analyses of functional enrichment of fine-mapped SNPs produced consistent results.

#### Biological significance

Our results provide information about which genes impact disease risk, distinguishing specific types of genes that play a greater role in genetic risk of disease (and have not previously been implicated in playing a greater role in genetic risk of disease). In some ways, our results distinguishing genes that are important for disease provide a quantitative improvement over previous work (e.g. vs. EDS-binary, a previously proposed enhancer-related gene score^24^). However, in other ways, our results provide qualitatively new findings (e.g. candidate master-regulator genes, and genes that interact with enhancer-related genes or master-regulator genes without being directly implicated). Our characterization of genes that are important for disease is validated by their enrichment in gold-standard gene sets, including autoimmune disease drug targets and Mendelian genes related to immune dysregulation; these enrichments were higher than for previously published characterizations. Notably, 22 out of 196 drug target genes were uniquely implicated by PPI-enhancer gene score as compared to other enhancer-regulated gene scores (based on top 10% genes) (Supplementary Table S72). These include three genes, *CCL2*, *IFNA1*, and *IKBKB* ^79–81^, are known to be particularly important for autoimmune disease; for *IKBKB*, we further note that the SNP *rs4737010* (chromosome 8) is a fine-mapped SNP for lymphocyte count that is implicated by our PPI-enhancer *×* ABC annotation (the annotation that is most conditionally informative for disease in our combined joint model; Figure 5, Supplementary Table S67). Similarly, 34 of 196 drug target genes were uniquely implicated by PPI-master gene score as compared to other candidate master-regulator gene scores (Supplementary Table S72). Furthermore, although gold-standard gene sets may be viewed as positive controls, our results are expected to also implicate true disease genes that are not previously known. Genes uniquely implicated by PPI-enhancer that may be important for autoimmune disease include *CD70*, a known target for cancer immunotherapy^82, 83^, and the *STAT* family genes (*STAT4, STAT5A and STAT6*), which serve to organize the epigenetic landscape of immune cells^84^ — both of which were not implicated by known gold-standard gene sets. Our results provide a route to performing functional follow-up experiments to elucidate and validate specific biological mechanisms (see below).

#### Downstream implications

Our work has several downstream implications. First, the PPI-enhancer gene score, which attained a particularly strong enrichment for approved autoimmune disease drug targets, will aid prioritization of drug targets that share similar characteristics as previously discovered drugs, analogous to pLI^47^ and LOEUF^85, 86^. Second, it is not practical to perform functional experiments on every SNP or genomic locus in the genome; using our results, specific gene-linked regulatory regions implicated by our results can be targeted for functional follow-up experiments (e.g. CRISPR base editing targeted at GWAS fine-mapped autoimmune disease SNPs linked to genes implicated by our gene scores) to elucidate and validate specific biological mechanisms. Third, our results implicate the ABC and Roadmap SNP-to-gene (S2G) linking strategies as highly informative distal S2G strategies, and TSS as a highly informative proximal S2G strategy, when linking SNPs to genes in analyses prioritizing genes or pathways; these S2G strategies should be used instead of or in combination with standard gene window-based S2G strategies. Fourth, our framework for disease heritability analysis incorporating regulatory S2G strategies (instead of conventional window-based approaches) is broadly applicable to other gene sets e.g. characterizing cell types and cellular processes, as in our more recent work^87^. Fifth, at the level of genes, our findings have immediate potential for improving gene-level probabilistic fine-mapping of transcriptome-wide association studies^88^ and gene-based association statistics^89^, using the gene scores as gene-level features to inform gene-level priors based on functional similarity of genes. Sixth, at the level of SNPs, our findings have immediate potential for improving functionally informed fine-mapping^51, 90–92^ (including experimental follow-up^93^), polygenic localization^51^, and polygenic risk prediction^94, 95^; specifically, SNP annotations derived from SNPs linked to high-scoring genes can be used to inform SNP-level priors used in these applications.

#### Limitations of study

Our work has several limitations, representing important directions for future research. First, we caution the readers that the terms ”enhancer-related genes” and ”candidate master-regulator” genes are inherently broad, and individual gene scores and annotations should be interpreted based on their specific meanings. Second, our results do not provide an understanding of specific biological mechanisms at individual disease loci, necessitating functional follow-up. Third, our findings distinguish specific types of genes that play a greater role in genetic risk of disease, but do not localize disease risk to a small number of genes, motivating more precise gene-level characterizations. Fourth, we restricted our analyses to enhancer-related and candidate master-regulator genes in blood, focusing on autoimmune diseases and blood-related traits; this choice was primarily motivated by the better representation of blood cell types in functional genomics assays and trans-eQTL studies. However, it will be valuable to extend our analyses to other tissues and traits as more functional data becomes available. Fifth, the trans-eQTL data from eQTLGen consortium^56^ is restricted to 10,317 previously disease-associated SNPs; we modified our analyses to account for this bias. However, it would be valuable to extend our analyses to genome-wide trans-eQTL data at large sample sizes, if that data becomes available. Sixth, we investigated the 10 S2G strategies separately, instead of constructing a single optimal combined strategy. A comprehensive evaluation of S2G strategies, and a method to combine them, will be provided elsewhere (S. Gazal, unpublished data). Seventh, the forward stepwise elimination procedure that we use to identify jointly significant annotations^29^ is a heuristic procedure whose choice of prioritized annotations may be close to arbitrary in the case of highly correlated annotations; however, the correlations between the gene scores, S2G strategies, and annotations that we analyzed were modest. Eigth, the potential of the gene scores implicated in this study to aid prioritization of future drug targets—based on observed gene-level enrichments for approved autoimmune disease drug targets—is subject to the limitation that novel drug targets that do not adhere to existing patterns may be missed; encouragingly, we also identify gene-level enrichments of the gene scores implicated in this study for 4 other “gold-standard” disease-related gene sets. Despite all these limitations, our findings expand and enhance our understanding of which gene-level characterizations of enhancer-related and candidate master-regulatory architecture and their corresponding gene-linked regions impact autoimmune diseases.

### Methods

#### Genomic annotations and the baseline-LD model

We define an annotation as an assignment of a numeric value to each SNP in a predefined reference panel (e.g., 1000 Genomes Project^31^; see URLs). Binary annotations can have value 0 or 1 only. Continuous-valued annotations can have any real value; our focus is on continuous-valued annotations with values between 0 and 1. Annotations that correspond to known or predicted function are referred to as functional annotations. The baseline-LD model (v.2.1) contains 86 functional annotations (see URLs). These annotations include binary coding, conserved, and regulatory annotations (e.g., promoter, enhancer, histone marks, TFBS) and continuous-valued linkage disequilibrium (LD)-related annotations.

#### Gene Scores

We define a gene score as an assignment of a numeric value between 0 and 1 to each gene; we primarily focus on binary gene sets defined by the top 10% of genes. We analyze a total of 11 gene scores (Table 1): 7 enhancer-related gene scores, 2 candidate master-regulator gene scores and 2 PPI-based gene scores (PPI-master, PPI-enhancer) that aggregate information across enhancer-related and candidate master-regulator gene scores. We scored 22,020 genes on chromosomes 1-22 from ref.^7^ (see URLs). When selecting the top 10% of genes for a given score, we rounded the number of genes to 2,200. We used the top 10% of genes in our primary analyses to be consistent with previous work^9^, who also defined gene scores using the top 10% of genes for a given metric, and to ensure that all SNP annotations (gene scores x S2G strategies) analyzed were of reasonable size (0.2% of SNPs or larger).

The 7 enhancer-related gene scores are as follows:

*•* **ABC-G**: A binary gene score denoting genes that are in top 10% of the number of ’intergenic’ and ’genic’ Activity-by-Contact (ABC) enhancer to gene links in blood cell types, with average HiC score fraction *>* 0.015^32^ (see URLs).
*•* **ATAC-distal**: A probabilistic gene score denoting the proportion of gene expression variance in 86 immune cell types in mouse, that is explained by the patterns of chromatin covariance of distal enhancer OCRs (open chromatin regions) to the gene, compared to chromatin covariance of OCRs that are near TSS of the gene and unexplained variances (see Figure 2 from^48^). The genes were mapped to their human orthologs using Ensembl biomaRt^96^.
*•* **EDS-binary**: A binary gene score denoting genes that are in top 10% of the blood-specific Activity-based Enhancer Domain Score (EDS)^24^ that reflects the number of conserved bases in enhancers that are linked to genes in blood related cell types as per the Roadmap Epigenomics Project^97, 98^ (see URLs).
*•* **eQTL-CTS**: A probabilistic gene score denoting the proportion of immune cell-type-specific eQTLs (with FDR adjusted p-value *<* 0.05 in one or two cell-types) across 15 different immune cell-types from the DICEdb project^99^ (see URLs). We consider this to be an enhancer-related gene score, as cell type specificity is often characterized by different enhancer activation status in different cell types^100, 101^.
*•* **Expecto-MVP**: A binary gene score denoting genes that are in top 10% in terms of the magnitude of variation potential (MVP) in GTEx Whole Blood, which is the sum of the absolute values of all directional mutation effects within 1kb of the TSS upstream and downstream, as evaluated by the Expecto method^7^ (see URLs). We consider this to be an enhancer-related gene score, as this score has been reported to be indicative of tissue specificity of expression and activation/repression status^7^.
*•* **PC-HiC-distal**: A binary gene score denoting genes that are in top 10% in terms of the total number of Promoter-capture HiC connections across 17 primary blood cell-types.
*•* **SEG-GTEx**: A binary gene score denoting genes that are in top 10% in terms of the SEG t-statistic^9^ score in GTEx Whole Blood. We consider this to be an enhancer-related gene score, as tissue specificity is often characterized by different enhancer activation status in different tissues^100, 101^.

The 2 candidate master-regulator gene scores are as follows:

*•* **Trans-master**: A binary gene score denoting genes with significant trait-associated cis-eQTLs in blood that also act as significant trans-eQTLs for at least 3 other genes based on data from eQTLGen Consortium^56^. We used the threshold of trans-regulating *≥*3 genes in our primary analyses because this results in a gene score spanning *≈*10% of genes, analogous to other gene scores.
*•* **TF**: A binary gene score denoting genes that act as human transcription factors^57^. The 2 PPI-based gene scores are as follows:
*•* **PPI-enhancer**: A binary gene score denoting genes in top 10% in terms of closeness centrality measure to the disease informative enhancer-regulated gene scores. To get the closeness centrality metric, we first perform a Random Walk with Restart (RWR) algorithm^54^ on the STRING protein-protein interaction (PPI) network^53, 102^(see URLs) with seed nodes defined by genes in top 10% of the 4 enhancer-regulated gene scores with jointly significant disease informativeness (ABC-G, ATAC-distal, EDS-binary and SEG-GTEx). The closeness centrality score was defined as the average network connectivity of the protein products from each gene based on the RWR method.
*•* **PPI-master**: A binary gene score denoting genes in top 10% in terms of closeness centrality measure to the 2 disease informative candidate master-regulator gene scores (Trans-master and TF). The algorithm was same as that of PPI-enhancer.

#### S2G strategies

We define a SNP-to-gene (S2G) linking strategy as an assignment of 0, 1 or more linked genes to each SNP with minor allele count *≥* 5 in a 1000 Genomes Project European reference panel^31^. We explored 10 SNP-to-gene linking strategies, including both distal and proximal strategies (Table 2). The proximal strategies included gene body *±* 5kb; gene body *±* 100kb; predicted TSS (by Segway^103, 104^) ; coding SNPs; and promoter SNPs (as defined by UCSC^105, 106^). The distal strategies included regions predicted to be distally linked to the gene by Activity-by-Contact (ABC) score^32, 33^ *>* 0.015 as suggested in ref.^33^ (see below); regions predicted to be enhancer-gene links based on Roadmap Epigenomics data (Roadmap)^97, 98, 107^; regions in ATAC-seq peaks that are highly correlated (*>* 50% as recommended in ref.^48^) to expression of a gene in mouse immune cell-types (ATAC)^48^; regions distally connected through promoter-capture Hi-C links (PC-HiC)^108^; and SNPs with fine-mapped causal posterior probability (CPP)^37^ *>* 0.001 (we chose this threshold to ensure that the SNP annotations generated after combining the gene scores with the eQTL S2G strategy were of reasonable size (0.2% of SNPs or larger) for all gene scores analyzed) in GTEx whole blood (we use this thresholding on CPP to ensure adequate annotation size for annotations resulting from combining this S2G strategy with the gene scores studied in this paper).

#### Activity-by-Contact model predictions

We used the Activity-by-Contact (ABC) model (https://github.com/broadinstitute/ABC-Enhancer-Gene-Prediction) to predict enhancer-gene connections in each cell type, based on measurements of chromatin accessibility (ATAC-seq or DNase-seq) and histone modifications (H3K27ac ChIP-seq), as previously described^32, 33^. In a given cell type, the ABC model reports an “ABC score” for each element-gene pair, where the element is within 5 Mb of the TSS of the gene.

For each cell type, we:

*•* Called peaks on the chromatin accessibility data using MACS2 with a lenient p-value cutoff of 0.1.
*•* Counted chromatin accessibility reads in each peak and retained the top 150,000 peaks with the most read counts. We then resized each of these peaks to be 500bp centered on the peak summit. To this list we added 500bp regions centered on all gene TSS’s and removed any peaks overlapping blacklisted regions^109, 110^ (https://sites.google.com/site/anshulkundaje/projects/blacklists). Any resulting overlapping peaks were merged. We call the resulting peak set candidate elements.
*•* Calculated element Activity as the geometric mean of quantile normalized chromatin accessibility and H3K27ac ChIP-seq counts in each candidate element region.
*•* Calculated element-promoter Contact using the average Hi-C signal across 10 human Hi-C datasets as described below.
*•* Computed the ABC Score for each element-gene pair as the product of Activity and Contact, normalized by the product of Activity and Contact for all other elements within 5 Mb of that gene.

To generate a genome-wide averaged Hi-C dataset, we downloaded KR normalized Hi-C matrices for 10 human cell types (GM12878, NHEK, HMEC, RPE1, THP1, IMR90, HUVEC, HCT116, K562, KBM7). This Hi-C matrix (5 Kb) resolution is available here: ftp://ftp.broadinstitute.org/outgoing/lincRNA/average_hic/average_hic.v2.191020. tar.gz^32, 111^. For each cell type we performed the following steps.

*•* Transformed the Hi-C matrix for each chromosome to be doubly stochastic.
*•* We then replaced the entries on the diagonal of the Hi-C matrix with the maximum of its four neighboring bins.
*•* We then replaced all entries of the Hi-C matrix with a value of NaN or corresponding to Knight–Ruiz matrix balancing (KR) normalization factors ¡ 0.25 with the expected contact under the power-law distribution in the cell type.
*•* We then scaled the Hi-C signal for each cell type using the power-law distribution in that cell type as previously described.
*•* We then computed the “average” Hi-C matrix as the arithmetic mean of the 10 cell-type specific Hi-C matrices.

In each cell type, we assign enhancers only to genes whose promoters are “active” (i.e., where the gene is expressed and that promoter drives its expression). We defined active promoters as those in the top 60% of Activity (geometric mean of chromatin accessibility and H3K27ac ChIP-seq counts). We used the following set of TSSs (one per gene symbol) for ABC predictions: https://github.com/broadinstitute/ ABC-Enhancer-Gene-Prediction/blob/v0.2.1/reference/RefSeqCurated.170308.bed. CollapsedGeneBounds.bed. We note that this approach does not account for cases where genes have multiple TSSs either in the same cell type or in different cell types.

For intersecting ABC predictions with variants, we took the predictions from the ABC Model and applied the following additional processing steps: (i) We considered all distal element-gene connections with an ABC score *≥* 0.015, and all distal or proximal promoter-gene connections with an ABC score *≥* 0.1. (ii) We shrunk the *∼*500-bp regions by 150-bp on either side, resulting in a *∼*200-bp region centered on the summit of the accessibility peak. This is because, while the larger region is important for counting reads in H3K27ac ChIP-seq, which occur on flanking nucleosomes, most of the DNA sequences important for enhancer function are likely located in the central nucleosome-free region. (iii) We included enhancer-gene connections spanning up to 2 Mb.

#### Stratified LD score regression

Stratified LD score regression (S-LDSC) is a method that assesses the contribution of a genomic annotation to disease and complex trait heritability^28, 29^. S-LDSC assumes that the per-SNP heritability or variance of effect size (of standardized genotype on trait) of each SNP is equal to the linear contribution of each annotation

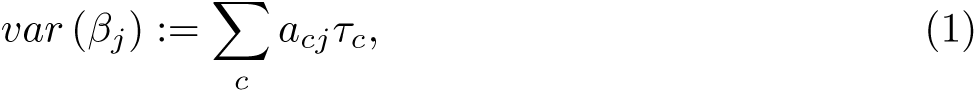

where *a_cj_* is the value of annotation *c* for SNP *j*, where *a_cj_* may be binary (0/1), continuous or probabilistic, and *τ_c_* is the contribution of annotation *c* to per-SNP heritability conditioned on other annotations. S-LDSC estimates the *τ_c_* for each annotation using the following equation

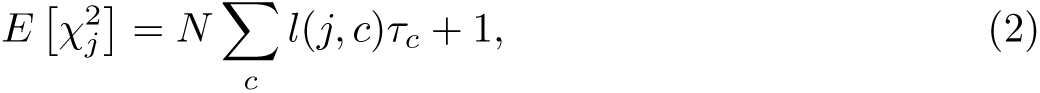

Where 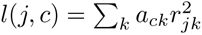 is the *stratified LD score* of SNP *j* with respect to annotation *c* and *r_jk_* is the genotypic correlation between SNPs *j* and *k* computed using data from 1000 Genomes Project^31^ (see URLs); N is the GWAS sample size.

We assess the informativeness of an annotation *c* using two metrics. The first metric is enrichment (*E*), defined as follows (for binary and probabilistic annotations only):

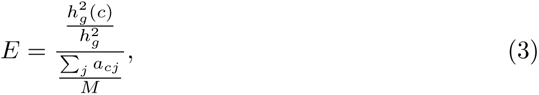

where 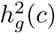 is the heritability explained by the SNPs in annotation *c*, weighted by the annotation values.

The second metric is standardized effect size (*τ\**) defined as follows (for binary, probabilistic, and continuous-valued annotations):

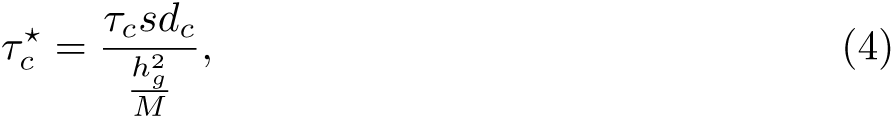

where *sd_c_* is the standard error of annotation 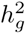 the total SNP heritability and *M* is the total number of SNPs on which this heritability is computed (equal to 5, 961, 159 in our analyses). 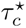 represents the proportionate change in per-SNP heritability associated to a 1 standard deviation increase in the value of the annotation.

#### Combined *τ^*^*

We defined a new metric quantifying the conditional informativeness of a heritability model (combined *τ^∗^*, generalizing the combined *τ^*^* metric of ref.^63^ to more than two annotations. In detail, given a joint model defined by *M* annotations (conditional on a published set of annotations such as the baseline-LD model), we define

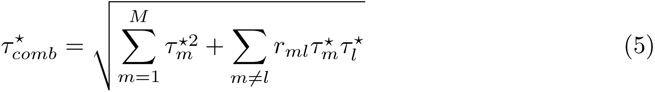

Here *r_ml_* is the pairwise correlation of the annotations *m* and *l*, and 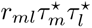 is expected to be positive since two positively correlated annotations typically have the same direction of effect (resp. two negatively correlated annotations typically have opposite directions of effect). We caclulate standard errors for 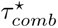 using a genomic block-jackknife with 200 blocks.

#### Evaluating heritability model fit using SumHer log***l_SS_***

Given a heritability model (e.g. the baseline-LD model or the combined joint model of Figure 5), we define the Δlog*l*_SS_ of that heritability model as the log*l*_SS_ of that heritability model minus the log*l*_SS_ of a model with no functional annotations (baseline-LD-nofunct; 17 LD and MAF annotations from the baseline-LD model^29^), where log*l*_SS_^64^ is an approximate likelihood metric that has been shown to be consistent with the exact likelihood from restricted maximum likelihood (REML). We compute p-values for Δlog*l*_SS_ using the asymptotic distribution of the Likelihood Ratio Test (LRT) statistic: *−*2 log*l*_SS_ follows a *χ*^2^ distribution with degrees of freedom equal to the number of annotations in the focal model, so that *−*2Δlog*l*_SS_ follows a *χ*^2^ distribution with degrees of freedom equal to the difference in number of annotations between the focal model and the baseline-LD-nofunct model. We used UK10K as the LD reference panel and analyzed 4,631,901 HRC (haplotype reference panel^34^) well-imputed SNPs with MAF *≥* and INFO *≥* 0.99 in the reference panel; We removed SNPs in the MHC region, SNPs explaining *>* 1% of phenotypic variance and SNPs in LD with these SNPs.

We computed Δlog*l*_SS_ for 8 heritability models:

• **baseline-LD model**: annotations from the baseline-LD model^29^ (86 annotations).
• **baseline-LD+ model**: baseline-LD model plus 7 new S2G annotations not included in the baseline-LD model (93 annotations).
• **baseline-LD+Enhancer model**: baseline-LD model+ plus 6 jointly significant S2G annotations c corresponding to enhancer-related gene scores from Supplementary Figure S11 (99 annotations).
• **baseline-LD+PPI-enhancer model**: baseline-LD model+ plus 7 jointly significant S2G annotations c corresponding to enhancer-related and PPI-enhancer gene scores from Figure 3D (100 annotations).
• **baseline-LD+cis model**: baseline-LD+ plus 20 S2G annotations used to correct for confounding in evaluation of Trans-master gene score (see Results) (113 annotations).
• **baseline-LD+Master model**: baseline-LD+cis plus 4 jointly significant candidate master-regulator S2G annotations from Supplementary Figure S14 (117 annotations).
• **baseline-LD+PPI-master model**: baseline-LD+cis plus 4 jointly significant candidate master-regulator and PPI-master S2G annotations from Figure 4D (117 annotations).
• **baseline-LD+PPI-master model**: baseline-LD+cis plus 8 jointly significant enhancer-related, candidate master-regulator, PPI-enhancer and PPI-master S2G annotations from the final joint model in Figure 5 (121 annotations).

#### Data Availability

All summary statistics used in this paper are publicly available (see URLs). This work used summary statistics from the UK Biobank study (http://www.ukbiobank.ac.uk/). The summary statistics for UK Biobank is available online (see URLs). All gene scores, S2G links and SNP annotations analyzed in this study are publicly available here: https://data.broadinstitute.org/alkesgroup/LDSCORE/Dey_Enhancer_MasterReg. Supplementary Tables S14 and S67 are provided as excel files in the above link. We have also included annotations for 93 million Haplotype Reference Consortium (HRC) SNPs and 170 million TOPMed SNPs (Freeze 3A).

#### Code Availability

The codes used to generate SNP annotations from gene sets, and for performing PPI-informed integration of gene sets are available on Github: https://github.com/kkdey/ GSSG.

#### URLs

• Gene scores, S2G links, annotations

https://data.broadinstitute.org/alkesgroup/LDSCORE/Dey_Enhancer_MasterReg

• Github code repository and data

https://github.com/kkdey/GSSG

• Activity-by-Contact (ABC) S2G links:

https://www.engreitzlab.org/resources

• 1000 Genomes Project Phase 3 data:

ftp://ftp.1000genomes.ebi.ac.uk/vol1/ftp/release/20130502

• UK Biobank summary statistics:

https://data.broadinstitute.org/alkesgroup/UKBB/

• baseline-LD model annotations:

https://data.broadinstitute.org/alkesgroup/LDSCORE/

• BOLT-LMM software:

https://data.broadinstitute.org/alkesgroup/BOLT-LMM

• S-LDSC software:

https://github.com/bulik/ldsc

## Acknowledgments and Funding

We thank Ran Cui, Hilary Finucane, Sebastian Pott, John Platig, Xinchen Wang and Soumya Raychaudhuri for helpful discussions. This research was funded by NIH grants U01 HG009379, U01 HG012009, R01 MH101244, K99HG010160, R37

MH107649, R01 MH115676 and R01 MH109978. S.S.Kim was supported by NIH award F31HG010818. K.K.Dey is supported by an NIH Pathway to Independence (K99/R00) Award (K99HG012203). J.M.Engreitz was supported by an NHGRI Genomic Innovator Award (R35HG011324); by Gordon and Betty Moore and the BASE Research Initiative at the Lucile Packard Children’s Hospital at Stanford University; and an NIH Pathway to Independence Award (R00HG009917). This research was conducted using the UK Biobank Resource under application 16549.

## 1 Supplementary Tables

**Table S1.**
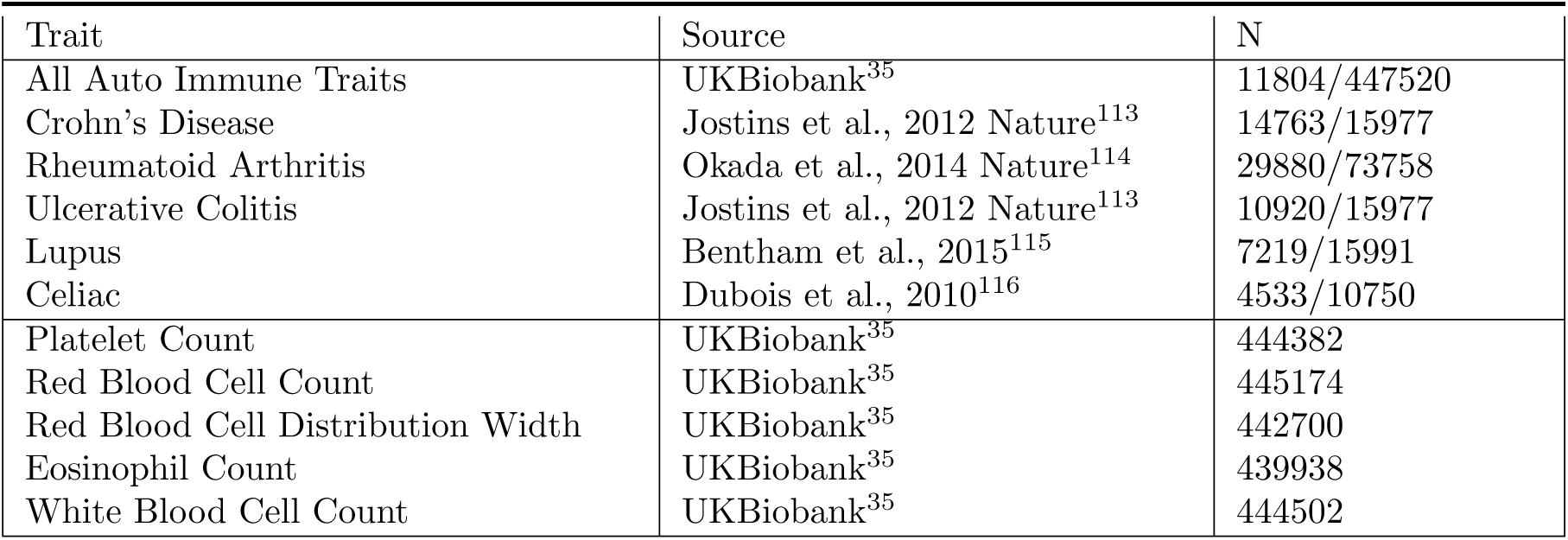
List of all blood-related traits: List of 11 blood-related traits (6 autoimmune diseases and 5 blood cell traits) analyzed in this paper. The disease called “All Auto Immune Traits” is based on the following codes and disease names that characterize autoimmune physiopathogenic etiology - 1222 (t1d; type 1 diabetes); 1256 (guillainBarre); 1260 (myasthenia); 1261 (ms); 1372 (vasculitis); 1378 (Wegener’s); 1381 (sle); 1382 (Sjogren); 1384 (sysSclerosis); 1437 (myasthenia); 1456 (celiac); 1464 (ra); 1522 (grave); 1661 (vitiligo)^37, 38^. For autoimmune disease case/control GWAS-es, we report *N* as *XXX/Y Y Y* where *XXX* represents the number of cases and *Y Y Y* represents the number of controls. The average number of cases for the autoimmune diseases in this study is equal to 13,186 and the average number of samples (N) for blood cell traits is 443K.

| Trait | Source | N |
| --- | --- | --- |
| All Auto Immune Traits | UKBiobank<sup>35</sup> | 11804/447520 |
| Crohn’s Disease | Jostins et al., 2012 Nature<sup>113</sup> | 14763/15977 |
| Rheumatoid Arthritis | Okada et al., 2014 Nature<sup>114</sup> | 29880/73758 |
| Ulcerative Colitis | Jostins et al., 2012 Nature<sup>113</sup> | 10920/15977 |
| Lupus | Bentham et al., 2015<sup>115</sup> | 7219/15991 |
| Celiac | Dubois et al., 2010<sup>116</sup> | 4533/10750 |
| Platelet Count | UKBiobank<sup>35</sup> | 444382 |
| Red Blood Cell Count | UKBiobank<sup>35</sup> | 445174 |
| Red Blood Cell Distribution Width | UKBiobank<sup>35</sup> | 442700 |
| Eosinophil Count | UKBiobank<sup>35</sup> | 439938 |
| White Blood Cell Count | UKBiobank<sup>35</sup> | 444502 |

**Table S2.** S-LDSC results for SNPs linked to all genes conditional on baseline-LD model: Standardized Effect sizes (*τ\**) and Enrichment (E) of 10 SNP annotations corresponding to all genes. The analysis is conditional on 86 baseline-LD (v2.1) annotations. Reports are meta-analyzed across 11 blood-related traits.

| All genes |  |  |  |  |  |  |
| --- | --- | --- | --- | --- | --- | --- |
| Annotation | $\tau^*$ | $se(\tau^*)$ | $p(\tau^*)$ | $E$ | $se(E)$ | $p(E)$ |
| ABC (1.3%) | 0.61 | 0.12 | 1.4e-07 | 11 | 1.1 | 5.4e-08 |
| TSS (1.6%) | 1.4 | 0.2 | 2e-12 | 14 | 0.8 | 5.3e-08 |
| Coding (1.6%) | 0.70 | 0.15 | 4.5e-06 | 6.2 | 0.70 | 8e-05 |
| ATAC (1.6%) | 0.42 | 0.1 | 5.1e-05 | 6.5 | 0.75 | 1.3e-07 |
| eQTL (2.4%) | 0.22 | 0.061 | 0.00034 | 4.6 | 0.35 | 1.9e-06 |
| Roadmap (3.1%) | 1 | 0.15 | 3.6e-11 | 8.9 | 0.74 | 5e-11 |
| Promoter (4.2%) | 0.063 | 0.079 | 0.36 | 3.8 | 0.38 | 7.1e-05 |
| PC-HiC (27.3%) | 0.13 | 0.041 | 0.0018 | 2.1 | 0.065 | 1.4e-10 |
| 5kb (52%) | 0.0013 | 0.023 | 0.95 | 1.4 | 0.025 | 6.9e-08 |
| 100kb (81%) | -0.009 | 0.0091 | 0.31 | 1.2 | 0.0067 | 8.9e-10 |

**Table S3.** Standardized enrichment of SNP annotations linked to all genes: Standardized enrichment of SNP annotations for SNPs linked to all genes, conditional on 86 baseline-LD annotations. Reports are meta-analyzed across 11 blood-related traits.

| All genes |  |  |  |
| --- | --- | --- | --- |
| | $StdE$ | $se(StdE)$ | $p(StdE)$ |
| ABC (1.3%) | 1.3 | 0.12 | 7e-08 |
| TSS (1.6%) | 1.8 | 0.12 | 2.3e-07 |
| Coding (1.6%) | 0.87 | 0.092 | 4.7e-05 |
| ATAC (1.6%) | 0.81 | 0.093 | 1.2e-07 |
| eQTL (2.4%) | 0.69 | 0.055 | 2.9e-06 |
| Roadmap (3.1%) | 1.5 | 0.12 | 1e-10 |
| Promoter (4.2%) | 0.88 | 0.072 | 2.9e-05 |
| PC-HiC (27.3%) | 0.93 | 0.028 | 1.6e-10 |
| 5kb (52%) | 0.69 | 0.013 | 5.3e-08 |
| 100kb (81%) | 0.44 | 0.0026 | 8e-10 |

**Table S4.** S-LDSC results of a joint analysis of all S2G annotations for SNPs linked to all genes conditional on baseline-LD annotations: Standardized Effect sizes (*τ\**) and Enrichment (E) of SNP annotations in a joint analysis comprising of 10 SNP annotations corresponding to all genes. The analysis is conditional on 86 baseline-LD annotations. Reports are meta-analyzed across 11 blood-related traits.

| All genes |  |  |  |  |  |  |
| --- | --- | --- | --- | --- | --- | --- |
| Annotation | $\tau^*$ | $se(\tau^*)$ | $p(\tau^*)$ | $E$ | $se(E)$ | $p(E)$ |
| ABC (1.3%) | 0.44 | 0.097 | 4.8e-06 | 9.1 | 0.82 | 8.8e-07 |
| TSS (1.6%) | 0.97 | 0.2 | 1.6e-06 | 12 | 0.9 | 3e-06 |
| Coding (1.6%) | 0.33 | 0.15 | 0.022 | 5.1 | 0.69 | 0.00038 |
| ATAC (1.6%) | 0.25 | 0.091 | 0.0058 | 5.5 | 0.67 | 1.3e-06 |
| eQTL (2.4%) | 0.14 | 0.066 | 0.033 | 4.1 | 0.38 | 9.9e-06 |
| Roadmap (3.1%) | 0.84 | 0.14 | 6.1e-09 | 8 | 0.67 | 2.5e-10 |
| Promoter (4.2%) | -0.39 | 0.16 | 0.01 | 3.2 | 0.47 | 0.00078 |
| PC-HiC (27.3%) | 0.081 | 0.04 | 0.043 | 2.1 | 0.06 | 2e-10 |
| 5kb (52%) | -0.036 | 0.029 | 0.21 | 1.4 | 0.028 | 8.4e-08 |
| 100kb (81%) | -0.014 | 0.013 | 0.27 | 1.2 | 0.0067 | 6.3e-10 |

**Table S5.** S-LDSC results for SNP annotations based on (random genes *×* S2G strategies): Maximum standardized Effect sizes (*τ\**) (with significance) and maximum Enrichment (E) (with significance) of SNP annotations corresponding to 10 different negative control gene sets where each gene set is generated by taking 10% genes randomly sampled from the list of protein coding genes without replacing. All results are conditional on 93 baseline-LD+ annotations. Reports are meta-analyzed across 11 blood-related traits. Size of the annotation is reported in bracket.

| Negative control |  |  |  |  |  |  |
| --- | --- | --- | --- | --- | --- | --- |
| | $\tau^*$ | $se(\tau^*)$ | $p(\tau^*)$ | $E$ | $se(E)$ | $p(E)$ |
| ABC (0.58%) | 0.34 | 0.15 | 0.019 | 14 | 1.8 | 1.2e-05 |
| TSS (0.33%) | 0.21 | 0.1 | 0.04 | 16 | 2.2 | 0.00035 |
| Coding (0.24%) | 0.1 | 0.08 | 0.2 | 7.7 | 1.8 | 0.00338 |
| ATAC (0.41%) | 0.036 | 0.091 | 0.69 | 6 | 2 | 0.05 |
| eQTL (0.38%) | 0.008 | 0.04 | 0.83 | 4.6 | 0.78 | 0.00258 |
| Roadmap (1.05%) | 0.056 | 0.082 | 0.5 | 8.6 | 1.2 | 2.1e-05 |
| Promoter (0.64%) | 0.092 | 0.076 | 0.22 | 3.9 | 0.88 | 0.011 |
| PC-HiC (8.66%) | -0.006 | 0.03 | 0.83 | 2.4 | 0.13 | 7.8e-08 |
| 5kb (6.45%) | 0.043 | 0.019 | 0.022 | 1.7 | 0.068 | 4.4e-07 |
| 100kb (17.3%) | 0.018 | 0.016 | 0.27 | 1.7 | 0.044 | 2.1e-09 |

**Table S6.** Enrichment of gene scores for 5 “gold-standard” disease-related gene sets: Enrichment and bootstrap standard error of all probabilistic gene scores with respect to (a) 195 genes that are Phase2+ drug targets for immune related diseases^10, 43^, (b) 550 Mendelian genes related to “immune dysregulation” as per ref.^44^ (c) 390 Mendelian genes related to blood disorders curated in ref.^45^, (d) 146 “Bone Marrow/Immune” genes defined by the Developmental Disorders Database/Genotype-Phenotype Database (DDD/G2P)^46^ and (e) top 10 % pLI genes^47^.

| Gene Score | Immune drug targets | Mendelian (Freund) | Mendelian (Vuckovic) | DDD/G2P (Bone Marrow, Immune) | Top 10% pLI |
| --- | --- | --- | --- | --- | --- |
| ABC-G | 1.39<br>(0.07) | 1.76<br>(0.08) | 1.58<br>(0.05) | 1.35<br>(0.19) | 1.11<br>(0.04) |
| ATAC-distal | 1.79<br>(0.04) | 1.91<br>(0.07) | 1.37<br>(0.06) | 1.15<br>(0.08) | 1.21<br>(0.02) |
| EDS-binary | 2.14<br>(0.10) | 2.34<br>(0.09) | 1.74<br>(0.09) | 1.57<br>(0.19) | 1.61<br>(0.06) |
| eQTL-CTS | 1.16<br>(0.03) | 1.28<br>(0.06) | 1.14<br>(0.04) | 1.23<br>(0.06) | 1.07<br>(0.01) |
| Expecto-MVP | 1.67<br>(0.09) | 1.98<br>(0.06) | 1.37<br>(0.05) | 0.97<br>(0.15) | 0.60<br>(0.04) |
| PC-HiC-distal | 0.91<br>(0.07) | 1.46<br>(0.07) | 1.63<br>(0.05) | 1.57<br>(0.22) | 1.57<br>(0.05) |
| SEG-GTEx | 2.42<br>(0.11) | 2.39<br>(0.10) | 1.52<br>(0.10) | 0.75<br>(0.16) | 0.30<br>(0.03) |
| Trans-master | 1.42<br>(0.07) | 2.06<br>(0.06) | 1.49<br>(0.06) | 1.42<br>(0.18) | 1.20<br>(0.05) |
| TF | 0.53<br>(0.08) | 1.53<br>(0.07) | 1.15<br>(0.07) | 1.01<br>(0.23) | 1.16<br>(0.06) |
| PPI-enhancer | 5.26<br>(0.12) | 4.65<br>(0.14) | 3.22<br>(0.13) | 2.03<br>(0.22) | 1.72<br>(0.05) |
| PPI-master | 2.71<br>(0.11) | 3.38<br>(0.12) | 2.24<br>(0.11) | 2.03<br>(0.22) | 1.90<br>(0.06) |

**Table S7.** Excess overlap of 6 of the enhancer-related gene scores with the EDS-binary gene score: We report excess overlap of genes, as defined in ref^11^, in each of 6 enhancer-related gene scores (ABC-G, ATAC-distal, eQTL-CTS, Expecto-MVP, PC-HiC-binary and SEG-GTEx) as well as the PPI-enhancer gene score with respect to the EDS-binary gene score derived from the well-established Enhancer Domain Score (EDS) that characterizes enhancer-regulation in genes^24^. The excess overlap metric assess the relative enrichment in gene scores in the genes annotated in EDS-binary, compared to all genes. We also report excess overlap for a control score defined by a gene set with randomly chosen 10% genes.

| Gene Score | ENR | p(ENR) |
| --- | --- | --- |
| ABC-G | 1.72 | 2.1e-08 |
| ATAC-distal | 1.78 | 6.2e-08 |
| eQTL-CTS | 1.31 | 1.2e-04 |
| Expecto-MVP | 1.59 | 6.3e-06 |
| PC-HiC-distal | 1.68 | 2.3e-06 |
| SEG-GTE<sub>x</sub> | 1.64 | 4.4e-07 |
| PPI-enhancer | 3.4 | 1.4e-97 |
| Control | 0.98 | 0.74 |

**Table S8.** S-LDSC results for SNP annotations corresponding to enhancer-related gene scores: Standardized Effect sizes (*τ\**) and Enrichment (E) of 70 SNP annotations corresponding to 7 enhancer-related gene scores and 10 S2G strategies, conditional on 93 baseline-LD+ annotations. Reports are meta-analyzed across 11 blood-related traits.

| ABC-G |  |  |  |  |  |  |
| --- | --- | --- | --- | --- | --- | --- |
| | $\tau^*$ | $se(\tau^*)$ | $p(\tau^*)$ | $E$ | $se(E)$ | $p(E)$ |
| ABC (0.57%) | 0.74 | 0.16 | 5.9e-06 | 15 | 1.5 | 2.4e-07 |
| TSS (0.38%) | 0.64 | 0.11 | 2.5e-08 | 19 | 1.8 | 2.7e-06 |
| Coding (0.23%) | 0.54 | 0.11 | 3.6e-07 | 16 | 2.1 | 6.7e-05 |
| ATAC (0.30%) | 0.28 | 0.073 | 0.00015 | 12 | 1.8 | 4.6e-06 |
| eQTL (0.41%) | 0.19 | 0.057 | 0.0012 | 7.4 | 0.98 | 2.7e-05 |
| Roadmap (0.90%) | 0.38 | 0.1 | 0.00017 | 12 | 1.3 | 1.4e-09 |
| Promoter (0.60%) | 0.44 | 0.082 | 8.3e-08 | 10 | 1.1 | 2.8e-05 |
| PC-HiC (8.86%) | 0.2 | 0.052 | 9.1e-05 | 3.1 | 0.17 | 8.2e-11 |
| 5kb (5.95%) | 0.081 | 0.029 | 0.0057 | 3.1 | 0.13 | 1.1e-09 |
| 100kb (14.2%) | 0.12 | 0.022 | 3.8e-08 | 2.6 | 0.095 | 1.2e-10 |
| ATAC-distal |  |  |  |  |  |  |
| | $\tau^*$ | $se(\tau^*)$ | $p(\tau^*)$ | $E$ | $se(E)$ | $p(E)$ |
| ABC (0.93%) | 1.0 | 0.24 | 1.7e-05 | 12 | 1.1 | 1e-07 |
| TSS (0.85%) | 0.95 | 0.11 | 2.8e-17 | 16 | 1.1 | 2.4e-06 |
| Coding (0.72%) | 0.66 | 0.11 | 3.6e-09 | 11 | 1 | 2.5e-05 |
| ATAC (1.02%) | 0.63 | 0.15 | 1.9e-05 | 7.9 | 1 | 1.9e-07 |
| eQTL (1.11%) | 0.19 | 0.048 | 6.3e-05 | 5.4 | 0.44 | 4.6e-06 |
| Roadmap (1.91%) | 0.55 | 0.12 | 5.4e-06 | 9.6 | 0.86 | 1.6e-10 |
| Promoter (1.84%) | 0.63 | 0.073 | 3.3e-18 | 7.2 | 0.53 | 9.8e-06 |
| PC-HiC (17.4%) | 0.22 | 0.062 | 0.00027 | 2.5 | 0.089 | 1.3e-10 |
| 5kb (19.4%) | 0.079 | 0.019 | 3.6e-05 | 2.1 | 0.037 | 1.2e-09 |
| 100kb (43.0%) | 0.066 | 0.018 | 0.00021 | 1.8 | 0.036 | 1.4e-10 |
| EDS-binary |  |  |  |  |  |  |
| | $\tau^*$ | $se(\tau^*)$ | $p(\tau^*)$ | $E$ | $se(E)$ | $p(E)$ |
| ABC (0.44%) | 0.91 | 0.21 | 1.4e-05 | 20 | 2.5 | 2.6e-07 |
| TSS (0.26%) | 0.64 | 0.12 | 1.6e-07 | 23 | 2.6 | 2.2e-06 |
| Coding (0.22%) | 0.66 | 0.1 | 1.6e-10 | 19 | 2.1 | 2.7e-05 |
| ATAC (0.28%) | 0.56 | 0.16 | 0.00042 | 15 | 2.8 | 2.9e-06 |
| eQTL (0.32%) | 0.18 | 0.042 | 2.6e-05 | 7.9 | 0.72 | 4.5e-06 |
| Promoter (0.52%) | 0.53 | 0.086 | 8.4e-10 | 12 | 1.2 | 2.3e-06 |
| Roadmap (1.15%) | 0.48 | 0.089 | 9.1e-08 | 11 | 0.98 | 2e-10 |
| PC-HiC (7.63%) | 0.15 | 0.041 | 0.00029 | 3 | 0.17 | 1.6e-10 |
| 5kb (5.66%) | 0.096 | 0.02 | 1.6e-06 | 2.7 | 0.1 | 9e-10 |
| 100kb (14.6%) | 0.15 | 0.02 | 6.4e-14 | 2.5 | 0.088 | 8.3e-11 |
| eQTL-CTS |  |  |  |  |  |  |
| | $\tau^*$ | $se(\tau^*)$ | $p(\tau^*)$ | $E$ | $se(E)$ | $p(E)$ |
| ABC (0.92%) | -0.047 | 0.22 | 0.83 | 9.8 | 0.9 | 2.6e-06 |
| TSS (0.81%) | 0.69 | 0.14 | 3.1e-07 | 14 | 1.2 | 9.9e-06 |
| Coding (0.81%) | 0.26 | 0.11 | 0.015 | 7.2 | 0.9 | 0.00025 |
| ATAC (0.78%) | 0.12 | 0.089 | 0.18 | 5.6 | 0.65 | 1.4e-05 |
| eQTL (1.09%) | 0.16 | 0.059 | 0.008 | 4.5 | 0.55 | 6.1e-05 |
| Promoter (2.06%) | 0.22 | 0.1 | 0.029 | 4.1 | 0.61 | 0.00072 |
| Roadmap (1.56%) | 0.16 | 0.11 | 0.15 | 8.3 | 0.68 | 5.3e-09 |
| PC-HiC (17.0%) | 0.18 | 0.046 | 0.00016 | 2.3 | 0.063 | 6.4e-10 |
| 5kb (22.2%) | 0.053 | 0.02 | 0.0068 | 1.7 | 0.046 | 3.9e-08 |
| 100kb (50.2%) | -0.013 | 0.016 | 0.42 | 1.5 | 0.028 | 4.5e-10 |
| Expecto-MVP |  |  |  |  |  |  |
| | $\tau^*$ | $se(\tau^*)$ | $p(\tau^*)$ | $E$ | $se(E)$ | $p(E)$ |
| ABC (0.49%) | 0.59 | 0.17 | 0.0004 | 16 | 2.1 | 3.2e-07 |
| TSS (0.29%) | 0.25 | 0.11 | 0.023 | 15 | 1.9 | 4.3e-05 |
| Coding (0.21%) | 0.41 | 0.15 | 0.0047 | 13 | 2.8 | 0.00044 |
| ATAC (0.27%) | 0.35 | 0.1 | 0.00069 | 13 | 2.3 | 3e-06 |
| eQTL (0.38%) | 0.11 | 0.047 | 0.018 | 6.3 | 0.71 | 1.8e-05 |
| Roadmap (0.80%) | 0.31 | 0.13 | 0.012 | 12 | 1.5 | 1.8e-09 |
| Promoter (0.57%) | 0.4 | 0.11 | 0.00015 | 9.1 | 1.1 | 1.6e-05 |
| PC-HiC (7.08%) | 0.03 | 0.05 | 0.55 | 2.8 | 0.18 | 7.9e-10 |
| 5kb (4.9%) | 0.093 | 0.03 | 0.0021 | 2.9 | 0.16 | 3e-10 |
| 100kb (15.7%) | 0.074 | 0.027 | 0.0064 | 2.3 | 0.086 | 3.8e-11 |
| PC-HiC-distal |  |  |  |  |  |  |
| | $\tau^*$ | $se(\tau^*)$ | $p(\tau^*)$ | $E$ | $se(E)$ | $p(E)$ |
| ABC (0.35%) | 0.21 | 0.11 | 0.067 | 13 | 1.7 | 9.8e-06 |
| TSS (0.24%) | 0.26 | 0.099 | 0.0079 | 16 | 2 | 3.6e-05 |
| Coding (0.20%) | 0.14 | 0.1 | 0.18 | 9.3 | 2.2 | 0.002 |
| ATAC (0.16%) | 0.27 | 0.077 | 0.00052 | 13 | 1.8 | 2.1e-05 |
| eQTL (0.34%) | 0.04 | 0.049 | 0.41 | 4.5 | 0.65 | 8.4e-05 |
| Roadmap (0.47%) | 0.28 | 0.074 | 0.00017 | 11 | 1.1 | 3.4e-07 |
| Promoter (0.49%) | 0.14 | 0.079 | 0.083 | 5.9 | 0.96 | 0.00035 |
| PC-HiC (12.0%) | 0.029 | 0.042 | 0.49 | 2.4 | 0.098 | 3.4e-10 |
| 5kb (4.70%) | 0.0091 | 0.018 | 0.61 | 2.2 | 0.083 | 1.5e-08 |
| 100kb (12.2%) | 0.034 | 0.02 | 0.087 | 2 | 0.054 | 2.2e-09 |

| SEG-GTE<sub>x</sub> |  |  |  |  |  |  |
| --- | --- | --- | --- | --- | --- | --- |
| | $\tau^*$ | se($\tau^*$) | p($\tau^*$) | $E$ | se($E$) | p($E$) |
| ABC (0.40%) | 0.9 | 0.13 | 1.9e-11 | 20 | 1.8 | 1.7e-07 |
| TSS (0.21%) | 0.72 | 0.15 | 9.8e-07 | 26 | 3.1 | 1.6e-06 |
| Coding (0.17%) | 0.89 | 0.15 | 8.7e-09 | 24 | 3.2 | 1.8e-05 |
| ATAC (0.26%) | 0.51 | 0.15 | 0.00052 | 16 | 3 | 4.1e-06 |
| eQTL (0.30%) | 0.23 | 0.052 | 8.9e-06 | 8.9 | 1.1 | 4.8e-06 |
| Roadmap (0.81%) | 0.43 | 0.11 | 7.5e-05 | 12 | 1.3 | 1.7e-09 |
| Promoter (0.50%) | 0.51 | 0.073 | 4.1e-12 | 11 | 1 | 6e-06 |
| PC-HiC (5.58%) | 0.095 | 0.033 | 0.0044 | 3.2 | 0.2 | 9.9e-10 |
| 5kb (3.54%) | 0.16 | 0.031 | 2.1e-07 | 3.7 | 0.26 | 9.1e-10 |
| 100kb (13.4%) | 0.16 | 0.023 | 2.2e-11 | 2.4 | 0.096 | 2.5e-10 |

**Table S9.** Standardized enrichment of SNP annotations based on enhancer-related gene scores: Standardized enrichment of the enhancer-related SNP annotations, conditional on 93 baseline-LD+ annotations. Reports are meta-analyzed across 11 blood-related traits.

| ABC-G |  |  |  |
| --- | --- | --- | --- |
|  | *StdE* | *se(StdE)* | *p(StdE)* |
| ABC (0.57%) | 1.1 | 0.11 | 2.4e-07 |
| TSS (0.38%) | 1.2 | 0.11 | 2.7e-06 |
| Coding (0.23%) | 0.79 | 0.1 | 6.7e-05 |
| ATAC (0.30%) | 0.64 | 0.1 | 4.6e-06 |
| eQTL (0.41%) | 0.48 | 0.063 | 2.7e-05 |
| Roadmap (0.90%) | 1.1 | 0.12 | 1.4e-09 |
| Promoter (0.60%) | 0.77 | 0.082 | 2.8e-05 |
| PC-HiC (8.86%) | 0.88 | 0.049 | 8.2e-11 |
| 5kb (5.95%) | 0.74 | 0.031 | 1.1e-09 |
| 100kb (14.2%) | 0.92 | 0.033 | 1.2e-10 |
| ATAC-distal |  |  |  |
|  | *StdE* | *se(StdE)* | *p(StdE)* |
| ABC (0.93%) | 1.2 | 0.11 | 1e-07 |
| TSS (0.85%) | 1.4 | 0.098 | 2.4e-06 |
| Coding (0.72%) | 0.89 | 0.085 | 2.5e-05 |
| ATAC (1.02%) | 0.79 | 0.1 | 1.9e-07 |
| eQTL (1.11%) | 0.55 | 0.045 | 4.6e-06 |
| Roadmap (1.91%) | 1.3 | 0.12 | 1.6e-10 |
| Promoter (1.84%) | 0.95 | 0.069 | 9.8e-06 |
| PC-HiC (17.4%) | 0.91 | 0.033 | 1.3e-10 |
| 5kb (19.4%) | 0.82 | 0.014 | 1.2e-09 |
| 100kb (43.0%) | 0.87 | 0.017 | 1.4e-10 |
| EDS-binary |  |  |  |
|  | *StdE* | *se(StdE)* | *p(StdE)* |
| ABC (0.44%) | 1.3 | 0.16 | 2.6e-07 |
| TSS (0.26%) | 1.2 | 0.13 | 2.2e-06 |
| Coding (0.22%) | 0.89 | 0.098 | 2.7e-05 |
| ATAC (0.28%) | 0.83 | 0.16 | 3.1e-06 |
| eQTL (0.32%) | 0.44 | 0.04 | 4.5e-06 |
| Promoter (0.52%) | 0.83 | 0.09 | 2.3e-06 |
| Roadmap (1.15%) | 1.1 | 0.1 | 2e-10 |
| PC-HiC (7.63%) | 0.79 | 0.045 | 1.6e-10 |
| 5kb (5.66%) | 0.63 | 0.024 | 9e-10 |
| 100kb (14.6%) | 0.86 | 0.031 | 8.3e-11 |
| eQTL-CTS |  |  |  |
|  | *StdE* | *se(StdE)* | *p(StdE)* |
| ABC (0.92%) | 0.9 | 0.082 | 2.6e-06 |
| TSS (0.81%) | 0.61 | 0.075 | 0.00025 |
| Coding (0.81%) | 0.61 | 0.075 | 0.00025 |
| ATAC (0.78%) | 0.45 | 0.052 | 1.4e-05 |
| eQTL (1.09%) | 0.41 | 0.05 | 6.1e-05 |
| Promoter (2.06%) | 0.54 | 0.081 | 0.00072 |
| Roadmap (1.56%) | 0.93 | 0.075 | 5.3e-09 |
| PC-HiC (17.0%) | 0.81 | 0.022 | 6.4e-10 |
| 5kb (22.2%) | 0.62 | 0.017 | 3.9e-08 |
| 100kb (50.2%) | 0.72 | 0.013 | 4.5e-10 |
| Expecto-MVP |  |  |  |
|  | *StdE* | *se(StdE)* | *p(StdE)* |
| ABC (0.49%) | 1.1 | 0.15 | 3.2e-07 |
| TSS (0.29%) | 0.81 | 0.1 | 4.3e-05 |
| Coding (0.21%) | 0.62 | 0.13 | 0.00044 |
| ATAC (0.27%) | 0.67 | 0.12 | 3e-06 |
| eQTL (0.38%) | 0.39 | 0.044 | 1.8e-05 |
| Roadmap (0.80%) | 1 | 0.14 | 1.8e-09 |
| Promoter (0.57%) | 0.68 | 0.082 | 1.6e-05 |
| PC-HiC (7.08%) | 0.71 | 0.047 | 7.9e-10 |
| 5kb (4.9%) | 0.62 | 0.034 | 3e-10 |
| 100kb (15.7%) | 0.82 | 0.031 | 3.8e-11 |
| PC-HiC-distal |  |  |  |
|  | *StdE* | *se(StdE)* | *p(StdE)* |
| ABC (0.35%) | 0.75 | 0.1 | 9.8e-06 |
| TSS (0.24%) | 0.77 | 0.096 | 3.6e-05 |
| Coding (0.20%) | 0.42 | 0.097 | 0.002 |
| ATAC (0.16%) | 0.51 | 0.073 | 2.1e-05 |
| eQTL (0.34%) | 0.26 | 0.038 | 8.4e-05 |
| Roadmap (0.47%) | 0.76 | 0.074 | 3.4e-07 |
| Promoter (0.49%) | 0.41 | 0.067 | 0.00035 |
| PC-HiC (12.0%) | 0.77 | 0.032 | 3.4e-10 |
| 5kb (4.70%) | 0.47 | 0.018 | 1.5e-08 |
| 100kb (12.2%) | 0.65 | 0.017 | 2.2e-09 |
| SEG-GTE<sub>x</sub> |  |  |  |
|  | *StdE* | *se(StdE)* | *p(StdE)* |
| ABC (0.40%) | 1.3 | 0.11 | 1.7e-07 |
| TSS (0.21%) | 1.2 | 0.14 | 1.6e-06 |
| Coding (0.17%) | 0.98 | 0.13 | 1.8e-05 |
| ATAC (0.26%) | 0.8 | 0.15 | 4.1e-06 |
| eQTL (0.30%) | 0.49 | 0.058 | 4.8e-06 |
| Roadmap (0.81%) | 1.1 | 0.12 | 1.7e-09 |
| Promoter (0.50%) | 0.74 | 0.072 | 6e-06 |
| PC-HiC (5.58%) | 0.74 | 0.045 | 9.9e-10 |
| 5kb (3.54%) | 0.68 | 0.048 | 9.1e-10 |
| 100kb (13.4%) | 0.82 | 0.032 | 2.5e-10 |

**Table S10.** S-LDSC results for SNP annotations corresponding to enhancer-related gene scores meta-analyzed across 5 blood cell traits: Standardized Effect sizes (*τ\**) and Enrichment (E) of 80 SNP annotations corresponding to 7 enhancer-related gene scores and 1 PPI-enhancer, and 10 S2G strategies, conditional on 93 baseline-LD+ annotations. Reports are meta-analyzed across 5 blood cell traits.

| ABC-G |  |  |  |  |  |  |
| --- | --- | --- | --- | --- | --- | --- |
| | $\tau^*$ | $se(\tau^*)$ | $p(\tau^*)$ | $E$ | $se(E)$ | $p(E)$ |
| ABC (0.57%) | 0.43 | 0.16 | 0.0067 | 12 | 1.5 | 7.4e-13 |
| TSS (0.38%) | 0.57 | 0.14 | 5.2e-05 | 18 | 2.4 | 1.6e-15 |
| Coding (0.23%) | 0.58 | 0.13 | 7.3e-06 | 17 | 2.5 | 9.4e-10 |
| ATAC (0.30%) | 0.22 | 0.088 | 0.014 | 8.7 | 2 | 2.4e-06 |
| eQTL (0.41%) | 0.22 | 0.069 | 0.0011 | 8.7 | 0.99 | 5.1e-125 |
| Roadmap (0.90%) | 0.2 | 0.14 | 0.17 | 8.5 | 1.5 | 3.9e-07 |
| Promoter (0.60%) | 0.5 | 0.1 | 1.4e-06 | 11 | 1.3 | 1.1e-12 |
| PC-HiC (8.86%) | 0.092 | 0.05 | 0.063 | 2.7 | 0.14 | 5.3e-17 |
| 5kb (5.95%) | 0.077 | 0.031 | 0.013 | 3 | 0.13 | 5e-20 |
| 100kb (14.2%) | 0.1 | 0.028 | 0.00033 | 2.4 | 0.096 | 2.9e-26 |
| ATAC-distal |  |  |  |  |  |  |
| | $\tau^*$ | $se(\tau^*)$ | $p(\tau^*)$ | $E$ | $se(E)$ | $p(E)$ |
| ABC (0.93%) | 0.69 | 0.24 | 0.0035 | 11 | 1.2 | 2.2e-18 |
| TSS (0.85%) | 1.0 | 0.13 | 7.1e-15 | 16 | 1.2 | 1.7e-27 |
| Coding (0.72%) | 0.78 | 0.17 | 2.5e-06 | 11 | 1.2 | 2.8e-17 |
| ATAC (1.02%) | 0.41 | 0.19 | 0.028 | 5.9 | 1.2 | 2.6e-06 |
| eQTL (1.11%) | 0.16 | 0.072 | 0.024 | 6 | 0.59 | 3.1e-10 |
| Roadmap (1.91%) | 0.38 | 0.17 | 0.027 | 7.7 | 1 | 1.8e-11 |
| Promoter (1.84%) | 0.60 | 0.086 | 2e-12 | 7.4 | 0.61 | 5.3e-11 |
| PC-HiC (17.4%) | 0.12 | 0.053 | 0.026 | 2.3 | 0.075 | 3e-31 |
| 5kb (19.4%) | 0.08 | 0.022 | 0.00032 | 2.1 | 0.053 | 1.1e-22 |
| 100kb (43.0%) | 0.067 | 0.021 | 0.0016 | 1.8 | 0.048 | 1.5e-24 |
| EDS-binary |  |  |  |  |  |  |
| | $\tau^*$ | $se(\tau^*)$ | $p(\tau^*)$ | $E$ | $se(E)$ | $p(E)$ |
| ABC (0.44%) | 0.57 | 0.17 | 0.00084 | 16 | 2.3 | 1.6e-12 |
| TSS (0.26%) | 0.65 | 0.2 | 0.001 | 23 | 4.1 | 1.2e-10 |
| Coding (0.22%) | 0.68 | 0.12 | 1.8e-08 | 20 | 2.5 | 7e-13 |
| ATAC (0.28%) | 0.05 | 0.13 | 0.71 | 6.1 | 2.6 | 0.05 |
| eQTL (0.32%) | 0.15 | 0.052 | 0.0043 | 7.9 | 0.86 | 4.1e-15 |
| Promoter (0.52%) | 0.55 | 0.13 | 3.3e-05 | 12 | 1.9 | 2.1e-06 |
| Roadmap (1.15%) | 0.39 | 0.13 | 0.0024 | 8.5 | 1.1 | 3.1e-10 |
| PC-HiC (7.63%) | 0.065 | 0.042 | 0.12 | 2.6 | 0.17 | 2.8e-14 |
| 5kb (5.66%) | 0.095 | 0.024 | 7.9e-05 | 2.6 | 0.12 | 1.1e-16 |
| 100kb (14.6%) | 0.13 | 0.029 | 2.8e-06 | 2.3 | 0.11 | 2.7e-18 |
| eQTL-CTS |  |  |  |  |  |  |
| | $\tau^*$ | $se(\tau^*)$ | $p(\tau^*)$ | $E$ | $se(E)$ | $p(E)$ |
| ABC (0.92%) | 0.088 | 0.27 | 0.75 | 9.5 | 1 | 6.3e-15 |
| TSS (0.81%) | 0.83 | 0.16 | 2.7e-07 | 16 | 1.4 | 3.9e-21 |
| Coding (0.81%) | 0.34 | 0.12 | 0.0049 | 7.5 | 0.99 | 3.6e-10 |
| ATAC (0.78%) | 0.14 | 0.1 | 0.16 | 5 | 0.82 | 5e-08 |
| eQTL (1.09%) | 0.22 | 0.071 | 0.0016 | 5.8 | 0.44 | 6.1e-17 |
| Promoter (2.06%) | 0.39 | 0.1 | 0.00012 | 5.5 | 0.83 | 7.6e-06 |
| Roadmap (1.56%) | 0.21 | 0.13 | 0.12 | 7.4 | 0.82 | 1.8e-11 |
| PC-HiC (17.0%) | 0.18 | 0.056 | 0.0011 | 2.3 | 0.07 | 9.5e-24 |
| 5kb (22.2%) | 0.084 | 0.024 | 0.00048 | 1.8 | 0.054 | 1.2e-18 |
| 100kb (50.2%) | -0.0038 | 0.019 | 0.84 | 1.5 | 0.039 | 5.3e-25 |
| Expecto-MVP |  |  |  |  |  |  |
| | $\tau^*$ | $se(\tau^*)$ | $p(\tau^*)$ | $E$ | $se(E)$ | $p(E)$ |
| ABC (0.49%) | 0.2 | 0.16 | 0.19 | 11 | 1.8 | 1.8e-08 |
| TSS (0.29%) | 0.19 | 0.16 | 0.26 | 14 | 2.9 | 4.5e-08 |
| Coding (0.21%) | 0.14 | 0.14 | 0.34 | 8.9 | 2.8 | 0.0048 |
| ATAC (0.27%) | 0.2 | 0.15 | 0.18 | 8.8 | 3 | 0.0082 |
| eQTL (0.38%) | 0.057 | 0.061 | 0.35 | 6.4 | 0.91 | 1e-08 |
| Roadmap (0.80%) | -0.0024 | 0.16 | 0.99 | 7.3 | 1.8 | 0.00057 |
| Promoter (0.57%) | 0.15 | 0.11 | 0.17 | 7.2 | 1.2 | 2.4e-05 |
| PC-HiC (7.08%) | -0.066 | 0.042 | 0.11 | 2.4 | 0.17 | 1.5e-10 |
| 5kb (4.9%) | 0.031 | 0.037 | 0.39 | 2.5 | 0.18 | 6.5e-11 |
| 100kb (15.7%) | 0.014 | 0.029 | 0.63 | 2.1 | 0.096 | 6.7e-17 |
| PC-HiC-distal |  |  |  |  |  |  |
| | $\tau^*$ | $se(\tau^*)$ | $p(\tau^*)$ | $E$ | $se(E)$ | $p(E)$ |
| ABC (0.35%) | 0.077 | 0.13 | 0.57 | 10 | 1.7 | 1.8e-07 |
| TSS (0.24%) | 0.18 | 0.12 | 0.14 | 15 | 2.3 | 1e-08 |
| Coding (0.20%) | 0.11 | 0.11 | 0.31 | 9 | 2.3 | 0.00077 |
| ATAC (0.16%) | 0.26 | 0.11 | 0.017 | 11 | 2.2 | 2e-05 |
| eQTL (0.34%) | -0.015 | 0.078 | 0.85 | 4.7 | 1 | 0.0014 |
| Roadmap (0.47%) | 0.26 | 0.09 | 0.0043 | 9.8 | 1.3 | 1.2e-09 |
| Promoter (0.49%) | 0.021 | 0.081 | 0.8 | 5.2 | 1.1 | 0.00063 |
| PC-HiC (12.0%) | -0.065 | 0.041 | 0.11 | 2.2 | 0.1 | 2.1e-22 |
| 5kb (4.70%) | -0.0062 | 0.021 | 0.77 | 2.2 | 0.096 | 9.8e-26 |
| 100kb (12.2%) | 0.03 | 0.025 | 0.23 | 2 | 0.064 | 7.2e-33 |

| SEG-GTE<sub>x</sub> |  |  |  |  |  |  |
| --- | --- | --- | --- | --- | --- | --- |
| | $\tau^*$ | $se(\tau^*)$ | $p(\tau^*)$ | $E$ | $se(E)$ | $p(E)$ |
| ABC (0.40%) | 0.74 | 0.17 | 9.3e-06 | 18 | 2.2 | 2.4e-13 |
| TSS (0.21%) | 0.62 | 0.23 | 0.0062 | 24 | 5.1 | 1.5e-12 |
| Coding (0.17%) | 0.68 | 0.14 | 1.4e-06 | 21 | 3.1 | 1.1e-11 |
| ATAC (0.26%) | 0.13 | 0.14 | 0.35 | 8 | 3.1 | 0.016 |
| eQTL (0.30%) | 0.24 | 0.088 | 0.0077 | 9.3 | 1.7 | 1e-07 |
| Roadmap (0.81%) | 0.22 | 0.12 | 0.078 | 8.8 | 1.6 | 3.9e-08 |
| Promoter (0.50%) | 0.45 | 0.09 | 4.5e-07 | 10 | 1.3 | 1.1e-12 |
| PC-HiC (5.58%) | 0.038 | 0.036 | 0.28 | 2.8 | 0.2 | 6.5e-21 |
| 5kb (3.54%) | 0.095 | 0.04 | 0.016 | 3 | 0.31 | 1.2e-16 |
| 100kb (13.4%) | 0.11 | 0.026 | 3.2e-05 | 2.2 | 0.1 | 4.6e-37 |

| PPI-enhancer |  |  |  |  |  |  |
| --- | --- | --- | --- | --- | --- | --- |
| | $\tau^*$ | $se(\tau^*)$ | $p(\tau^*)$ | $E$ | $se(E)$ | $p(E)$ |
| ABC (0.58%) | 1.2 | 0.25 | 2.8e-06 | 17 | 2.4 | 1.2e-15 |
| TSS (0.33%) | 0.84 | 0.18 | 3.4e-06 | 23 | 3.1 | 1.1e-15 |
| Coding (0.24%) | 1.0 | 0.23 | 4e-06 | 25 | 3.7 | 1.2e-11 |
| ATAC (0.41%) | 0.47 | 0.21 | 0.022 | 10 | 2.9 | 0.0008 |
| eQTL (0.38%) | 0.17 | 0.097 | 0.086 | 7.7 | 1.5 | 4e-06 |
| Roadmap (1.05%) | 0.6 | 0.19 | 0.0017 | 10 | 1.7 | 2.8e-08 |
| Promoter (0.64%) | 0.72 | 0.13 | 1.4e-08 | 13 | 1.1 | 4.3e-15 |
| PC-HiC (8.66%) | 0.18 | 0.064 | 0.0055 | 2.8 | 0.17 | 4.3e-17 |
| 5kb (6.45%) | 0.19 | 0.027 | 6.9e-12 | 2.9 | 0.12 | 2.7e-19 |
| 100kb (17.3%) | 0.26 | 0.03 | 1.1e-18 | 2.6 | 0.1 | 1.4e-23 |

**Table S11.** S-LDSC results for SNP annotations corresponding to enhancer-related gene scores meta-analyzed across 6 autoimmune diseases: Standardized Effect sizes (*τ\**) and Enrichment (E) of 80 SNP annotations corresponding to 7 enhancer-related gene scores and 1 PPI-enhancer, and 10 S2G strategies, conditional on 93 baseline-LD+ annotations. Reports are meta-analyzed across 6 autoimmune diseases.

| ABC-G |  |  |  |  |  |  |
| --- | --- | --- | --- | --- | --- | --- |
| | $\tau^*$ | $se(\tau^*)$ | $p(\tau^*)$ | $E$ | $se(E)$ | $p(E)$ |
| ABC (0.57%) | 1.2 | 0.24 | 2.6e-07 | 19 | 2.5 | 7.3e-05 |
| TSS (0.38%) | 0.77 | 0.21 | 0.00018 | 21 | 3.2 | 0.00041 |
| Coding (0.23%) | 0.46 | 0.19 | 0.014 | 16 | 3.8 | 0.0056 |
| ATAC (0.30%) | 0.46 | 0.14 | 0.0014 | 15 | 2.5 | 0.00053 |
| eQTL (0.41%) | 0.15 | 0.097 | 0.11 | 6.3 | 1.6 | 0.0037 |
| Roadmap (0.90%) | 0.63 | 0.13 | 2e-06 | 14 | 1.3 | 1.2e-05 |
| Promoter (0.60%) | 0.34 | 0.13 | 0.012 | 8.6 | 1.8 | 0.0043 |
| PC-HiC (8.86%) | 0.33 | 0.072 | 5.3e-06 | 3.6 | 0.25 | 1e-05 |
| 5kb (5.95%) | 0.088 | 0.056 | 0.11 | 3.3 | 0.23 | 1.3e-05 |
| 100kb (14.2%) | 0.17 | 0.04 | 2.8e-05 | 2.9 | 0.13 | 4.8e-06 |
| ATAC-distal |  |  |  |  |  |  |
| | $\tau^*$ | $se(\tau^*)$ | $p(\tau^*)$ | $E$ | $se(E)$ | $p(E)$ |
| ABC (0.93%) | 1.7 | 0.37 | 7.1e-06 | 16 | 2 | 0.00011 |
| TSS (0.85%) | 0.79 | 0.23 | 0.00066 | 14 | 2.3 | 0.00057 |
| Coding (0.72%) | 0.39 | 0.2 | 0.057 | 9.1 | 2.2 | 0.0061 |
| ATAC (1.02%) | 0.94 | 0.2 | 2.1e-06 | 11 | 1.3 | 0.00015 |
| eQTL (1.11%) | 0.24 | 0.08 | 0.0027 | 4.7 | 0.62 | 0.0011 |
| Roadmap (1.91%) | 0.84 | 0.18 | 2e-06 | 11 | 0.94 | 2e-05 |
| Promoter (1.84%) | 0.72 | 0.14 | 2.4e-07 | 6.5 | 1.1 | 0.0036 |
| PC-HiC (17.4%) | 0.36 | 0.1 | 0.00048 | 2.7 | 0.14 | 6.2e-06 |
| 5kb (19.4%) | 0.075 | 0.037 | 0.041 | 2.1 | 0.081 | 7.5e-06 |
| 100kb (43.0%) | 0.067 | 0.036 | 0.065 | 1.8 | 0.06 | 2.4e-06 |
| EDS-binary |  |  |  |  |  |  |
| | $\tau^*$ | $se(\tau^*)$ | $p(\tau^*)$ | $E$ | $se(E)$ | $p(E)$ |
| ABC (0.44%) | 1.4 | 0.39 | 0.00032 | 26 | 4.6 | 0.00012 |
| TSS (0.26%) | 0.78 | 0.21 | 0.00018 | 25 | 4.1 | 0.0013 |
| Coding (0.22%) | 0.64 | 0.22 | 0.0036 | 18 | 4.3 | 0.0038 |
| ATAC (0.28%) | 1.1 | 0.2 | 7.3e-09 | 25 | 3.2 | 0.00016 |
| eQTL (0.32%) | 0.23 | 0.072 | 0.0012 | 7.9 | 1.3 | 0.001 |
| Promoter (0.52%) | 0.57 | 0.15 | 0.00021 | 11 | 2.2 | 0.0025 |
| Roadmap (1.15%) | 0.61 | 0.13 | 5.3e-06 | 13 | 1.2 | 1.9e-0 |
| PC-HiC (7.63%) | 0.28 | 0.059 | 3.3e-06 | 3.5 | 0.22 | 7.5e-06 |
| 5kb (5.66%) | 0.097 | 0.036 | 0.0065 | 3 | 0.17 | 3e-05 |
| 100kb (14.6%) | 0.2 | 0.037 | 5.9e-08 | 2.6 | 0.11 | 1.1e-05 |
| eQTL-CTS |  |  |  |  |  |  |
| | $\tau^*$ | $se(\tau^*)$ | $p(\tau^*)$ | $E$ | $se(E)$ | $p(E)$ |
| ABC (0.92%) | -0.29 | 0.37 | 0.43 | 11 | 1.9 | 0.00074 |
| TSS (0.81%) | 0.37 | 0.25 | 0.13 | 11 | 2.1 | 0.0021 |
| Coding (0.81%) | -0.048 | 0.24 | 0.84 | 6.2 | 2.1 | 0.061 |
| ATAC (0.78%) | 0.04 | 0.19 | 0.83 | 7.6 | 1.3 | 0.0014 |
| eQTL (1.09%) | 0.0069 | 0.11 | 0.95 | 3.2 | 0.74 | 0.014 |
| Promoter (2.06%) | -0.092 | 0.14 | 0.52 | 2.1 | 0.83 | 0.5 |
| Roadmap (1.56%) | 0.054 | 0.2 | 0.78 | 9.7 | 0.97 | 2.8e-05 |
| PC-HiC (17.0%) | 0.16 | 0.082 | 0.053 | 2.4 | 0.11 | 5.5e-06 |
| 5kb (22.2%) | -0.0066 | 0.033 | 0.84 | 1.5 | 0.056 | 1.8e-05 |
| 100kb (50.2%) | -0.041 | 0.033 | 0.22 | 1.5 | 0.04 | 2e-06 |
| Expecto-MVP |  |  |  |  |  |  |
| | $\tau^*$ | $se(\tau^*)$ | $p(\tau^*)$ | $E$ | $se(E)$ | $p(E)$ |
| ABC (0.49%) | 1.1 | 0.24 | 2e-06 | 22 | 2.9 | 6.5e-05 |
| TSS (0.29%) | 0.41 | 0.18 | 0.025 | 18 | 3.3 | 0.0042 |
| Coding (0.21%) | 0.75 | 0.22 | 0.00049 | 20 | 4.3 | 0.0045 |
| ATAC (0.27%) | 0.53 | 0.13 | 5.2e-05 | 17 | 2.5 | 0.00013 |
| eQTL (0.38%) | 0.18 | 0.072 | 0.01 | 6.2 | 1.1 | 0.00185 |
| Roadmap (0.80%) | 0.61 | 0.13 | 2.1e-06 | 15 | 1.4 | 2.3e-05 |
| Promoter (0.57%) | 0.74 | 0.14 | 9.2e-08 | 13 | 1.9 | 0.0011 |
| PC-HiC (7.08%) | 0.13 | 0.078 | 0.11 | 3.2 | 0.25 | 1.5e-05 |
| 5kb (4.9%) | 0.17 | 0.041 | 2e-05 | 3.3 | 0.2 | 3.9e-06 |
| 100kb (15.7%) | 0.16 | 0.037 | 2.3e-05 | 2.5 | 0.1 | 2.6e-06 |
| PC-HiC-distal |  |  |  |  |  |  |
| | $\tau^*$ | $se(\tau^*)$ | $p(\tau^*)$ | $E$ | $se(E)$ | $p(E)$ |
| ABC (0.35%) | 0.56 | 0.22 | 0.01 | 19 | 3.2 | 0.00088 |
| TSS (0.24%) | 0.43 | 0.17 | 0.012 | 18 | 4 | 0.0026 |
| Coding (0.20%) | 0.25 | 0.22 | 0.27 | 11 | 4.9 | 0.026 |
| ATAC (0.16%) | 0.3 | 0.14 | 0.025 | 16 | 3.3 | 0.002 |
| eQTL (0.34%) | 0.099 | 0.061 | 0.11 | 4.6 | 0.99 | 0.0049 |
| Roadmap (0.47%) | 0.32 | 0.13 | 0.013 | 13 | 1.8 | 0.00013 |
| Promoter (0.49%) | 0.33 | 0.14 | 0.019 | 7.5 | 1.8 | 0.0089 |
| PC-HiC (12.0%) | 0.13 | 0.057 | 0.028 | 2.6 | 0.13 | 6.2e-06 |
| 5kb (4.70%) | 0.052 | 0.035 | 0.14 | 2.3 | 0.17 | 3.1e-05 |
| 100kb (12.2%) | 0.043 | 0.034 | 0.21 | 2.1 | 0.12 | 5.9e-06 |

| SEG-GTE<sub>x</sub> |  |  |  |  |  |  |
| --- | --- | --- | --- | --- | --- | --- |
| | $\tau^*$ | $se(\tau^*)$ | $p(\tau^*)$ | $E$ | $se(E)$ | $p(E)$ |
| ABC (0.40%) | 1.2 | 0.23 | 1.2e-07 | 27 | 3.2 | 0.00011 |
| TSS (0.21%) | 0.85 | 0.17 | 1.3e-06 | 28 | 3.9 | 0.00047 |
| Coding (0.17%) | 1.1 | 0.28 | 4.5e-05 | 29 | 6.1 | 0.0026 |
| ATAC (0.26%) | 0.88 | 0.16 | 7.7e-08 | 23 | 3.2 | 0.00042 |
| eQTL (0.30%) | 0.26 | 0.08 | 0.0013 | 8.3 | 1.5 | 0.00088 |
| Roadmap (0.81%) | 0.69 | 0.13 | 1.6e-07 | 15 | 1.4 | 3.9e-05 |
| Promoter (0.50%) | 0.6 | 0.12 | 1.2e-06 | 12 | 1.8 | 0.0015 |
| PC-HiC (5.58%) | 0.22 | 0.06 | 0.00025 | 3.8 | 0.26 | 1.9e-05 |
| 5kb (3.54%) | 0.25 | 0.043 | 4.3e-09 | 4.3 | 0.26 | 9.4e-06 |
| 100kb (13.4%) | 0.23 | 0.036 | 2.3e-10 | 2.7 | 0.12 | 7.3e-06 |

| PPI-enhancer |  |  |  |  |  |  |
| --- | --- | --- | --- | --- | --- | --- |
| | $\tau^*$ | $se(\tau^*)$ | $p(\tau^*)$ | $E$ | $se(E)$ | $p(E)$ |
| ABC (0.58%) | 2.8 | 0.3 | 1.8e-21 | 31 | 3 | 2.6e-05 |
| TSS (0.33%) | 1.6 | 0.19 | 6e-17 | 33 | 3.6 | 5.2e-05 |
| Coding (0.24%) | 1.8 | 0.21 | 6.1e-18 | 38 | 4.4 | 7.2e-05 |
| ATAC (0.41%) | 1.6 | 0.17 | 6.7e-21 | 27 | 2.7 | 4.5e-05 |
| eQTL (0.38%) | 0.59 | 0.082 | 6.1e-13 | 12 | 1.4 | 0.0001 |
| Roadmap (1.05%) | 1.6 | 0.14 | 2.1e-30 | 19 | 1.4 | 1.4e-05 |
| Promoter (0.64%) | 1.4 | 0.17 | 2.5e-17 | 20 | 2.1 | 5.7e-05 |
| PC-HiC (8.66%) | 0.46 | 0.059 | 9.3e-15 | 3.8 | 0.22 | 9.1e-06 |
| 5kb (6.45%) | 0.25 | 0.038 | 4.3e-11 | 3.4 | 0.18 | 1.6e-05 |
| 100kb (17.3%) | 0.38 | 0.038 | 7.6e-23 | 2.9 | 0.11 | 7e-06 |

**Table S12.**
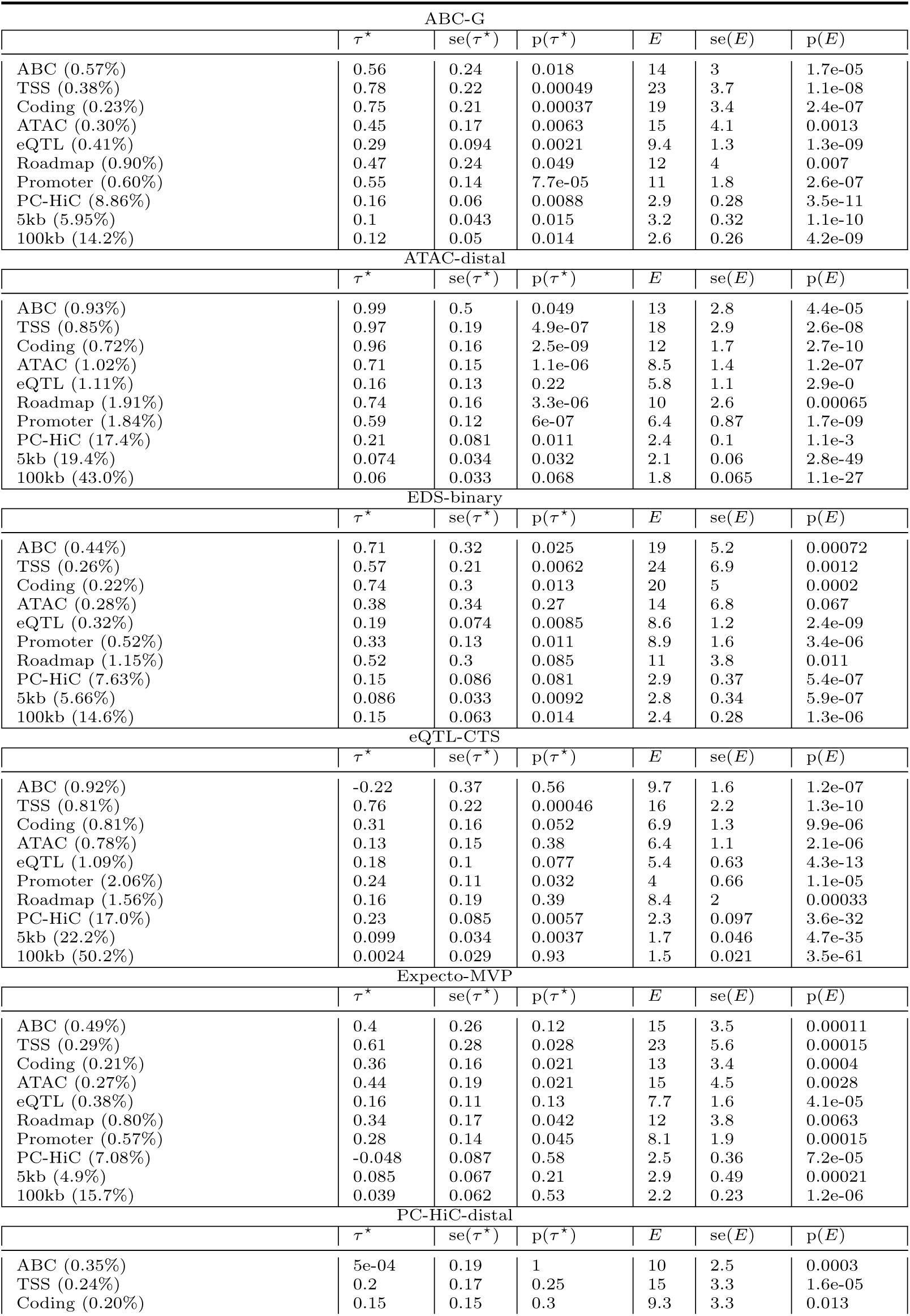
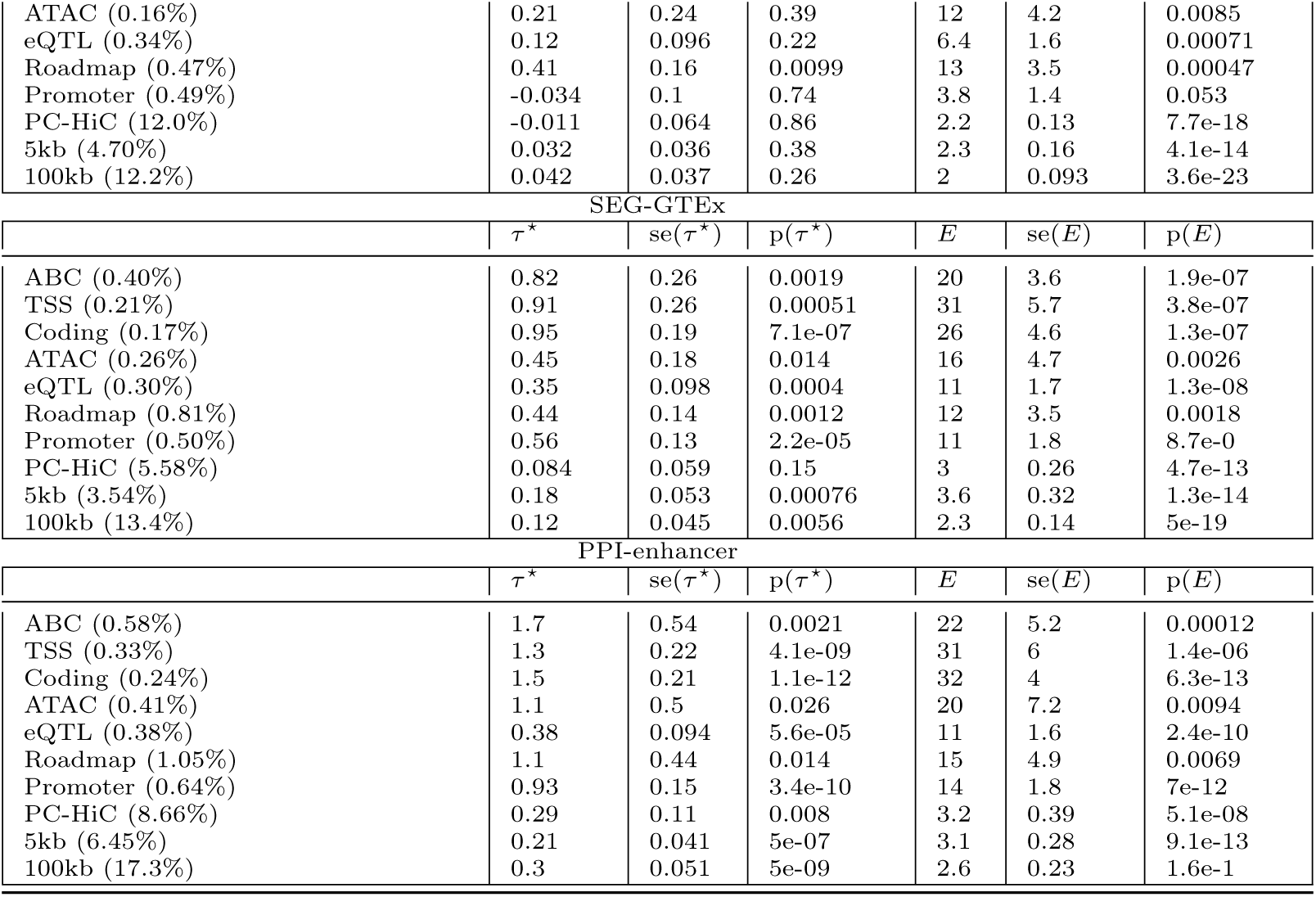
S-LDSC results for SNP annotations corresponding to enhancer-related gene scores meta-analyzed across 2 granulocyte related blood cell traits: Standardized Effect sizes (*τ\**) and Enrichment (E) of 80 SNP annotations corresponding to 7 enhancer-related gene scores and 1 PPI-enhancer gene score and 10 S2G strategies, conditional on 93 baseline-LD+ annotations. Reports are meta-analyzed across 2 granulocyte related traits (white blood cell count and eosinophil count).

**Table S13.**
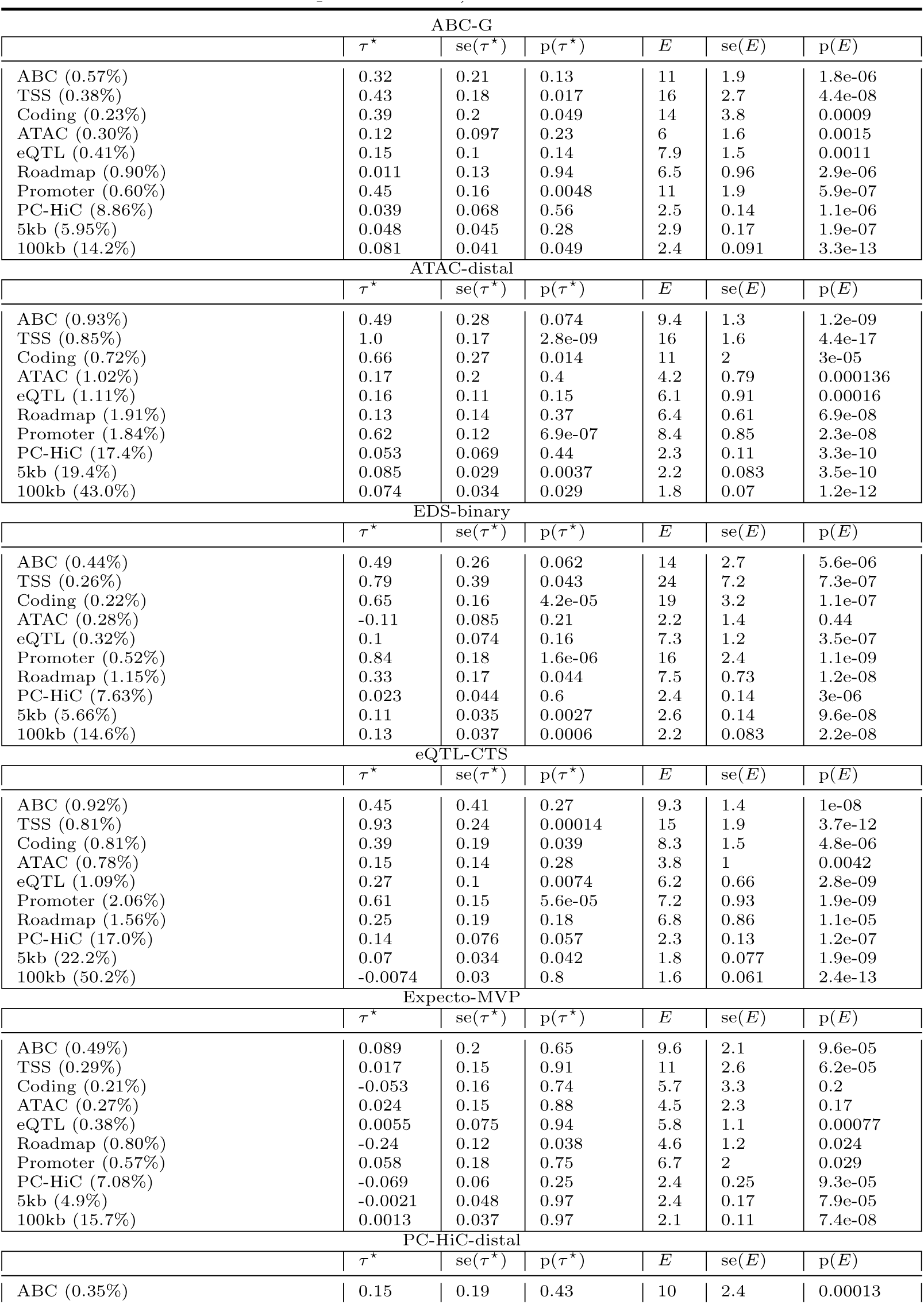
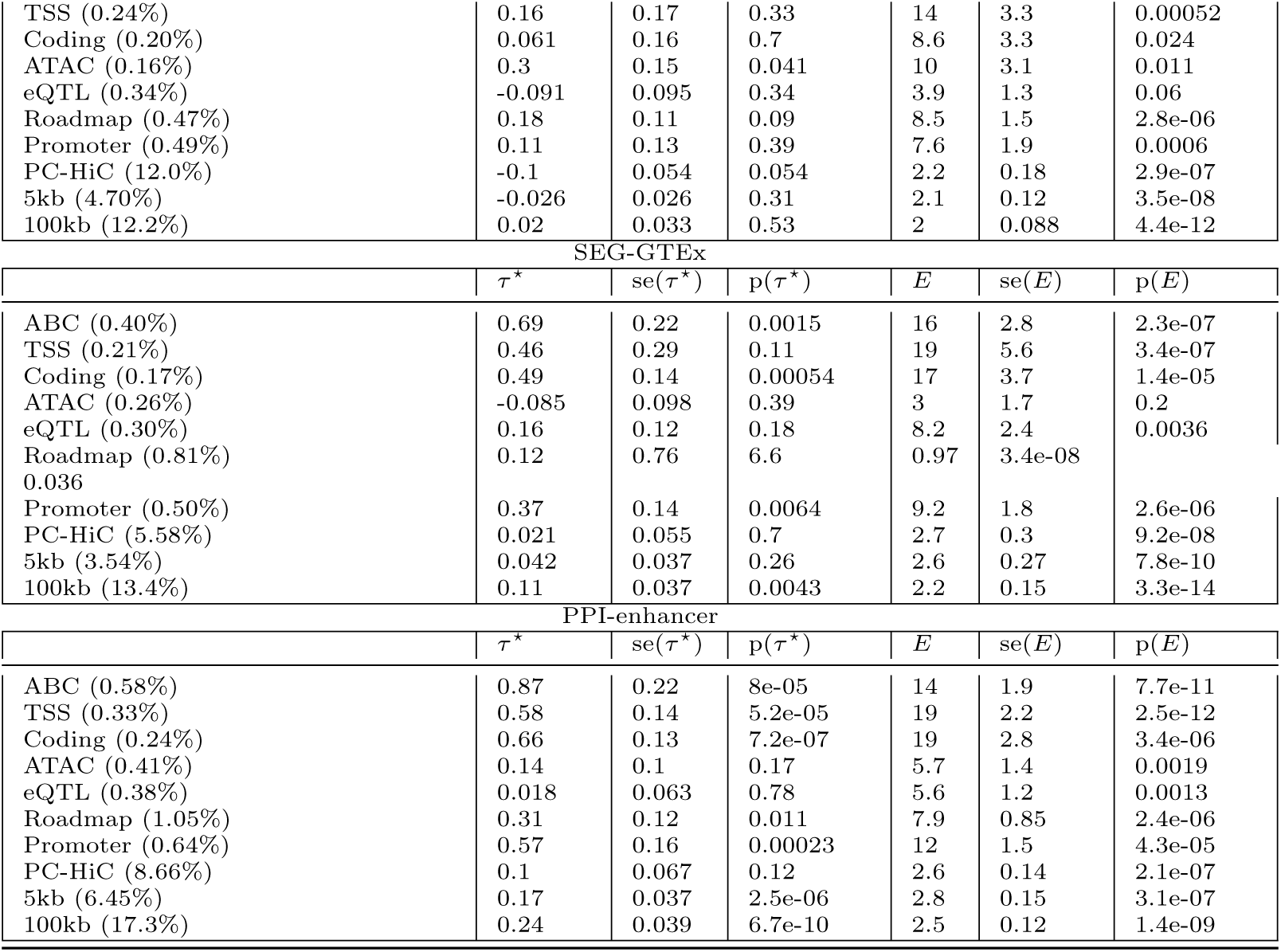
S-LDSC results for SNP annotations corresponding to enhancer-related gene scores meta-analyzed across 3 red blood cell or platelet-related blood cell traits: Standardized Effect sizes (*τ\**) and Enrichment (E) of 80 SNP annotations corresponding to 7 enhancer-related gene scores and 1 PPI-enhancer gene score and 10 S2G strategies, conditional on 93 baseline-LD+ annotations. Reports are meta-analyzed across 3 red blood cell or platelet related traits (red blood count, red blood distribution width and platelet count).

**Table S14.**
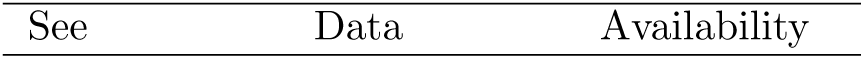
S-LDSC results for SNP annotations corresponding to all 11 gene scores for each of 11 autoimmune diseases and blood cell traits: Enrichment and standardized effect size estimates (*τ\**) corresponding to each of 11 blood-related traits and each of 110 annotations generated by combining the 11 gene scores described in the paper (7 enhancer-related, 1 PPI-enhancer, 2 candidate master-regulator and 1 PPI-master) with 10 2G strategies.

**Table S15.**
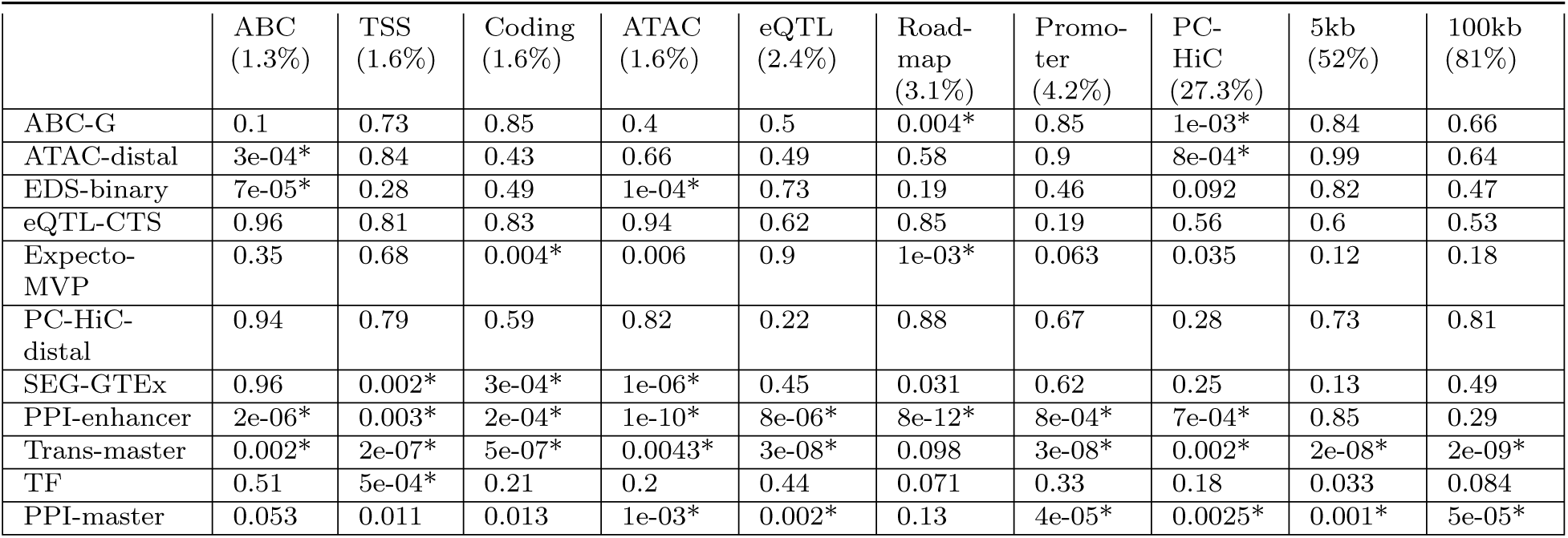
Results of test of heterogeneity of *τ^*^* values across traits. We report the significance (p-value) of the heterogeneity metric of standardized effect size (*τ\**) estimates across 11 blood-related traits, where heterogeneity is defined as per ref.^42^ (see Methods). Results are meta-analyzed across 11 blood-related traits.All FDR significant (*<* 5%) associations are marked by asterisk (*).

|  | ABC<br>(1.3%) | TSS<br>(1.6%) | Coding<br>(1.6%) | ATAC<br>(1.6%) | eQTL<br>(2.4%) | Road-<br>map<br>(3.1%) | Promo-<br>ter<br>(4.2%) | PC-HiC<br>(27.3%) | 5kb<br>(52%) | 100kb<br>(81%) |
| --- | --- | --- | --- | --- | --- | --- | --- | --- | --- | --- |
| ABC-G | 0.1 | 0.73 | 0.85 | 0.4 | 0.5 | 0.004* | 0.85 | 1e-03* | 0.84 | 0.66 |
| ATAC-distal | 3e-04* | 0.84 | 0.43 | 0.66 | 0.49 | 0.58 | 0.9 | 8e-04* | 0.99 | 0.64 |
| EDS-binary | 7e-05* | 0.28 | 0.49 | 1e-04* | 0.73 | 0.19 | 0.46 | 0.092 | 0.82 | 0.47 |
| eQTL-CTS | 0.96 | 0.81 | 0.83 | 0.94 | 0.62 | 0.85 | 0.19 | 0.56 | 0.6 | 0.53 |
| Expecto-MVP | 0.35 | 0.68 | 0.004* | 0.006 | 0.9 | 1e-03* | 0.063 | 0.035 | 0.12 | 0.18 |
| PC-HiC-distal | 0.94 | 0.79 | 0.59 | 0.82 | 0.22 | 0.88 | 0.67 | 0.28 | 0.73 | 0.81 |
| SEG-GTEx | 0.96 | 0.002* | 3e-04* | 1e-06* | 0.45 | 0.031 | 0.62 | 0.25 | 0.13 | 0.49 |
| PPI-enhancer | 2e-06* | 0.003* | 2e-04* | 1e-10* | 8e-06* | 8e-12* | 8e-04* | 7e-04* | 0.85 | 0.29 |
| Trans-master | 0.002* | 2e-07* | 5e-07* | 0.0043* | 3e-08* | 0.098 | 3e-08* | 0.002* | 2e-08* | 2e-09* |
| TF | 0.51 | 5e-04* | 0.21 | 0.2 | 0.44 | 0.071 | 0.33 | 0.18 | 0.033 | 0.084 |
| PPI-master | 0.053 | 0.011 | 0.013 | 1e-03* | 0.002* | 0.13 | 4e-05* | 0.0025* | 0.001* | 5e-05* |

**Table S16.** S-LDSC results for joint analysis of all marginally significant SNP annotations for enhancer-related genes. Standardized Effect sizes (*τ\**) and Enrichment (E) of the jointly significant enhancer-related SNP annotations from Table S8, conditional on the baseline-LD+ model and all SNP annotations in the enhancer-related joint model. Results are meta-analyzed across 11 blood-related traits.

| Annotation | $\tau^*$ | $\text{se}(\tau^*)$ | $\text{p}(\tau^*)$ | $E$ | $\text{se}(E)$ | $\text{p}(E)$ |
| --- | --- | --- | --- | --- | --- | --- |
| ABC-G × TSS<br>(0.38%) | 0.56 | 0.11 | 6.1e-07 | 18 | 1.7 | 3.1e-06 |
| ATAC-distal × Promoter<br>(1.84%) | 0.56 | 0.072 | 7.7e-15 | 6.9 | 0.51 | 1.2e-05 |
| EDS-binary × Roadmap<br>(1.15%) | 0.31 | 0.088 | 0.00041 | 8.6 | 0.89 | 3.2e-09 |
| EDS-binary × 100kb<br>(14.6%) | 0.096 | 0.02 | 2.4e-06 | 2.5 | 0.089 | 8.2e-11 |
| SEG-GTE<sub>x</sub> × ABC<br>(0.40%) | 0.68 | 0.13 | 2.1e-07 | 17 | 1.8 | 9.1e-07 |
| SEG-GTE<sub>x</sub> × Coding<br>(0.17%) | 0.68 | 0.15 | 8.3e-06 | 20 | 3.2 | 8.6e-05 |

**Table S17.** Enrichment of SNP annotations from different joint models with respect to fine-mapped SNPs for blood-related traits: We report the enrichment (E) and Jack-knife standard error with respect to 8,741 fine-mapped SNPs in autoimmune traits (Farh^50^) and 1,429 genome-wide functionally fine-mapped SNPs (PIP ¿ 0.95) in blood-related traits^51^. Annotations in the combined joint model in Figure 5 are marked in bold.

| enhancer-related joint model annotations: Supplementary Figure S11 |  |  |  |  |
| --- | --- | --- | --- | --- |
|  | Farh SNPs |  | Omer SNPs |  |
| Annotation | E | se(E) | E | se(E) |
| ABC-G × TSS | 5.67 | 0.26 | 12.5 | 0.41 |
| **ATAC-distal × Promoter** | 3.02 | 0.1 | 7.61 | 0.13 |
| EDS-binary × Roadmap | 5.67 | 0.17 | 5.60 | 0.13 |
| **EDS-binary × 100kb** | 2.22 | 0.04 | 2.00 | 0.02 |
| SEG-GTEx × ABC | 6.23 | 0.30 | 6.12 | 0.28 |
| SEG-GTEx × Coding | 7.60 | 0.41 | 14.0 | 1.16 |
| Master regulator joint model annotations: Supplementary Figure S14 |  |  |  |  |
|  | Farh SNPs |  | Omer SNPs |  |
| Annotation | E | se(E) | E | se(E) |
| **Trans-master × Roadmap** | 10.6 | 0.43 | 12.4 | 0.52 |
| **Trans-master × PC-HiC** | 5.1 | 0.22 | 3.5 | 0.08 |
| Trans-master × 5kb | 5.4 | 0.14 | 4.4 | 0.08 |
| TF × ATAC | 6.3 | 0.37 | 8.6 | 0.40 |
| TF × Roadmap | 7.3 | 0.36 | 8.3 | 0.35 |
| PPI-enhancer-related joint model annotations: Figure 3D |  |  |  |  |
|  | Farh SNPs |  | Omer SNPs |  |
| Annotation | E | se(E) | E | se(E) |
| **PPI-enhancer × ABC** | 9.0 | 0.50 | 8.7 | 0.32 |
| **PPI-enhancer × Coding** | 6.9 | 0.28 | 27.2 | 0.82 |
| PPI-enhancer × ATAC | 7.2 | 0.20 | 9.8 | 0.41 |
| PPI-enhancer × TSS | 6.5 | 0.28 | 16.1 | 0.51 |
| PPI-master-regulator joint model annotations: Figure 4D |  |  |  |  |
|  | Farh SNPs |  | Omer SNPs |  |
| Annotation | E | se(E) | E | se(E) |
| PPI-master × Coding | 7.1 | 0.37 | 22.9 | 0.95 |
| **PPI-master × ATAC** | 9.8 | 0.32 | 14.8 | 0.90 |
| **PPI-master × Roadmap** | 10.4 | 0.50 | 10.3 | 0.40 |

**Table S18.** Pathway enrichment analysis of the different gene scores: Pathway enrichment analysis of the top 10% genes for 7 enhancer-related gene scores (colored red), 2 candidate master-regulator gene scores (colored blue), PPI-enhancer and PPI-master gene scores (Methods). Pathway enrichment is performed using the ConsensusPathDB database^52, 117^. Only the top 5 non-redundant and statistically significant (q-value *<* 0.05) pathways for a gene set are reported.

| Gene Set | Top pathways |
| --- | --- |
| ABC-G | EGFR1 (7.1e-10), T cell receptor (8.5e-10), Neutrophil degranulation (9.8e-10), Interleukin signaling (2.2e-09), B cell receptor (4.2e-09) |
| ATAC-distal | Immune system (1.6e-11), Pathways in cancer(6.8e-10), White fat cell differentiation (5.5e-08),Cytokine signaling immune system (4.9e-07), EGFR1 (2.9e-06) |
| EDS-binary | T cell signaling (1.3e-12), T-cell receptor (1.3e-11), Th17 cell differentiation (8.8e-10), Hematopoietic cell lineage (1.5e-09), Pathways in cancer (1.9e-09) |
| eQTL-CTS | Olfactory Receptor Activity (3e-03), GPCR signaling (0.01), RUNX1 HSC differentiation (0.01) |
| Expecto-MVP | Immune system (7.3e-38), cytokine signaling (3.8e-26), interferon signaling (1.8e-13), Innate immune system (2.0e-13), Neutrophil degranulation (8.4e-13) |
| PC-HiC-distal | HDACs deacetylate histones (1.0e-20), HATs acetylate histones (3.7e-20), ERCC6 and EHM2 rRNA regulation (5.8e-19), chromatin organization (1.9e-18), RNA Pol 1 transcription (1.1e-17) |
| SEG-GTEx | Immune system (5.9e-99), Neutrophil degranulation (6.8e-55), Cytokine signaling (1.5e-24), NK cell mediated toxicity (3.6e-18), Hematopoietic cell lineage (1.6e-17) |
| Trans-master | Antigen processing and presentation (1.4e-17), Immune system (1.5e-13), Interferon gamma signaling (9.5e-11), Graft-versus-host disease (2.9e-10), Cytokine signaling (3.7e-10) |
| TF | Transcription pathway (2.4e-265), Neural receptor (1.9e-38), Transcription mis-regulation cancer (5.9e-25), Neural crest differentiation (4.3e-23), Adipogenesis (6.8e-22) |
| PPI-enhancer | Pathways in cancer (1.4e-119), Immune system (2.5e-116), Signal transduction (9.5e-112), Cytokine signaling immune system (7.7e-91), Innate immune system (5.5e-72) |
| PPI-master | Transcription pathway (8.1e-123), T-cell leukemia virus (7.0e-57), Viral carcinogenesis (5.7e-45), Hepatitis B (4.8e-43), JAK-STAT (6.3e-30) |

**Table S19.** S-LDSC results for SNP annotations corresponding to enhancer-related gene scores, using mean across genes of the gene scores linked to a SNP: Standardized Effect sizes (*τ\**) and Enrichment (E) of 70 SNP annotations corresponding to 7 enhancer-related gene scores and 10 S2G strategies, where SNPs linked to multiple genes are weighted by the mean of the gene scores of the genes they are linked to. The results are conditional on 93 baseline-LD+ annotations for enhancer-related annotaions and 113 baseline-LD+cis annotations for candidate master-regulator

| ABC-G |  |  |  |  |  |  |  |
| --- | --- | --- | --- | --- | --- | --- | --- |
| | $\tau^*$ | $se(\tau^*)$ | $p(\tau^*)$ | $p(\Delta\tau^*)$ | $E$ | $se(E)$ | $p(E)$ |
| ABC (0.55%) | 0.72 | 0.15 | 7.9e-07 | 0.49 | 16 | 1.5 | 6.8e-08 |
| ATAC (0.30%) | 0.26 | 0.069 | 0.0001 | 0.32 | 11 | 1.7 | 3.7e-06 |
| eQTL (0.38%) | 0.20 | 0.056 | 0.0001 | 0.22 | 7.3 | 0.98 | 3.6e-05 |
| Roadmap (0.80%) | 0.35 | 0.09 | 5.6e-05 | 0.30 | 10 | 1.4 | 5.6e-07 |
| PC-HiC (8.84%) | 0.22 | 0.054 | 2.4e-05 | 0.48 | 3.3 | 0.17 | 8.9e-11 |
| 5kb (5.94%) | 0.079 | 0.030 | 0.0054 | 0.38 | 3.2 | 0.15 | 4.6e-07 |
| 100kb (14.2%) | 0.11 | 0.025 | 5.4e-06 | 0.35 | 2.4 | 0.094 | 1.2e-06 |
| ATAC-distal |  |  |  |  |  |  |  |
| | $\tau^*$ | $se(\tau^*)$ | $p(\tau^*)$ | $p(\Delta\tau^*)$ | $E$ | $se(E)$ | $p(E)$ |
| ABC (0.77%) | 0.95 | 0.25 | 7.2e-05 | 0.29 | 10 | 1.1 | 1e-06 |
| ATAC (1.01%) | 0.26 | 0.06 | 7.4e-06 | 0.36 | 7.2 | 1 | 8.6e-06 |
| eQTL (1.05%) | 0.18 | 0.044 | 1.6e-05 | 0.20 | 5.4 | 0.44 | 4.6e-06 |
| Roadmap (1.67%) | 0.52 | 0.14 | 1.1e-04 | 0.25 | 9.0 | 0.82 | 3.4e-07 |
| PC-HiC (14.4%) | 0.20 | 0.057 | 2.2e-04 | 0.41 | 2.2 | 0.084 | 6.2e-07 |
| 5kb (18.5%) | 0.076 | 0.019 | 3.2e-05 | 0.35 | 2.0 | 0.037 | 6.4e-08 |
| 100kb (36.4%) | 0.066 | 0.018 | 0.0001 | 0.42 | 1.8 | 0.036 | 1.4e-10 |
| EDS-binary |  |  |  |  |  |  |  |
| | $\tau^*$ | $se(\tau^*)$ | $p(\tau^*)$ | $p(\Delta\tau^*)$ | $E$ | $se(E)$ | $p(E)$ |
| ABC (0.43%) | 0.87 | 0.22 | 3.8e-05 | 0.31 | 17 | 2.6 | 3.5e-06 |
| ATAC (0.28%) | 0.52 | 0.18 | 0.0008 | 0.30 | 15 | 2.9 | 3.3e-06 |
| eQTL (0.29%) | 0.17 | 0.042 | 2.6e-05 | 0.19 | 7.8 | 0.72 | 5.6e-06 |
| Roadmap (1.03%) | 0.47 | 0.088 | 4.6e-08 | 0.22 | 10 | 0.93 | 4.2e-09 |
| PC-HiC (7.62%) | 0.14 | 0.038 | 0.0001 | 0.27 | 2.7 | 0.16 | 3.7e-07 |
| 5kb (5.65%) | 0.094 | 0.02 | 2.2e-06 | 0.30 | 2.7 | 0.1 | 9e-10 |
| 100kb (14.6%) | 0.15 | 0.02 | 4.2e-14 | 0.39 | 2.5 | 0.087 | 5.6e-10 |
| eQTL-CTS |  |  |  |  |  |  |  |
| | $\tau^*$ | $se(\tau^*)$ | $p(\tau^*)$ | $p(\Delta\tau^*)$ | $E$ | $se(E)$ | $p(E)$ |
| ABC (0.76%) | -0.066 | 0.28 | 0.59 | 0.17 | 9.6 | 0.9 | 4.9e-06 |
| ATAC (0.75%) | 0.10 | 0.083 | 0.11 | 0.29 | 5.4 | 0.63 | 4.6e-05 |
| eQTL (1.04%) | 0.14 | 0.056 | 0.006 | 0.27 | 4.5 | 0.55 | 5.4e-05 |
| Roadmap (1.33%) | 0.14 | 0.12 | 0.12 | 0.20 | 7.9 | 0.66 | 6.2e-07 |
| PC-HiC (14.6%) | 0.16 | 0.048 | 0.0004 | 0.32 | 2.2 | 0.063 | 5.7e-09 |
| 5kb (20.2%) | 0.052 | 0.02 | 0.0062 | 0.39 | 1.7 | 0.045 | 5.7e-09 |
| 100kb (37.7%) | -0.022 | 0.014 | 0.82 | 0.54 | 1.3 | 0.027 | 6.8e-09 |
| Expecto-MVP |  |  |  |  |  |  |  |
| | $\tau^*$ | $se(\tau^*)$ | $p(\tau^*)$ | $p(\Delta\tau^*)$ | $E$ | $se(E)$ | $p(E)$ |
| ABC (0.48%) | 0.56 | 0.16 | 0.0002 | 0.42 | 16 | 2.1 | 4.6e-07 |
| ATAC (0.26%) | 0.33 | 0.1 | 0.0005 | 0.34 | 10 | 2.2 | 2.7e-05 |
| eQTL (0.36%) | 0.10 | 0.047 | 0.018 | 0.35 | 6.3 | 0.71 | 4.7e-05 |
| Roadmap (0.64%) | 0.30 | 0.13 | 0.01 | 0.28 | 11 | 1.5 | 4.6e-08 |
| PC-HiC (7.08%) | 0.09 | 0.07 | 0.15 | 0.38 | 2.4 | 0.17 | 1.2e-08 |
| 5kb (4.9%) | 0.089 | 0.03 | 0.0015 | 0.40 | 2.8 | 0.16 | 4.4e-09 |
| 100kb (15.7%) | 0.068 | 0.029 | 0.0091 | 0.38 | 2.1 | 0.084 | 2.6e-08 |
| PC-HiC-distal |  |  |  |  |  |  |  |
| | $\tau^*$ | $se(\tau^*)$ | $p(\tau^*)$ | $p(\Delta\tau^*)$ | $E$ | $se(E)$ | $p(E)$ |
| ABC (0.35%) | 0.20 | 0.11 | 0.034 | 0.42 | 11 | 1.9 | 2.8e-05 |
| ATAC (0.16%) | 0.25 | 0.076 | 0.0005 | 0.40 | 13 | 1.7 | 5.6e-05 |
| eQTL (0.32%) | 0.06 | 0.059 | 0.16 | 0.48 | 4.4 | 0.65 | 6.8e-05 |
| Roadmap (0.39%) | 0.25 | 0.077 | 0.0005 | 0.29 | 11 | 1.1 | 4.6e-08 |
| PC-HiC (12.0%) | 0.027 | 0.041 | 0.26 | 0.47 | 2.3 | 0.098 | 4.2e-09 |
| 5kb (4.70%) | 0.009 | 0.018 | 0.58 | 0.43 | 2.1 | 0.083 | 4.6e-07 |
| 100kb (12.0%) | 0.036 | 0.02 | 0.034 | 0.48 | 2.2 | 0.059 | 1.6e-08 |
| SEG-GTEx |  |  |  |  |  |  |  |
| | $\tau^*$ | $se(\tau^*)$ | $p(\tau^*)$ | $p(\Delta\tau^*)$ | $E$ | $se(E)$ | $p(E)$ |
| ABC (0.39%) | 0.88 | 0.12 | 1.1e-13 | 0.27 | 18 | 1.7 | 2.4e-06 |
| ATAC (0.26%) | 0.48 | 0.16 | 0.001 | 0.23 | 13 | 3.1 | 8.5e-05 |
| eQTL (0.27%) | 0.24 | 0.053 | 3.0e-06 | 0.24 | 8.8 | 1.1 | 4.2e-06 |
| Roadmap (0.66%) | 0.40 | 0.12 | 0.0004 | 0.18 | 10 | 1.3 | 4.4e-07 |
| PC-HiC (5.58%) | 0.094 | 0.031 | 0.001 | 0.28 | 3.0 | 0.2 | 7.4e-09 |
| 5kb (3.54%) | 0.15 | 0.031 | 3.4e-06 | 0.38 | 3.5 | 0.26 | 5.6e-09 |
| 100kb (13.4%) | 0.16 | 0.022 | 1.5e-10 | 0.32 | 2.4 | 0.096 | 3.4e-10 |
| PPI-enhancer |  |  |  |  |  |  |  |
| | $\tau^*$ | $se(\tau^*)$ | $p(\tau^*)$ | $p(\Delta\tau^*)$ | $E$ | $se(E)$ | $p(E)$ |
| ABC (0.58%) | 1.8 | 0.28 | 6.4e-11 | 0.28 | 19 | 2.5 | 4.8e-07 |
| ATAC (0.40%) | 1.0 | 0.22 | 2.7e-06 | 0.27 | 17 | 2.8 | 5.6e-07 |
| eQTL (0.35%) | 0.35 | 0.083 | 1.3e-05 | 0.16 | 9.5 | 1.2 | 4.2e-08 |
| Roadmap (0.89%) | 1.3 | 0.20 | 4.1e-11 | 0.33 | 13 | 1.6 | 7.2e-10 |
| PC-HiC (8.66%) | 0.28 | 0.052 | 3.6e-08 | 0.36 | 3.1 | 0.19 | 9.5e-09 |
| 5kb (6.45%) | 0.22 | 0.021 | 5.5e-26 | 0.42 | 3.1 | 0.11 | 3.8e-10 |
| 100kb (17.3%) | 0.30 | 0.026 | 6.4e-30 | 0.39 | 2.7 | 0.087 | 4.2e-10 |
| Trans-master |  |  |  |  |  |  |  |
| | $\tau^*$ | $se(\tau^*)$ | $p(\tau^*)$ | $p(\Delta\tau^*)$ | $E$ | $se(E)$ | $p(E)$ |
| ABC (0.33%) | 1.3 | 0.26 | 2.9e-07 | 0.18 | 26 | 2.4 | 1e-07 |
| ATAC (0.21%) | 0.88 | 0.15 | 2.2e-09 | 0.22 | 22 | 3.5 | 1.1e-05 |
| eQTL (0.51%) | 0.58 | 0.12 | 6.7e-07 | 0.22 | 10 | 1.5 | 8.2e-06 |
| Roadmap (0.49%) | 1.1 | 0.11 | 7.6e-24 | 0.23 | 20 | 1.4 | 5.3e-07 |
| PC-HiC (5.1%) | 0.45 | 0.042 | 4.4e-27 | 0.32 | 4.9 | 0.18 | 2.2e-08 |
| 5kb (4.1%) | 0.38 | 0.054 | 9.9e-13 | 0.36 | 5.3 | 0.29 | 2.5e-08 |
| 100kb (9.7%) | 0.36 | 0.046 | 2.5e-15 | 0.39 | 3.5 | 0.17 | 6.9e-07 |
| TF |  |  |  |  |  |  |  |
| | $\tau^*$ | $se(\tau^*)$ | $p(\tau^*)$ | $p(\Delta\tau^*)$ | $E$ | $se(E)$ | $p(E)$ |
| ABC (0.32%) | 0.44 | 0.12 | 0.0001 | 0.28 | 15 | 1.7 | 1.1e-05 |
| ATAC (0.14%) | 0.64 | 0.12 | 4.2e-08 | 0.26 | 18 | 3.0 | 6.8e-06 |
| eQTL (0.15%) | 0.05 | 0.063 | 0.21 | 0.42 | 3.8 | 0.76 | 0.0004 |
| Roadmap (0.25%) | 0.8 | 0.12 | 1.4e-11 | 0.26 | 20 | 2.1 | 2.2e-08 |
| PC-HiC (4.8%) | 0.22 | 0.035 | 3.6e-10 | 0.29 | 3.6 | 0.13 | 4.5e-10 |
| 5kb (3.4%) | 0.17 | 0.030 | 7.2e-09 | 0.34 | 2.9 | 0.16 | 3.3e-09 |
| 100kb (10.6%) | 0.16 | 0.027 | 5.4e-09 | 0.36 | 2.2 | 0.055 | 6.3e-08 |
| PPI-master |  |  |  |  |  |  |  |
| | $\tau^*$ | $se(\tau^*)$ | $p(\tau^*)$ | $p(\Delta\tau^*)$ | $E$ | $se(E)$ | $p(E)$ |
| ABC (0.44%) | 1.4 | 0.16 | 1.1e-18 | 0.36 | 22 | 1.8 | 2.2e-07 |
| ATAC (0.25%) | 1.2 | 0.17 | 8.4e-13 | 0.30 | 23 | 3.4 | 2.1e-06 |
| eQTL (0.34%) | 0.44 | 0.055 | 6.2e-16 | 0.15 | 10 | 1.1 | 8.4e-06 |
| Roadmap (0.49%) | 1.4 | 0.14 | 7.6e-24 | 0.32 | 17 | 1.8 | 2.6e-06 |
| PC-HiC (7.09%) | 0.36 | 0.07 | 1.4e-07 | 0.26 | 3.6 | 0.12 | 7.5e-07 |
| 5kb (4.81%) | 0.37 | 0.032 | 3.2e-32 | 0.40 | 4.4 | 0.12 | 5.3e-09 |
| 100kb (14.3%) | 0.37 | 0.029 | 1.4e-37 | 0.48 | 3.2 | 0.076 | 6.8e-09 |

**Table S20.** S-LDSC results for SNP annotations corresponding to enhancer-related gene scores, using sum across genes of the gene scores linked to a SNP: Standardized Effect sizes (*τ\**) and Enrichment (E) of 70 SNP annotations corresponding to 7 enhancer-related gene scores and 10 S2G strategies, where SNPs linked to multiple genes are weighted by the sum of the gene scores of the genes they are linked to. The results are conditional on 93 baseline-LD+ annotations for enhancer-related annotaions and 113 baseline-LD+cis annotations for candidate master-regulator

| ABC-G |  |  |  |  |  |  |  |
| --- | --- | --- | --- | --- | --- | --- | --- |
| | $\tau^*$ | $se(\tau^*)$ | $p(\tau^*)$ | $p(\Delta\tau^*)$ | $E$ | $se(E)$ | $p(E)$ |
| ABC | 0.43 | 0.13 | 0.00088 | 8e-04 | 26 | 3.7 | 3.4e-05 |
| ATAC | 0.36 | 0.1 | 0.00058 | 0.20 | 15 | 2.6 | 3.7e-06 |
| eQTL | 0.2 | 0.06 | 0.00099 | 0.76 | 7.9 | 1.1 | 2.6e-05 |
| Roadmap | 0.4 | 0.14 | 0.0036 | 0.38 | 19 | 3.1 | 3.9e-06 |
| PC-HiC | 0.16 | 0.045 | 0.00033 | 0.14 | 3.5 | 0.2 | 2.4e-09 |
| 5kb | 0.065 | 0.037 | 0.085 | 0.59 | 3.3 | 0.17 | 2.7e-09 |
| 100kb | 0.081 | 0.024 | 0.00091 | 0.66 | 2.8 | 0.086 | 1.4e-09 |
| ATAC-distal |  |  |  |  |  |  |  |
| | $\tau^*$ | $se(\tau^*)$ | $p(\tau^*)$ | $p(\Delta\tau^*)$ | $E$ | $se(E)$ | $p(E)$ |
| ABC (0.77%) | 0.39 | 0.12 | 0.001 | 0.02 | 21 | 2.6 | 7.8e-06 |
| ATAC (1.01%) | 0.61 | 0.16 | 0.0002 | 0.02 | 11 | 1.7 | 2.3e-07 |
| eQTL (1.05%) | 0.19 | 0.049 | 9.4e-05 | 0.34 | 5.6 | 0.49 | 5.5e-06 |
| Roadmap (1.67%) | 0.31 | 0.18 | 0.084 | 0.03 | 15 | 2.6 | 2.4e-06 |
| PC-HiC (14.4%) | 0.075 | 0.049 | 0.13 | 0.11 | 2.8 | 0.12 | 1.3e-09 |
| 5kb (18.5%) | 0.099 | 0.024 | 4e-05 | 0.24 | 2.4 | 0.064 | 1.1e-09 |
| 100kb (36.4%) | 0.12 | 0.028 | 1.1e-05 | 0.01 | 2.6 | 0.068 | 1.2e-09 |
| EDS-binary |  |  |  |  |  |  |  |
| | $\tau^*$ | $se(\tau^*)$ | $p(\tau^*)$ | $p(\Delta\tau^*)$ | $E$ | $se(E)$ | $p(E)$ |
| ABC | 0.43 | 0.16 | 0.0059 | 0.02 | 27 | 4.8 | 5.8e-05 |
| ATAC | 0.67 | 0.19 | 0.00044 | 0.06 | 19 | 4 | 1.8e-05 |
| eQTL | 0.18 | 0.044 | 5e-05 | 0.32 | 8.1 | 0.77 | 6.2e-06 |
| Roadmap | 0.39 | 0.13 | 0.0035 | 0.07 | 18 | 2.8 | 3.7e-06 |
| PC-HiC | 0.07 | 0.032 | 0.031 | 0.24 | 3.3 | 0.2 | 4.3e-10 |
| 5kb | 0.044 | 0.019 | 0.022 | 0.01 | 2.8 | 0.1 | 8.9e-10 |
| 100kb | 0.15 | 0.032 | 6e-06 | 0.28 | 3.1 | 0.16 | 1.8e-09 |
| eQTL-CTS |  |  |  |  |  |  |  |
| | $\tau^*$ | $se(\tau^*)$ | $p(\tau^*)$ | $p(\Delta\tau^*)$ | $E$ | $se(E)$ | $p(E)$ |
| ABC | 0.15 | 0.12 | 0.24 | 0.56 | 17 | 2.7 | 0.00014 |
| ATAC | 0.14 | 0.084 | 0.096 | 0.48 | 7.6 | 1.2 | 1.5e-05 |
| eQTL | 0.18 | 0.062 | 0.0031 | 0.30 | 4.8 | 0.61 | 4.4e-05 |
| Roadmap | 0.23 | 0.14 | 0.089 | 0.22 | 13 | 2.1 | 1e-05 |
| PC-HiC | -0.041 | 0.036 | 0.25 | 0.32 | 2.5 | 0.099 | 6.3e-08 |
| 5kb | 0.02 | 0.03 | 0.5 | 0.40 | 1.9 | 0.075 | 2.3e-08 |
| 100kb | -0.037 | 0.04 | 0.36 | 0.53 | 1.9 | 0.099 | 1.5e-07 |
| Expecto-MVP |  |  |  |  |  |  |  |
| | $\tau^*$ | $se(\tau^*)$ | $p(\tau^*)$ | $p(\Delta\tau^*)$ | $E$ | $se(E)$ | $p(E)$ |
| ABC | 0.32 | 0.13 | 0.019 | 0.08 | 22 | 4 | 0.00016 |
| ATAC | 0.54 | 0.14 | 0.00016 | 0.12 | 19 | 3.6 | 2.6e-06 |
| eQTL | 0.13 | 0.051 | 0.0089 | 0.26 | 6.7 | 0.84 | 2.8e-05 |
| Roadmap | 0.28 | 0.14 | 0.048 | 0.18 | 17 | 3.7 | 6.3e-05 |
| PC-HiC | 0.011 | 0.05 | 0.82 | 0.36 | 3 | 0.24 | 2.2e-08 |
| 5kb | 0.062 | 0.026 | 0.018 | 0.25 | 2.9 | 0.16 | 2.7e-09 |
| 100kb | 0.063 | 0.032 | 0.052 | 0.28 | 2.7 | 0.14 | 8.5e-11 |
| PC-HiC-distal |  |  |  |  |  |  |  |
| | $\tau^*$ | $se(\tau^*)$ | $p(\tau^*)$ | $p(\Delta\tau^*)$ | $E$ | $se(E)$ | $p(E)$ |
| ABC | -0.037 | 0.11 | 0.74 | 0.22 | 13 | 4.2 | 0.026 |
| ATAC | 0.2 | 0.079 | 0.012 | 0.46 | 12 | 2.2 | 0.00019 |
| eQTL | 0.042 | 0.044 | 0.34 | 0.58 | 4.8 | 0.67 | 6.8e-05 |
| Roadmap | 0.29 | 0.091 | 0.0013 | 0.30 | 18 | 3 | 4.8e-05 |
| PC-HiC | -0.055 | 0.031 | 0.078 | 0.26 | 2.7 | 0.13 | 2.7e-08 |
| 5kb | -0.031 | 0.019 | 0.1 | 0.22 | 2.3 | 0.13 | 9.9e-08 |
| 100kb | 0.016 | 0.026 | 0.55 | 0.50 | 2.7 | 0.14 | 1.8e-07 |
| SEG-GTEx |  |  |  |  |  |  |  |
| | $\tau^*$ | $se(\tau^*)$ | $p(\tau^*)$ | $p(\Delta\tau^*)$ | $E$ | $se(E)$ | $p(E)$ |
| ABC (0.39%) | 0.7 | 0.14 | 5.9e-07 | 0.07 | 34 | 4.2 | 1e-05 |
| ATAC (0.26%) | 0.59 | 0.17 | 0.00051 | 0.12 | 21 | 4.3 | 1.1e-05 |
| eQTL (0.27%) | 0.24 | 0.056 | 2.4e-05 | 0.23 | 9.3 | 1.2 | 6.4e-06 |
| Roadmap (0.66%) | 0.55 | 0.12 | 1.1e-05 | 0.24 | 21 | 2.7 | 3.2e-06 |
| PC-HiC (5.58%) | 0.039 | 0.03 | 0.2 | 0.48 | 3.5 | 0.25 | 7.1e-09 |
| 5kb (3.54%) | 0.12 | 0.026 | 1.8e-06 | 0.08 | 3.6 | 0.18 | 4.1e-08 |
| 100kb (13.4%) | 0.079 | 0.022 | 0.00026 | 0.02 | 2.6 | 0.11 | 5.1e-08 |
| PPI-enhancer |  |  |  |  |  |  |  |
| | $\tau^*$ | $se(\tau^*)$ | $p(\tau^*)$ | $p(\Delta\tau^*)$ | $E$ | $se(E)$ | $p(E)$ |
| ABC | 1.0 | 0.2 | 9e-07 | 0.008 | 39 | 5.7 | 1.3e-07 |
| ATAC | 1.2 | 0.23 | 1.5e-07 | 0.12 | 25 | 4 | 3.7e-09 |
| eQTL | 0.41 | 0.088 | 3.6e-06 | 0.28 | 10 | 1.3 | 6.2e-0 |
| Roadmap | 0.99 | 0.19 | 1.2e-07 | 0.08 | 28 | 3.9 | 1.5e-07 |
| PC-HiC | 0.25 | 0.056 | 7.9e-06 | 0.23 | 3.6 | 0.23 | 6.5e-11 |
| 5kb | 0.2 | 0.023 | 6.7e-18 | 0.27 | 3.2 | 0.12 | 2.7e-10 |
| 100kb | 0.27 | 0.047 | 6.2e-09 | 0.32 | 3.4 | 0.19 | 1.4e-10 |
| Trans-master |  |  |  |  |  |  |  |
| | $\tau^*$ | $se(\tau^*)$ | $p(\tau^*)$ | $p(\Delta\tau^*)$ | $E$ | $se(E)$ | $p(E)$ |
| ABC (0.33%) | 0.9 | 0.21 | 2.7e-05 | 0.04 | 60 | 8.1 | 1.8e-05 |
| ATAC | 1.1 | 0.22 | 2.8e-07 | 0.08 | 36 | 5.7 | 1.5e-06 |
| eQTL | 0.54 | 0.11 | 1.7e-06 | 0.26 | 12 | 1.8 | 9.2e-07 |
| Roadmap | 1.7 | 0.2 | 1.1e-17 | 0.002 | 57 | 5.3 | 8.3e-07 |
| PC-HiC | 0.42 | 0.051 | 2.4e-16 | 0.40 | 6.1 | 0.27 | 9.4e-08 |
| 5kb | 0.3 | 0.066 | 4.3e-06 | 0.34 | 5.9 | 0.47 | 2.2e-08 |
| 100kb | 0.46 | 0.097 | 2.5e-06 | 0.10 | 6 | 0.69 | 7.1e-06 |
| TF |  |  |  |  |  |  |  |
| | $\tau^*$ | $se(\tau^*)$ | $p(\tau^*)$ | $p(\Delta\tau^*)$ | $E$ | $se(E)$ | $p(E)$ |
| ABC (0.32%) | 0.19 | 0.11 | 0.088 | 0.37 | 22 | 3.9 | 0.00075 |
| ATAC | 0.53 | 0.083 | 2.3e-10 | 0.10 | 22 | 2.9 | 1.1e-05 |
| eQTL | 0.0033 | 0.036 | 0.93 | 0.66 | 3.6 | 0.95 | 0.036 |
| Roadmap | 0.75 | 0.12 | 5.4e-10 | 0.27 | 37 | 4.8 | 2.3e-0 |
| PC-HiC | -0.027 | 0.038 | 0.47 | 0.44 | 2.4 | 0.41 | 0.012 |
| 5kb | 0.12 | 0.024 | 1.8e-06 | 0.11 | 2.7 | 0.16 | 5.7e-07 |
| 100kb | -0.0061 | 0.019 | 0.74 | 0.001 | 1.6 | 0.16 | 0.015 |
| PPI-master |  |  |  |  |  |  |  |
| | $\tau^*$ | $se(\tau^*)$ | $p(\tau^*)$ | $p(\Delta\tau^*)$ | $E$ | $se(E)$ | $p(E)$ |
| ABC | 1.2 | 0.19 | 7.3e-10 | 0.13 | 58 | 6.2 | 7.6e-06 |
| ATAC | 1.4 | 0.2 | 9.5e-13 | 0.16 | 37 | 4.5 | 1.2e-07 |
| eQTL | 0.46 | 0.06 | 9e-15 | 0.22 | 13 | 1.2 | 1.6e-06 |
| Roadmap | 1.7 | 0.18 | 5.4e-21 | 0.32 | 57 | 5.5 | 4.6e-07 |
| PC-HiC | 0.3 | 0.052 | 1e-08 | 0.58 | 4.7 | 0.26 | 1.1e-07 |
| 5kb | 0.3 | 0.052 | 1e-08 | 0.34 | 4.7 | 0.26 | 1.1e-07 |
| 100kb | 0.21 | 0.055 | 0.00013 | 0.36 | 4.3 | 0.33 | 2e-07 |

**Table S21.**
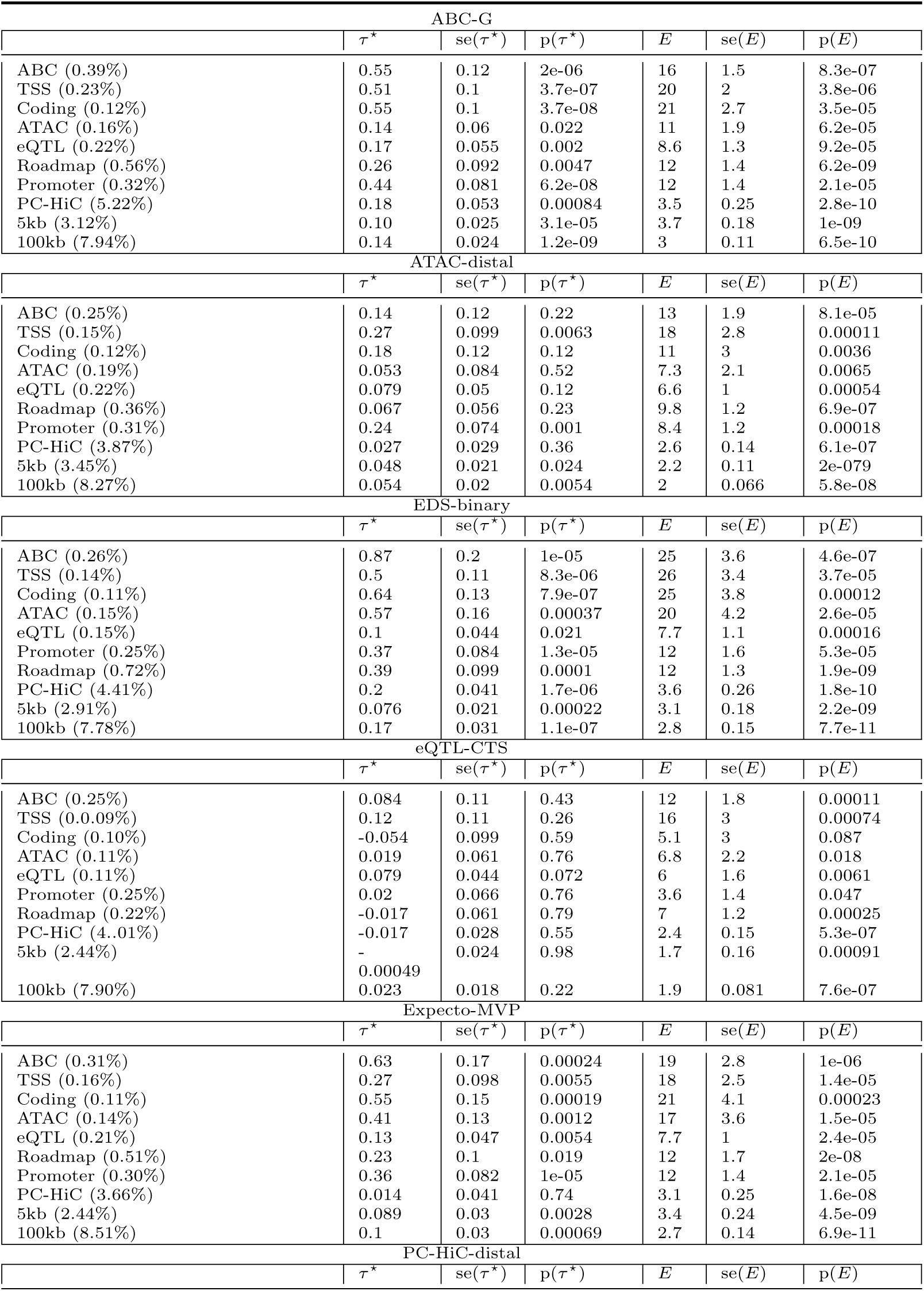
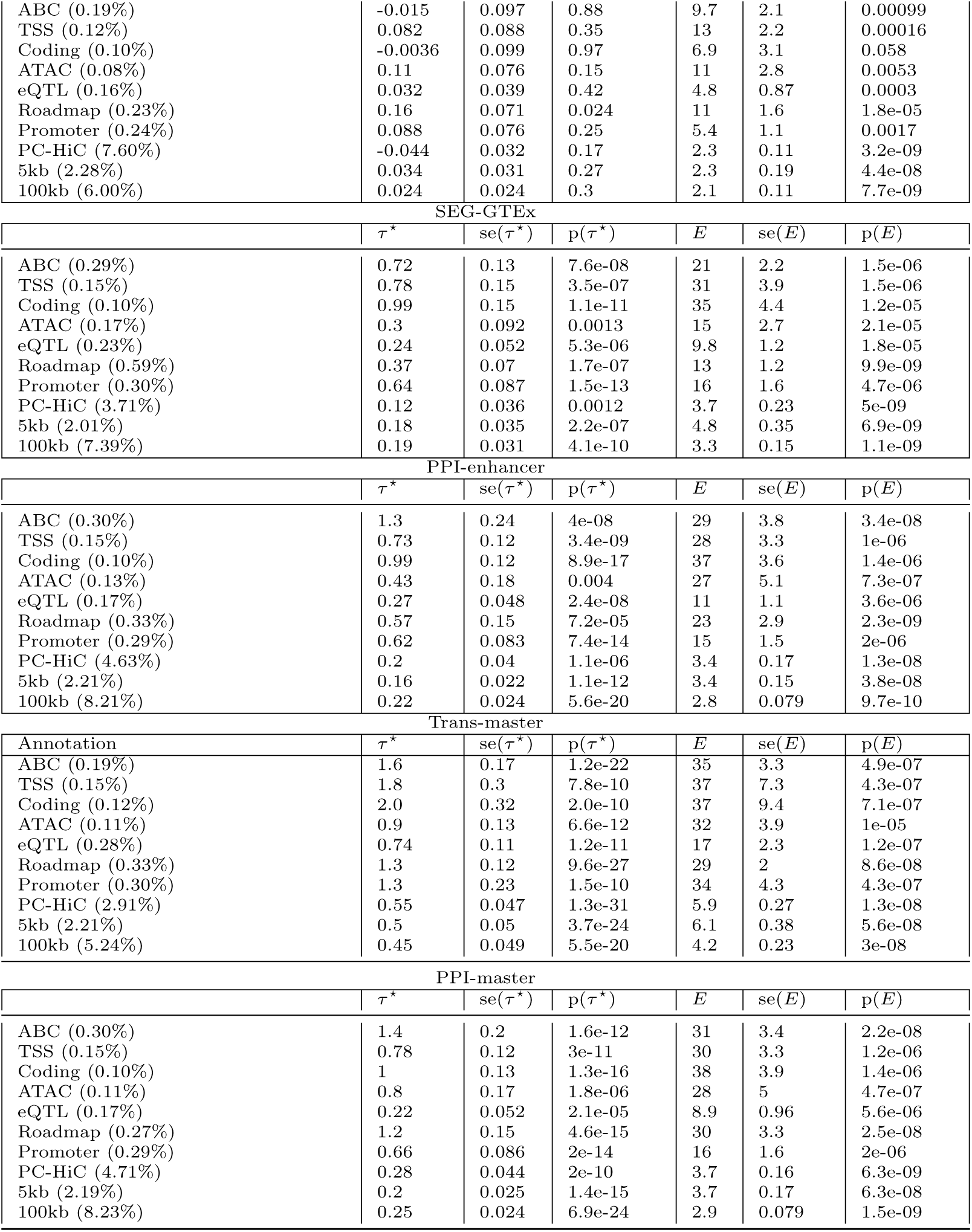
S-LDSC results for SNP annotations corresponding to all gene scores based on top 5% of genes (instead of top 10%): Standardized Effect sizes (*τ\**) and Enrichment (E) of 80 SNP annotations corresponding to 8 gene scores with 10% genes down-sampled to top 5% genes and 10 S2G strategies, conditional on 93 baseline-LD+ annotations for enhancer-related or PPI-enhancer gene scores and 113 baseline-LD+cis annotations for PPI-master. Reports are meta-analyzed across 11 blood-related traits.

**Table S22.**
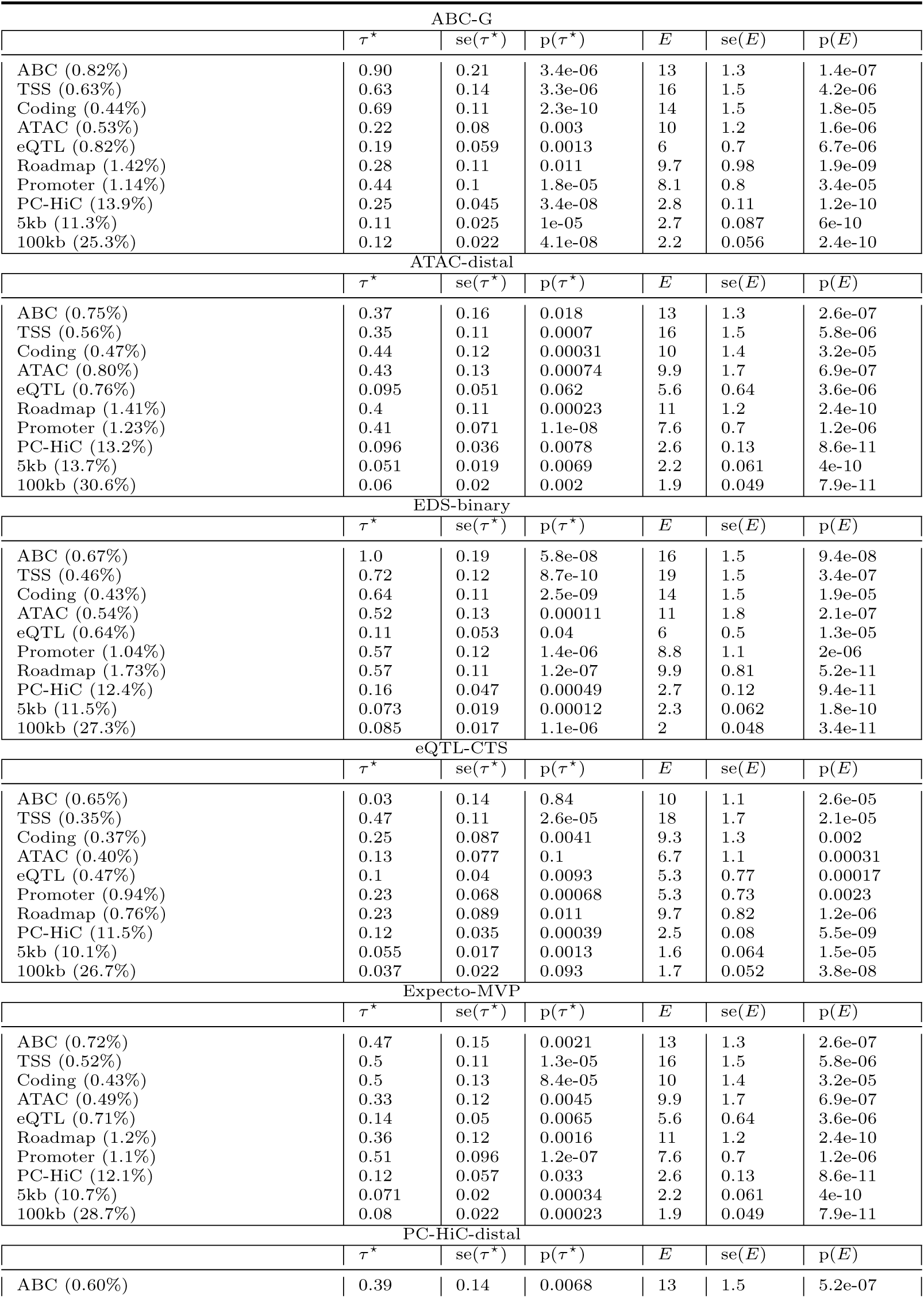
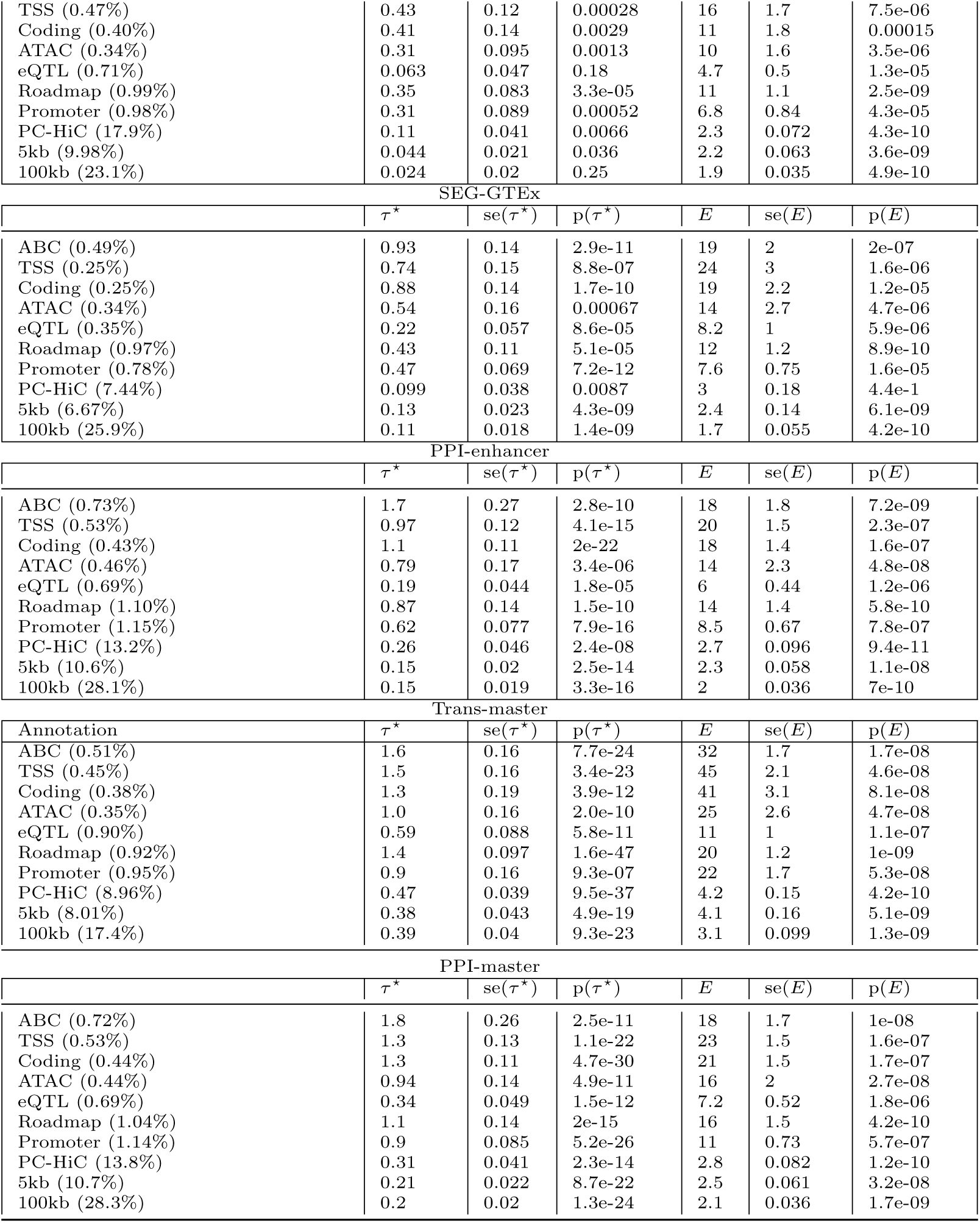
S-LDSC results for SNP annotations corresponding to all gene scores based on top 20% of genes (instead of top 10%): Standardized Effect sizes (*τ\**) and Enrichment (E) of 80 SNP annotations corresponding to 8 gene scores with 10% genes up-sampled to top 20% genes and 10 S2G strategies, conditional on 93 baseline-LD+ annotations for enhancer-related or PPI-enhancer gene scores and 113 baseline-LD+cis annotations for PPI-master. Reports are meta-analyzed across 11 blood-related traits.

**Table S23.** S-LDSC results for SNP annotations corresponding to the PPI-enhancer gene score conditional on the baseline-LD+ model: Standardized Effect sizes (*τ\**) and Enrichment (E) of 10 SNP annotations corresponding to the PPI-enhancer gene score, conditional on 93 baseline-LD+ annotations. Reports are meta-analyzed across 11 blood-related traits.

| PPI-enhancer |  |  |  |  |  |  |
| --- | --- | --- | --- | --- | --- | --- |
| | $\tau^*$ | $\text{se}(\tau^*)$ | $\text{p}(\tau^*)$ | $E$ | $\text{se}(E)$ | $\text{p}(E)$ |
| ABC (0.58%) | 2.0 | 0.29 | 8.2e-13 | 24 | 2.6 | 3.4e-09 |
| TSS (0.33%) | 1.2 | 0.17 | 2.6e-13 | 28 | 2.8 | 3.3e-08 |
| Coding (0.24%) | 1.4 | 0.18 | 4.8e-15 | 29 | 3 | 2.6e-08 |
| ATAC (0.41%) | 1.2 | 0.21 | 2.9e-08 | 20 | 2.9 | 2.9e-09 |
| eQTL (0.38%) | 0.39 | 0.085 | 4e-06 | 9.7 | 1.2 | 6.8e-08 |
| Roadmap (1.05%) | 1.2 | 0.18 | 3.4e-11 | 15 | 1.6 | 9e-12 |
| Promoter (0.64%) | 1.1 | 0.14 | 1e-14 | 15 | 1.3 | 2.1e-08 |
| PC-HiC (8.66%) | 0.32 | 0.056 | 1.1e-08 | 3.3 | 0.18 | 7.3e-11 |
| 5kb (6.45%) | 0.21 | 0.022 | 4.3e-21 | 3.1 | 0.11 | 4.3e-10 |
| 100kb (17.3%) | 0.31 | 0.026 | 5.1e-32 | 2.7 | 0.087 | 8.3e-11 |

**Table S24.**
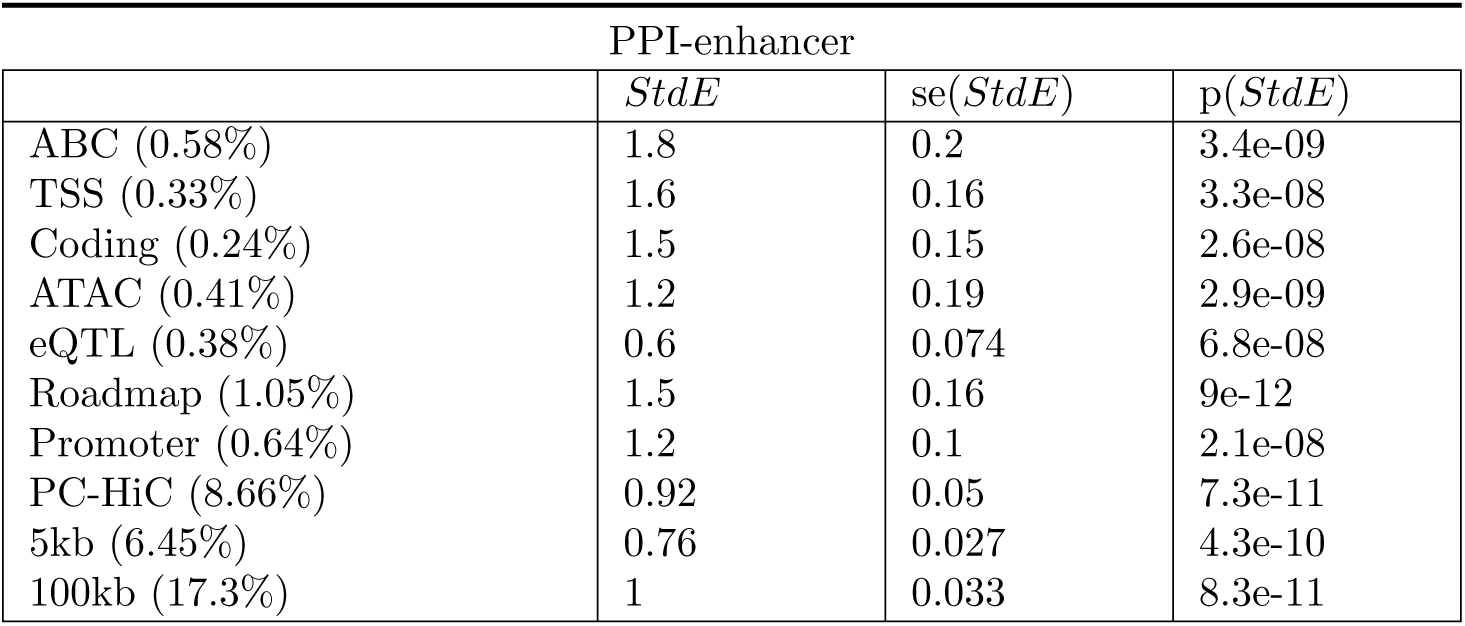
Standardized enrichment of SNP annotations corresponding to PPI-enhancer gene score conditional on the baseline-LD+ model: Standardized enrichment of the 10 SNP annotations corresponding to the PPI-enhancer gene score, conditional on 93 baseline-LD+ annotations respectively. Reports are meta-analyzed across 11 blood-related traits.

| PPI-enhancer |  |  |  |
| --- | --- | --- | --- |
| | $StdE$ | $se(StdE)$ | $p(StdE)$ |
| ABC (0.58%) | 1.8 | 0.2 | 3.4e-09 |
| TSS (0.33%) | 1.6 | 0.16 | 3.3e-08 |
| Coding (0.24%) | 1.5 | 0.15 | 2.6e-08 |
| ATAC (0.41%) | 1.2 | 0.19 | 2.9e-09 |
| eQTL (0.38%) | 0.6 | 0.074 | 6.8e-08 |
| Roadmap (1.05%) | 1.5 | 0.16 | 9e-12 |
| Promoter (0.64%) | 1.2 | 0.1 | 2.1e-08 |
| PC-HiC (8.66%) | 0.92 | 0.05 | 7.3e-11 |
| 5kb (6.45%) | 0.76 | 0.027 | 4.3e-10 |
| 100kb (17.3%) | 1 | 0.033 | 8.3e-11 |

**Table S25.** S-LDSC results for SNP annotations corresponding to the PPI-enhancer gene score conditional on the enhancer-related joint model: Standardized Effect sizes (*τ\**) and Enrichment (E) of 10 SNP annotations corresponding to the PPI-enhancer gene score, conditional on the enhancer-related joint model from Table S16. Reports are meta-analyzed across 11 blood-related traits.

| PPI-enhancer |  |  |  |  |  |  |
| --- | --- | --- | --- | --- | --- | --- |
| | $\tau^*$ | $se(\tau^*)$ | $p(\tau^*)$ | $E$ | $se(E)$ | $p(E)$ |
| ABC (0.58%) | 1.8 | 0.3 | 3.5e-09 | 21 | 2.5 | 8.4e-09 |
| TSS (0.33%) | 0.85 | 0.15 | 4.6e-08 | 24 | 2.6 | 9.9e-08 |
| Coding (0.24%) | 0.93 | 0.16 | 3.1e-09 | 25 | 2.9 | 1.2e-07 |
| ATAC (0.41%) | 0.96 | 0.2 | 1.8e-06 | 17 | 2.8 | 3.6e-08 |
| eQTL (0.38%) | 0.29 | 0.08 | 0.00024 | 8.7 | 1.2 | 2.9e-07 |
| Roadmap (1.05%) | 1 | 0.18 | 4.5e-08 | 13 | 1.5 | 4.9e-11 |
| Promoter (0.64%) | 0.73 | 0.14 | 2e-07 | 13 | 1.3 | 7.7e-08 |
| PC-HiC (8.66%) | 0.24 | 0.053 | 4.5e-06 | 3.1 | 0.17 | 8e-11 |
| 5kb (6.45%) | 0.13 | 0.023 | 1.6e-08 | 3.2 | 0.11 | 4e-10 |
| 100kb (17.3%) | 0.23 | 0.024 | 9.8e-23 | 2.7 | 0.087 | 8.6e-11 |

**Table S26.** S-LDSC results for the joint analalysis of SNP annotations corresponding to the enhancer-related and PPI-enhancer gene scores. Standardized Effect sizes (*τ\**) and Enrichment (E) of the significant SNP annotations in a joint model comprising of marginally significant enhancer-related SNP annotations and PPI-enhancer SNP annotations and from Figure 3. We report results on the annotations that are significant in the joint model. Also marked in red are annotations that ere jointly Bonferroni significant in the enhancer-related joint model (Supplementary Figure S11) but not Bonferroni significant in this PPI-enhancer-related joint model. All analysis are conditional on 93 baseline-LD+ annotations. Results are meta-analyzed across 11 blood-related traits.

| Annotation | $\tau^*$ | se($\tau^*$) | p($\tau^*$) | $E$ | se ($E$) | p($E$) |
| --- | --- | --- | --- | --- | --- | --- |
| ATAC-distal × Promoter<br>(1.8%) | 0.43 | 0.071 | 1.1e-09 | 6.7 | 0.51 | 1.6e-05 |
| EDS-binary × 100kb<br>(14.6%) | 0.10 | 0.02 | 8.8e-07 | 2.5 | 0.089 | 6.7e-11 |
| SEG-GTEx × Coding<br>(0.17%) | 0.62 | 0.15 | 4.3e-05 | 22 | 3.2 | 4.6e-05 |
| PPI-enhancer × ABC<br>(0.58%) | 1.2 | 0.21 | 3.5e-09 | 18 | 2 | 1.2e-07 |
| PPI-enhancer × TSS<br>(0.33%) | 0.38 | 0.1 | 0.00017 | 19 | 1.8 | 3.6e-06 |
| PPI-enhancer × Coding<br>(0.24%) | 0.38 | 0.097 | 7.2e-05 | 18 | 1.9 | 1.3e-05 |
| PPI-enhancer × ATAC<br>(0.41%) | 0.6 | 0.17 | 0.00038 | 14 | 2.6 | 8.8e-07 |
| ABC-G × TSS<br>(0.38%) | 0.34 | 0.12 | 0.0054 | 19 | 1.8 | 2.9e-06 |
| EDS-binary × Roadmap<br>(1.15%) | 0.066 | 0.082 | 0.42 | 7.1 | 0.9 | 2.5e-05 |
| SEG-GTEx × ABC<br>(0.40%) | 0.17 | 0.15 | 0.26 | 16 | 1.8 | 2.4e-06 |

**Table S27.** S-LDSC results for SNP annotations corresponding to the Weighted Enhancer gene score conditional on the baseline-LD+ model: Standardized Effect sizes (*τ\**) and Enrichment (E) of 10 SNP annotations corresponding to the Weighted Enhancer gene score. The analysis is conditional on 93 baseline-LD+ annotations. Reports are meta-analyzed across 11 blood-related traits.

| Annotation | $\tau^*$ | $se(\tau^*)$ | $p(\tau^*)$ | $E$ | $se(E)$ | $p(E)$ |
| --- | --- | --- | --- | --- | --- | --- |
| ABC (0.58%) | 1 | 0.13 | 3.2e-14 | 17 | 1.3 | 8.2e-08 |
| TSS (0.36%) | 0.91 | 0.11 | 6.8e-16 | 22 | 1.7 | 1.1e-06 |
| Coding (0.25%) | 0.98 | 0.1 | 1.4e-20 | 23 | 2 | 1.5e-06 |
| ATAC (0.44%) | 0.64 | 0.15 | 2.2e-05 | 14 | 2.3 | 1.5e-07 |
| eQTL (0.44%) | 0.29 | 0.057 | 3.7e-07 | 8.6 | 0.9 | 2.8e-06 |
| Roadmap (1.31%) | 0.81 | 0.12 | 3.3e-11 | 12 | 1.1 | 4.3e-11 |
| Promoter (0.64%) | 0.86 | 0.087 | 4.4e-23 | 14 | 1.1 | 7.1e-07 |
| PC-HiC (9.26%) | 0.17 | 0.044 | 0.0001 | 3.1 | 0.17 | 1e-10 |
| 5kb (6.22%) | 0.15 | 0.025 | 2.2e-09 | 3.4 | 0.14 | 2.1e-10 |
| 100kb (16.8%) | 0.23 | 0.023 | 2.3e-23 | 2.8 | 0.088 | 1.2e-10 |

**Table S28.** S-LDSC results for SNP annotations corresponding to the Weighted Enhancer gene score conditional on the PPI-enhancer-related joint model: Standardized Effect sizes (*τ\**) and Enrichment (E) of SNP annotations corresponding to the Weighted Enhancer gene score. The analysis is conditional on 93 baseline-LD+ annotations and 7 annotations from the PPI-enhancer-related joint model in Table S26. Reports are meta-analyzed across 11 blood-related traits.

| Annotation | $\tau^*$ | $se(\tau^*)$ | $p(\tau^*)$ | $E$ | $se(E)$ | $p(E)$ |
| --- | --- | --- | --- | --- | --- | --- |
| ABC (0.58%) | -0.11 | 0.18 | 0.54 | 14 | 1.3 | 1.1e-06 |
| TSS (0.36%) | 0.2 | 0.14 | 0.13 | 19 | 1.7 | 8.7e-06 |
| Coding (0.25%) | 0.092 | 0.15 | 0.55 | 17 | 1.9 | 3.7e-05 |
| ATAC (0.44%) | -0.083 | 0.083 | 0.32 | 11 | 2.1 | 2.6e-05 |
| eQTL (0.44%) | 0.11 | 0.055 | 0.04 | 7.1 | 0.89 | 1.8e-05 |
| Roadmap (1.31%) | 0.25 | 0.083 | 0.0021 | 9 | 0.97 | 7.3e-10 |
| Promoter (0.64%) | 0.23 | 0.092 | 0.014 | 11 | 1 | 1.8e-05 |
| PC-HiC (9.26%) | 0.014 | 0.033 | 0.68 | 2.9 | 0.16 | 1.4e-10 |
| 5kb (6.22%) | -0.036 | 0.028 | 0.2 | 3.5 | 0.14 | 2.8e-10 |
| 100kb (16.8%) | 0.051 | 0.028 | 0.075 | 2.8 | 0.089 | 1.2e-10 |

**Table S29.** S-LDSC results for SNP annotations corresponding to the PPI-control gene score conditional on the baseline-LD+ model: Standardized Effect sizes (*τ\**) and Enrichment (E) of 10 SNP annotations corresponding to the PPI-control gene score, conditional on 93 baseline-LD+ annotations. Reports are meta-analyzed across 11 blood-related traits.

| PPI-control-4 |  |  |  |  |  |  |
| --- | --- | --- | --- | --- | --- | --- |
| | $\tau^*$ | $\text{se}(\tau^*)$ | $\text{p}(\tau^*)$ | $E$ | $\text{se}(E)$ | $\text{p}(E)$ |
| ABC (0.46%) | 0.59 | 0.13 | 3e-06 | 16 | 1.4 | 7.9e-07 |
| TSS (0.25%) | 0.6 | 0.11 | 1.3e-08 | 21 | 2 | 6.4e-06 |
| Coding (0.23%) | 0.39 | 0.093 | 3.2e-05 | 13 | 1.8 | 0.00024 |
| ATAC (0.24%) | 0.25 | 0.077 | 0.0013 | 9.6 | 1.6 | 0.00013 |
| eQTL (0.31%) | 0.17 | 0.043 | 5e-05 | 6.9 | 0.78 | 2.8e-05 |
| Roadmap (0.49%) | 0.32 | 0.075 | 2.3e-05 | 12 | 0.97 | 6.4e-07 |
| Promoter (0.59%) | 0.44 | 0.069 | 1.2e-10 | 8.7 | 0.88 | 2.7e-05 |
| PC-HiC (7.55%) | 0.098 | 0.033 | 0.0027 | 2.7 | 0.098 | 6.5e-08 |
| 5kb (5.86%) | 0.089 | 0.018 | 8.2e-07 | 2 | 0.073 | 3.8e-07 |
| 100kb (16.7%) | 0.11 | 0.018 | 6.8e-09 | 2 | 0.058 | 9e-09 |

**Table S30.** S-LDSC results for SNP annotations corresponding to the PPI-control gene score conditional on the PPI-enhancer-related joint model: Standardized Effect sizes (*τ\**) and Enrichment (E) of 10 SNP annotations corresponding to the PPI-control gene score, conditional on the PPI-enhancer-related joint model from Table S26. Reports are meta-analyzed across 11 blood-related traits.

| PPI-control-4 |  |  |  |  |  |  |
| --- | --- | --- | --- | --- | --- | --- |
| | $\tau^*$ | $\text{se}(\tau^*)$ | $\text{p}(\tau^*)$ | $E$ | $\text{se}(E)$ | $\text{p}(E)$ |
| ABC (0.46%) | 0.27 | 0.16 | 0.1 | 15 | 1.4 | 1.3e-06 |
| TSS (0.25%) | 0.27 | 0.11 | 0.014 | 18 | 1.9 | 2.2e-05 |
| Coding (0.23%) | 0.015 | 0.14 | 0.92 | 10 | 2 | 0.0032 |
| ATAC (0.24%) | 0.05 | 0.072 | 0.49 | 8.3 | 1.8 | 0.00049 |
| eQTL (0.31%) | 0.076 | 0.045 | 0.087 | 5.8 | 0.88 | 0.00019 |
| Roadmap (0.49%) | 0.25 | 0.077 | 0.0014 | 10 | 1 | 4.3e-06 |
| Promoter (0.59%) | 0.14 | 0.07 | 0.039 | 7 | 0.87 | 0.0002 |
| PC-HiC (7.55%) | 0.09 | 0.033 | 0.0071 | 2.7 | 0.1 | 3.1e-08 |
| 5kb (5.86%) | 0.024 | 0.02 | 0.23 | 2.1 | 0.092 | 6.5e-07 |
| 100kb (16.7%) | 0.07 | 0.027 | 0.01 | 2.1 | 0.071 | 3.2e-09 |

**Table S31.** S-LDSC results for SNP annotations corresponding to the RegNet-Enhancer gene score conditional on the baseline-LD+ model: Standardized Effect sizes (*τ\**) and Enrichment (E) of SNP annotations corresponding to the RegNet-Enhancer gene score. The analysis is conditional on 93 baseline-LD+ annotations. Reports are meta-analyzed across 11 blood-related traits.

| Annotation | $\tau^*$ | $se(\tau^*)$ | $p(\tau^*)$ | $E$ | $se(E)$ | $p(E)$ |
| --- | --- | --- | --- | --- | --- | --- |
| ABC (0.57%) | 1.3 | 0.23 | 5.9e-09 | 19 | 2.2 | 3e-08 |
| TSS (0.36%) | 0.9 | 0.15 | 1.9e-09 | 23 | 2.5 | 4.6e-08 |
| Coding (0.26%) | 0.97 | 0.15 | 1e-10 | 21 | 2.4 | 5.2e-07 |
| ATAC (0.36%) | 0.79 | 0.17 | 4.8e-06 | 17 | 2.7 | 8.1e-08 |
| eQTL (0.39%) | 0.23 | 0.05 | 4.7e-06 | 8.1 | 0.76 | 3.5e-06 |
| Roadmap (1.10%) | 0.8 | 0.18 | 5.3e-06 | 13 | 1.5 | 3.9e-11 |
| Promoter (0.68%) | 0.73 | 0.087 | 4e-17 | 12 | 0.98 | 2.5e-07 |
| PC-HiC (9.49%) | 0.33 | 0.05 | 1.8e-11 | 3.4 | 0.17 | 8.6e-11 |
| 5kb (6.63%) | 0.16 | 0.023 | 2e-11 | 3.1 | 0.11 | 4.1e-10 |
| 100kb (18.0%) | 0.22 | 0.025 | 1.8e-19 | 2.6 | 0.083 | 6.9e-11 |

**Table S32.**
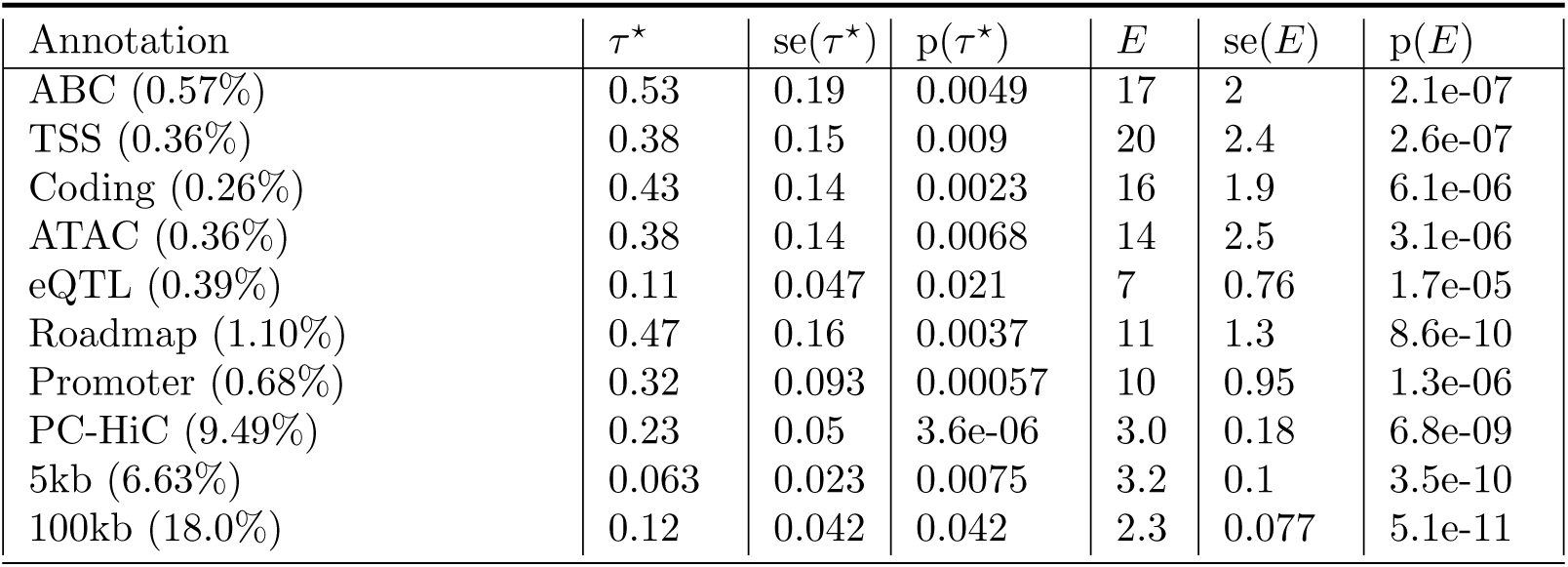
S-LDSC results for SNP annotations corresponding to the RegNet-Enhancer gene score conditional on the PPI-enhancer-related joint model: Standardized Effect sizes (*τ\**) and Enrichment (E) of SNP annotations corresponding to the RegNet-Enhancer gene score. The analysis is conditional on 93 baseline-LD+ annotations and 7 annotations from the PPI-enhancer-related joint model in Table S26. Reports are meta-analyzed across 11 blood-related traits.

| Annotation | $\tau^*$ | $se(\tau^*)$ | $p(\tau^*)$ | $E$ | $se(E)$ | $p(E)$ |
| --- | --- | --- | --- | --- | --- | --- |
| ABC (0.57%) | 0.53 | 0.19 | 0.0049 | 17 | 2 | 2.1e-07 |
| TSS (0.36%) | 0.38 | 0.15 | 0.009 | 20 | 2.4 | 2.6e-07 |
| Coding (0.26%) | 0.43 | 0.14 | 0.0023 | 16 | 1.9 | 6.1e-06 |
| ATAC (0.36%) | 0.38 | 0.14 | 0.0068 | 14 | 2.5 | 3.1e-06 |
| eQTL (0.39%) | 0.11 | 0.047 | 0.021 | 7 | 0.76 | 1.7e-05 |
| Roadmap (1.10%) | 0.47 | 0.16 | 0.0037 | 11 | 1.3 | 8.6e-10 |
| Promoter (0.68%) | 0.32 | 0.093 | 0.00057 | 10 | 0.95 | 1.3e-06 |
| PC-HiC (9.49%) | 0.23 | 0.05 | 3.6e-06 | 3.0 | 0.18 | 6.8e-09 |
| 5kb (6.63%) | 0.063 | 0.023 | 0.0075 | 3.2 | 0.1 | 3.5e-10 |
| 100kb (18.0%) | 0.12 | 0.042 | 0.042 | 2.3 | 0.077 | 5.1e-11 |

**Table S33.** List of 97 known master-regulator genes from 18 master-regulator families: We list 97 genes spanning 18 master-regulator families curated from existing literature^58–62^ that we use as a validation of the candidate master-regulator gene scores.

| Family | Genes |
| --- | --- |
| CDX | *CDX1, CDX2, CDX4* |
| CDK | *CDK1, CDK2* |
| HSF | *HSF1, HSF2, HSFA1* |
| MTA | *MTA1, MTA2, MTA3, MTAP* |
| SREBP | *SREBF1, SREBF2* |
| HNF | *HNF1A, HNF4A, HNF4G* |
| IL | *IL-1, IL-2, IL-6, IL-7, IL-10, IL-12, IL-21, IL-23, IL-27, ILC3* |
| PLK | *PLK1, PLK2, PLK3, PLK4, PLK5* |
| PAX | *PAX1, PAX2, PAX3, PAX4, PAX5, PAX6, PAX7, PAX8, PAX9* |
| TBX | *TBX1, TBX5, TBX21* |
| SOX | *SOX2, SOX3, SOX4, SOX5, SOX6, SOX9, SOXB1, SOXE, SOXF* |
| RUNX | *RUNX1, RUNX2* |
| IRF | *IRF1, IRF4, IRF7, IRF8* |
| BCL | *BCL2, BCL6, BCL11B, BCL2L1, BCL2L2, BCL2L10, BCL2L11, BCL2L12, BCL2L13, BCL2L14, BCL2L15, BCL2A1* |
| CEBP | *CEBPE, CEBPZ, CEBPG, CEBPB, CEBPD, CEBPA* |
| HLA | *HLA-A, HLA-B, HLA-C, HLA-E, HLA-G, HLA-F* |
| GATA | *GATA1, GATA2, GATA3, GATA4, GATA5, GATA6* |
| NFKB | *NFKB1, NFKB2, NFKBIA, NFKBIB, NFKBIZ, NFKBIE, NFKBID, NFKBIL1* |

**Table S34.** Excess overlap of candidate master-regulator gene scores with 97 known master-regulator genes: Excess overlap (with p-value) of the Trans-master, TF and PPI-master gene scores with respect to 97 genes from 18 master-regulator families are curated from known literature^58–62^ (see Table S33. For the control, we averaged over 10 different control gene scores as reported in Table S5.

| Gene Scores | ENR | p(ENR) |
| --- | --- | --- |
| Trans-master | 3.49 | 5.6e-72 |
| TF | 5.36 | 2.2e-160 |
| PPI-master | 7.22 | 2.3e-214 |

**Table S35.** S-LDSC results for Trans-master and Transcription Factor (TF) SNP annotations conditional on the baseline-LD+cis model: Standardized Effect sizes (*τ\**) and Enrichment (E) of 20 SNP annotations corresponding to Trans-master and Transcription Factor genes. The analysis is conditional on 113 baseline-LD+cis model annotations. Reports are meta-analyzed across 11 blood-related traits.

| Trans-master |  |  |  |  |  |  |
| --- | --- | --- | --- | --- | --- | --- |
| Annotation | $\tau^*$ | $se(\tau^*)$ | $p(\tau^*)$ | $E$ | $se(E)$ | $p(E)$ |
| ABC (0.33%) | 1.5 | 0.25 | 8e-10 | 32 | 2.5 | 8e-07 |
| TSS (0.27%) | 1.6 | 0.25 | 2.2e-10 | 44 | 3.3 | 5.7e-07 |
| Coding (0.22%) | 1.6 | 0.28 | 1.2e-08 | 32 | 4.9 | 1.8e-05 |
| ATAC (0.21%) | 0.9 | 0.17 | 6.4e-08 | 24 | 3.7 | 2.1e-06 |
| eQTL (0.54%) | 0.58 | 0.1 | 5.2e-09 | 11 | 1.5 | 1.7e-06 |
| Roadmap (0.58%) | 1.3 | 0.12 | 1.3e-25 | 22 | 1.4 | 3.6e-08 |
| Promoter (0.57%) | 1.1 | 0.18 | 1.9e-09 | 21 | 2.2 | 1.7e-06 |
| PC-HiC (5.1%) | 0.48 | 0.043 | 1.1e-29 | 5 | 0.18 | 5.3e-09 |
| 5kb (4.2%) | 0.39 | 0.053 | 1.3e-13 | 5.3 | 0.29 | 1.9e-08 |
| 100kb (9.7%) | 0.37 | 0.046 | 3.1e-16 | 3.8 | 0.17 | 8.7e-09 |

| TF |  |  |  |  |  |  |
| --- | --- | --- | --- | --- | --- | --- |
| Annotation | $\tau^*$ | $se(\tau^*)$ | $p(\tau^*)$ | $E$ | $se(E)$ | $p(E)$ |
| ABC (0.32%) | 0.48 | 0.13 | 0.00022 | 17 | 1.6 | 1.2e-06 |
| TSS (0.16%) | 0.88 | 0.18 | 1.2e-06 | 30 | 4.3 | 8.2e-07 |
| Coding (0.16%) | 0.32 | 0.086 | 0.00022 | 12 | 1.8 | 8.1e-05 |
| ATAC (0.14%) | 0.68 | 0.11 | 1.8e-10 | 23 | 3.1 | 8.3e-07 |
| eQTL (0.17%) | 0.03 | 0.037 | 0.41 | 4.1 | 0.87 | 0.0009 |
| Roadmap (0.30%) | 0.8 | 0.12 | 8.6e-12 | 21 | 2.3 | 1.1e-07 |
| Promoter (0.37%) | 0.31 | 0.08 | 0.00012 | 9.1 | 1.1 | 9e-05 |
| PC-HiC (4.9%) | 0.22 | 0.037 | 2.5e-09 | 3.3 | 0.14 | 1.5e-08 |
| 5kb (3.4%) | 0.19 | 0.031 | 2.9e-10 | 2.8 | 0.16 | 2.7e-08 |
| 100kb (10.6%) | 0.16 | 0.027 | 1.5e-09 | 2.3 | 0.055 | 1.2e-08 |

**Table S36.** Standardized Enrichment S-LDSC results for SNP annotations generated from Trans-master and Transcription Factor gene scores conditional on the baseline-LD+cis model.: Standardized Enrichment (StdE) of 20 SNP annotations corresponding to Trans-master and Transcription Factor genes. The analysis is conditional on the 113 baseline-LD+cis (93 baseline-LD+ and 10 Cis1 and 10 Cis3LD) annotations. Reports are meta-analyzed across 11 blood-related traits.

| Trans-master |  |  |  |
| --- | --- | --- | --- |
| Annotation | $StdE$ | $se(StdE)$ | $p(StdE)$ |
| ABC (0.33%) | 1.8 | 0.14 | 8e-07 |
| TSS (0.27%) | 2.3 | 0.17 | 5.7e-07 |
| Coding (0.22%) | 1.5 | 0.23 | 1.8e-05 |
| ATAC (0.21%) | 1.1 | 0.17 | 2.1e-06 |
| eQTL (0.54%) | 0.8 | 0.11 | 1.7e-06 |
| Roadmap (0.58%) | 1.7 | 0.11 | 3.6e-08 |
| Promoter (0.57%) | 1.6 | 0.16 | 1.7e-06 |
| PC-HiC (5.1%) | 1.1 | 0.039 | 5.3e-09 |
| 5kb (4.2%) | 1.1 | 0.059 | 1.9e-08 |
| 100kb (9.7%) | 1.1 | 0.049 | 8.7e-09 |

| TF |  |  |  |
| --- | --- | --- | --- |
| | $StdE$ | $se(StdE)$ | $p(StdE)$ |
| ABC (0.32%) | 0.97 | 0.09 | 1.2e-06 |
| TSS (0.16%) | 1.2 | 0.18 | 8.2e-07 |
| Coding (0.16%) | 0.49 | 0.073 | 8.1e-05 |
| ATAC (0.14%) | 0.84 | 0.11 | 8.3e-07 |
| eQTL (0.17%) | 0.17 | 0.035 | 0.0099 |
| Roadmap (0.30%) | 1.2 | 0.13 | 1.1e-07 |
| Promoter (0.37%) | 0.55 | 0.067 | 9e-05 |
| PC-HiC (4.9%) | 0.71 | 0.029 | 1.5e-08 |
| 5kb (3.4%) | 0.51 | 0.028 | 2.7e-08 |
| 100kb (10.6%) | 0.68 | 0.017 | 1.2e-08 |

**Table S37.** S-LDSC results for Trans-master and Transcription Factor (TF) SNP annotations conditional on the baseline-LD+ model: Standardized Effect sizes (*τ\**) and Enrichment (E) of 20 SNP annotations corresponding to Trans-master and Transcription Factor genes. The analysis is conditional on 93 baseline-LD+ annotations. Reports are meta-analyzed across 11 blood-related traits.

| Trans-master |  |  |  |  |  |  |
| --- | --- | --- | --- | --- | --- | --- |
| Annotation | $\tau^*$ | $se(\tau^*)$ | $p(\tau^*)$ | $E$ | $se(E)$ | $p(E)$ |
| ABC (0.33%) | 2.3 | 0.17 | 3.5e-44 | 42 | 2.6 | 6.8e-08 |
| TSS (0.27%) | 2.6 | 0.23 | 6.4e-30 | 55 | 4 | 6.9e-08 |
| Coding (0.22%) | 2.4 | 0.28 | 2e-18 | 52 | 6 | 1.8e-07 |
| ATAC (0.21%) | 1.4 | 0.18 | 1.6e-14 | 33 | 4 | 1.3e-07 |
| eQTL (0.54%) | 0.9 | 0.11 | 3.4e-16 | 14 | 1.7 | 1.2e-07 |
| Roadmap (0.58%) | 1.7 | 0.11 | 2.1e-52 | 27 | 1.5 | 7.8e-09 |
| Promoter (0.57%) | 1.8 | 0.18 | 4.7e-22 | 27 | 2.4 | 8.4e-08 |
| PC-HiC (5.1%) | 0.69 | 0.041 | 3.9e-62 | 5.3 | 0.18 | 4.2e-09 |
| 5kb (4.2%) | 0.59 | 0.049 | 1.1e-32 | 5.2 | 0.3 | 1.3e-08 |
| 100kb (9.7%) | 0.56 | 0.044 | 2.5e-38 | 3.8 | 0.17 | 5.4e-09 |

| TF |  |  |  |  |  |  |
| --- | --- | --- | --- | --- | --- | --- |
| Annotation | $\tau^*$ | $se(\tau^*)$ | $p(\tau^*)$ | $E$ | $se(E)$ | $p(E)$ |
| ABC (0.32%) | 0.51 | 0.13 | 5.1e-05 | 17 | 1.7 | 2.5e-06 |
| TSS (0.16%) | 0.88 | 0.19 | 2.5e-06 | 30 | 4.4 | 8.6e-07 |
| Coding (0.16%) | 0.33 | 0.093 | 0.0004 | 12 | 1.9 | 0.00011 |
| ATAC (0.14%) | 0.7 | 0.11 | 1e-10 | 24 | 3.2 | 8.7e-07 |
| eQTL (0.17%) | 0.0086 | 0.039 | 0.83 | 4.5 | 0.9 | 0.0062 |
| Roadmap (0.30%) | 0.86 | 0.12 | 8.3e-13 | 23 | 2.4 | 4.1e-08 |
| Promoter (0.37%) | 0.34 | 0.084 | 6e-05 | 8.8 | 1.2 | 0.00018 |
| PC-HiC (4.9%) | 0.25 | 0.038 | 5.5e-11 | 3.3 | 0.14 | 1.2e-08 |
| 5kb (3.4%) | 0.19 | 0.031 | 1.4e-09 | 2.8 | 0.16 | 3.2e-08 |
| 100kb (10.6%) | 0.18 | 0.027 | 9.7e-11 | 2.2 | 0.058 | 8.3e-09 |

**Table S38.** S-LDSC results for Trans-master SNP annotations conditional on the baseline-LD+Cis1 model: Standardized Effect sizes (*τ\**) and Enrichment (E) of 10 Trans-master SNP annotations conditional on 103 baseline-LD+Cis1 (93 baseline-LD+ and 10 Cis1) annotations where Cis1 represents S2G annotations linked to genes with at least 1 trait-associated cis-eQTL. Reports are meta-analyzed across 11 blood-related traits.

| Trans-master |  |  |  |  |  |  |
| --- | --- | --- | --- | --- | --- | --- |
| Annotation | $\tau^*$ | $se(\tau^*)$ | $p(\tau^*)$ | $E$ | $se(E)$ | $p(E)$ |
| ABC (0.33%) | 1.9 | 0.18 | 1.5e-26 | 37 | 2.7 | 5.5e-07 |
| TSS (0.27%) | 1.8 | 0.22 | 7.8e-17 | 48 | 4.1 | 6.5e-07 |
| Coding (0.22%) | 1.8 | 0.29 | 2e-09 | 41 | 6 | 3.5e-06 |
| ATAC (0.21%) | 0.95 | 0.12 | 1e-15 | 27 | 3.6 | 1.4e-06 |
| eQTL (0.54%) | 0.7 | 0.12 | 3.4e-09 | 12 | 1.8 | 4.5e-07 |
| Roadmap (0.58%) | 1.4 | 0.12 | 1.8e-31 | 25 | 1.5 | 2.7e-08 |
| Promoter (0.57%) | 1.3 | 0.18 | 2.4e-13 | 24 | 2.8 | 7.2e-07 |
| PC-HiC (5.1%) | 0.51 | 0.041 | 4.1e-35 | 5 | 0.17 | 4.7e-09 |
| 5kb (4.2%) | 0.45 | 0.05 | 2.8e-19 | 5.8 | 0.31 | 1.5e-08 |
| 100kb (9.7%) | 0.41 | 0.046 | 8.3e-19 | 4 | 0.19 | 1.1e-08 |

**Table S39.** S-LDSC results for Trans-master SNP annotations conditional on the baseline-LD+Cis1+Cis2LD model: Standardized Effect sizes (*τ\**) and Enrichment (E) of 10 Trans-master SNP annotations conditional on 113 baseline-LD+Cis1+Cis2LD (93 baseline-LD+ and 10 Cis1 and 10 Cis2LD) annotations where Cis1 represents S2G annotations linked to genes with at least 1 trait-associated cis-eQTL and Cis2LD represents S2G lined to genes with 2 unlinked trait-associated cis-eQTLs. Reports are meta-analyzed across 11 blood-related traits.

| Trans-master |  |  |  |  |  |  |
| --- | --- | --- | --- | --- | --- | --- |
| Annotation | $\tau^*$ | $se(\tau^*)$ | $p(\tau^*)$ | $E$ | $se(E)$ | $p(E)$ |
| ABC (0.33%) | 1.7 | 0.2 | 2.4e-16 | 32 | 2.4 | 5.6e-07 |
| TSS (0.27%) | 1.8 | 0.22 | 8.3e-16 | 45 | 3.5 | 5e-07 |
| Coding (0.22%) | 1.7 | 0.27 | 1.6e-10 | 34 | 4.9 | 9.7e-06 |
| ATAC (0.21%) | 1 | 0.17 | 1.1e-09 | 24 | 3.6 | 1.1e-06 |
| eQTL (0.54%) | 0.66 | 0.1 | 6.1e-11 | 11 | 1.5 | 9.2e-07 |
| Roadmap (0.58%) | 1.3 | 0.12 | 2e-28 | 22 | 1.4 | 2.4e-08 |
| Promoter (0.57%) | 1.2 | 0.17 | 9e-13 | 21 | 2.2 | 1.1e-06 |
| PC-HiC (5.1%) | 0.48 | 0.04 | 2e-32 | 5 | 0.17 | 5.3e-09 |
| 5kb (4.2%) | 0.4 | 0.053 | 1.5e-14 | 5.3 | 0.3 | 1.6e-08 |
| 100kb (9.7%) | 0.39 | 0.045 | 1.4e-17 | 3.8 | 0.17 | 7.5e-09 |

**Table S40.** S-LDSC results for Trans-master SNP annotations conditional on the baseline-LD+Cis1+Cis2 model: Standardized Effect sizes (*τ\**) and Enrichment (E) of 10 Trans-master SNP annotations conditional on 113 baseline-LD+Cis1+Cis2 (93 baseline-LD+ and 10 Cis1 and 10 Cis2) annotations where Cis1 represents S2G annotations linked to genes with at least 1 trait-associated cis-eQTL and Cis2 represents S2G lined to genes with 2 not LD-corrected trait-associated cis-eQTLs. Reports are meta-analyzed across 11 blood-related traits.

| Trans-master |  |  |  |  |  |  |
| --- | --- | --- | --- | --- | --- | --- |
| Annotation | $\tau^*$ | $se(\tau^*)$ | $p(\tau^*)$ | $E$ | $se(E)$ | $p(E)$ |
| ABC (0.33%) | 1.5 | 0.21 | 6.7e-13 | 33 | 2.6 | 1.7e-06 |
| TSS (0.27%) | 1.3 | 0.3 | 5.8e-06 | 33 | 6 | 5e-05 |
| Coding (0.22%) | 1.3 | 0.3 | 5.8e-06 | 33 | 6 | 5e-05 |
| ATAC (0.21%) | 0.74 | 0.12 | 3.8e-10 | 23 | 3.4 | 5.3e-06 |
| eQTL (0.54%) | 0.57 | 0.12 | 1.3e-06 | 11 | 1.8 | 1.4e-06 |
| Roadmap (0.58%) | 1.1 | 0.12 | 4.1e-21 | 23 | 1.5 | 4.2e-08 |
| Promoter (0.57%) | 1 | 0.18 | 3.5e-08 | 21 | 3 | 2.6e-06 |
| PC-HiC (5.1%) | 0.4 | 0.04 | 3.1e-24 | 4.9 | 0.16 | 6.8e-09 |
| 5kb (4.2%) | 0.34 | 0.056 | 1e-09 | 5.8 | 0.33 | 1.8e-08 |
| 100kb (9.7%) | 0.31 | 0.053 | 5.5e-09 | 4 | 0.19 | 1.1e-08 |

**Table S41.**
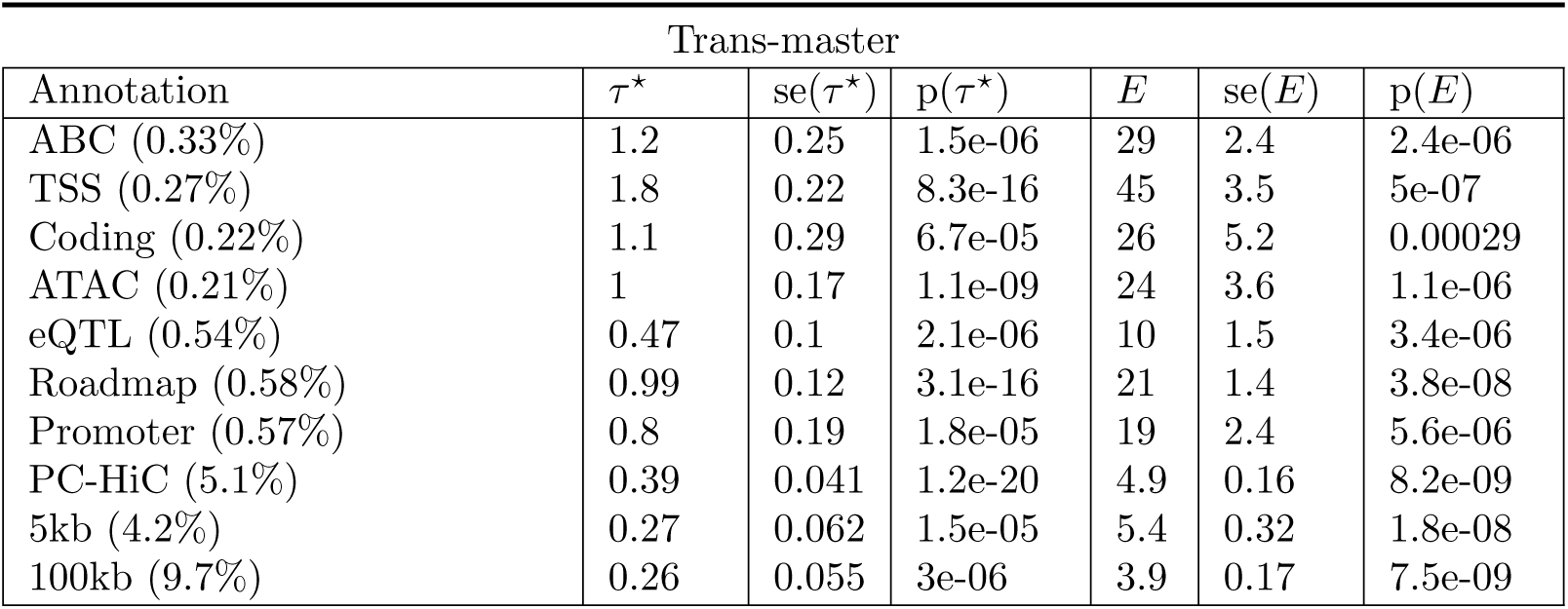
S-LDSC results for Trans-master SNP annotations conditional on the baseline-LD+Cis1+Cis3 model: Standardized Effect sizes (*τ\**) and Enrichment (E) of 10 Trans-master SNP annotations conditional on 113 baseline-LD+Cis1+Cis3 (93 baseline-LD+ and 10 Cis1 + 10 Cis3) annotations where Cis1 represents S2G annotations linked to genes with at least 1 trait-associated cis-eQTL and Cis3 represents S2G lined to genes with 3 not LD-corrected trait-associated cis-eQTLs. Reports are meta-analyzed across 11 blood-related traits.

**Table S42.** S-LDSC results for Trans-master SNP annotations conditional on the baseline-LD+Cis1+Cis4 model: Standardized Effect sizes (*τ\**) and Enrichment (E) of 10 Trans-master SNP annotations conditional on 113 baseline-LD+Cis1+Cis4 (93 baseline-LD+ and 10 Cis1 + 10 Cis4) annotations where Cis1 represents S2G annotations linked to genes with at least 1 trait-associated cis-eQTL and Cis4 represents S2G lined to genes with 4 not LD-corrected trait-associated cis-eQTLs. Reports are meta-analyzed across 11 blood-related traits.

| Trans-master |  |  |  |  |  |  |
| --- | --- | --- | --- | --- | --- | --- |
| Annotation | $\tau^*$ | $se(\tau^*)$ | $p(\tau^*)$ | $E$ | $se(E)$ | $p(E)$ |
| ABC (0.33%) | 1 | 0.28 | 0.00021 | 28 | 2.4 | 2.9e-06 |
| TSS (0.27%) | 1.1 | 0.27 | 4.4e-05 | 37 | 3.5 | 9.2e-06 |
| Coding (0.22%) | 0.88 | 0.29 | 0.0028 | 25 | 5.1 | 0.00032 |
| ATAC (0.21%) | 0.59 | 0.13 | 5.1e-06 | 19 | 3.3 | 1.4e-05 |
| eQTL (0.54%) | 0.38 | 0.11 | 0.00034 | 9.9 | 1.5 | 4.3e-06 |
| Roadmap (0.58%) | 0.86 | 0.12 | 1.7e-12 | 21 | 1.4 | 5.1e-08 |
| Promoter (0.57%) | 0.63 | 0.2 | 0.0012 | 18 | 2.4 | 1.7e-05 |
| PC-HiC (5.1%) | 0.36 | 0.044 | 6.2e-16 | 4.9 | 0.16 | 8.4e-09 |
| 5kb (4.2%) | 0.24 | 0.063 | 0.0002 | 5.4 | 0.31 | 2.2e-08 |
| 100kb (9.7%) | 0.23 | 0.058 | 7.3e-05 | 3.9 | 0.17 | 7.7e-09 |

**Table S43.** S-LDSC results for Trans-master SNP annotations conditional on the baseline-LD+cis model, but with all trait-associated trans-eQTLs removed.: Standardized Effect sizes (*τ\**) and Enrichment (E) of 10 Trans-master SNP annotations, but with SNPs that are among the 3,853 trait-associated trans-eQTL SNPs that were used to construct the Trans-master gene score. The analysis is conditional on 113 baseline-LD+cis annotations. Reports are meta-analyzed across 11 blood-related traits.

| Trans-master |  |  |  |  |  |  |
| --- | --- | --- | --- | --- | --- | --- |
| Annotation | $\tau^*$ | $se(\tau^*)$ | $p(\tau^*)$ | $E$ | $se(E)$ | $p(E)$ |
| ABC (0.33%) | 1 | 0.21 | 1.8e-06 | 30 | 2.5 | 2.2e-06 |
| TSS (0.27%) | 0.85 | 0.28 | 0.002 | 39 | 4.1 | 2.7e-06 |
| Coding (0.22%) | 0.99 | 0.29 | 0.00051 | 29 | 5.7 | 0.00012 |
| ATAC (0.21%) | 0.56 | 0.13 | 2.3e-05 | 19 | 3.2 | 2e-05 |
| eQTL (0.54%) | 0.43 | 0.12 | 0.00023 | 10 | 1.8 | 5.1e-06 |
| Roadmap (0.58%) | 0.98 | 0.11 | 2.4e-19 | 21 | 1.5 | 9.6e-08 |
| Promoter (0.57%) | 0.8 | 0.18 | 1.2e-05 | 20 | 2.7 | 4.9e-06 |
| PC-HiC (5.1%) | 0.41 | 0.04 | 2.7e-24 | 4.7 | 0.16 | 6.5e-09 |
| 5kb (4.2%) | 0.34 | 0.049 | 4.7e-12 | 5.4 | 0.29 | 3.8e-08 |
| 100kb (9.7%) | 0.32 | 0.046 | 5.2e-12 | 3.9 | 0.18 | 1.8e-08 |

**Table S44.**
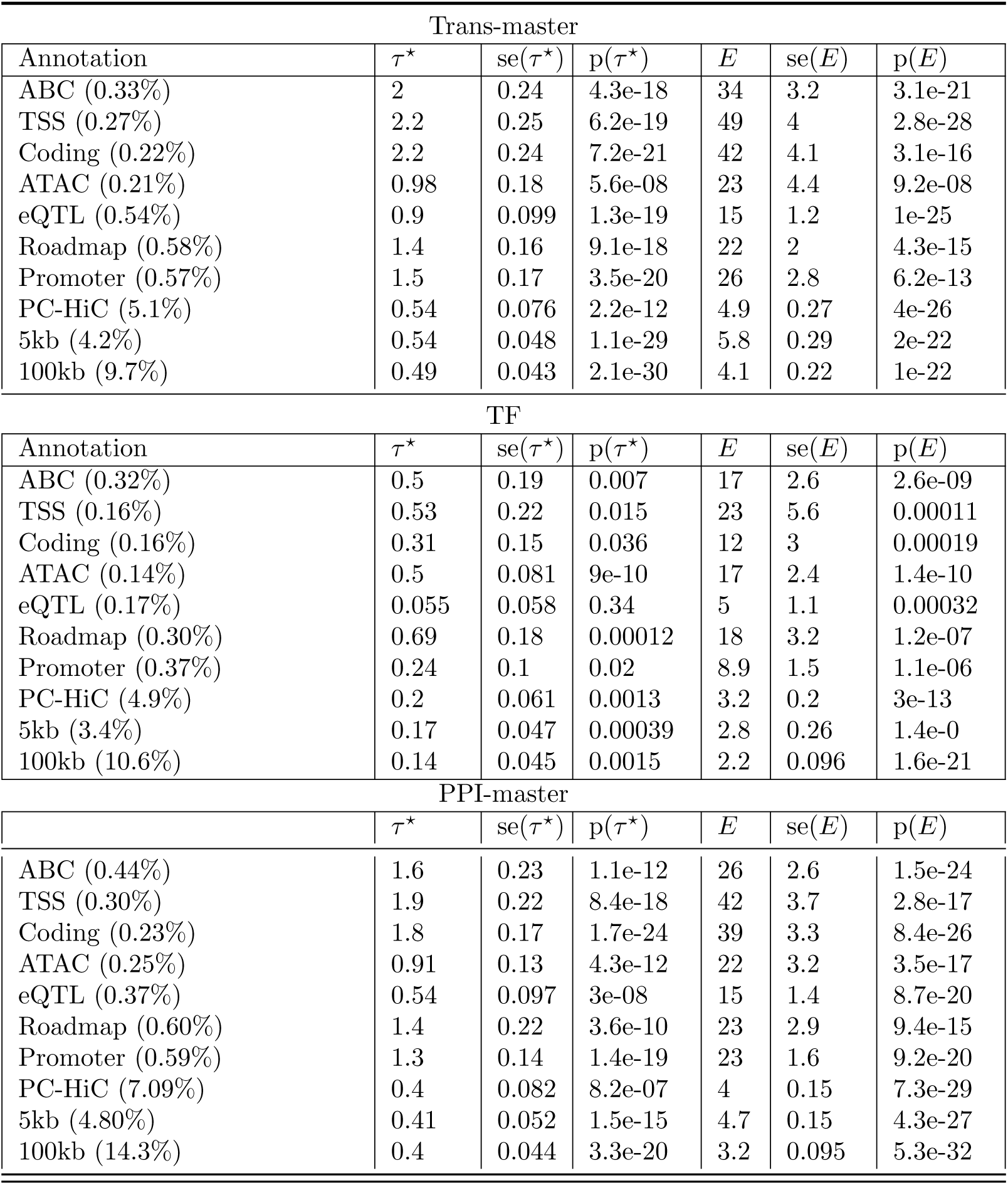
S-LDSC results for SNP annotations corresponding to candidate master-regulator gene scores meta-analyzed across 5 blood cell traits: Standardized Effect sizes (*τ\**) and Enrichment (E) of 30 SNP annotations corresponding to 2 candidate master-regulator gene scores and 1 PPI-master score and 10 S2G strategies, conditional on 113 baseline-LD+cis annotations. Reports are meta-analyzed across 5 blood cell traits.

**Table S45.** S-LDSC results for SNP annotations corresponding to candidate master-regulator gene scores meta-analyzed across 6 autoimmune diseases: Standardized Effect sizes (*τ\**) and Enrichment (E) of 30 SNP annotations corresponding to 2 candidate master-regulator gene scores and 1 PPI-master and 10 S2G strategies, conditional on 113 baseline-LD+cis annotations. Reports are meta-analyzed across 6 autoimmune diseases.

| Trans-master |  |  |  |  |  |  |
| --- | --- | --- | --- | --- | --- | --- |
| Annotation | $\tau^*$ | $se(\tau^*)$ | $p(\tau^*)$ | $E$ | $se(E)$ | $p(E)$ |
| ABC (0.33%) | 0.89 | 0.33 | 0.0069 | 28 | 4 | 0.0002 |
| TSS (0.27%) | 1 | 0.27 | 0.00016 | 37 | 5.4 | 8.5e-05 |
| Coding (0.22%) | 0.91 | 0.3 | 0.0024 | 20 | 6 | 0.0053 |
| ATAC (0.21%) | 0.81 | 0.29 | 0.0052 | 25 | 6.4 | 0.0006 |
| eQTL (0.54%) | 0.37 | 0.11 | 0.00067 | 7.6 | 1.6 | 0.00031 |
| Roadmap (0.58%) | 1.1 | 0.18 | 1.3e-09 | 23 | 2.3 | 4.5e-05 |
| Promoter (0.57%) | 0.66 | 0.18 | 0.00037 | 16 | 2.2 | 0.0003 |
| PC-HiC (5.1%) | 0.41 | 0.075 | 5.8e-08 | 5.2 | 0.32 | 9.4e-06 |
| 5kb (4.2%) | 0.27 | 0.054 | 6.5e-07 | 4.8 | 0.41 | 7.4e-06 |
| 100kb (9.7%) | 0.25 | 0.047 | 8.6e-08 | 3.5 | 0.22 | 5.5e-06 |

| TF |  |  |  |  |  |  |
| --- | --- | --- | --- | --- | --- | --- |
| Annotation | $\tau^*$ | $se(\tau^*)$ | $p(\tau^*)$ | $E$ | $se(E)$ | $p(E)$ |
| ABC (0.32%) | 0.48 | 0.23 | 0.037 | 22 | 3.4 | 0.00085 |
| TSS (0.16%) | 1.3 | 0.23 | 6e-09 | 40 | 5.8 | 0.00039 |
| Coding (0.16%) | 0.44 | 0.16 | 0.0046 | 17 | 3.9 | 0.0098 |
| ATAC (0.14%) | 1.2 | 0.2 | 6.1e-09 | 37 | 5.7 | 0.00033 |
| eQTL (0.17%) | 0.018 | 0.059 | 0.76 | 2.8 | 1.3 | 0.44 |
| Roadmap (0.30%) | 1 | 0.17 | 2.2e-09 | 25 | 3 | 6e-05 |
| Promoter (0.37%) | 0.47 | 0.14 | 0.00057 | 11 | 2.3 | 0.02 |
| PC-HiC (4.9%) | 0.27 | 0.059 | 4e-06 | 3.7 | 0.26 | 7.1e-05 |
| 5kb (3.4%) | 0.24 | 0.04 | 2.4e-09 | 3 | 0.22 | 8.4e-05 |
| 100kb (10.6%) | 0.2 | 0.036 | 4.3e-08 | 2.3 | 0.11 | 1.6e-05 |

| PPI-master |  |  |  |  |  |  |
| --- | --- | --- | --- | --- | --- | --- |
| | $\tau^*$ | $se(\tau^*)$ | $p(\tau^*)$ | $E$ | $se(E)$ | $p(E)$ |
| ABC (0.44%) | 1.5 | 0.29 | 1.7e-07 | 29 | 3.4 | 5.7e-05 |
| TSS (0.30%) | 1.5 | 0.24 | 3.3e-10 | 38 | 4.6 | 5.8e-05 |
| Coding (0.23%) | 1.5 | 0.23 | 1.9e-10 | 33 | 5.2 | 0.00032 |
| ATAC (0.25%) | 1.8 | 0.24 | 1.3e-13 | 37 | 4.3 | 4.2e-05 |
| eQTL (0.37%) | 0.43 | 0.078 | 2.8e-08 | 9.4 | 1.3 | 0.00045 |
| Roadmap (0.60%) | 1.6 | 0.17 | 6.1e-21 | 28 | 2.3 | 2.3e-05 |
| Promoter (0.59%) | 0.93 | 0.16 | 1.2e-08 | 17 | 2.3 | 0.00021 |
| PC-HiC (7.09%) | 0.35 | 0.064 | 3.4e-08 | 4.1 | 0.25 | 9.8e-06 |
| 5kb (4.80%) | 0.34 | 0.045 | 1e-13 | 4.3 | 0.25 | 5.3e-06 |
| 100kb (14.3%) | 0.33 | 0.04 | 1.3e-16 | 3 | 0.13 | 4.5e-06 |

**Table S46.** S-LDSC results for SNP annotations corresponding to candidate master-regulator gene scores meta-analyzed across 2 granulocyte-related blood cell traits: Standardized Effect sizes (*τ\**) and Enrichment (E) of 30 SNP annotations corresponding to 2 candidate master-regulator gene scores and 1 PPI-master score and 10 S2G strategies, conditional on 113 baseline-LD+cis annotations. Reports are meta-analyzed across 2 granulocyte related traits (white blood cell count and eosinophil count).

| Trans-master |  |  |  |  |  |  |
| --- | --- | --- | --- | --- | --- | --- |
| Annotation | $\tau^*$ | $se(\tau^*)$ | $p(\tau^*)$ | $E$ | $se(E)$ | $p(E)$ |
| ABC (0.33%) | 2.3 | 0.58 | 0.00011 | 37 | 8.5 | 4.1e-05 |
| TSS (0.27%) | 2.6 | 0.43 | 1.2e-09 | 52 | 7 | 2.9e-11 |
| Coding (0.22%) | 2.5 | 0.61 | 6.1e-05 | 41 | 6.9 | 5.3e-08 |
| ATAC (0.21%) | 1.6 | 0.64 | 0.013 | 33 | 11 | 0.0035 |
| eQTL (0.54%) | 0.88 | 0.15 | 7.4e-09 | 14 | 1.7 | 7.1e-13 |
| Roadmap (0.58%) | 1.6 | 0.37 | 1.1e-05 | 26 | 6.2 | 0.00015 |
| Promoter (0.57%) | 1.6 | 0.3 | 3.8e-08 | 21 | 3 | 8.2e-10 |
| PC-HiC (5.1%) | 0.54 | 0.076 | 2.2e-12 | 4.9 | 0.27 | 4e-26 |
| 5kb (4.2%) | 0.62 | 0.13 | 2e-06 | 5.6 | 0.54 | 1.5e-15 |
| 100kb (9.7%) | 0.55 | 0.12 | 4.7e-06 | 4 | 0.41 | 8.6e-12 |

| TF |  |  |  |  |  |  |
| --- | --- | --- | --- | --- | --- | --- |
| Annotation | $\tau^*$ | $se(\tau^*)$ | $p(\tau^*)$ | $E$ | $se(E)$ | $p(E)$ |
| ABC (0.32%) | 0.6 | 0.44 | 0.17 | 20 | 7.7 | 0.01 |
| TSS (0.16%) | 0.82 | 0.44 | 0.062 | 30 | 12 | 0.018 |
| Coding (0.16%) | 0.48 | 0.28 | 0.087 | 15 | 5.6 | 0.01 |
| ATAC (0.14%) | 0.58 | 0.21 | 0.0067 | 21 | 6.6 | 0.0024 |
| eQTL (0.17%) | 0.26 | 0.15 | 0.079 | 9.8 | 3.5 | 0.014 |
| Roadmap (0.30%) | 1.0 | 0.55 | 0.063 | 25 | 11 | 0.036 |
| Promoter (0.37%) | 0.31 | 0.17 | 0.062 | 7.9 | 2.3 | 0.0034 |
| PC-HiC (4.9%) | 0.27 | 0.12 | 0.022 | 3.4 | 0.47 | 7.3e-07 |
| 5kb (3.4%) | 0.2 | 0.078 | 0.0088 | 2.8 | 0.38 | 4.1e-06 |
| 100kb (10.6%) | 0.2 | 0.06 | 0.0012 | 2.2 | 0.17 | 2.4e-12 |

| PPI-master |  |  |  |  |  |  |
| --- | --- | --- | --- | --- | --- | --- |
| | $\tau^*$ | $se(\tau^*)$ | $p(\tau^*)$ | $E$ | $se(E)$ | $p(E)$ |
| ABC (0.44%) | 2.0 | 0.39 | 3.5e-07 | 30 | 5.7 | 9.3e-07 |
| TSS (0.30%) | 2.3 | 0.33 | 1.4e-11 | 49 | 5.8 | 1.3e-13 |
| Coding (0.23%) | 2.0 | 0.28 | 3.2e-12 | 40 | 4.9 | 1.3e-13 |
| ATAC (0.25%) | 1.1 | 0.39 | 0.0036 | 28 | 8.1 | 0.0012 |
| eQTL (0.37%) | 0.8 | 0.15 | 1.6e-07 | 17 | 2.3 | 2.3e-10 |
| Roadmap (0.60%) | 1.8 | 0.41 | 1.3e-05 | 28 | 6.9 | 0.00016 |
| Promoter (0.59%) | 1.4 | 0.22 | 1.5e-10 | 20 | 2.6 | 1.5e-11 |
| PC-HiC (7.09%) | 0.54 | 0.07 | 7.1e-15 | 4.2 | 0.23 | 1.8e-33 |
| 5kb (4.80%) | 0.51 | 0.075 | 8.8e-12 | 4.8 | 0.36 | 3.5e-23 |
| 100kb (14.3%) | 0.48 | 0.085 | 1.8e-08 | 3.2 | 0.26 | 9.7e-16 |

**Table S47.** S-LDSC results for SNP annotations corresponding to candidate master-regulator gene scores meta-analyzed across across 3 red blood cell or platelet-related blood cell traits: Standardized Effect sizes (*τ\**) and Enrichment (E) of 30 SNP annotations corresponding to 2 candidate master-regulator gene scores and 1 PPI-master score and 10 S2G strategies, conditional on 113 baseline-LD+cis annotations. Reports are meta-analyzed across 3 red blood cell or platelet related traits (red blood count, red blood distribution width and platelet count).

| Trans-master |  |  |  |  |  |  |
| --- | --- | --- | --- | --- | --- | --- |
| Annotation | $\tau^*$ | $se(\tau^*)$ | $p(\tau^*)$ | $E$ | $se(E)$ | $p(E)$ |
| ABC (0.33%) | 2 | 0.3 | 3.5e-11 | 32 | 4.1 | 2.4e-12 |
| TSS (0.27%) | 2.1 | 0.31 | 5.2e-11 | 48 | 4.8 | 1.3e-18 |
| Coding (0.22%) | 2.2 | 0.3 | 6.4e-13 | 43 | 5.9 | 1.1e-08 |
| ATAC (0.21%) | 0.98 | 0.18 | 5.6e-08 | 23 | 4.4 | 9.2e-08 |
| eQTL (0.54%) | 0.91 | 0.13 | 2.9e-12 | 16 | 1.6 | 2.1e-18 |
| Roadmap (0.58%) | 1.2 | 0.21 | 4.2e-09 | 20 | 2.3 | 3.3e-08 |
| Promoter (0.57%) | 1.5 | 0.23 | 2.4e-11 | 30 | 2.8 | 7e-17 |
| PC-HiC (5.1%) | 0.42 | 0.064 | 6.9e-11 | 4.6 | 0.26 | 1.2e-09 |
| 5kb (4.2%) | 0.5 | 0.066 | 4.2e-14 | 6.1 | 0.3 | 5.1e-13 |
| 100kb (9.7%) | 0.47 | 0.053 | 4.7e-19 | 4.2 | 0.31 | 2.4e-13 |

| TF |  |  |  |  |  |  |
| --- | --- | --- | --- | --- | --- | --- |
| Annotation | $\tau^*$ | $se(\tau^*)$ | $p(\tau^*)$ | $E$ | $se(E)$ | $p(E)$ |
| ABC (0.32%) | 0.49 | 0.25 | 0.054 | 16 | 2.7 | 1.5e-08 |
| TSS (0.16%) | 0.38 | 0.28 | 0.17 | 19 | 7 | 0.015 |
| Coding (0.16%) | 0.24 | 0.2 | 0.23 | 11 | 3.9 | 0.015 |
| ATAC (0.14%) | 0.49 | 0.11 | 9.4e-06 | 16 | 2.8 | 8e-06 |
| eQTL (0.17%) | 0.018 | 0.058 | 0.75 | 4.4 | 1.2 | 0.0039 |
| Roadmap (0.30%) | 0.57 | 0.2 | 0.0041 | 15 | 2.9 | 3.3e-06 |
| Promoter (0.37%) | 0.21 | 0.14 | 0.15 | 9.6 | 2.3 | 0.00048 |
| PC-HiC (4.9%) | 0.15 | 0.084 | 0.068 | 3.1 | 0.27 | 1.1e-05 |
| 5kb (3.4%) | 0.15 | 0.07 | 0.033 | 2.7 | 0.4 | 0.0012 |
| 100kb (10.6%) | 0.11 | 0.065 | 0.1 | 2.2 | 0.15 | 7.9e-11 |

| PPI-master |  |  |  |  |  |  |
| --- | --- | --- | --- | --- | --- | --- |
| | $\tau^*$ | $se(\tau^*)$ | $p(\tau^*)$ | $E$ | $se(E)$ | $p(E)$ |
| ABC (0.44%) | 1.4 | 0.24 | 4.8e-09 | 23 | 2.8 | 2.1e-13 |
| TSS (0.30%) | 1.9 | 0.22 | 8.4e-18 | 42 | 3.7 | 2.8e-17 |
| Coding (0.23%) | 1.6 | 0.23 | 1.3e-12 | 38 | 4.5 | 8.7e-14 |
| ATAC (0.25%) | 0.91 | 0.13 | 4.3e-12 | 22 | 3.2 | 3.5e-17 |
| eQTL (0.37%) | 0.4 | 0.11 | 0.00016 | 13 | 1.8 | 5.4e-11 |
| Roadmap (0.60%) | 1.1 | 0.23 | 6.4e-07 | 19 | 1.9 | 3.1e-09 |
| Promoter (0.59%) | 1.2 | 0.19 | 1.3e-10 | 25 | 2.1 | 6e-13 |
| PC-HiC (7.09%) | 0.31 | 0.1 | 0.0019 | 3.8 | 0.17 | 8.2e-10 |
| 5kb (4.80%) | 0.35 | 0.049 | 9.3e-13 | 4.7 | 0.2 | 1.2e-10 |
| 100kb (14.3%) | 0.36 | 0.042 | 5.3e-17 | 3.2 | 0.088 | 1.4e-20 |

**Table S48.** S-LDSC results for jointly significant Trans-master and Transcription Factor SNP annotations: Standardized Effect sizes (*τ\**) and Enrichment (E) of SNP annotations that are significant in a joint analysis of all marginally significant Trans-master and Transcription Factor (TF) SNP annotations. The analysis is conditional on 113 baseline-LD+cis annotations. Reports are meta-analyzed across 11 blood-related traits.

| Annotation | $\tau^*$ | $se(\tau^*)$ | $p(\tau^*)$ | $E$ | $se(E)$ | $p(E)$ |
| --- | --- | --- | --- | --- | --- | --- |
| Trans-master × Roadmap<br>(0.58%) | 0.81 | 0.13 | 5.4e-10 | 19 | 1.4 | 8.4e-08 |
| Trans-master × PC-HiC<br>(5.1%) | 0.2 | 0.044 | 8.3e-06 | 4.6 | 0.16 | 5.9e-09 |
| Trans-master × 5kb<br>(4.2%) | 0.19 | 0.049 | 1.1e-04 | 5.2 | 0.29 | 2.3e-08 |
| TF × ATAC<br>(0.14%) | 0.4 | 0.08 | 7.2e-07 | 17 | 2.5 | 7e-06 |
| TF × Roadmap<br>(0.30%) | 0.58 | 0.11 | 2.2e-07 | 19 | 2.2 | 2.7e-07 |

**Table S49.** S-LDSC results for Trans-regulated SNP annotations conditional on the baseline-LD+cis model: Standardized Effect sizes (*τ\**) and Enrichment (E) of 10 SNP annotations corresponding to Trans-regulated genes. The analysis is conditional on 113 baseline-LD+cis annotations. Reports are meta-analyzed across 11 blood-related traits.

| Trans-Regulated |  |  |  |  |  |  |
| --- | --- | --- | --- | --- | --- | --- |
| Annotation | $\tau^*$ | $se(\tau^*)$ | $p(\tau^*)$ | $E$ | $se(E)$ | $p(E)$ |
| ABC (0.87%) | -0.12 | 0.15 | 0.43 | 10 | 0.97 | 1.1e-06 |
| TSS (0.70%) | 0.068 | 0.11 | 0.54 | 14 | 1.3 | 2.1e-05 |
| Coding (0.56%) | 0.2 | 0.095 | 0.04 | 9.3 | 1.3 | 0.0004 |
| ATAC (0.72%) | 0.12 | 0.077 | 0.13 | 7.5 | 1.1 | 9.2e-06 |
| eQTL (1.07%) | 0.016 | 0.049 | 0.74 | 4 | 0.54 | 0.00028 |
| Roadmap (1.67%) | -0.091 | 0.11 | 0.41 | 7.8 | 0.75 | 2.2e-08 |
| Promoter (1.47%) | 0.02 | 0.072 | 0.78 | 6.1 | 0.6 | 0.00012 |
| PC-HiC (16.0%) | -0.057 | 0.044 | 0.2 | 2.4 | 0.075 | 3.2e-09 |
| 5kb (14.9%) | -0.021 | 0.02 | 0.3 | 2.2 | 0.046 | 4.9e-09 |
| 100kb (34.7%) | -0.023 | 0.019 | 0.23 | 1.9 | 0.044 | 1.8e-09 |

**Table S50.** S-LDSC results for Trans-regulated SNP annotations conditional on the baseline-LD+ model: Standardized Effect sizes (*τ\**) and Enrichment (E) of 10 SNP annotations corresponding to Trans-regulated genes. The analysis is conditional on 93 baseline-LD+ annotations. Reports are meta-analyzed across 11 blood-related traits.

| Trans-Regulated |  |  |  |  |  |  |
| --- | --- | --- | --- | --- | --- | --- |
| Annotation | $\tau^*$ | $se(\tau^*)$ | $p(\tau^*)$ | $E$ | $se(E)$ | $p(E)$ |
| ABC (0.87%) | 0.37 | 0.16 | 0.018 | 11 | 0.98 | 4.2e-07 |
| TSS (0.70%) | 0.5 | 0.11 | 5.6e-06 | 15 | 1.2 | 9.1e-06 |
| Coding (0.56%) | 0.55 | 0.098 | 2.3e-08 | 12 | 1.2 | 6.4e-05 |
| ATAC (0.72%) | 0.34 | 0.099 | 0.00053 | 8.2 | 1.1 | 1.5e-06 |
| eQTL (1.07%) | 0.091 | 0.051 | 0.075 | 4.8 | 0.53 | 2.9e-05 |
| Roadmap (1.67%) | 0.23 | 0.1 | 0.028 | 8.6 | 0.76 | 4.6e-09 |
| Promoter (1.47%) | 0.38 | 0.072 | 8e-08 | 6.7 | 0.57 | 5.7e-05 |
| PC-HiC (16.0%) | 0.033 | 0.04 | 0.4 | 2.4 | 0.076 | 2.6e-09 |
| 5kb (14.9%) | 0.033 | 0.02 | 0.1 | 2.2 | 0.049 | 2.6e-09 |
| 100kb (34.7%) | 0.016 | 0.019 | 0.41 | 1.9 | 0.042 | 8.4e-10 |

**Table S51.** S-LDSC results restricted to 6 autoimmune traits for Trans-master SNP annotations conditional on baseline-LD+cis model: Standardized Effect sizes (*τ\**) and Enrichment (E) of 10 Trans-master SNP annotations conditional on 113 baseline-LD+cis annotations, with results meta-analyzed across 5 Autoimmune traits.

| Trans-master |  |  |  |  |  |  |
| --- | --- | --- | --- | --- | --- | --- |
| Annotation | $\tau^*$ | $se(\tau^*)$ | $p(\tau^*)$ | $E$ | $se(E)$ | $p(E)$ |
| ABC (0.33%) | 0.89 | 0.33 | 0.0069 | 28 | 4 | 0.0002 |
| TSS (0.27%) | 1 | 0.27 | 0.00016 | 37 | 5.4 | 8.5e-05 |
| Coding (0.22%) | 0.91 | 0.3 | 0.0024 | 20 | 6 | 0.0053 |
| ATAC (0.21%) | 0.81 | 0.29 | 0.0052 | 25 | 6.4 | 0.0006 |
| eQTL (0.54%) | 0.37 | 0.11 | 0.00067 | 7.6 | 1.6 | 0.00031 |
| Roadmap (0.58%) | 1.1 | 0.18 | 1.3e-09 | 23 | 2.3 | 4.5e-05 |
| Promoter (0.57%) | 0.66 | 0.18 | 0.00037 | 16 | 2.2 | 0.00031 |
| PC-HiC (5.1%) | 0.41 | 0.075 | 5.8e-08 | 5.2 | 0.32 | 9.4e-06 |
| 5kb (4.2%) | 0.27 | 0.054 | 6.5e-07 | 4.8 | 0.41 | 7.4e-06 |
| 100kb (9.7%) | 0.25 | 0.047 | 8.6e-08 | 3.5 | 0.22 | 5.5e-06 |

**Table S52.** S-LDSC results for SNP annotations corresponding to Trans-master gene score based on genes that regulate *≥* 10 genes or *≥* 1 gene (instead of *≥* 3 genes): Standardized Effect sizes (*τ\**) and Enrichment (E) of 20 SNP annotations corresponding to Trans-master gene score but instead of trans-regulating *≥* 3 genes by any significant cis-eQTL of the focal gene as per original definition, we define two versions corresponding to trans-regulating *≥* 10 (3717) genes or *≥* 1 (1170) genes. Results are conditional on 113 baseline-LD+cis annotations for PPI-master. Reports are meta-analyzed across 11 blood-related traits.

| Trans-master ($\geq 1$) | | | | | | |
| --- | --- | --- | --- | --- | --- | --- |
| Annotation | $\tau^*$ | $se(\tau^*)$ | $p(\tau^*)$ | $E$ | $se(E)$ | $p(E)$ |
| ABC (0.19%) | 1.6 | 0.17 | 1.2e-22 | 35 | 3.3 | 4.9e-07 |
| TSS (0.15%) | 1.8 | 0.3 | 7.8e-10 | 37 | 7.3 | 4.3e-07 |
| Coding (0.12%) | 2.0 | 0.32 | 2.0e-10 | 37 | 9.4 | 7.1e-07 |
| ATAC (0.11%) | 0.9 | 0.13 | 6.6e-12 | 32 | 3.9 | 1e-05 |
| eQTL (0.28%) | 0.74 | 0.11 | 1.2e-11 | 17 | 2.3 | 1.2e-07 |
| Roadmap (0.33%) | 1.3 | 0.12 | 9.6e-27 | 29 | 2 | 8.6e-08 |
| Promoter (0.30%) | 1.3 | 0.23 | 1.5e-10 | 34 | 4.3 | 4.3e-07 |
| PC-HiC (2.91%) | 0.55 | 0.047 | 1.3e-31 | 5.9 | 0.27 | 1.3e-08 |
| 5kb (2.21%) | 0.5 | 0.05 | 3.7e-24 | 6.1 | 0.38 | 5.6e-08 |
| 100kb (5.24%) | 0.45 | 0.049 | 5.5e-20 | 4.2 | 0.23 | 3e-08 |

| Trans-master ($\geq 10$) | | | | | | |
| --- | --- | --- | --- | --- | --- | --- |
| Annotation | $\tau^*$ | $se(\tau^*)$ | $p(\tau^*)$ | $E$ | $se(E)$ | $p(E)$ |
| ABC (0.51%) | 1.6 | 0.16 | 7.7e-24 | 32 | 1.7 | 1.7e-08 |
| TSS (0.45%) | 1.5 | 0.16 | 3.4e-23 | 45 | 2.1 | 4.6e-08 |
| Coding (0.38%) | 1.3 | 0.19 | 3.9e-12 | 41 | 3.1 | 8.1e-08 |
| ATAC (0.35%) | 1.0 | 0.16 | 2.0e-10 | 25 | 2.6 | 4.7e-08 |
| eQTL (0.90%) | 0.59 | 0.088 | 5.8e-11 | 11 | 1 | 1.1e-07 |
| Roadmap (0.92%) | 1.4 | 0.097 | 1.6e-47 | 20 | 1.2 | 1e-09 |
| Promoter (0.95%) | 0.9 | 0.16 | 9.3e-07 | 22 | 1.7 | 5.3e-08 |
| PC-HiC (8.96%) | 0.47 | 0.039 | 9.5e-37 | 4.2 | 0.15 | 4.2e-10 |
| 5kb (8.01%) | 0.38 | 0.043 | 4.9e-19 | 4.1 | 0.16 | 5.1e-09 |
| 100kb (17.4%) | 0.39 | 0.04 | 9.3e-23 | 3.1 | 0.099 | 1.3e-09 |

**Table S53.** S-LDSC results for SNP annotations corresponding to the PPI-master gene score conditional on the baseline-LD+cis model: Standardized Effect sizes (*τ\**) and Enrichment (E) of 10 SNP annotations corresponding to the PPI-master gene score, conditional on 113 baseline-LD+cis annotations. Reports are meta-analyzed across 11 blood-related traits.

| PPI-master |  |  |  |  |  |  |
| --- | --- | --- | --- | --- | --- | --- |
| | $\tau^*$ | $\text{se}(\tau^*)$ | $\text{p}(\tau^*)$ | $E$ | $\text{se}(E)$ | $\text{p}(E)$ |
| ABC (0.44%) | 1.6 | 0.17 | 5.1e-21 | 27 | 1.9 | 9.8e-08 |
| TSS (0.30%) | 1.7 | 0.16 | 5.6e-27 | 41 | 2.8 | 2.1e-07 |
| Coding (0.23%) | 1.7 | 0.14 | 4.2e-33 | 37 | 2.8 | 1.2e-06 |
| ATAC (0.25%) | 1.3 | 0.17 | 4.6e-15 | 29 | 3.3 | 7.6e-09 |
| eQTL (0.37%) | 0.47 | 0.058 | 2.5e-16 | 12 | 1.1 | 2.4e-06 |
| Roadmap (0.60%) | 1.5 | 0.12 | 4.4e-33 | 25 | 1.9 | 5.5e-09 |
| Promoter (0.59%) | 1.1 | 0.11 | 4.3e-26 | 21 | 1.4 | 8e-07 |
| PC-HiC (7.09%) | 0.38 | 0.05 | 2.5e-14 | 4 | 0.12 | 5.8e-09 |
| 5kb (4.80%) | 0.38 | 0.031 | 2.4e-35 | 4.6 | 0.13 | 8.6e-09 |
| 100kb (14.3%) | 0.37 | 0.028 | 1.1e-40 | 3.2 | 0.075 | 4.7e-09 |

**Table S54.** Standardized enrichment of SNP annotations corresponding to PPI-master gene score conditional on the baseline-LD+cis model: Standardized enrichment of the 10 SNP annotations corresponding to the PPI-master gene score, conditional on 113 baseline-LD+cis annotations respectively. Reports are meta-analyzed across 11 blood-related traits.

| PPI-master |  |  |  |
| --- | --- | --- | --- |
| | $StdE$ | $se(StdE)$ | $p(StdE)$ |
| ABC (0.44%) | 1.8 | 0.12 | 9.8e-08 |
| TSS (0.30%) | 2.2 | 0.15 | 2.1e-07 |
| Coding (0.23%) | 1.8 | 0.13 | 1.2e-06 |
| ATAC (0.25%) | 1.5 | 0.17 | 7.6e-09 |
| eQTL (0.37%) | 0.72 | 0.069 | 2.4e-06 |
| Roadmap (0.60%) | 1.9 | 0.15 | 5.5e-09 |
| Promoter (0.59%) | 1.6 | 0.11 | 8e-07 |
| PC-HiC (7.09%) | 1 | 0.031 | 5.8e-09 |
| 5kb (4.80%) | 0.98 | 0.028 | 8.6e-09 |
| 100kb (14.3%) | 1.1 | 0.026 | 4.7e-09 |

**Table S55.** S-LDSC results for SNP annotations corresponding to the PPI-master gene score conditional on the baseline-LD+ model: Standardized Effect sizes (*τ\**) and Enrichment (E) of 10 SNP annotations corresponding to the PPI-master gene score, conditional on 93 baseline-LD+ annotations. Reports are meta-analyzed across 11 blood-related traits.

| PPI-master |  |  |  |  |  |  |
| --- | --- | --- | --- | --- | --- | --- |
| | $\tau^*$ | $se(\tau^*)$ | $p(\tau^*)$ | $E$ | $se(E)$ | $p(E)$ |
| ABC (0.44%) | 2.2 | 0.15 | 1.5e-47 | 33 | 1.9 | 1.8e-08 |
| TSS (0.30%) | 2.5 | 0.16 | 3e-55 | 50 | 2.8 | 4.2e-08 |
| Coding (0.23%) | 2.4 | 0.14 | 3.3e-61 | 50 | 2.9 | 5e-08 |
| ATAC (0.25%) | 1.7 | 0.19 | 5.7e-20 | 34 | 3.4 | 1.1e-09 |
| eQTL (0.37%) | 0.77 | 0.061 | 3.4e-36 | 15 | 1.2 | 2.2e-07 |
| Roadmap (0.60%) | 1.8 | 0.11 | 9.2e-64 | 28 | 1.8 | 2e-09 |
| Promoter (0.59%) | 1.7 | 0.11 | 6.7e-51 | 25 | 1.4 | 8.4e-08 |
| PC-HiC (7.09%) | 0.56 | 0.038 | 2.8e-47 | 4.2 | 0.12 | 3.4e-09 |
| 5kb (4.80%) | 0.51 | 0.029 | 3e-68 | 4.6 | 0.13 | 5.5e-09 |
| 100kb (14.3%) | 0.52 | 0.025 | 3.9e-93 | 3.2 | 0.073 | 2.6e-09 |

**Table S56.** S-LDSC results for SNP annotations corresponding to the PPI-master gene score conditional on the candidate master-regulator joint model: Standardized Effect sizes (*τ\**) and Enrichment (E) of 10 SNP annotations corresponding to the PPI-master gene score, conditional on the candidate master-regulator joint model from Table S48. Reports are meta-analyzed across 11 blood-related traits.

| PPI-master |  |  |  |  |  |  |
| --- | --- | --- | --- | --- | --- | --- |
| | $\tau^*$ | $se(\tau^*)$ | $p(\tau^*)$ | $E$ | $se(E)$ | $p(E)$ |
| ABC (0.44%) | 0.8 | 0.14 | 1.8e-08 | 21 | 1.8 | 7.7e-07 |
| TSS (0.30%) | 0.85 | 0.16 | 5e-08 | 30 | 2.7 | 2.9e-06 |
| Coding (0.23%) | 0.84 | 0.13 | 9.5e-11 | 23 | 2.7 | 4.4e-05 |
| ATAC (0.25%) | 0.73 | 0.15 | 1.5e-06 | 23 | 3 | 6e-08 |
| eQTL (0.37%) | 0.23 | 0.057 | 6.5e-05 | 9.3 | 0.99 | 1.5e-05 |
| Roadmap (0.60%) | 0.9 | 0.12 | 5.9e-15 | 23 | 1.8 | 9.6e-09 |
| Promoter (0.59%) | 0.49 | 0.11 | 3.9e-06 | 15 | 1.5 | 1.6e-05 |
| PC-HiC (7.09%) | 0.15 | 0.051 | 0.003 | 3.8 | 0.11 | 8.3e-09 |
| 5kb (4.80%) | 0.2 | 0.028 | 2e-12 | 4.6 | 0.13 | 7.8e-09 |
| 100kb (14.3%) | 0.18 | 0.024 | 6.7e-15 | 3.2 | 0.072 | 4.9e-09 |

**Table S57.** S-LDSC results for a joint analysis of all SNP annotations corresponding to the candidate master-regulator and PPI-master gene scores. Standardized Effect sizes (*τ\**) and Enrichment (E) of the significant SNP annotations in a joint analysis comprising of marginally significant candidate master-regulator SNP annotations from Figure 4B and PPI-master SNP annotations. Reported are the results for the jointly significant annotations only. Also marked in red are annotations that are jointly Bonferroni significant in the candidate master-regulator joint model in Supplementary Figure S14 but not Bonferroni significant in this model. All analyses are conditional on 113 baseline-LD+cis annotations. Results are meta-analyzed across 11 blood-related traits.

| Annotation | $\tau^*$ | se($\tau^*$) | p($\tau^*$) | $E$ | se ($E$) | p($E$) |
| --- | --- | --- | --- | --- | --- | --- |
| Trans-master × Roadmap<br>(0.58%) | 0.50 | 0.14 | 0.0003 | 21 | 1.4 | 5.1e-08 |
| Trans-master × PCHiC<br>(5.1%) | 0.24 | 0.049 | 1.2e-06 | 4.7 | 0.17 | 7.5e-09 |
| PPI-master × Coding<br>(0.23%) | 0.54 | 0.13 | 2.4e-05 | 19 | 2.7 | 0.00067 |
| PPI-master × ATAC<br>(0.25%) | 0.63 | 0.14 | 7.4e-06 | 20 | 2.9 | 1.7e-07 |
| PPI-master × Roadmap<br>(0.60%) | 0.93 | 0.14 | 2.8e-11 | 23 | 1.8 | 1.5e-08 |
| Trans-master × 5kb<br>(4.2%) | 0.34 | 0.12 | 0.0054 | 19 | 1.8 | 2.9e-06 |
| TF × ATAC<br>(0.14%) | 0.066 | 0.082 | 0.42 | 7.1 | 0.9 | 2.5e-05 |
| TF × Roadmap<br>(0.30%) | 0.17 | 0.15 | 0.26 | 16 | 1.8 | 2.4e-06 |

**Table S58.** S-LDSC results for the combined joint analysis of SNP annotations corresponding to enhancer-related, PPI-enhancer, candidate master-regulator and PPI-master gene scores conditional on the baseline-LD+cis model. Standardized Effect sizes (*τ\**) and Enrichment (E) of the significant SNP annotations in a combined joint model comprising of marginally significant enhancer-related, PPI-enhancer, candidate master-regulator and PPI-master SNP annotations. All analysis are conditional on 113 baseline-LD+cis annotations. Results are meta-analyzed across 11 blood-related traits.

| Annotation | $\tau^*$ | se($\tau^*$) | p($\tau^*$) | $E$ | se ($E$) | p($E$) |
| --- | --- | --- | --- | --- | --- | --- |
| ATAC-distal × Promoter<br>(1.8%) | 0.25 | 0.07 | 0.0003 | 6.4 | 0.53 | 2.3e-05 |
| EDS-binary × 100kb<br>(14.6%) | 0.077 | 0.019 | 4.3e-05 | 2.5 | 0.088 | 1.4e-10 |
| PPI-enhancer × ABC<br>(0.58%) | 0.99 | 0.23 | 1.2e-05 | 18 | 2.2 | 6.7e-08 |
| PPI-enhancer × Coding<br>(0.24%) | 0.57 | 0.13 | 1.3e-05 | 17 | 2.2 | 9.1e-06 |
| Trans-master × Roadmap<br>(0.58%) | 0.51 | 0.14 | 0.0003 | 20 | 1.4 | 7.1e-08 |
| Trans-master × PC-HiC<br>(5.1%) | 0.25 | 0.049 | 2.5e-07 | 4.8 | 0.17 | 6.6e-09 |
| PPI-master × ATAC<br>(0.25%) | 0.61 | 0.13 | 2.3e-06 | 20 | 2.7 | 2e-07 |
| PPI-master × Roadmap<br>(0.60%) | 0.91 | 0.12 | 7.1e-14 | 22 | 1.5 | 2.3e-08 |

**Table S59.**
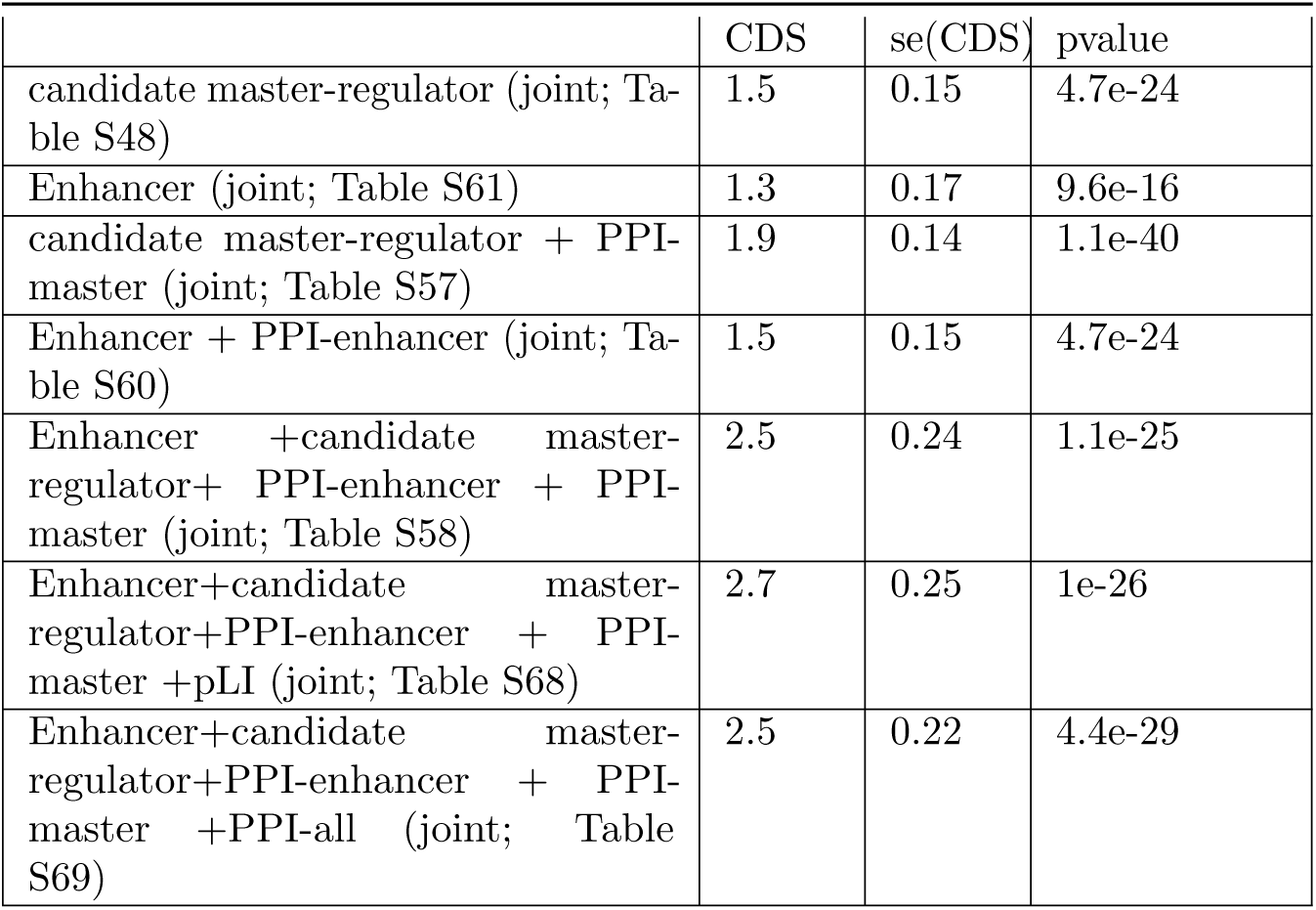
Combined *τ^*^* of various joint models. We report combined *τ^*^* scores (as well as standard errors, t statistic and p-value) for the several joint models studied.

|  | CDS | se(CDS) | pvalue |
| --- | --- | --- | --- |
| candidate master-regulator (joint; Table S48) | 1.5 | 0.15 | 4.7e-24 |
| Enhancer (joint; Table S61) | 1.3 | 0.17 | 9.6e-16 |
| candidate master-regulator + PPI-master (joint; Table S57) | 1.9 | 0.14 | 1.1e-40 |
| Enhancer + PPI-enhancer (joint; Table S60) | 1.5 | 0.15 | 4.7e-24 |
| Enhancer + candidate master-regulator + PPI-enhancer + PPI-master (joint; Table S58) | 2.5 | 0.24 | 1.1e-25 |
| Enhancer + candidate master-regulator + PPI-enhancer + PPI-master + pLI (joint; Table S68) | 2.7 | 0.25 | 1e-26 |
| Enhancer + candidate master-regulator + PPI-enhancer + PPI-master + PPI-all (joint; Table S69) | 2.5 | 0.22 | 4.4e-29 |

**Table S60.**
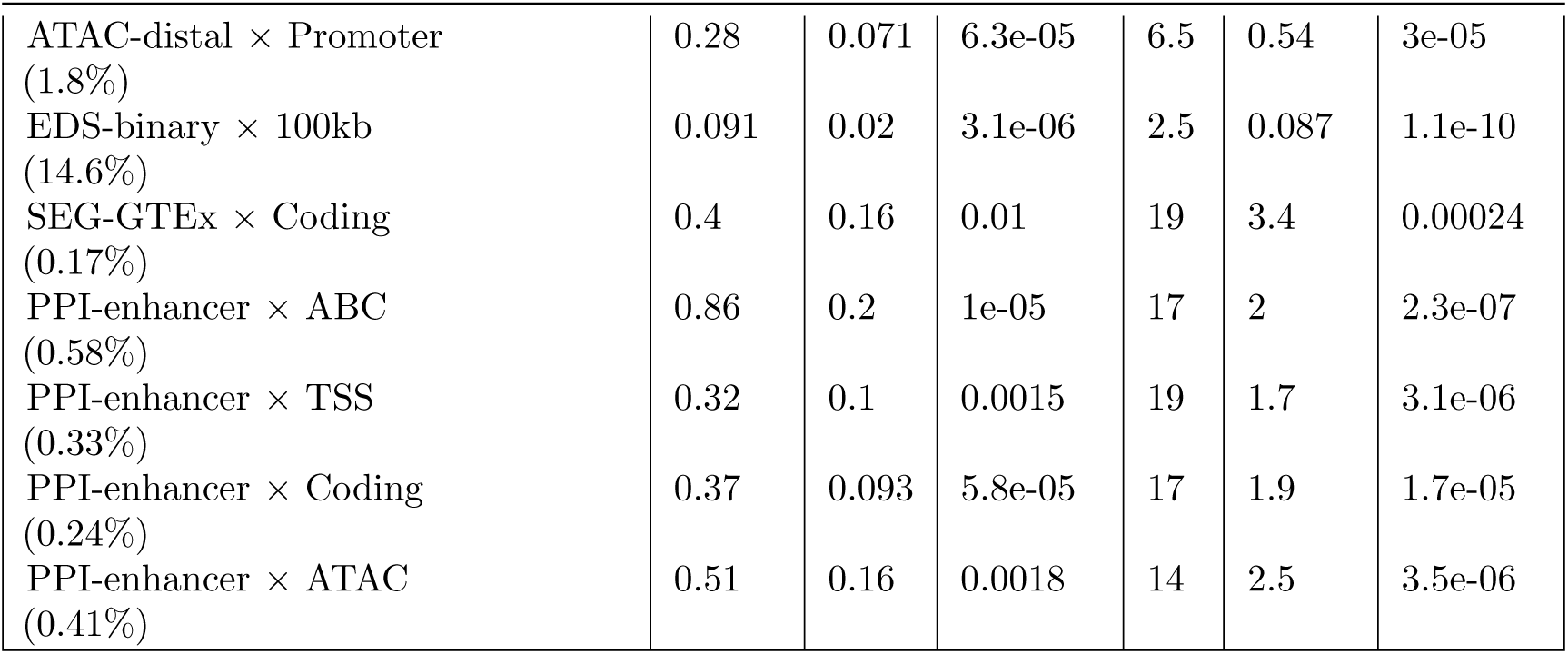
S-LDSC results for a joint analysis of SNP annotations corresponding to enhancer-related and PPI-enhancer gene scores conditional on baseline-LD+cis model. Standardized Effect sizes (*τ\**) and Enrichment (E) of the significant SNP annotations in the PPI-enhancer-related joint model in Table S26 conditional on 113 baseline-LD+cis annotations. Results are meta-analyzed across 11 blood-related traits.

**Table S61.** S-LDSC results for a joint analysis of all SNP annotations corresponding to enhancer-related gene scores conditional on baseline-LD+cis model. Standardized Effect sizes (*τ\**) and Enrichment (E) of the significant SNP annotations in the Enhancer driven joint model in Table S16 conditional on 113 baseline-LD+cis annotations. Results are meta-analyzed across 11 blood-related traits.

| Annotation | $\tau^*$ | se($\tau^*$) | p($\tau^*$) | $E$ | se ($E$) | p($E$) |
| --- | --- | --- | --- | --- | --- | --- |
| ABC-G × TSS<br>(0.38%) | 0.4 | 0.11 | 0.00021 | 17 | 1.7 | 4.5e-06 |
| ATAC-distal × Promoter<br>(1.84%) | 0.29 | 0.071 | 3.2e-05 | 6.7 | 0.54 | 2.3e-05 |
| EDS-binary × Roadmap<br>(1.15%) | 0.28 | 0.093 | 0.003 | 8.4 | 0.88 | 5e-09 |
| EDS-binary × 100kb<br>(14.6%) | 0.088 | 0.02 | 1.1e-05 | 2.5 | 0.087 | 1.2e-10 |
| SEG-GTEx × ABC<br>(0.40%) | 0.33 | 0.13 | 0.011 | 16 | 1.7 | 2.7e-06 |
| SEG-GTEx × Coding<br>(0.17%) | 0.5 | 0.16 | 0.0016 | 18 | 3.4 | 0.00034 |

**Table S62.** Δlog*l_SS_* results for the different heritability models. We report Δlog*l_SS_* derived from the log*l*_SS_ metric, proposed in ref.^64^, for the different heritability models studied in this paper: baseline-LD, baseline-LD+, baseline-LD+Enhancer (86 baseline-LD annotations and the 6 jointly significant annotations from Supplementary Figure S11), baseline-LD+PPI-enhancer (86 baseline-LD annotations and 7 jointly significant annotations from Figure 3D), baseline-LD+cis, baseline-LD+master (113 baseline-LD+cis and 4 jointly significant annotations from Supplementary Figure S14), baseline-LD+PPI-master (113 baseline-LD+cis and 4 jointly significant annotations from Figure 4D) and baseline-LD+final (113 baseline-LD+cis and 8 jointly significant annotations from Figure 5). We compute Δlog*l_SS_* as the difference in log*l*_SS_ for each model with respect to s baselineLD-no-funct model with 17 annotations that include no functional annotations^42, 64^. We also report the percentage increase in Δlog*l_SS_* for each model over the baseline-LD model. We do not report AIC as the number of annotations are not too different to alter conclusions based on just the log*l*_SS_. We report three summary log*l*_SS_ results - one averaged across 30 UK Biobank traits^42^ (All), one averaged across 6 blood-related traits from UK Biobank (Blood) and one averaged across the other 24 non blood related traits from UK Biobank (Non-blood) (Table S63). Averaged across all traits, we observe a +22.4% improvement from the annotations of the final joint model (baseline-LD+final) over baseline-LD. Of this 22.4%, we observe a +5.1% larger improvement in this metric from the 7 new S2G annotations constituting baseline-LD+ and +13% larger improvement from the 7 new S2G, plus the 20 annotations in baseline-LD+cis model. The remainder of the improvement (22.4% - 13% = 9.4%) comes from the 8 jointly significant annotations in the final joint model in Figure 5. The percentage increase was higher for the non-blood-related traits case as the log*l*_SS_ values, on average, were considerably smaller in comparison to the blood related traits.

| Model | $\Delta\log l_{SS}$<br>(All) | % incr.<br>over<br>baseline-LD<br>(All) | pval<br>(All) | $\Delta\log l_{SS}$<br>(Blood) | % incr.<br>over<br>baseline-LD<br>(Blood) | pval<br>(Blood) | $\Delta\log l_{SS}$<br>(Non-blood) | % incr.<br>over<br>baseline-LD<br>(Non-blood) | pval<br>(Non-blood) |
| --- | --- | --- | --- | --- | --- | --- | --- | --- | --- |
| baseline-LD<br>(n=86) | 1379 | 0 | - | 2668 | 0 | - | 1121 | 0 | - |
| baseline-LD+<br>(n=93) | 1449 | 5.1% | 5e-27 | 2879 | 7.8% | 1e-86 | 1163 | 3.7% | 3e-115 |
| baseline-LD+<br>Enhancer (n=99) | 1507 | 9.3% | 3e-47 | 3081 | 15% | 5e-186 | 1193 | 6.3% | 8e-24 |
| baseline-LD+<br>PPI-enhancer (n=100) | 1537 | 11.4% | 5e-59 | 3208 | 20% | 1e-220 | 1202 | 7.2% | 2e-27 |
| baseline-LD+cis<br>(n=113) | 1564 | 13.4% | 6e-62 | 2532 | -5.0% | 1.00 | 1370 | 22.2% | 3e-88 |
| baseline-LD+<br>Master (n=117) | 1669 | 21% | 2e-102 | 2952 | 10.6% | 1e-99 | 1412 | 25.9% | 1e-102 |
| baseline-LD+<br>PPI-master (n=117) | 1681 | 21.8% | 2e-107 | 2986 | 11.9% | 9e-114 | 1419 | 26.6% | 9e-106 |
| baseline-LD+final<br>(n=121) | 1688 | 22.4% | 9e-108 | 2998 | 12.3% | 3e-116 | 1425 | 27.2% | 9e-106 |

**Table S63.** List of UKBiobank traits used for log*l*_SS_ calculations. The list consists of 6 blood-related traits and 24 non blood-related traits.

| Trait | Category | N |
| --- | --- | --- |
| disease AID Sure | Blood | 459324 |
| blood Eosinophil count | Blood | 439938 |
| Blood Platelet count | Blood | 444382 |
| Blood Red count | Blood | 445174 |
| Blood RBC distr. width | Blood | 442700 |
| blood White count | Blood | 444502 |
| reproduction Menarche Age | Non-blood | 242278 |
| reproduction Menopause Age | Non-blood | 143025 |
| Body balding | Non-blood | 208336 |
| Body BMIz | Non-blood | 457824 |
| cov EDU Years | Non-blood | 454813 |
| disease Dermatology | Non-blood | 459324 |
| disease Allergy Eczema | Non-blood | 458699 |
| lung FVCzSmoke | Non-blood | 371949 |
| lung FEV1FVCzSmoke | Non-blood | 371949 |
| pigment Hair | Non-blood | 452720 |
| bmd Heel Tscorez | Non-blood | 445921 |
| body Heightz | Non-blood | 458303 |
| disease Hi-chol-self-rep | Non-blood | 459324 |
| disease Hypothyroidism self rep | Non-blood | 459324 |
| Other Morning-person | Non-blood | 41050 |
| Mental Neuroticism | Non-blood | 372066 |
| disease Respiratory ENT | Non-blood | 459324 |
| pigment Skin | Non-blood | 453609 |
| cov Smoking Status | Non-blood | 457683 |
| pigment Sunburn | Non-blood | 344229 |
| bp SystolicadjMedz | Non-blood | 422771 |
| pigment Tanning | Non-blood | 449984 |
| disease T2D | Non-blood | 459324 |
| body WHRadjBMIz | Non-blood | 458417 |

**Table S64.** List of 41 diseases and complex traits analyzed. We first list the 11 blood-related traits (6 autoimmune diseases and 5 blood cell traits), followed by 10 brain traits (with 2 traits analyzed by two different datasets). Overall, for 6 traits we analyzed two different data sets, leading to a total of 47 data sets.

| Trait | Source | N |
| --- | --- | --- |
| Platelet Count | UKBiobank<sup>35</sup> | 444382 |
| Red Blood Cell Count | UKBiobank<sup>35</sup> | 445174 |
| Red Blood Cell Distribution Width | UKBiobank<sup>35</sup> | 442700 |
| Eosinophil Count | UKBiobank<sup>35</sup> | 439938 |
| White Blood Cell Count | UKBiobank<sup>35</sup> | 444502 |
| Auto Immune Traits (Sure) | UKBiobank<sup>35</sup> | 459324 |
| Crohn's Disease | Jostins et al., 2012 Nature<sup>113</sup> | 20883 |
| Rheumatoid Arthritis | Okada et al., 2014 Nature<sup>114</sup> | 37681 |
| Ulcerative Colitis | Jostins et al., 2012 Nature<sup>113</sup> | 27432 |
| Lupus | Bentham et al., 2015<sup>115</sup> | 14267 |
| Celiac | Dubois et al., 2010<sup>116</sup> | 15283 |
| Age at Menarche | UKBiobank<sup>35</sup> | 242278 |
| BMI | Speliotes et al., 2010 Nat Genet<sup>118</sup> | 122033 |
| BMI | UKBiobank<sup>35</sup> | 457824 |
| Depressive symptoms | Okbay et al., 2016 Nat Genet<sup>119</sup> | 161460 |
| Neuroticism | UKBiobank<sup>35</sup> | 372066 |
| Schizophrenia | SCZ Working Group of the PGC, 2014 Nature<sup>120</sup> | 70100 |
| Years of Education | Okbay et al., 2016 Nature<sup>119</sup> | 328917 |
| Ever Smoked | TAG Consortium, 2010 Nat Genet<sup>121</sup> | 74035 |
| Smoking Status | UKBiobank<sup>35</sup> | 457683 |
| Bipolar Disorder | PGC Bipolar Disorder Group<sup>122</sup> | 16731 |
| Age at Menopause | UKBiobank<sup>35</sup> | 143025 |
| Age first birth | Barban et al., 2016 Nat Genet<sup>123</sup> | 222037 |
| Anorexia | Boraska et al., 2014 Mol Psych<sup>124</sup> | 32143 |
| Autism Spectrum | PGC Cross-Disorder Group, 2013 Lancet<sup>125</sup> | 10263 |
| College Education | UKBiobank<sup>35</sup> | 454813 |
| Coronary Artery Disease | Schunkert et al., 2011 Nat Genet<sup>126</sup> | 77210 |
| Dermatologic Diseases | UKBiobank<sup>35</sup> | 459324 |
| Eczema | UKBiobank<sup>35</sup> | 458699 |
| FEV1-FVC Ratio | UKBiobank<sup>35</sup> | 371949 |
| Forced Vital Capacity (FVC) | UKBiobank<sup>35</sup> | 371949 |
| Hair Color | UKBiobank<sup>35</sup> | 452720 |
| HDL | Teslovich et al., 2010 Nature<sup>127</sup> | 97749 |
| Heel T Score | UKBiobank<sup>35</sup> | 445921 |
| Height | Lango Allen et al., 2010 Nature<sup>128</sup> | 131547 |
| Height | UKBiobank<sup>35</sup> | 458303 |
| High Cholesterol | UKBiobank<sup>35</sup> | 459324 |
| Hypothyroidism | UKBiobank<sup>35</sup> | 459324 |
| LDL | Teslovich et al., 2010 Nature<sup>127</sup> | 93354 |
| Morning Person | UKBiobank<sup>35</sup> | 410520 |
| Number children ever born | Barban et al., 2016 Nat Genet<sup>123</sup> | 318863 |
| Respiratory and Ear-nose-throat Diseases | UKBiobank<sup>35</sup> | 459324 |
| Skin Color | UKBiobank<sup>35</sup> | 453609 |
| Sunburn Occasion | UKBiobank<sup>35</sup> | 344229 |
| Tanning | UKBiobank<sup>35</sup> | 449984 |
| Type 2 Diabetes | Morris et al., 2012 Nat Genet<sup>129</sup> | 60786 |
| Type 2 Diabetes | UKBiobank<sup>35</sup> | 459324 |

**Table S65.**
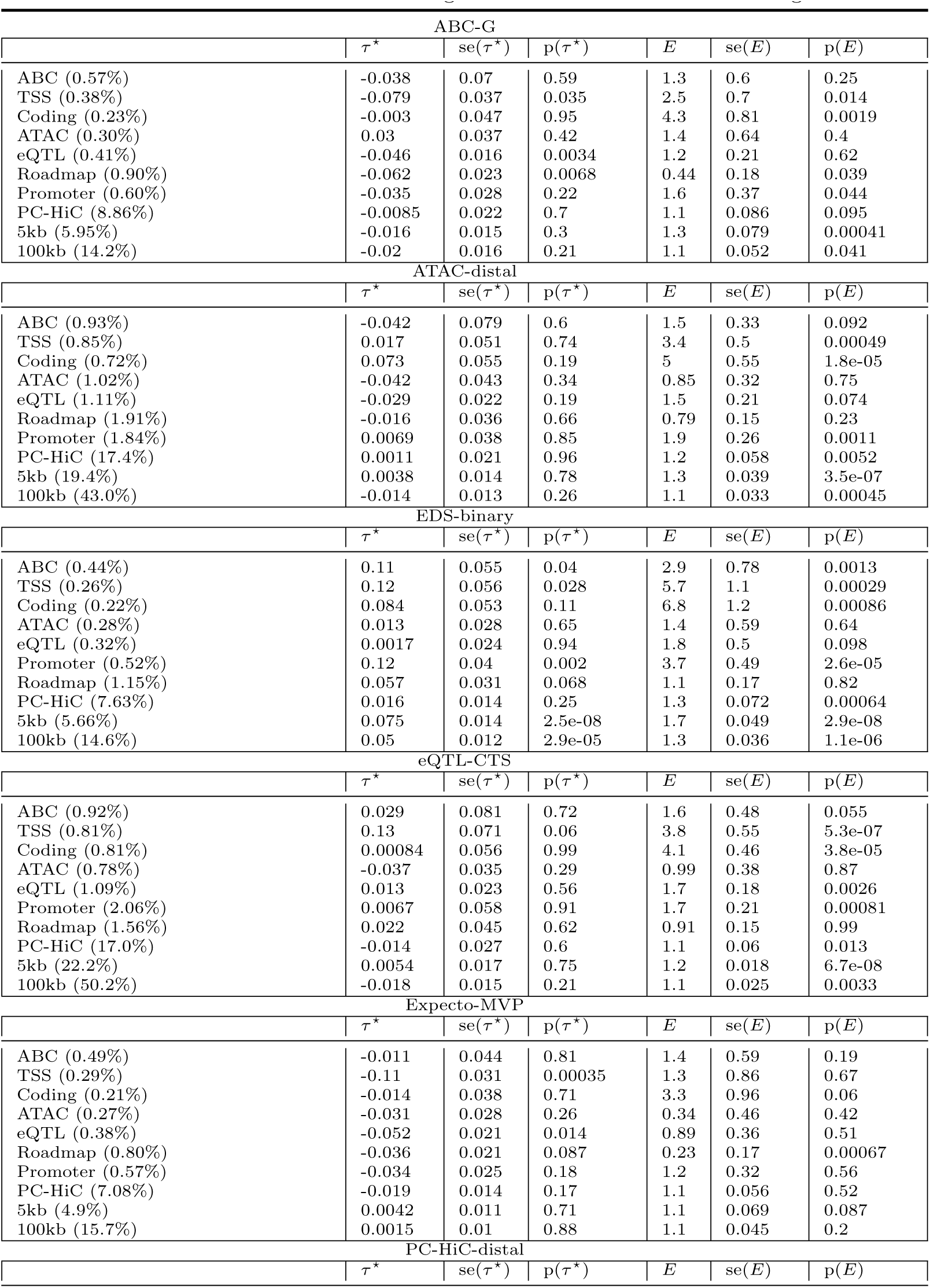
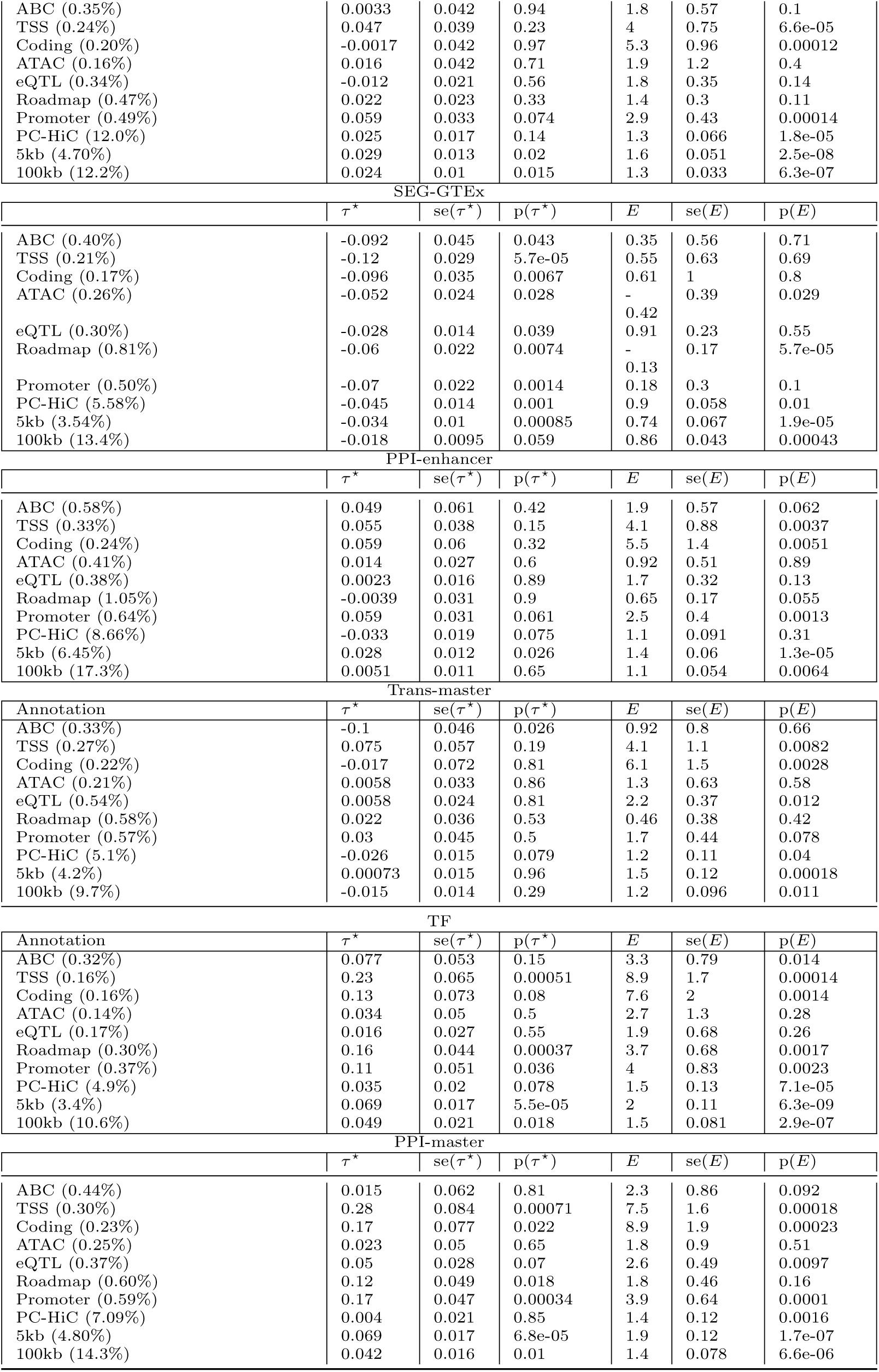
S-LDSC results for SNP annotations corresponding to enhancer-related and candidate master-regulator gene scores meta-analyzed across 8 brain-related traits: Standardized Effect sizes (*τ\**) and Enrichment (E) of SNP annotations corresponding to all enhancer-related and candidate master-regulator gene scores and 10 S2G strategies, meta-analyzed across 8 relatively independent brain-related traits. Results are conditional on 93 baseline-LD+ annotations for the 7 enhancer- related genes and 1 PPI-enhancer gene score, and conditional on 113 baseline-LD+cis annotations for the 2 candidate master-regulator scores and 1 PPI-master gene score.

**Table S66.** S-LDSC results for SNP annotations corresponding to enhancer-related and candidate master-regulator gene scores meta-analyzed across 28 non-blood and non-brain related traits: Standardized Effect sizes (*τ\**) and Enrichment (E) of SNP annotations corresponding to all enhancer-related and candidate master-regulator gene scores and 10 S2G strategies, meta-analyzed across 28 relatively independent non-blood and non-brain related traits. Results are conditional on 93 baseline-LD+ annotations for the 7 enhancer-related genes and 1 PPI-enhancer gene score, and conditional on 113 baseline-LD+cis annotations for the 2 candidate master-regulator scores and 1 PPI-master gene score.

| ABC-G |  |  |  |  |  |  |
| --- | --- | --- | --- | --- | --- | --- |
| | $\tau^*$ | se($\tau^*$) | p($\tau^*$) | $E$ | se($E$) | p($E$) |
| ABC (0.57%) | -0.022 | 0.056 | 0.7 | 2.2 | 0.66 | 0.026 |
| TSS (0.38%) | 0.058 | 0.044 | 0.19 | 6.2 | 0.91 | 8.1e-08 |
| Coding (0.23%) | 0.072 | 0.041 | 0.079 | 6.3 | 0.85 | 6.7e-06 |
| ATAC (0.30%) | 0.028 | 0.037 | 0.44 | 2.4 | 0.65 | 0.028 |
| eQTL (0.41%) | -0.027 | 0.015 | 0.076 | 2 | 0.29 | 0.0035 |
| Roadmap (0.90%) | -0.029 | 0.029 | 0.32 | 1.6 | 0.39 | 0.095 |
| Promoter (0.60%) | 0.043 | 0.032 | 0.17 | 3.1 | 0.41 | 1.1e-06 |
| PC-HiC (8.86%) | 0.018 | 0.019 | 0.33 | 1.7 | 0.097 | 1.9e-12 |
| 5kb (5.95%) | 0.0027 | 0.016 | 0.86 | 1.9 | 0.1 | 1.2e-16 |
| 100kb (14.2%) | 0.0043 | 0.015 | 0.78 | 1.6 | 0.074 | 6.8e-18 |
| ATAC-distal |  |  |  |  |  |  |
| | $\tau^*$ | se($\tau^*$) | p($\tau^*$) | $E$ | se($E$) | p($E$) |
| ABC (0.93%) | 0.25 | 0.13 | 0.044 | 3.2 | 0.66 | 0.00037 |
| TSS (0.85%) | 0.19 | 0.073 | 0.011 | 6.3 | 0.68 | 2.4e-11 |
| Coding (0.72%) | 0.12 | 0.069 | 0.091 | 5.8 | 0.63 | 3.6e-08 |
| ATAC (1.02%) | 0.0052 | 0.066 | 0.94 | 1.9 | 0.45 | 0.014 |
| eQTL (1.11%) | -0.0096 | 0.021 | 0.65 | 2.1 | 0.21 | 6.9e-07 |
| Roadmap (1.91%) | 0.059 | 0.062 | 0.35 | 2.2 | 0.37 | 0.0035 |
| Promoter (1.84%) | 0.11 | 0.051 | 0.027 | 3.1 | 0.34 | 1.1e-08 |
| PC-HiC (17.4%) | 0.03 | 0.027 | 0.25 | 1.6 | 0.076 | 1.8e-15 |
| 5kb (19.4%) | 0.017 | 0.013 | 0.2 | 1.6 | 0.054 | 3.8e-22 |
| 100kb (43.0%) | 0.012 | 0.012 | 0.34 | 1.4 | 0.044 | 5.3e-25 |
| EDS-binary |  |  |  |  |  |  |
| | $\tau^*$ | se($\tau^*$) | p($\tau^*$) | $E$ | se($E$) | p($E$) |
| ABC (0.44%) | 0.22 | 0.044 | 5.6e-07 | 4.7 | 0.8 | 4.4e-06 |
| TSS (0.26%) | 0.27 | 0.05 | 3.1e-08 | 10 | 1.1 | 5e-10 |
| Coding (0.22%) | 0.21 | 0.048 | 8.4e-06 | 10 | 1.1 | 2.1e-07 |
| ATAC (0.28%) | 0.13 | 0.041 | 0.0011 | 4.2 | 0.81 | 0.00012 |
| eQTL (0.32%) | 0.026 | 0.016 | 0.11 | 2.6 | 0.32 | 0.00024 |
| Promoter (0.52%) | 0.2 | 0.032 | 5.6e-10 | 5.3 | 0.47 | 1.9e-07 |
| Roadmap (1.15%) | 0.14 | 0.041 | 0.00083 | 2.4 | 0.38 | 7.7e-05 |
| PC-HiC (7.63%) | 0.025 | 0.022 | 0.25 | 1.7 | 0.11 | 5.8e-11 |
| 5kb (5.66%) | 0.037 | 0.015 | 0.011 | 1.8 | 0.07 | 3.7e-17 |
| 100kb (14.6%) | 0.066 | 0.012 | 4.2e-08 | 1.6 | 0.064 | 1.3e-21 |
| eQTL-CTS |  |  |  |  |  |  |
| | $\tau^*$ | se($\tau^*$) | p($\tau^*$) | $E$ | se($E$) | p($E$) |
| ABC (0.92%) | 0.011 | 0.074 | 0.89 | 2.3 | 0.47 | 0.0015 |
| TSS (0.81%) | -0.013 | 0.048 | 0.79 | 5 | 0.45 | 2.9e-09 |
| Coding (0.81%) | -0.049 | 0.052 | 0.35 | 4.5 | 0.42 | 4.5e-06 |
| ATAC (0.78%) | -0.0049 | 0.038 | 0.9 | 1.9 | 0.39 | 0.022 |
| eQTL (1.09%) | -0.0083 | 0.027 | 0.76 | 2 | 0.15 | 1.3e-07 |
| Promoter (2.06%) | 0.01 | 0.032 | 0.75 | 2.2 | 0.19 | 2.8e-06 |
| Roadmap (1.56%) | 0.0095 | 0.034 | 0.78 | 1.7 | 0.3 | 0.0064 |
| PC-HiC (17.0%) | 0.0067 | 0.022 | 0.76 | 1.5 | 0.071 | 1.2e-13 |
| 5kb (22.2%) | 0.0021 | 0.011 | 0.84 | 1.4 | 0.034 | 2.6e-18 |
| 100kb (50.2%) | -0.025 | 0.012 | 0.031 | 1.3 | 0.03 | 5.1e-21 |
| Expecto-MVP |  |  |  |  |  |  |
| | $\tau^*$ | se($\tau^*$) | p($\tau^*$) | $E$ | se($E$) | p($E$) |
| ABC (0.49%) | 0.03 | 0.043 | 0.49 | 2.5 | 0.63 | 0.0012 |
| TSS (0.29%) | -0.084 | 0.045 | 0.062 | 3.3 | 0.95 | 0.011 |
| Coding (0.21%) | -0.015 | 0.036 | 0.67 | 4.6 | 1 | 0.0089 |
| ATAC (0.27%) | -0.041 | 0.033 | 0.22 | 1.1 | 0.68 | 0.77 |
| eQTL (0.38%) | -0.023 | 0.02 | 0.25 | 1.7 | 0.35 | 0.055 |
| Roadmap (0.80%) | -0.041 | 0.035 | 0.24 | 0.78 | 0.42 | 0.55 |
| Promoter (0.57%) | -0.036 | 0.024 | 0.14 | 1.9 | 0.41 | 0.011 |
| PC-HiC (7.08%) | 0.014 | 0.016 | 0.36 | 1.5 | 0.1 | 1.7e-07 |
| 5kb (4.9%) | -0.023 | 0.015 | 0.13 | 1.5 | 0.087 | 9.4e-08 |
| 100kb (15.7%) | -0.014 | 0.012 | 0.24 | 1.4 | 0.063 | 6e-11 |
| PC-HiC-distal |  |  |  |  |  |  |

| | $\tau^*$ | $se(\tau^*)$ | $p(\tau^*)$ | $E$ | $se(E)$ | $p(E)$ |
| --- | --- | --- | --- | --- | --- | --- |
| ABC (0.35%) | 0.064 | 0.038 | 0.092 | 3.6 | 0.74 | 0.00016 |
| TSS (0.24%) | 0.14 | 0.045 | 0.0023 | 8 | 1 | 7.8e-08 |
| Coding (0.20%) | 0.064 | 0.042 | 0.12 | 6.9 | 0.92 | 2.1e-05 |
| ATAC (0.16%) | 0.036 | 0.029 | 0.23 | 3.5 | 0.67 | 2.8e-05 |
| eQTL (0.34%) | 0.011 | 0.016 | 0.49 | 2.4 | 0.31 | 0.0002 |
| Roadmap (0.47%) | 0.075 | 0.027 | 0.0058 | 3 | 0.53 | 1.4e-05 |
| Promoter (0.49%) | 0.097 | 0.03 | 0.0011 | 4 | 0.43 | 3e-07 |
| PC-HiC (12.0%) | 0.038 | 0.014 | 0.009 | 1.6 | 0.072 | 3.9e-16 |
| 5kb (4.70%) | 0.029 | 0.01 | 0.004 | 1.8 | 0.074 | 5.5e-16 |
| 100kb (12.2%) | 0.024 | 0.0094 | 0.012 | 1.6 | 0.058 | 1.6e-18 |

| SEG-GTE<sub>x</sub> |  |  |  |  |  |  |
| --- | --- | --- | --- | --- | --- | --- |
| | $\tau^*$ | $se(\tau^*)$ | $p(\tau^*)$ | $E$ | $se(E)$ | $p(E)$ |
| ABC (0.40%) | -0.017 | 0.039 | 0.67 | 1.7 | 0.72 | 0.13 |
| TSS (0.21%) | -0.058 | 0.039 | 0.14 | 3.6 | 0.93 | 0.0014 |
| Coding (0.17%) | 0.0015 | 0.035 | 0.97 | 4.2 | 1.1 | 0.033 |
| ATAC (0.26%) | -0.068 | 0.03 | 0.022 | 0.12 | 0.63 | 0.19 |
| eQTL (0.30%) | -0.04 | 0.015 | 0.0062 | 1.2 | 0.32 | 0.39 |
| Roadmap (0.81%) | -0.08 | 0.029 | 0.0053 | 0.56 | 0.39 | 0.22 |
| Promoter (0.50%) | -0.017 | 0.021 | 0.42 | 1.7 | 0.42 | 0.1 |
| PC-HiC (5.58%) | -0.062 | 0.019 | 0.0011 | 1.3 | 0.11 | 0.0081 |
| 5kb (3.54%) | -0.017 | 0.01 | 0.098 | 1.4 | 0.12 | 0.0041 |
| 100kb (13.4%) | -0.0012 | 0.011 | 0.92 | 1.3 | 0.073 | 1.1e-06 |

| PPI-enhancer |  |  |  |  |  |  |
| --- | --- | --- | --- | --- | --- | --- |
| | $\tau^*$ | $se(\tau^*)$ | $p(\tau^*)$ | $E$ | $se(E)$ | $p(E)$ |
| ABC (0.58%) | 0.3 | 0.083 | 0.00027 | 4.8 | 0.88 | 6.4e-06 |
| TSS (0.33%) | 0.36 | 0.065 | 2.6e-08 | 11 | 1.2 | 2e-13 |
| Coding (0.24%) | 0.27 | 0.075 | 0.00026 | 10 | 1.7 | 5.9e-07 |
| ATAC (0.41%) | 0.14 | 0.047 | 0.0029 | 3.6 | 0.74 | 7.6e-05 |
| eQTL (0.38%) | 0.024 | 0.021 | 0.26 | 2.4 | 0.36 | 0.00027 |
| Roadmap (1.05%) | 0.15 | 0.051 | 0.0035 | 2.7 | 0.48 | 0.00019 |
| Promoter (0.64%) | 0.26 | 0.053 | 1.6e-06 | 5.6 | 0.76 | 2e-08 |
| PC-HiC (8.66%) | 0.047 | 0.023 | 0.04 | 1.7 | 0.12 | 4.4e-12 |
| 5kb (6.45%) | 0.036 | 0.018 | 0.044 | 1.8 | 0.099 | 7.7e-16 |
| 100kb (17.3%) | 0.086 | 0.019 | 8.1e-06 | 1.7 | 0.086 | 1.7e-20 |

| Trans-master |  |  |  |  |  |  |
| --- | --- | --- | --- | --- | --- | --- |
| Annotation | $\tau^*$ | $se(\tau^*)$ | $p(\tau^*)$ | $E$ | $se(E)$ | $p(E)$ |
| ABC (0.33%) | 0.23 | 0.081 | 0.0036 | 5.6 | 1.3 | 0.00019 |
| TSS (0.27%) | 0.33 | 0.079 | 3.3e-05 | 11 | 1.5 | 2e-10 |
| Coding (0.22%) | 0.31 | 0.082 | 0.00014 | 8.3 | 0.99 | 2.3e-09 |
| ATAC (0.21%) | -0.0014 | 0.05 | 0.98 | 1.7 | 0.89 | 0.2 |
| eQTL (0.54%) | 0.055 | 0.032 | 0.083 | 3.5 | 0.39 | 4e-10 |
| Roadmap (0.58%) | 0.078 | 0.043 | 0.073 | 2 | 0.69 | 0.12 |
| Promoter (0.57%) | 0.18 | 0.052 | 0.00065 | 4.8 | 0.66 | 2.9e-09 |
| PC-HiC (5.1%) | 0.039 | 0.024 | 0.1 | 2.1 | 0.16 | 1.8e-12 |
| 5kb (4.2%) | 0.084 | 0.021 | 4.4e-05 | 2.5 | 0.17 | 1.8e-19 |
| 100kb (9.7%) | 0.074 | 0.02 | 0.00025 | 2.1 | 0.13 | 9.3e-21 |

| TF |  |  |  |  |  |  |
| --- | --- | --- | --- | --- | --- | --- |
| Annotation | $\tau^*$ | $se(\tau^*)$ | $p(\tau^*)$ | $E$ | $se(E)$ | $p(E)$ |
| ABC (0.32%) | 0.24 | 0.073 | 0.00093 | 6.3 | 1.2 | 5.5e-06 |
| TSS (0.16%) | 0.37 | 0.062 | 3.3e-09 | 14 | 1.7 | 8.5e-10 |
| Coding (0.16%) | 0.28 | 0.064 | 1.5e-05 | 11 | 1.6 | 3.4e-07 |
| ATAC (0.14%) | 0.17 | 0.042 | 8.3e-05 | 8 | 1.3 | 1.9e-08 |
| eQTL (0.17%) | 0.015 | 0.022 | 0.5 | 2.1 | 0.5 | 0.13 |
| Roadmap (0.30%) | 0.22 | 0.047 | 3.4e-06 | 5.5 | 0.89 | 3.4e-08 |
| Promoter (0.37%) | 0.25 | 0.045 | 2.3e-08 | 6.7 | 0.72 | 8.1e-09 |
| PC-HiC (4.9%) | 0.087 | 0.021 | 4.7e-05 | 2 | 0.13 | 7.2e-14 |
| 5kb (3.4%) | 0.13 | 0.016 | 1.3e-16 | 2.4 | 0.1 | 9.7e-20 |
| 100kb (10.6%) | 0.11 | 0.018 | 9.2e-10 | 1.8 | 0.077 | 2.1e-22 |

| PPI-master |  |  |  |  |  |  |
| --- | --- | --- | --- | --- | --- | --- |
| | $\tau^*$ | $se(\tau^*)$ | $p(\tau^*)$ | $E$ | $se(E)$ | $p(E)$ |
| ABC (0.44%) | 0.32 | 0.099 | 0.0012 | 6.3 | 1.2 | 7.5e-06 |
| TSS (0.30%) | 0.7 | 0.092 | 2.6e-14 | 16 | 1.7 | 2.5e-15 |
| Coding (0.23%) | 0.53 | 0.099 | 1.2e-07 | 15 | 1.9 | 1.2e-10 |
| ATAC (0.25%) | 0.25 | 0.066 | 0.00013 | 6.4 | 1.3 | 1.9e-05 |
| eQTL (0.37%) | 0.059 | 0.024 | 0.015 | 3.6 | 0.42 | 1.5e-08 |
| Roadmap (0.60%) | 0.27 | 0.056 | 1.6e-06 | 4.4 | 0.77 | 1.4e-06 |
| Promoter (0.59%) | 0.39 | 0.058 | 1.4e-11 | 7.3 | 0.69 | 7.8e-14 |
| PC-HiC (7.09%) | 0.099 | 0.024 | 3.4e-05 | 2.1 | 0.13 | 5.4e-17 |
| 5kb (4.80%) | 0.15 | 0.02 | 7.2e-14 | 2.7 | 0.15 | 2.2e-24 |
| 100kb (14.3%) | 0.14 | 0.022 | 7.5e-10 | 2 | 0.1 | 2.5e-25 |

**Table S67.**
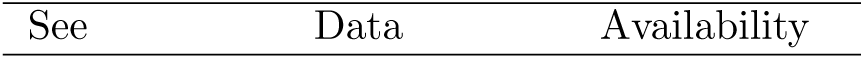
List of 194 fine-mapped SNPs belonging to Trans-master *×* Roadmap, PPI-enhancer *×* ABC, PPI-master *×* ATAC, and/or PPI-master *×* Roadmap annotations: Fine-mapped SNPs with PIP *>* 0.90 in UKBB All autoimmune traits, or either of the 5 blood cell traits belonging to Trans-master *×* Roadmap, PPI-enhancer *×* ABC, PPI-master *×* ATAC, and/or PPI-master *×* Roadmap annotations, annotations are regulatory and highly enriched (*≥* 18) in the combined joint model (Figure 5). The number of unique fine-mapped SNPs is 194 and the number of unique SNP-gene pairs for traits implicated by our analysis is 273.

**Table S68.**
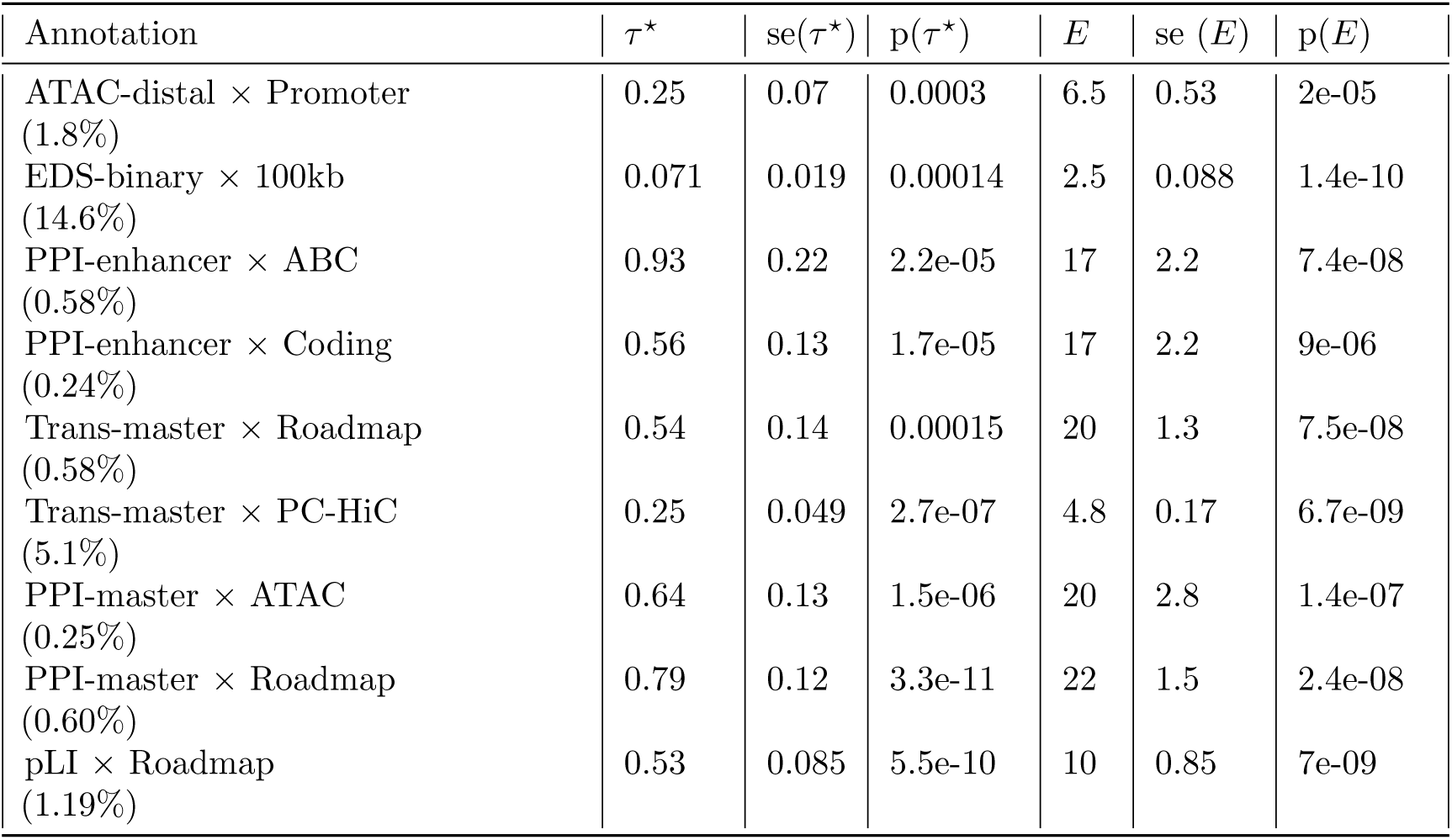
S-LDSC results for the combined joint model of all SNP annotations corresponding to enhancer-related, PPI-enhancer, candidate master-regulator, PPI-master and pLI gene scores conditional on baseline-LD+cis model. Standardized Effect sizes (*τ\**) and Enrichment (E) of the significant SNP annotations in a joint model comprising of enhancer-related, PPI-enhancer, candidate master-regulator, PPI-master and pLI SNP annotations. All analysis are conditional on 113 baseline-LD+cis annotations. Results are meta-analyzed across 11 blood-related traits.

| Annotation | $\tau^*$ | $se(\tau^*)$ | $p(\tau^*)$ | $E$ | $se(E)$ | $p(E)$ |
| --- | --- | --- | --- | --- | --- | --- |
| ATAC-distal × Promoter<br>(1.8%) | 0.25 | 0.07 | 0.0003 | 6.5 | 0.53 | 2e-05 |
| EDS-binary × 100kb<br>(14.6%) | 0.071 | 0.019 | 0.00014 | 2.5 | 0.088 | 1.4e-10 |
| PPI-enhancer × ABC<br>(0.58%) | 0.93 | 0.22 | 2.2e-05 | 17 | 2.2 | 7.4e-08 |
| PPI-enhancer × Coding<br>(0.24%) | 0.56 | 0.13 | 1.7e-05 | 17 | 2.2 | 9e-06 |
| Trans-master × Roadmap<br>(0.58%) | 0.54 | 0.14 | 0.00015 | 20 | 1.3 | 7.5e-08 |
| Trans-master × PC-HiC<br>(5.1%) | 0.25 | 0.049 | 2.7e-07 | 4.8 | 0.17 | 6.7e-09 |
| PPI-master × ATAC<br>(0.25%) | 0.64 | 0.13 | 1.5e-06 | 20 | 2.8 | 1.4e-07 |
| PPI-master × Roadmap<br>(0.60%) | 0.79 | 0.12 | 3.3e-11 | 22 | 1.5 | 2.4e-08 |
| pLI × Roadmap<br>(1.19%) | 0.53 | 0.085 | 5.5e-10 | 10 | 0.85 | 7e-09 |

**Table S69.** S-LDSC results for the combined joint model of all SNP annotations corresponding to enhancer-related, PPI-enhancer, candidate master-regulator, PPI-master and PPI-all gene scores conditional on baseline-LD+cis model. Standardized Effect sizes (*τ\**) and Enrichment (E) of the significant SNP annotations in a joint model comprising of enhancer-related, PPI-enhancer, candidate master-regulator, PPI-master and PPI-all SNP annotations. All analysis are conditional on 113 baseline-LD+cis annotations. Results are meta-analyzed across 11 blood-related traits.

| Annotation | $\tau^*$ | se($\tau^*$) | p($\tau^*$) | $E$ | se ($E$) | p($E$) |
| --- | --- | --- | --- | --- | --- | --- |
| ATAC-distal × Promoter<br>(1.8%) | 0.26 | 0.069 | 0.00013 | 6.4 | 0.53 | 2.3e-05 |
| EDS-binary × 100kb<br>(14.6%) | 0.087 | 0.019 | 2.5e-06 | 2.5 | 0.089 | 1.3e-10 |
| Trans-master × Roadmap<br>(0.58%) | 0.56 | 0.14 | 3.4e-05 | 20 | 1.3 | 7.2e-08 |
| Trans-master × PC-HiC<br>(5.1%) | 0.25 | 0.048 | 3.9e-07 | 4.7 | 0.17 | 5.9e-09 |
| PPI-master × ATAC<br>(0.25%) | 0.56 | 0.13 | 8e-06 | 19 | 2.7 | 3.3e-07 |
| PPI-master × Roadmap<br>(0.60%) | 0.58 | 0.13 | 4.6e-06 | 22 | 1.4 | 2.7e-08 |
| PPI-all × ABC<br>(0.53%) | 0.7 | 0.15 | 1.7e-06 | 18 | 1.9 | 1.4e-07 |
| PPI-all × Coding<br>(0.23%) | 0.56 | 0.1 | 5.2e-08 | 18 | 2.1 | 0.00012 |
| PPI-all × Roadmap<br>(0.81%) | 0.7 | 0.18 | 7.9e-05 | 19 | 1.9 | 3.6e-10 |

**Table S70.** S-LDSC results for the less restrictive combined joint model of all SNP annotations corresponding to both enhancer-related, PPI-enhancer, candidate master-regulator and PPI-master gene scores conditional on the baseline-LD+ model. Standardized Effect sizes (*τ\**) and Enrichment (E) of the significant SNP annotations in a joint model comprising of enhancer-related, PPI-enhancer, candidate master-regulator and PPI-master SNP annotations. All analysis are conditional on 113 baseline-LD+cis annotations. Results are meta-analyzed across 11 blood-related traits.

| Annotation | $\tau^*$ | se($\tau^*$) | p($\tau^*$) | $E$ | se ($E$) | p($E$) |
| --- | --- | --- | --- | --- | --- | --- |
| ATAC-distal × Promoter<br>(1.8%) | 0.42 | 0.07 | 2.5e-09 | 6.4 | 0.5 | 2e-05 |
| EDS-binary × 100kb<br>(14.6%) | 0.084 | 0.019 | 1.3e-05 | 2.5 | 0.09 | 1e-10 |
| SEG-GTEx × Coding<br>(0.17%) | 0.55 | 0.14 | 0.0003 | 20 | 3.3 | 7.6e-05 |
| PPI-enhancer × ABC<br>(0.58%) | 1.3 | 0.24 | 1.2e-07 | 18 | 2.2 | 4.5e-08 |
| PPI-enhancer × Coding<br>(0.24%) | 0.42 | 0.11 | 0.00022 | 17 | 2.2 | 6.1e-06 |
| Trans-master × Roadmap<br>(0.58%) | 0.82 | 0.16 | 1.5e-07 | 22 | 1.3 | 3.2e-08 |
| Trans-master × PC-HiC<br>(5.1%) | 0.34 | 0.037 | 1.7e-19 | 4.9 | 0.16 | 6.9e-09 |
| PPI-master × ATAC<br>(0.25%) | 0.69 | 0.14 | 4.1e-07 | 20 | 2.7 | 1.1e-07 |
| PPI-master × Roadmap<br>(0.60%) | 0.93 | 0.12 | 3.2e-14 | 23 | 1.3 | 1.3e-08 |

**Table S71.**
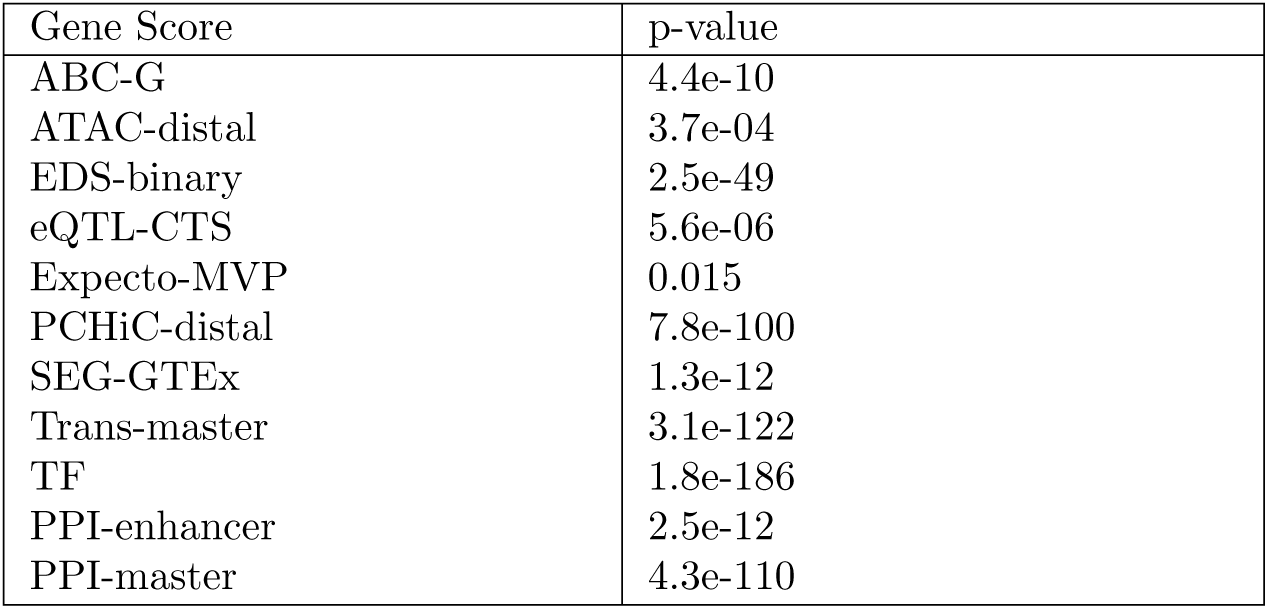
MAGMA gene-set analysis p-values for all gene scores, meta-analysed across blood-related traits: Meta-analysed p-value across 11 independent blood-related diseases and traits for binarized (taking top 10% if the actual gene score is probabilistic) enhancer-related, candidate master-regulator, PPI-enhancer and PPI-master gene scores in a MAGMA competitive gene sets analysis.

**Table S72.** List of drug target genes uniquely implicated by PPI-enhancer gene over other enhancer-regulated gene scores (based on top 10% genes in each gene score), and by PPI-master over other candidate master-regulator gene scores (based on top 10% genes in each gene score).

| Gene Score | Unique drug target genes |
| --- | --- |
| PPI-enhancer | FAS, PDE4A, ICAM1, IKBKB, ATP4A, SERPINE1, CCL2, IL12B, SPP1, CXCL10, PTEN, EPOR, IRAK4, MTOR, TLR9, SELE, ITGAE, PRKCE, RELA, HRH1, PTGIR, IFNA1 |
| PPI-master | CD4, FYN, FGFR3, PTGS2, PTGS1, JAK2, IKBKB, TGFB1, JAK3, PIK3CG, SERPINE1, IL2, IFNG, MAPK14, IL17A, IL12B, PDGFRB, IL1B, IL6, CDK9, TLR4, TLR2, KIT, JAK1, CSF2, SYK, PIK3CD, PTEN, MTOR, SMAD7, PRKCA, PRKCB, CD40LG, IL10 |

## 2 Supplementary Figures

**Figure S1.**
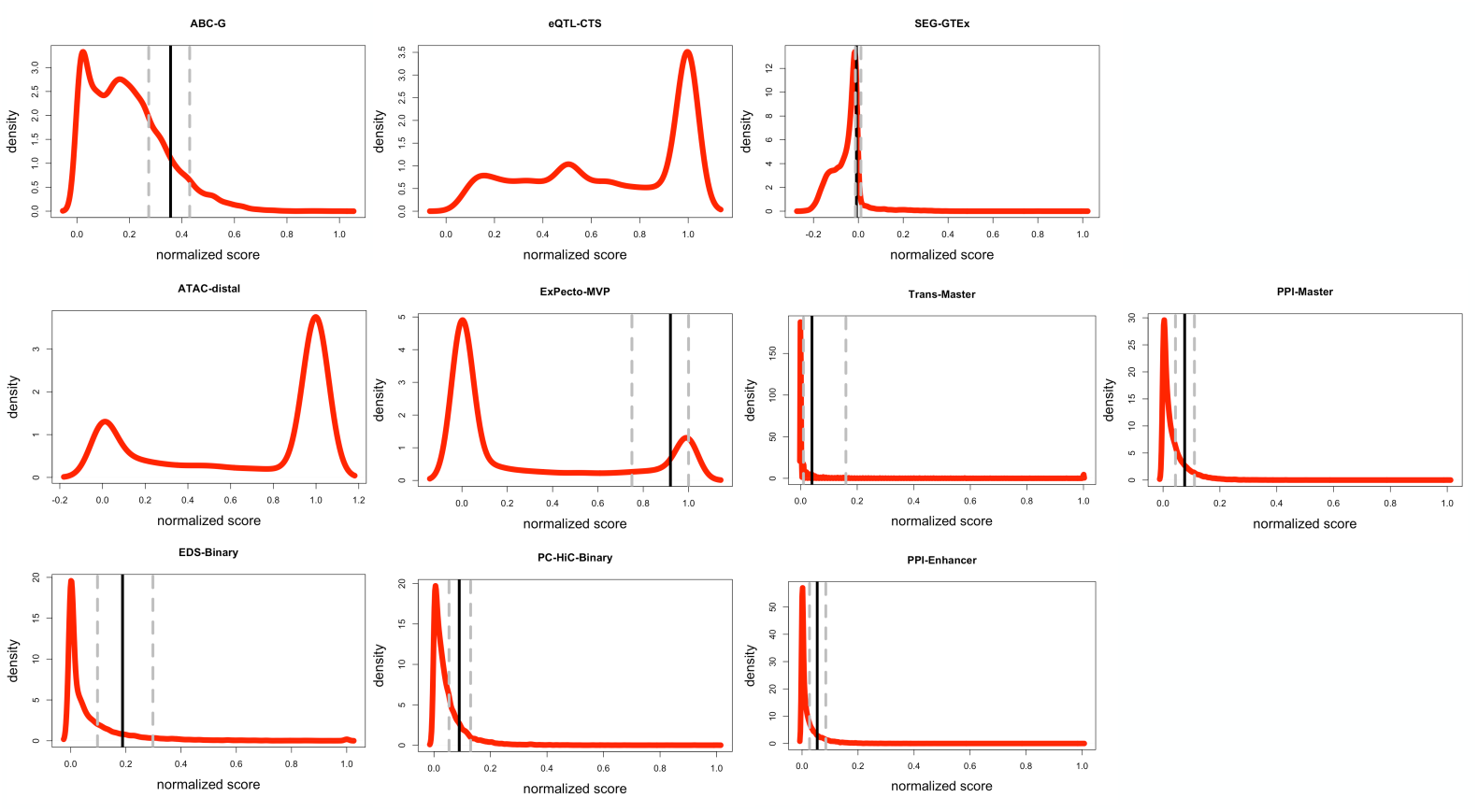
Density plots of the distribution of the metric underling each gene score. The density plots of the original gene scores (re-scaled between 0 and 1). The solid vertical line denotes the marker for top 10% genes for scores that are binarized in Table 1, while the dashed plots to its left and right correspond to top 5% and top 20% genes.

**Figure S2.**
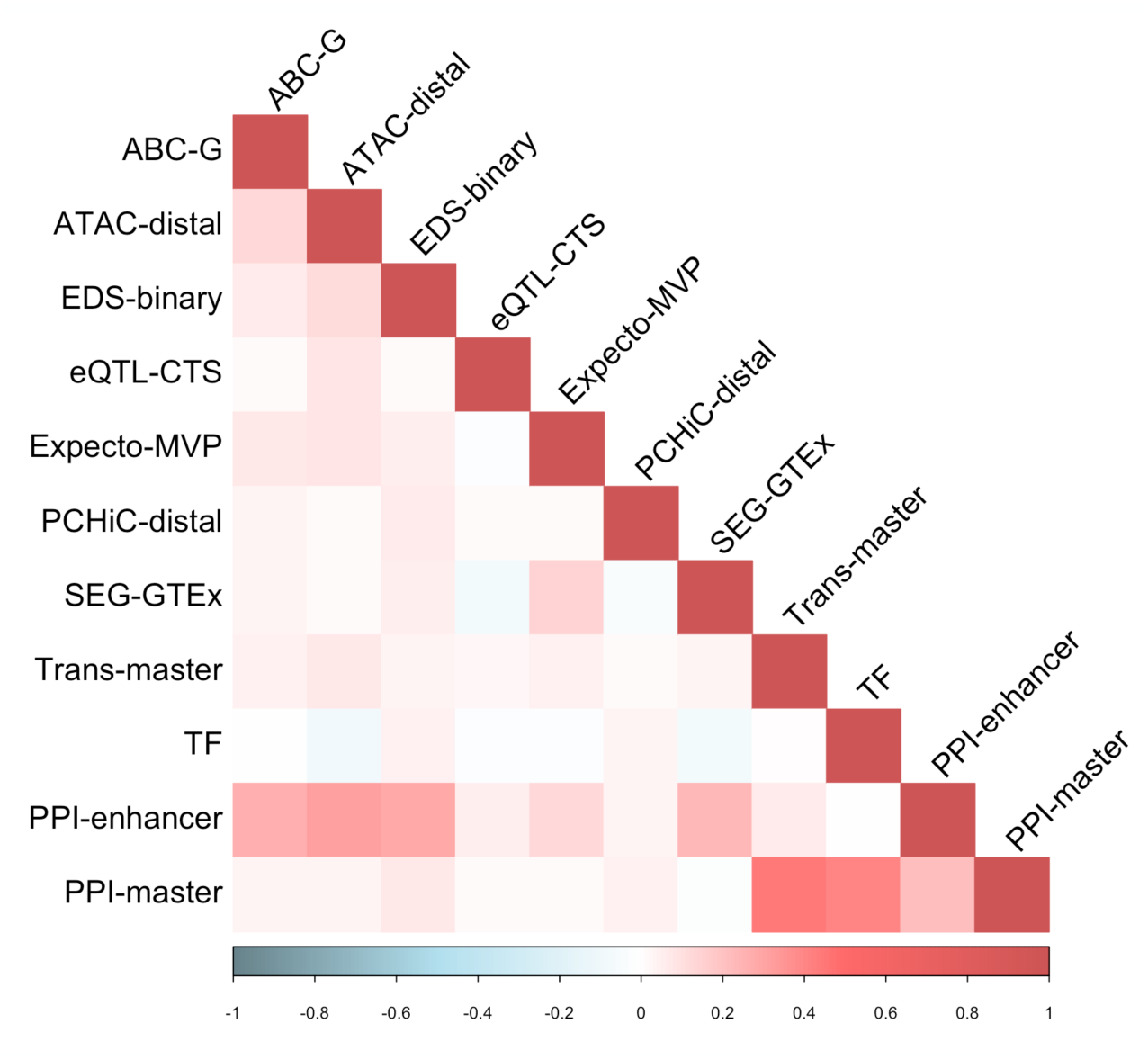
Correlation between different gene scores. Correlation matrix of all 11 gene scores (7 enhancer-related, 2 candidate master-regulator, 1 PPI-enhancer-related and 1 PPI-master) from Table 1 across all genes.

**Figure S3.**
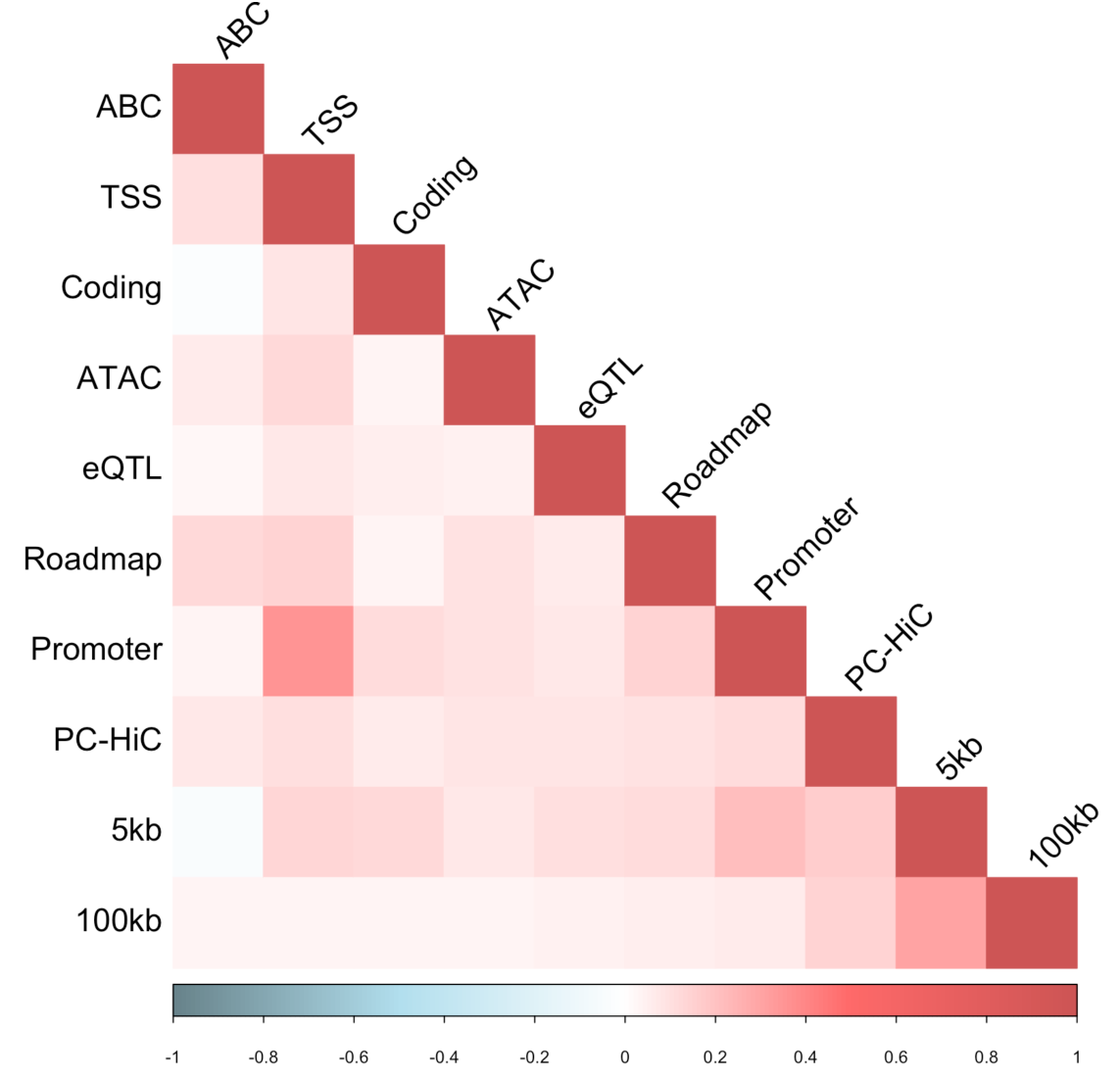
Correlation between SNP-to-gene (S2G) linking strategies. Correlation matrix of all 10 SNP-to-gene (S2G) linking strategies (Table 2), as assessed by the sets of SNPs linked to genes.

**Figure S4.**
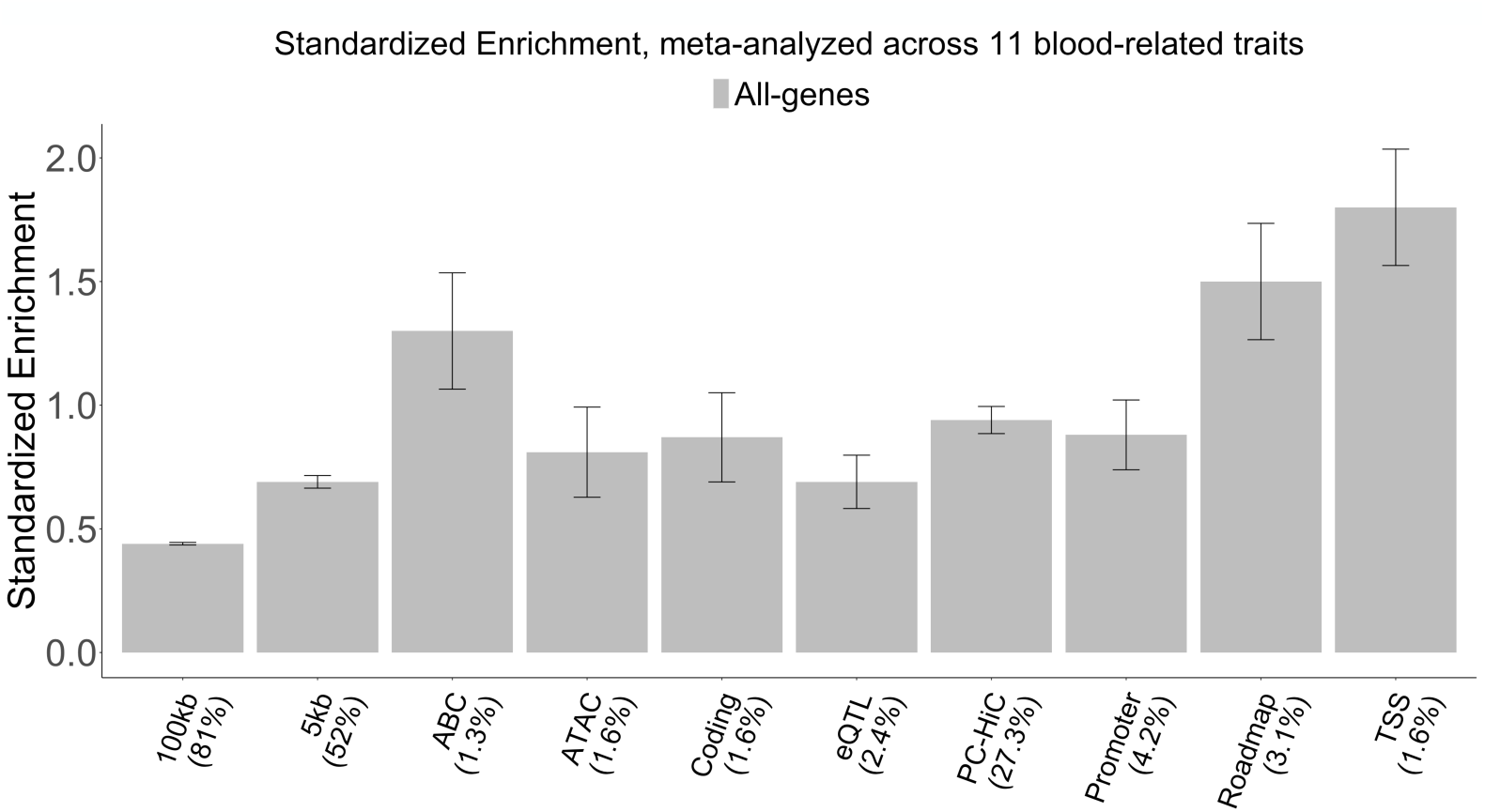
Standardized enrichment of SNP annotations for SNPs linked to all genes. Barplot representing standardized enrichment metric, as proposed in ref.^11^ for 10 SNP annotations corresponding SNPs linked to all genes by 10 S2G strategies from Table 2.

**Figure S5.**
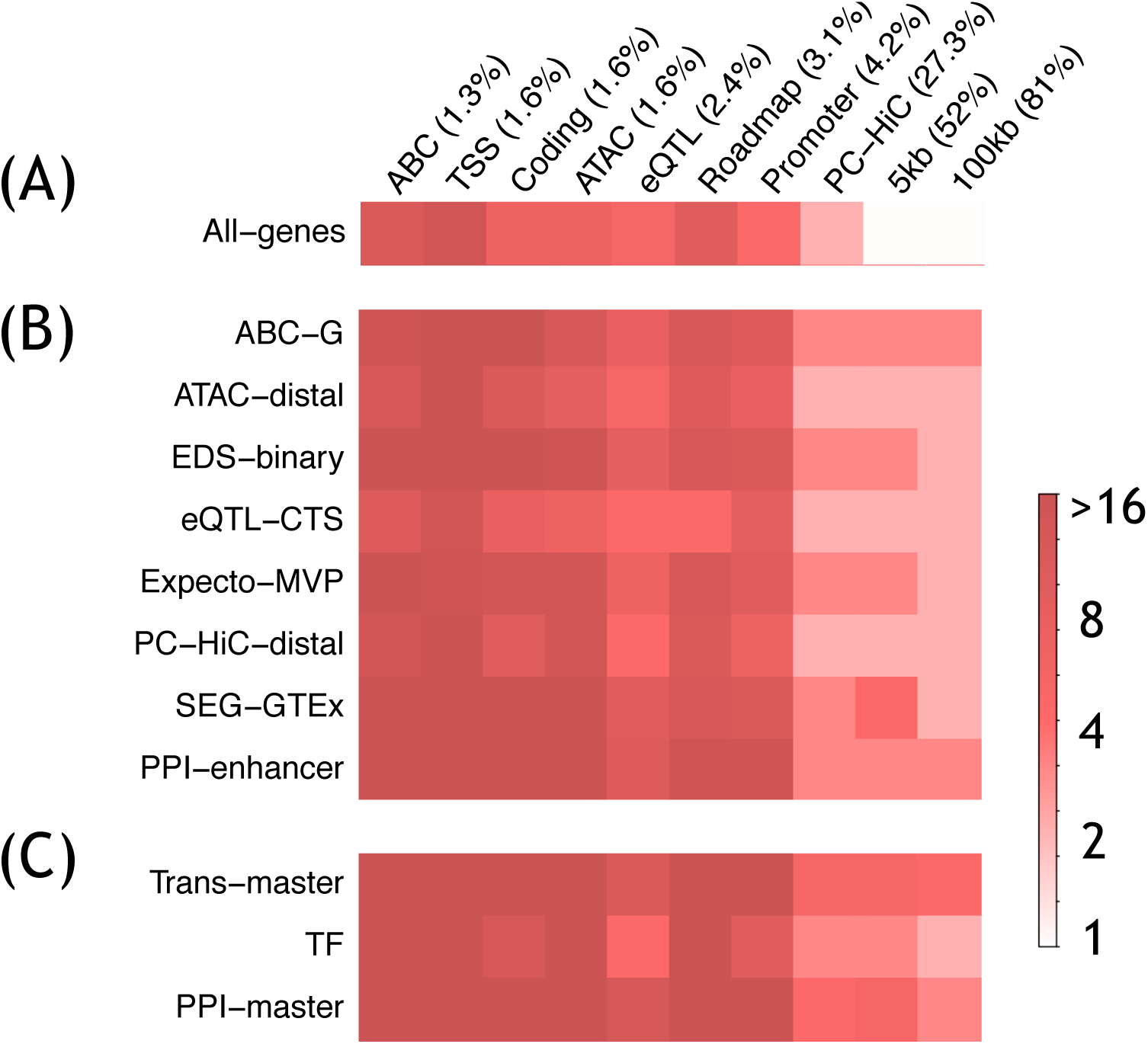
Heritability enrichment of SNP annotations corresponding to all gene scores. Heatmap representing heritability enrichment (log scale) metric for (A) SNP annotations corresponding to all genes, (B) SNP annotations corresponding to 7 enhancer-related and 1 PPI-enhancer gene scores, and (C) SNP annotations corresponding to 2 candidate master-regulator gene scores and 1 PPI-master gene score stacked sequentially from top to bottom..

**Figure S6.**
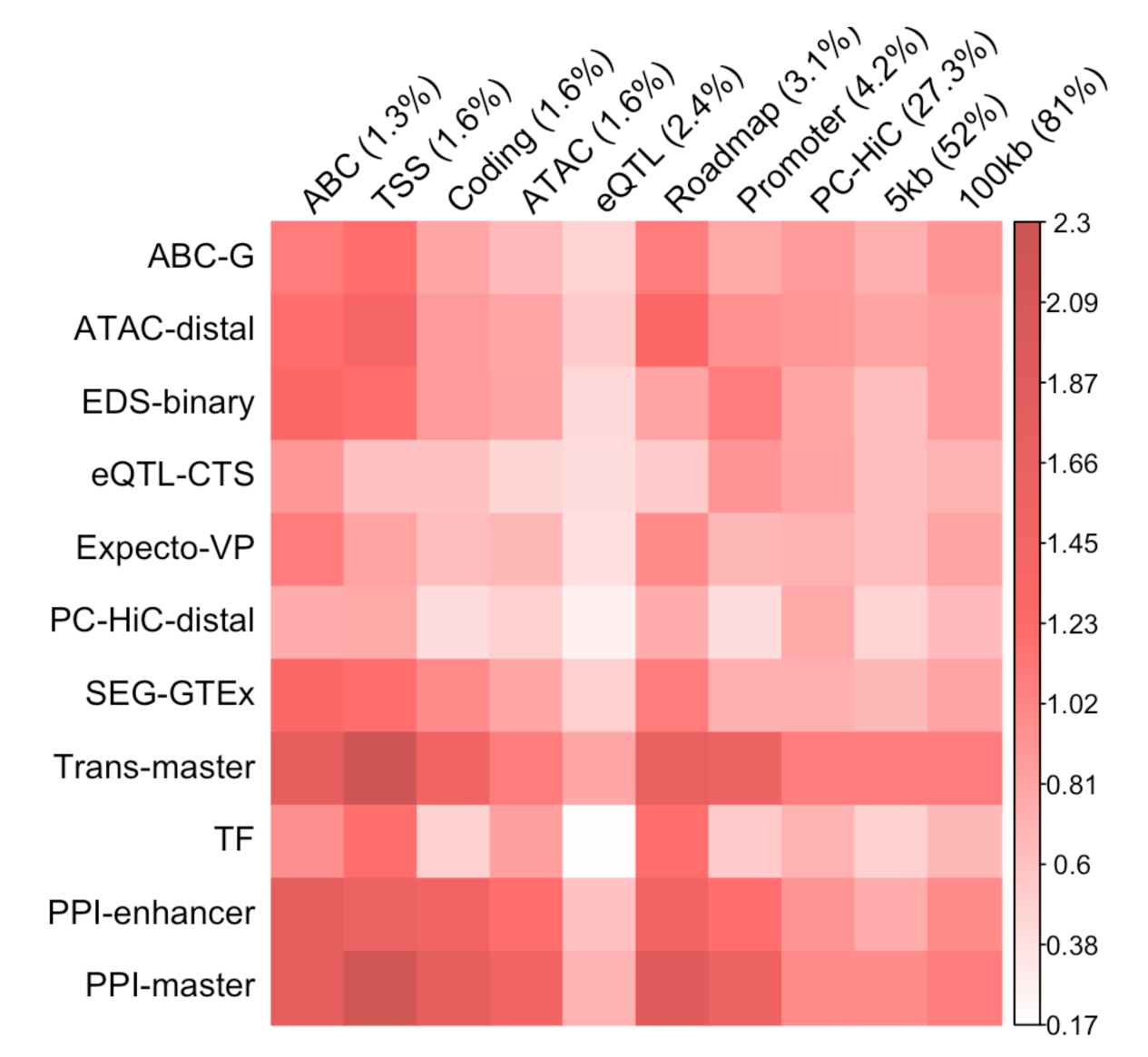
Standardized enrichment of SNP annotations corresponding to all 11 gene scores. Heatmap representing standardized enrichment metric, as proposed in ref.^11^ for all SNP annotations corresponding to all 11 gene scores (7 enhancer-related, 2 candidate master-regulator, 1 PPI-enhancer-related and 1 PPI-master) stacked sequentially from top to bottom.

**Figure S7.**
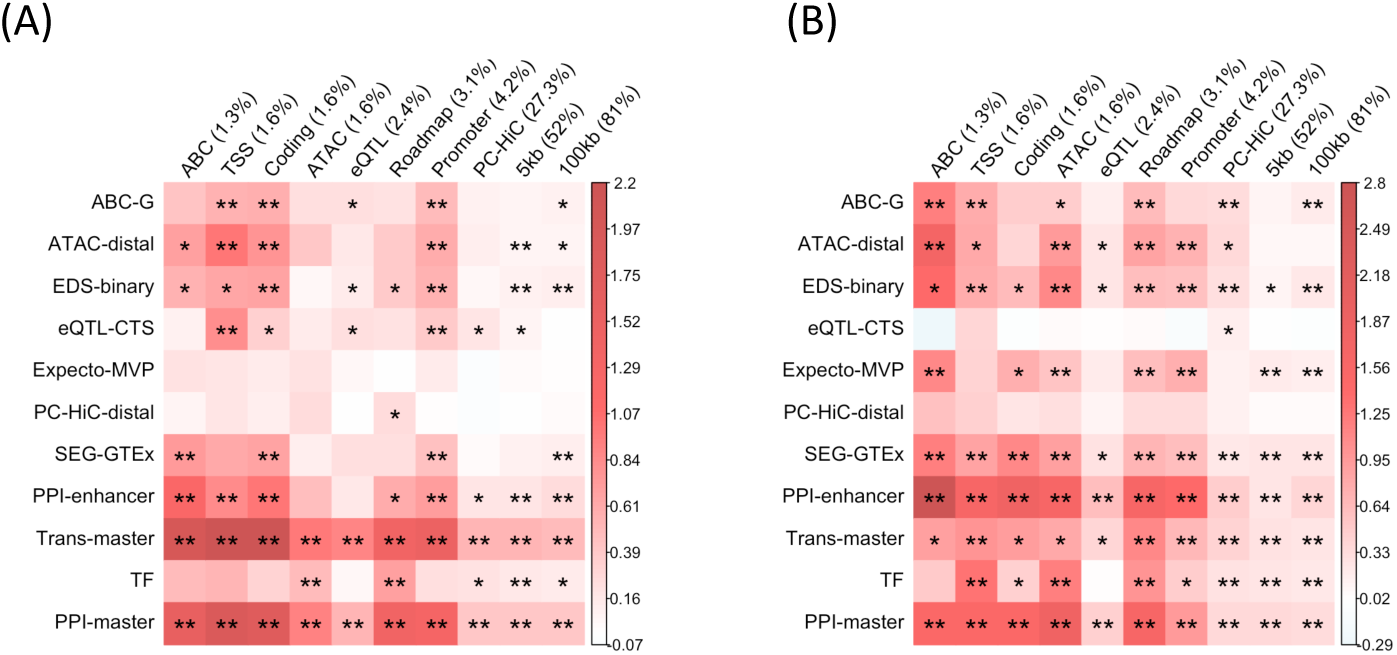
Standardized effect sizes of SNP annotations corresponding to all 11 genes scores, meta-analyzed across blood cell traits only or 5 autoimmune diseases only. Heatmap representing standardized effect size of all SNP annotations corresponding to all 11 gene scores (7 enhancer-related, 2 candidate master-regulator, 1 PPI-enhancer and 1 PPI-master; stacked sequentially from top to bottom) for two meta-analyses - (A) meta-analysis over 5 blood cell traits and (B) meta-analysis over 6 autoimmune diseases. The meta-analysis result for all 11 traits is reported in Figure 3B and Figure 4B. Numerical results are reported in Supplementary Tables S10, S11, S44, S45.

**Figure S8.**
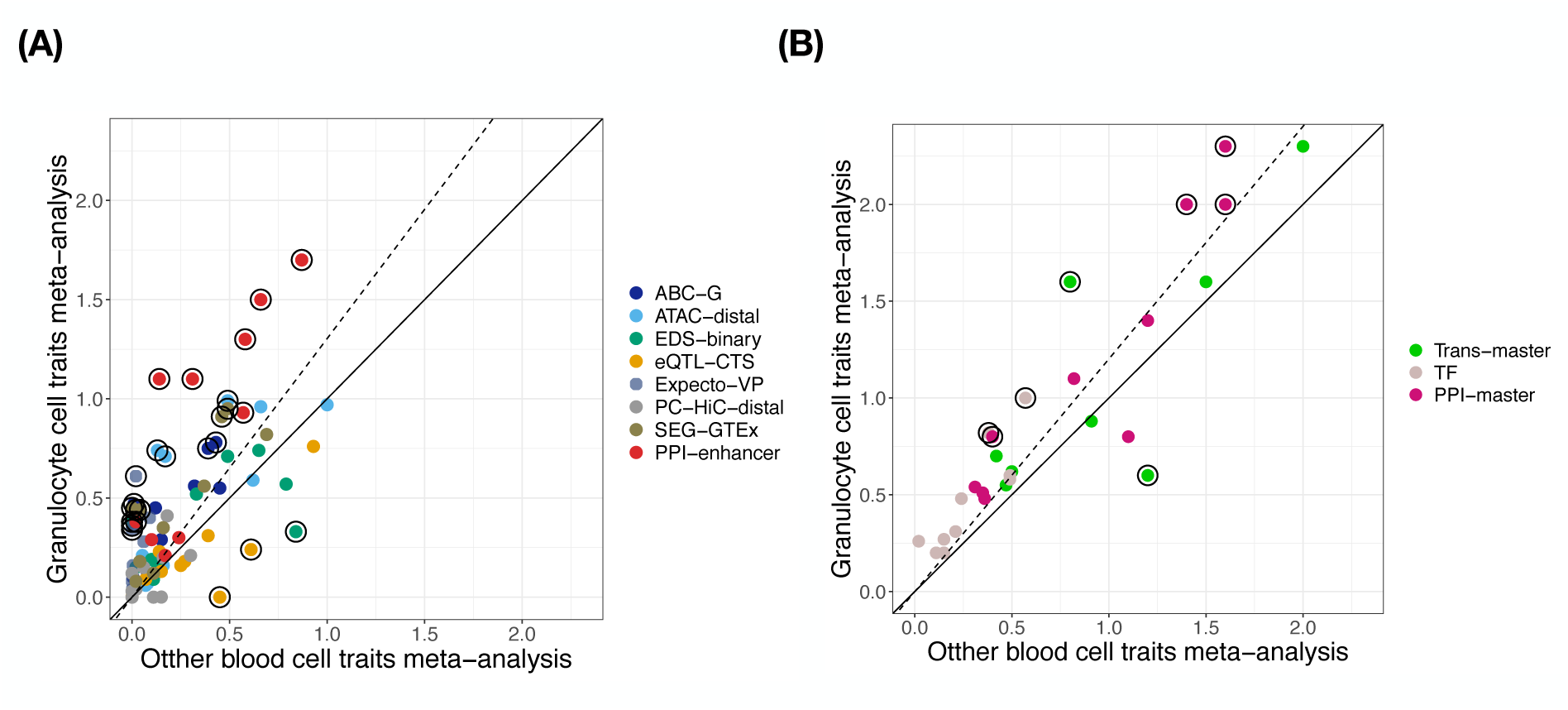
Standardized effect sizes of SNP annotations corresponding to all 11 genes scores, meta-analyzed across 2 granulocyte-related blood cell traits only or 3 red blood cell or platelet-related blood cell traits only. Scatter plot of the meta-analyzed standardized effect size *τ^*^* between two different meta-analysis - (i) across 2 granulocyte related blood cell traits (eosinophil count, white blood cell count) and (ii) 3 other blood cell traits (red blood cell counts, red blood cell distribution width, platelet count). The annotations with FDR (*<*0.05) significant difference in effect size between the two meta-analyses are circled. Numerical results are reported in Supplementary Tables S13, S12, S47 and S46.

**Figure S9.**
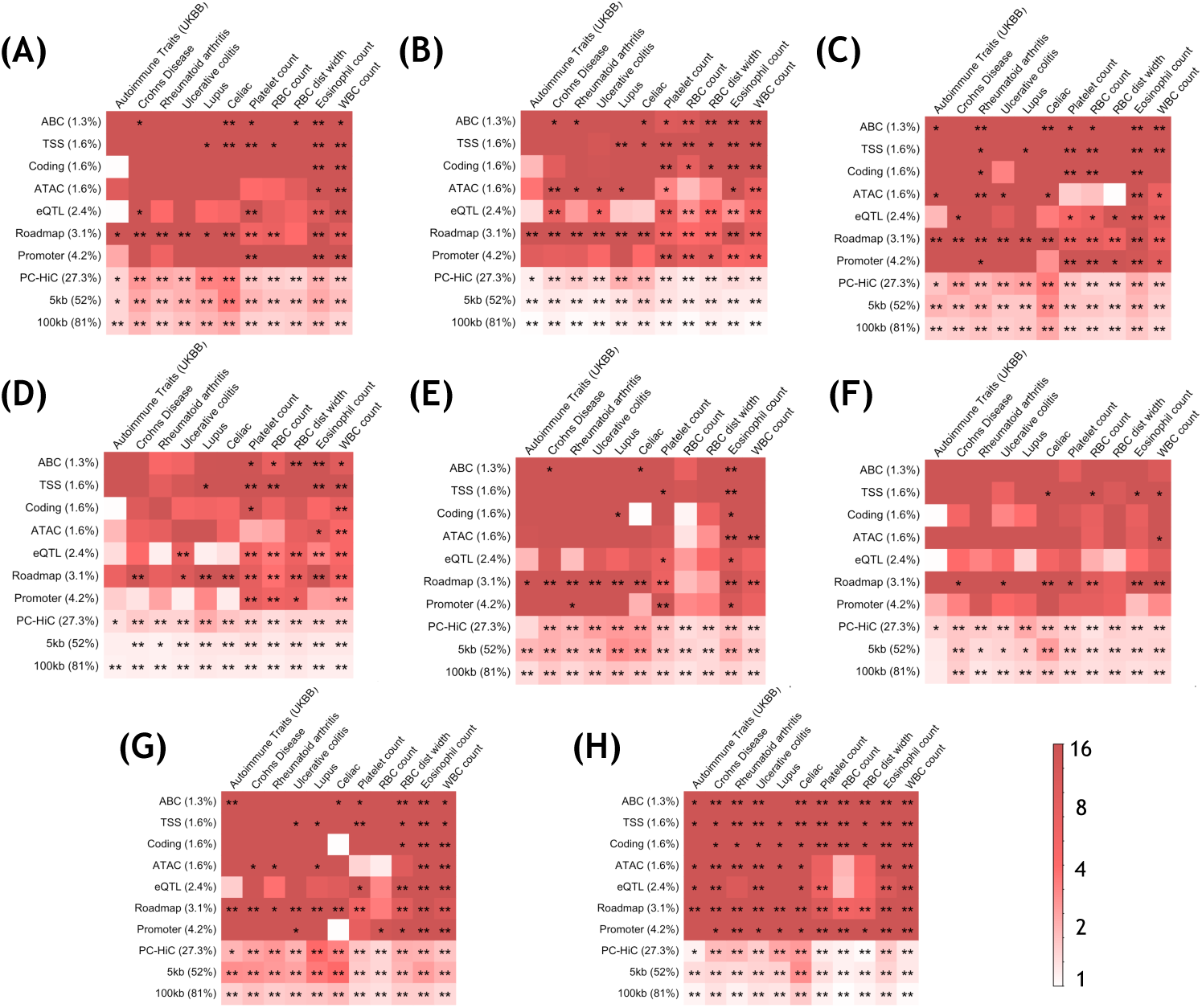
Heritability enrichment of SNP annotations corresponding to 7 enhancer-related and 1 PPI-enhancer gene scores for each of 11 autoimmune diseases and blood cell traits. Heatmap representing heritability enrichment of SNP annotations for each of 11 blood-related traits separately corresponding to (A) ABC-G, (B) ATAC-distal, (C) EDS-binary, (D) eQTL-CTS, (E) Expecto-MVP, (F) PC-HiC-distal, (G) SEG-GTEx and (H) PPI-enhancer gene scores. Numerical results are reported in Supplementary Table S14.

**Figure S10.**
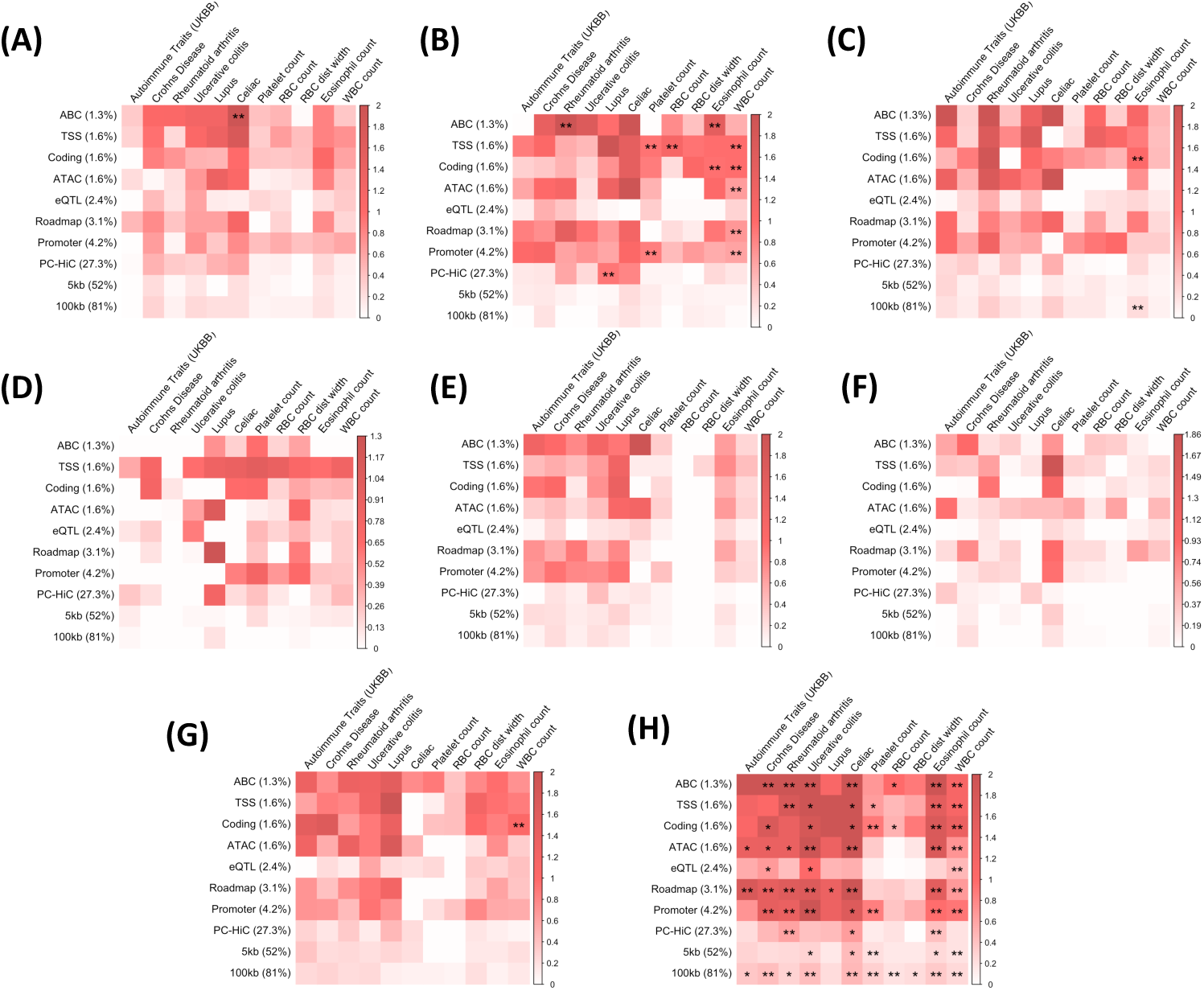
Standardized effect sizes of SNP annotations corresponding to 7 enhancer-related and 1 PPI-enhancer gene scores for each of 11 autoimmune diseases and blood cell traits. Heatmap representing standardized effect size of SNP annotations for each of 11 blood-related traits separately corresponding to (A) ABC-G, (B) ATAC-distal, (C) EDS-binary, (D) eQTL-CTS, (E) Expecto-MVP, (F) PC-HiC-distal, (G) SEG-GTEx and (H) PPI-enhancer gene scores. PPI-enhancer gene score showed consistently significant disease signal across multiple autoimmune diseases. Numerical results are reported in Supplementary Table S14.

**Figure S11.**
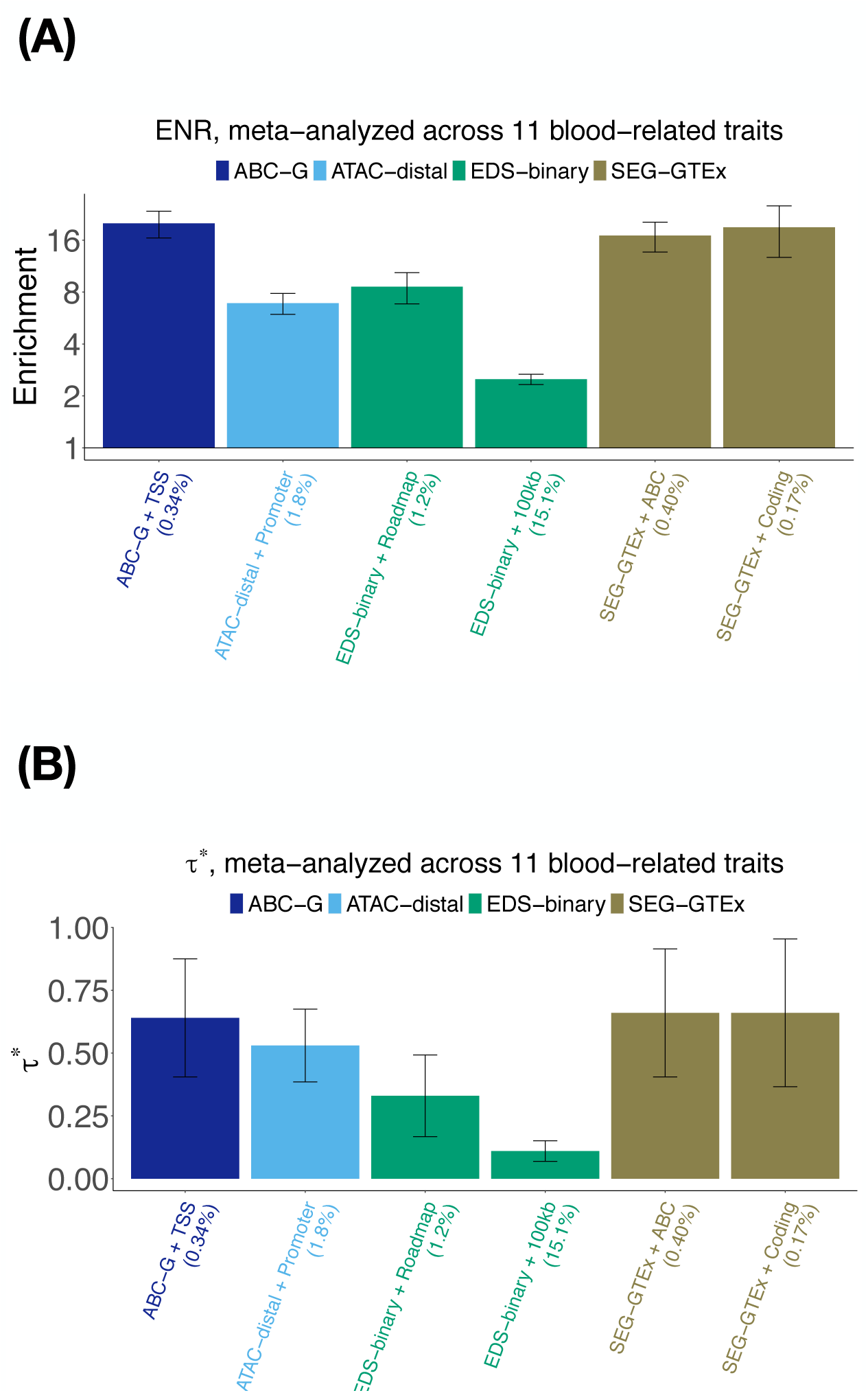
Disease informativeness of enhancer-related annotations in a joint analysis: (Panel A) Heritability enrichment (log scale), conditional on the enhancer-related joint model. Horizontal line denotes no enrichment. (Panel B) Standardized effect size *τ^*^* conditional on the same model. Results are shown only for the SNP annotations that are significant in the joint analysis after Bonferroni correction (0.05*/*110). Errors bars represent 95% confidence intervals. Numerical results are reported in Supplementary Table S16.

**Figure S12.**
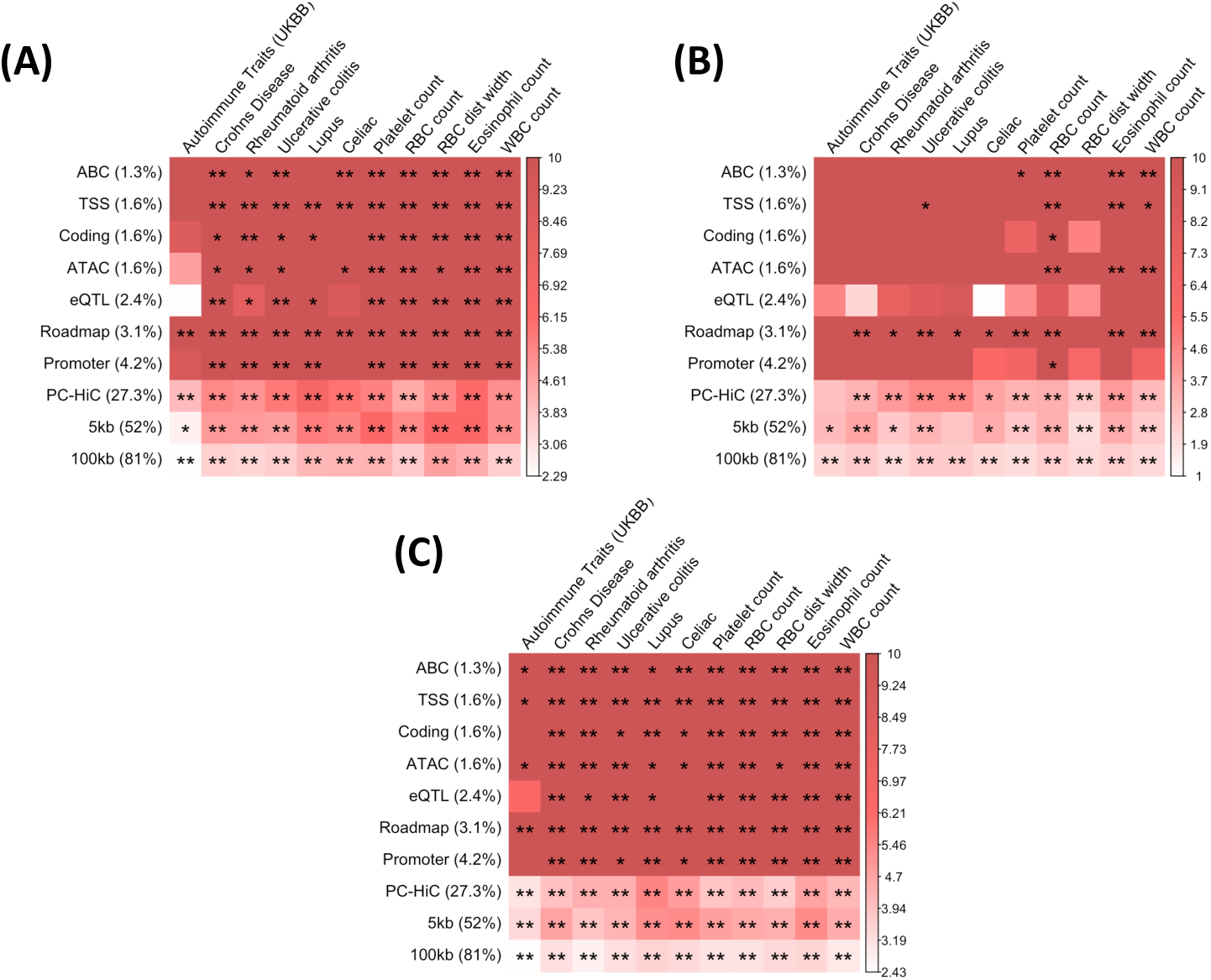
Heritability enrichment of SNP annotations corresponding to 2 candidate master-regulator and 1 PPI-master gene scores for each of 11 autoimmune diseases and blood cell traits. Heatmap representing heritability enrichment of SNP annotations for each of 11 blood-related traits separately corresponding to (A) Trans-master, (B) TF and (H) PPI-master gene scores. Numerical results are reported in Supplementary Table S14.

**Figure S13.**
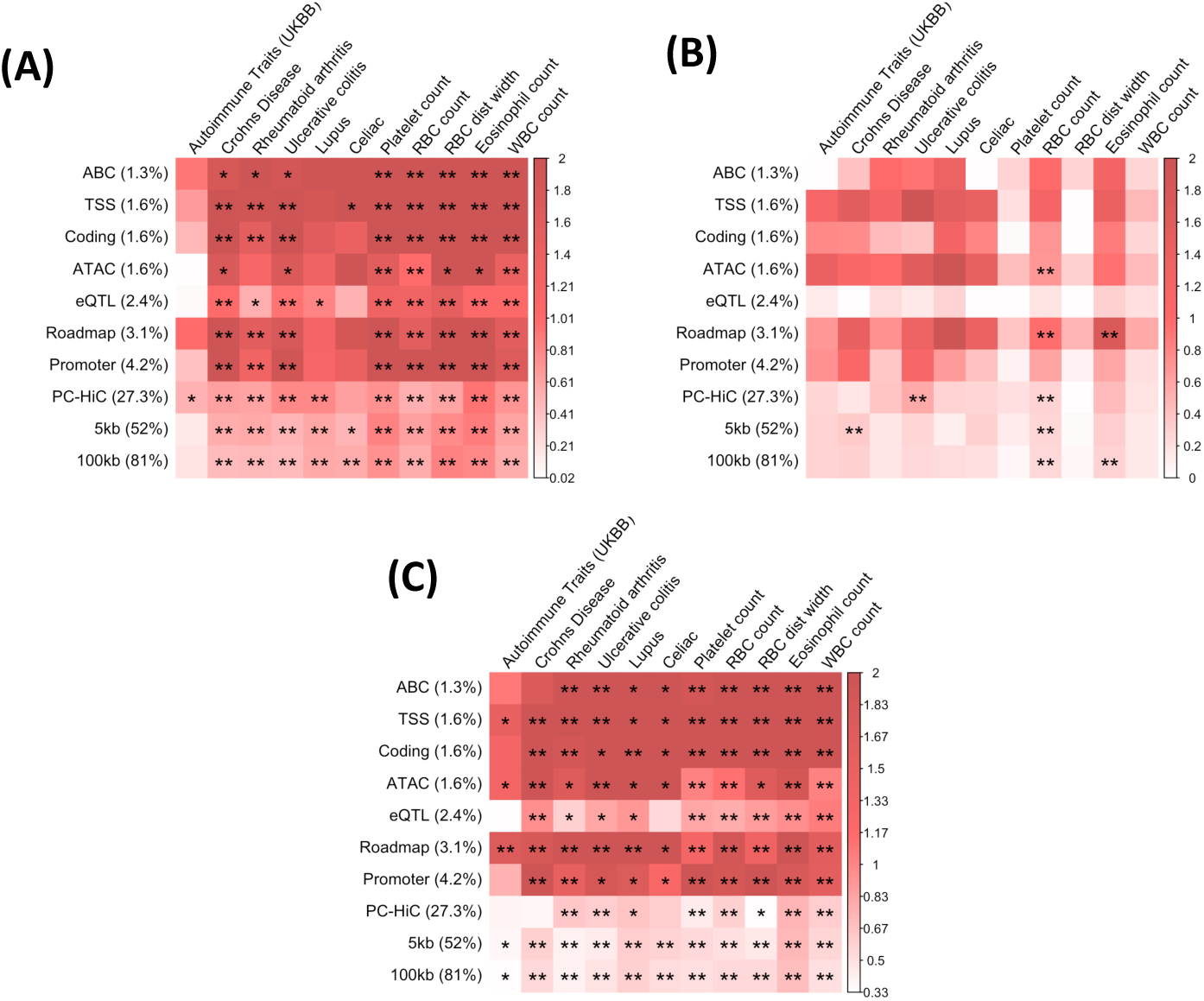
Standardized effect sizes of SNP annotations corresponding to 2 candidate master-regulator and 1 PPI-master gene scores for each of 11 autoimmune diseases and blood cell traits. Heatmap representing standardized effect size of SNP annotations for each of 11 blood-related traits separately corresponding to (A) Trans-master, (B) TF and (H) PPI-master gene scores. Trans-master and PPI-master gene scores showed consistently significant disease signal across multiple autoimmune diseases. Numerical results are reported in Supplementary Table S14.

**Figure S14.**
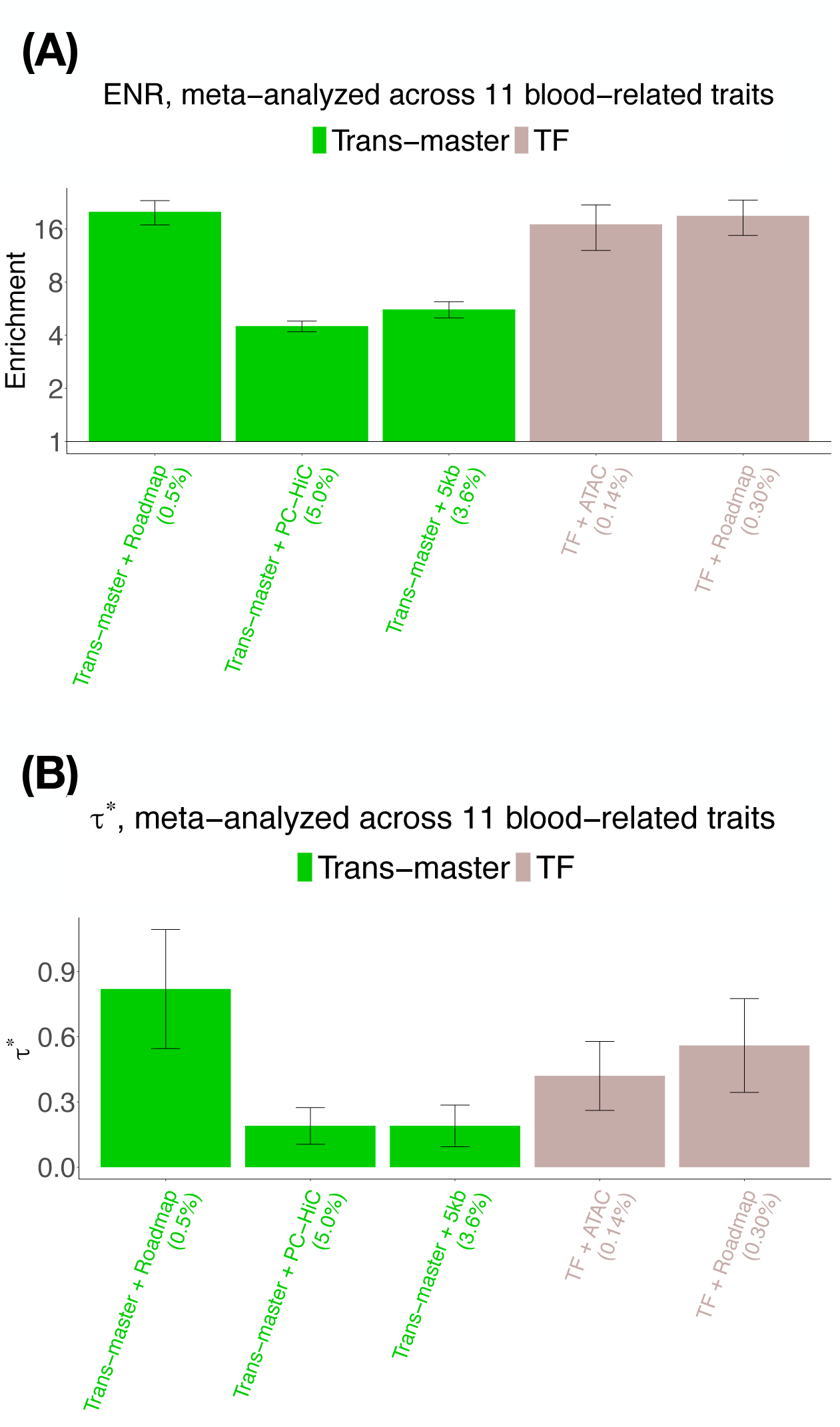
Disease informativeness of Master-regulator annotations in a joint analysis: (Panel A) Heritability enrichment (log scale), conditional on the candidate master-regulator joint model. Horizontal line denotes no enrichment. (Panel B) Standardized effect size *τ^*^* conditional on the same model. Results are shown only for the SNP annotations that are significant in the joint analysis after Bonferroni correction (0.05*/*110). Errors bars represent 95% confidence intervals. Numerical results are reported in Supplementary Table S48.

**Figure S15.**
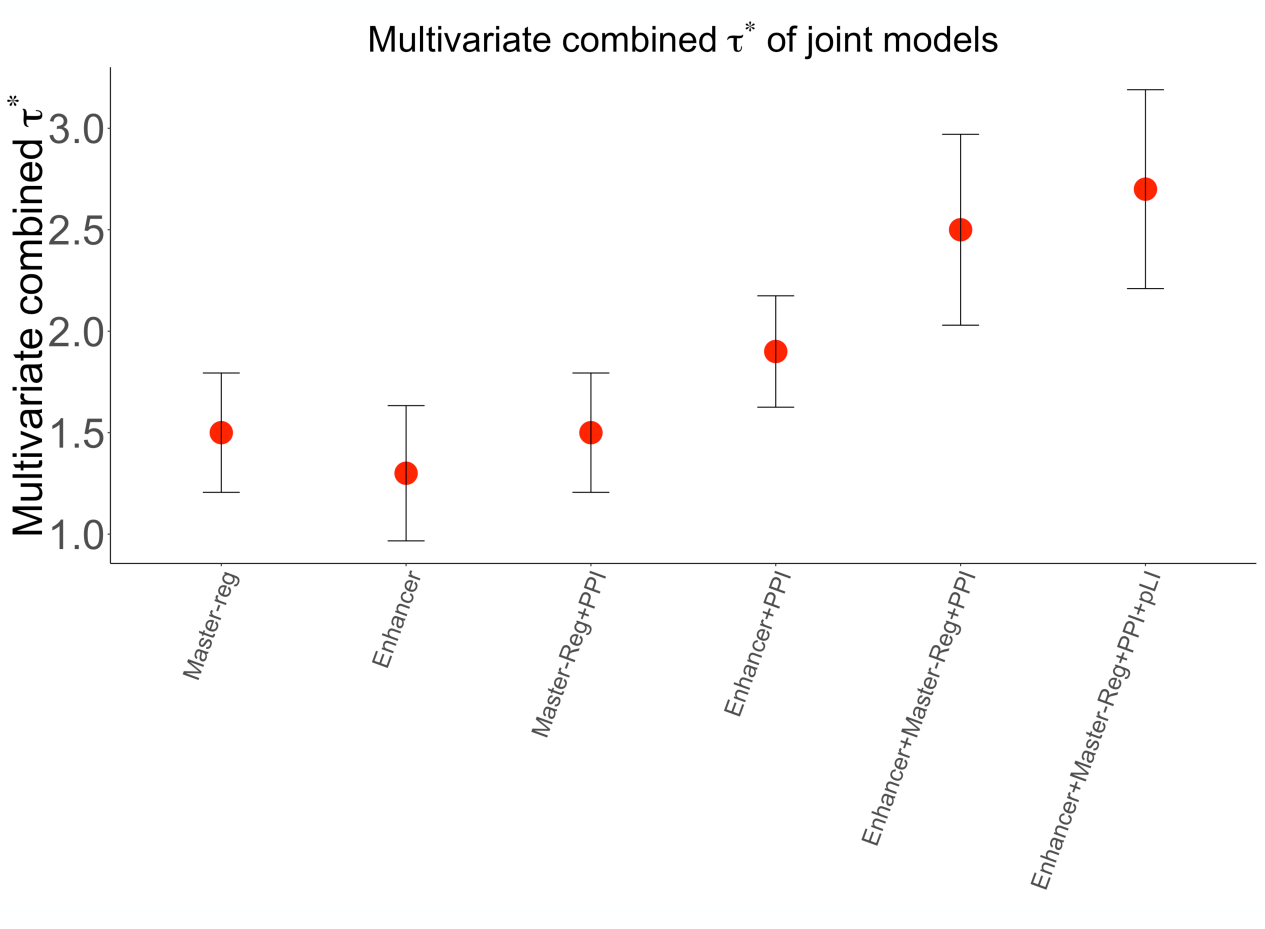
Combined *τ^*^* of different joint models studied: A plot of the combined *τ^*^* metric for different joint models. Numerical results are reported in Supplementary Table S59.

**Figure S16.**
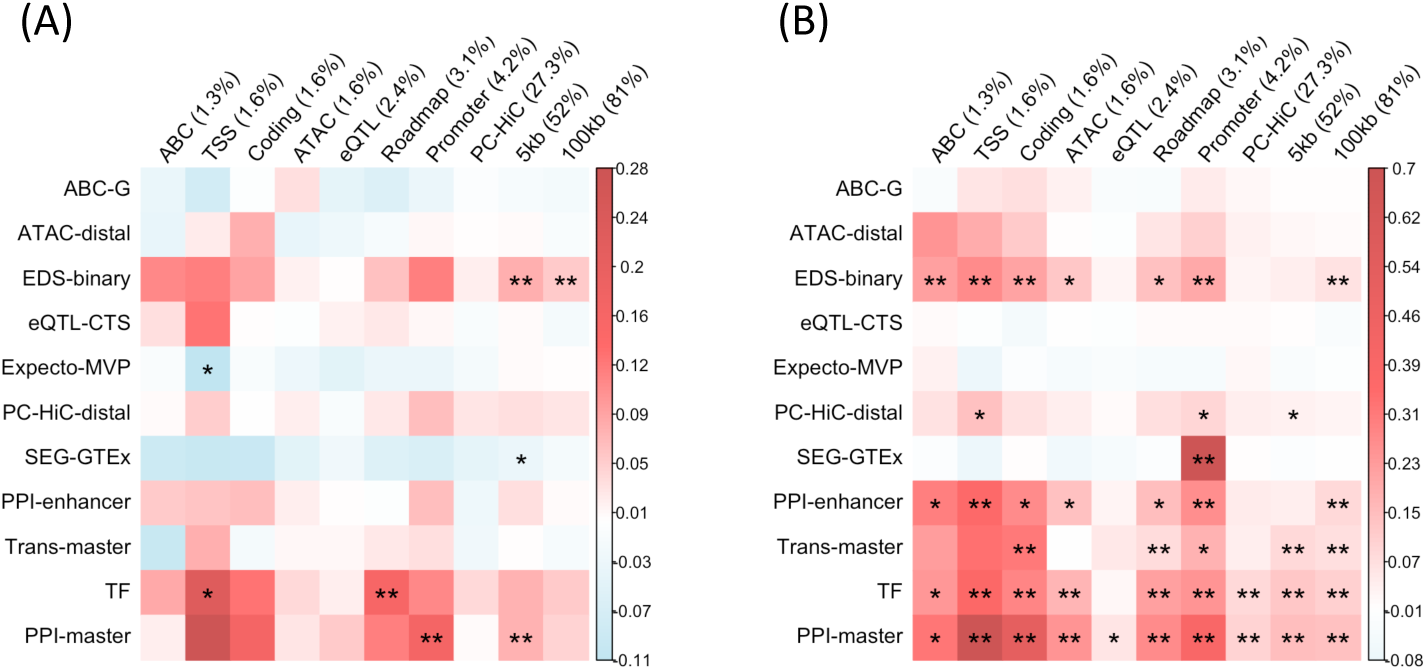
Standardized effect sizes of SNP annotations corresponding to all 11 gene scores, meta-analyzed across 8 brain-related traits or 28 non-brain and non-blood related traits. Heatmap representing standardized effect size of all SNP annotations corresponding to all 11 gene scores (7 enhancer-related, 2 candidate master-regulator, 1 PPI-enhancer and 1 PPI-master; stacked sequentially from top to bottom) for two meta-analyses -

**Figure S17.**
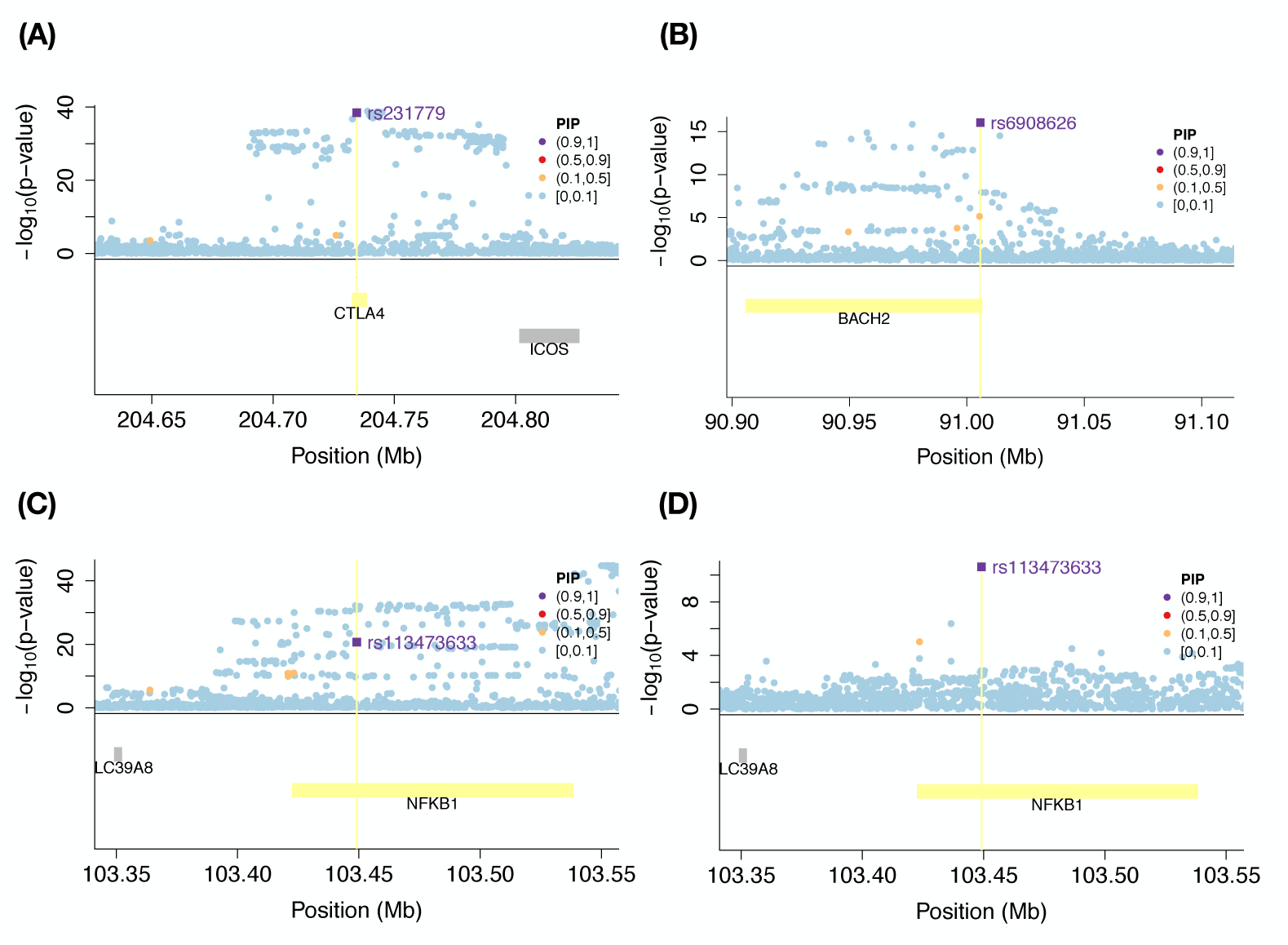
Illustrative examples of fine-mapped SNPs belonging to SNP annotations analyzed in this study: Illustration of four fine-mapped SNPs (PIP *>* 0.90), 2 in UKBB “All autoimmune traits”, and one variant that is fine-mapped for both WBC count and Eosinophil count traits (Table S1). These SNPs overlap with the region implicated by a S2G strategy linking to a gene in top 2% for some of the jointly informative S2G annotations in Figure 5. (A) rs231779 is linked to gene *CTLA4* by PPI-enhancer gene score (ranked 109) using ABC S2G strategy for “All autoimmune traits”, (B) rs6908626 is linked to gene *BACH2* by Trans-master gene score (ranked 311) using Roadmap S2G strategy for “All autoimmune traits”, (C, D) rs113473633 is linked to gene *NFKB1* by PPI-master (ranked 111) using Roadmap S2G strategy for WBC count (C) and Eosinophil count (D). Numerical results are reported in Supplementary Table S67.

**Figure S18.**
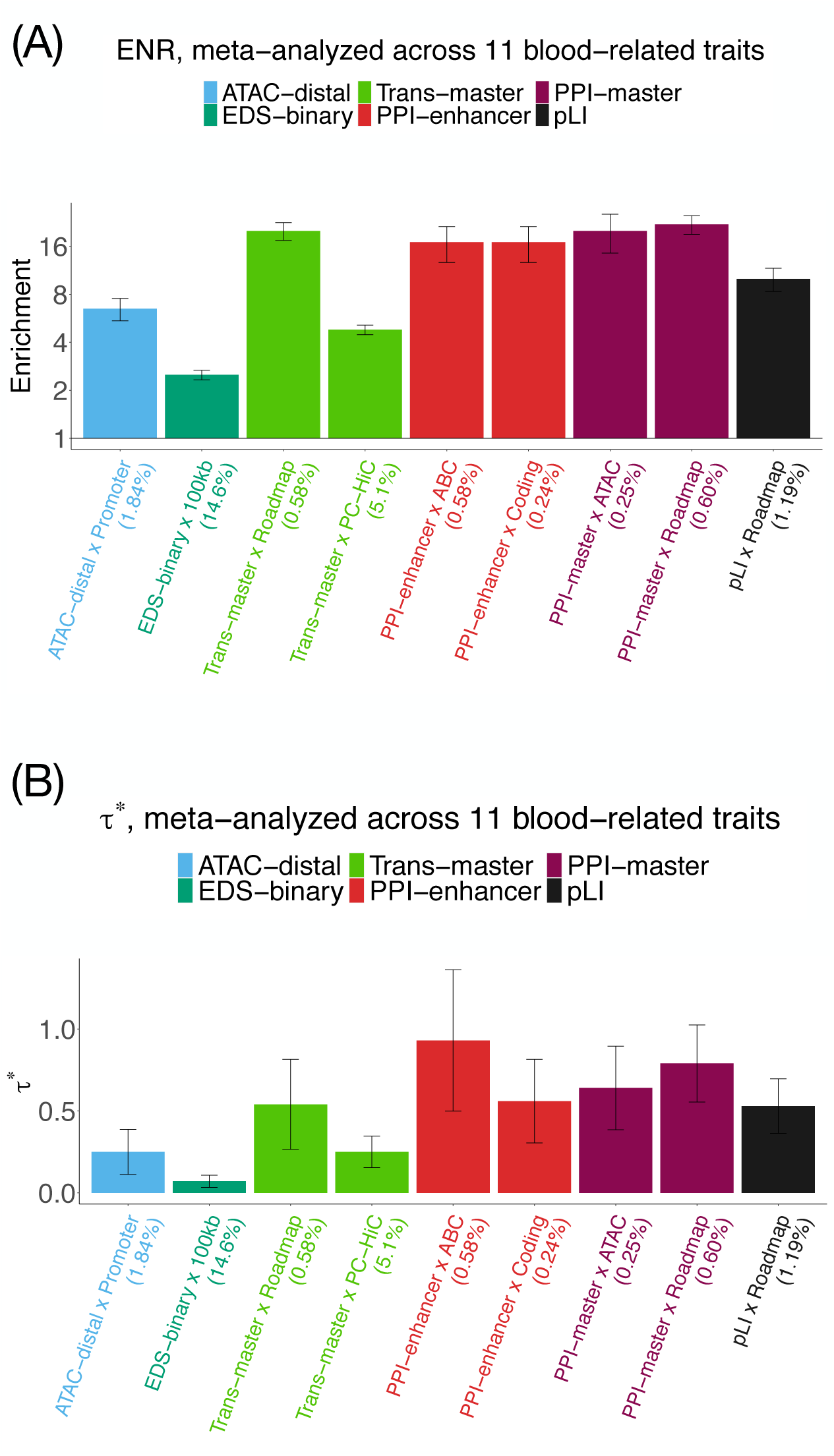
Disease informativeness of enhancer-related, PPI-enhancer, candidate master-regulator, PPI-master and pLI SNP annotations in a joint analysis: (Panel A) Heritability enrichment, conditional on the combined joint model of enhancer-related, PPI-enhancer, candidate master-regulator, PPI-master and pLI SNP annotations. Horizontal line denotes no enrichment. (Panel B) Standardized effect size *τ^*^* conditional on the same model. Results are shown only for the SNP annotations that are significant in the joint model after Bonferroni correction (0.05*/*110). Errors bars represent 95% confidence intervals. Numerical results are reported in Table S68.

**Figure S19.**
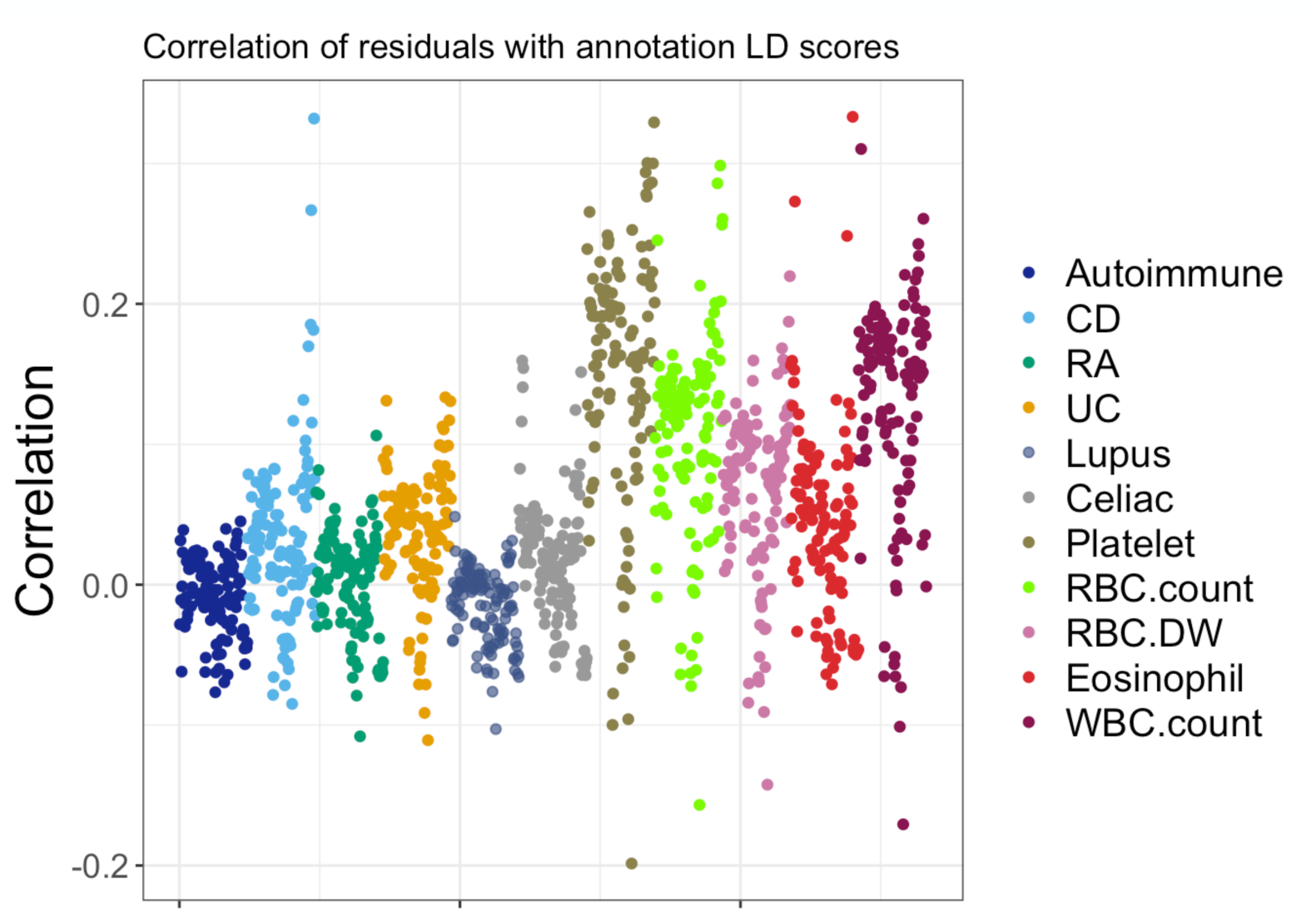
Correlations of residuals from stratified LD score regression with annotation-specific LD scores: A diagnostic plot assessing the model fit of the stratified LD score regression (across 11 blood-related traits) where we evaluate the correlation of the residuals from the final joint model with LD score predictors corresponding to 121 annotations in the final joint model (113 baseline-LD+cis and 8 annotations from Figure 5). The mean correlation was observed to be 0.05 which is close to 0.

## References

1. M.T. Maurano et al. Systematic localization of common disease-associated variation in regulatory DNA. Science, 337:1190–1195, 2012.

2. G. Trynka et al. Chromatin marks identify critical cell types for fine mapping complex trait variants. Nature genetics, 45(2):124–130, 2013.

3. J.K. Pickrell. Joint analysis of functional genomic data and genome-wide association studies of 18 human traits. The American Journal of Human Genetics, 94(4):559–573, 2014.

4. A.L. Price, C.C. Spencer, and P. Donnelly. Progress and promise in understanding the genetic basis of common diseases. Proceedings of the Royal Society B: Biological Sciences, 282(1821):20151684, 2015.

5. P.M. Visscher et al. 10 years of GWAS discovery: biology, function, and translation. The American Journal of Human Genetics, 101(1):5–22, 2017.

6. J. Shendure, G.M. Findlay, and M.W. Snyder. Genomic medicine–progress, pitfalls, and promise. Cell, 177:45–57, 2019.

7. J. Zhou, C.L. Theesfeld, K. Yao, K.M. Chen, A.K. Wong, and O.G. Troyanskaya. Deep learning sequence-based ab initio prediction of variant effects on expression and disease risk. Nature genetics, 50:1171–1179, 2018.

8. X. Zhu and M. Stephens. Large-scale genome-wide enrichment analyses identify new trait-associated genes and pathways across 31 human phenotypes. Nature communications, 9(1):4361, 2018.

9. H.K. Finucane, Y.A. Reshef, V. Anttila, K. Slowikowski, A. Gusev, A. Byrnes, et al. Heritability enrichment of specifically expressed genes identifies diseaserelevant tissues and cell types. Nature genetics, 50:621–629, 2018.

10. H. Fang et al. A genetics-led approach defines the drug target landscape of 30 immune-related traits. Nature genetics, 51:1082–1091, 2019.

11. S.S. Kim et al. Genes with high network connectivity are enriched for disease heritability. The American Journal of Human Genetics, 104:896–913, 2019.

12. Q. Wang et al. A Bayesian framework that integrates multi-omics data and gene networks predicts risk genes from schizophrenia GWAS data. Nature neuroscience, 22:691–699, 2019.

13. C.S. Smillie et al. Intra-and inter-cellular rewiring of the human colon during ulcerative colitis. Cell, 178:714–730, 2019.

14. M. Wainberg et al. Opportunities and challenges for transcriptome-wide association studies. Nature genetics, 51:592–599, 2019.

15. A.D. Sawle et al. Identification of master regulator genes in human periodontitis. Journal of dental research, 95:1010–1017, 2016.

16. E.A. Boyle, Y.I. Li, and J.K. Pritchard. An expanded view of complex traits: from polygenic to omnigenic. Cell, 169:1177–1186, 2017.

17. B. Brynedal et al. Large-scale trans-eQTLs affect hundreds of transcripts and mediate patterns of transcriptional co-regulation. The American Journal of Human Genetics, 100(4):581–591, 2017.

18. C. Yao et al. Dynamic role of trans regulation of gene expression in relation to complex traits. The American Journal of Human Genetics, 100:571–580, 2017.

19. D.M.D. Vargas et al. Alzheimer’s disease master regulators analysis: search for potential molecular targets and drug repositioning candidates. Alzheimer’s research & therapy, 10:59, 2018.

20. L.E. Montefiori et al. A promoter interaction map for cardiovascular disease genetics. Elife, 7:e35788, 2018.

21. X. Liu, Y.I. Li, and J.K. Pritchard. Trans effects on gene expression can drive omnigenic inheritance. Cell, 177:1022–1034, 2019.

22. A.D. Torshizi et al. Deconvolution of Transcriptional Networks Identifies TCF4 as a Master Regulator in Schizophrenia. Science Advances, 5:eaau4139, 2019.

23. R. Andersson and A. Sandelin. Determinants of enhancer and promoter activities of regulatory elements. Nature Reviews Genetics, pages 1–17, 2019.

24. X. Wang and D.B. Goldstein. Enhancer Domains Predict Gene Pathogenicity and Inform Gene Discovery in Complex Disease. The American Journal of Human Genetics, 106:215–233, 2020.

25. E.S. Emison et al. A common sex-dependent mutation in a ret enhancer underlies hirschsprung disease risk. Nature, 434(7035):857–863, 2005.

26. S. Chatterjee et al. Enhancer variants synergistically drive dysfunction of a gene regulatory network in hirschsprung disease. Cell, 167(2):pp.355–368, 2016.

27. K.S. Kobayashi and P.J. Van Den Elsen. Nlrc5: a key regulator of mhc class i-dependent immune responses. Nature Reviews Immunology, 12(12):813–820, 2012.

28. H.K. Finucane, B. Bulik-Sullivan, A. Gusev, G. Trynka, Y. Reshef, P.R. Loh, V. Anttila, H. Xu, C. Zang, K. Farh, and S. Ripke. Partitioning heritability by functional annotation using genome-wide association summary statistics. Nature genetics, 47:1228–1235, 2015.

29. S. Gazal et al. Linkage disequilibrium–dependent architecture of human complex traits shows action of negative selection. Nature genetics, 49(10):1421–1427, 2017.

30. S. Gazal, C. Marquez-Luna, H.K. Finucane, and A.L. Price. Reconciling s-ldsc and ldak models and functional enrichment estimates. Nature genetics, 51(8):1202–1204, 2019.

31. 1000 Genomes Project Consortium. A global reference for human genetic variation. Molecular cell, 526(7571):68–74, 2015.

32. C.P. Fulco et al. Activity-by-contact model of enhancer–promoter regulation from thousands of CRISPR perturbations. Nature Genetics, 51:1664–1669, 2019.

33. J. Nasser et al. Genome-wide enhancer maps link risk variants to disease genes. Nature, 593(7858):238–243, 2021.

34. S. McCarthy et al. A reference panel of 64,976 haplotypes for genotype imputation. Nature genetics, 48:1279–1283, 2016.

35. C. Bycroft et al. The uk biobank resource with deep phenotyping and genomic data. Nature, 562(7726):203–209, 2018.

36. D. Taliun et al. Sequencing of 53,831 diverse genomes from the nhlbi topmed program. Nature, 590(7845):290–299, 2021.

37. F. Hormozdiari et al. Leveraging molecular quantitative trait loci to understand the genetic architecture of diseases and complex traits. Nature genetics, 50(7):1041–1047, 2018.

38. S. Gazal et al. Functional architecture of low-frequency variants highlights strength of negative selection across coding and non-coding annotations. Nat. Genet, 50:1600–1607, 2018.

39. F.I. Hormozdiari et al. Functional disease architectures reveal unique biological role of transposable elements. Nature communications, 10(1):1–8, 2019.

40. P.F. Palamara et al. High-throughput inference of pairwise coalescence times identifies signals of selection and enriched disease heritability. Nature Genetics, 50(9):1311–1317, 2018.

41. K.K. Dey et al. Evaluating the informativeness of deep learning annotations for human complex diseases. Nature communications, 11(1):pp.1–9, 2020.

42. S.S. Kim et al. Improving the informativeness of Mendelian disease-derived pathogenicity scores for common disease. Nature communications, 11(1):1–15, 2020.

43. A. Gaulton et al. The ChEMBL database in 2017. Nucleic acids research, 45:D945–D954, 2016.

44. M.K. Freund et al. Phenotype-specific enrichment of Mendelian disorder genes near GWAS regions across 62 complex traits. The American Journal of Human Genetics, 103 (4):pp.535–552, 2018.

45. D. Vuckovic et al. The polygenic and monogenic basis of blood traits and diseases. Cell, 182 (5):pp.1214–1231, 2020.

46. C.F. Wright et al. Genetic diagnosis of developmental disorders in the DDD study: a scalable analysis of genome-wide research data. The Lancet, 385(9975):1305– 1314, 2015.

47. M. Lek, K.J. Karczewski, E.V. Minikel, K.E. Samocha, E. Banks, et al. Analysis of protein-coding genetic variation in 60,706 humans. Nature, 536:285–291, 2016.

48. H. Yoshida et al. The cis-Regulatory Atlas of the Mouse Immune System. Cell, 176:897–912, 2019.

49. A.P. Schoech et al. Quantification of frequency-dependent genetic architectures in 25 UK Biobank traits reveals action of negative selection. Nature communications, 10:1–10, 2019.

50. K.K.H. Farh et al. Genetic and epigenetic fine mapping of causal autoimmune disease variants. Nature, 518:337–343, 2015.

51. O. Weissbrod et al. Functionally-informed fine-mapping and polygenic localization of complex trait heritability. Nature Genetics, 52(12):p.1355–1363, 2020.

52. A. Kamburov et al. The ConsensusPathDB interaction database: 2013 update. Nucleic acids research, 41(D1):D793–D800, 2012.

53. D. Szklarczyk et al. The STRING database in 2017: quality-controlled protein–protein association networks, made broadly accessible. Nucleic acids research, 45(Database issue):D362–D368, 2017.

54. H. Tong, C. Faloutsos, and J.Y. Pan. Random walk with restart: fast solutions and applications. Knowledge and Information Systems, 14:327–346, 2008.

55. A.R. Sonawane et al. Understanding tissue-specific gene regulation. Cell reports, 21:1077–1088, 2017.

56. U. Võosa, et al. Unraveling the polygenic architecture of complex traits using blood eQTL meta-analysis. bioRxiv, page 447367, 2018.

57. S.A. Lambert et al. The human transcription factors. Cell, 172:650–665, 2018.

58. W. Cai et al. Master regulator genes and their impact on major diseases. PeerJ, 8:p.e9952, 2020.

59. M.C. Nakamura. CIITA: a master regulator of adaptive immunity shows its innate side in the bone. Journal of bone and mineral research: the official journal of the American Society for Bone and Mineral Research, 29(2):p.287, 2014.

60. L. Colomer, C.and Marruecos, A. Vert, A. Bigas, and L. Espinosa. NF-kB members left home: NF-kB-independent roles in cancer. Biomedicines, 5(2):p.26, 2017.

61. E.H. Bresnick, K.R. Katsumura, H.Y. Lee, K.D. Johnson, and A.S. Perkins. Master regulatory GATA transcription factors: mechanistic principles and emerging links to hematologic malignancies. Nucleic acids research, 40(13):pp.5819–5831, 2012.

62. S. Paul, P. Home, B. Bhattacharya, and S. Ray. GATA factors: Master regulators of gene expression in trophoblast progenitors. Placenta, 60:pp.S61–S66, 2017.

63. B. van de Geijn, H. Finucane, S. Gazal, F. Hormozdiari, T. Amariuta, and X Liu. Annotations capturing cell-type-specific TF binding explain a large fraction of disease heritability. Human Molecular Genetics, 29:1057–1067, 2020.

64. D. Speed, J. Holmes, and D.J Balding. Evaluating and improving heritability models using summary statistics. Nature Genetics, 52:458–462, 2020.

65. S. Chikuma. Ctla-4, an essential immune-checkpoint for t-cell activation. Emerging Concepts Targeting Immune Checkpoints in Cancer and Autoimmunity, pages 99–126, 2017.

66. Y. Zhao et al. Evolving roles for targeting ctla-4 in cancer immunotherapy. Cellular Physiology and Biochemistry, 47(2):721–734, 2018.

67. F. Liu et al. Ctla-4 correlates with immune and clinical characteristics of glioma. Cancer cell international, 20(1):1–10, 2020.

68. M.J. Richer et al. T cell fates zipped up: how the bach2 basic leucine zipper transcriptional repressor directs t cell differentiation and function. The Journal of Immunology, 197(4):1009–1015, 2016.

69. H. Zhang et al. Bach2 deficiency leads to spontaneous expansion of il-4-producing t follicular helper cells and autoimmunity. Frontiers in immunology, 10:2050, 2019.

70. R. Roychoudhuri et al. Bach2 represses effector programs to stabilize t regmediated immune homeostasis. Nature, 498(7455):506–510, 2013.

71. J.D. Cooper et al. Meta-analysis of genome-wide association study data identifies additional type 1 diabetes risk loci. Nature genetics, 40(12):1399–1401, 2008.

72. M.A. Ferreira et al. Identification of il6r and chromosome 11q13. 5 as risk loci for asthma. The Lancet, 378(9795):1006–1014, 2011.

73. D.L. Morris et al. Genome-wide association meta-analysis in chinese and european individuals identifies ten new loci associated with systemic lupus erythematosus. Nature genetics, 48(8):940, 2016.

74. A. Oeckinghaus and S. Ghosh. The nf-kb family of transcription factors and its regulation. Cold Spring Harbor perspectives in biology, 1(4):p.a000034, 2009.

75. R.J. Grumont et al. B lymphocytes differentially use the rel and nuclear factor kb1 (nfkb1) transcription factors to regulate cell cycle progression and apoptosis in quiescent and mitogen-activated cells. Journal of Experimental Medicine, 187(5):663–674, 1998.

76. S. Gerondakis and U. Siebenlist. Roles of the nf-kb pathway in lymphocyte development and function. Cold Spring Harbor perspectives in biology, 2(5):p.a000182, 2010.

77. M.L. Hujoel et al. Disease heritability enrichment of regulatory elements is concentrated in elements with ancient sequence age and conserved function across species. The American Journal of Human Genetics, 104:611–624, 2019.

78. C.A. de Leeuw, et al. MAGMA: generalized gene-set analysis of GWAS data. PLoS computational biology, 11(4):e1004219, 2015.

79. C. Daly and B. Rollins. Monocyte chemoattractant protein-1 (ccl2) in inflammatory disease and adaptive immunity: therapeutic opportunities and controversies. Microcirculation, 10(3-4):pp.247–257, 2003.

80. J. Plskova et al. Interferon-*α*: A key factor in autoimmune disease. Microcirculation, 47:pp.3946–2950, 2006.

81. C. Cardinez et al. Gain-of-function ikbkb mutation causes human combined immune deficiency. J Exp Med, 215(11):pp.2715–2724, 2018.

82. J. Jacobs et al. Cd70: An emerging target in cancer immunotherapy. Pharmacology & therapeutics, 155:pp.1–10, 2015.

83. D.R. Shaffer et al. T cells redirected against cd70 for the immunotherapy of cd70-positive malignancies. *Blood*, The Journal of the American Society of Hematology, 117(16):pp.4304–4314, 2011.

84. Y. Verhoeven et al. The potential and controversy of targeting stat family members in cancer. Seminars in cancer biology, 60:pp.41–56, 2020.

85. K.J. Karczewski et al. The mutational constraint spectrum quantified from variation in 141,456 humans. Nature, 581:434–443, 2020.

86. E.V. Minikel et al. Evaluating drug targets through human loss-of-function genetic variation. Nature, 581:459–464, 2020.

87. K.A. Jagadeesh, K.K. Dey, et al. Identifying disease-critical cell types and cellular processes across the human body by integration of single-cell profiles and human genetics. *bioRxiv, accepted in principle*, Nat Genet, 2021.

88. N. Mancuso et al. Probabilistic fine-mapping of transcriptome-wide association studies. Nature genetics, 51:675–682, 2019.

89. E.M. Weeks et al. Leveraging polygenic enrichments of gene features to predict genes underlying complex traits and diseases. medRxiv, 2020.

90. G. Kichaev et al. Integrating functional data to prioritize causal variants in statistical fine-mapping studies. PLoS genetics, 10(10):e1004722, 2014.

91. W. Chen et al. Incorporating functional annotations for fine-mapping causal variants in a Bayesian framework using summary statistics. Genetics, 204(3):933– 958, 2016.

92. G. Kichaev et al. Improved methods for multi-trait fine mapping of pleiotropic risk loci. Bioinformatics, 33(2):248–255, 2017.

93. J.P. Ray et al. Prioritizing disease and trait causal variants at the TNFAIP3 locus using functional and genomic features. Nature communications, 11(1):1–13, 2020.

94. Y. Hu et al. Leveraging functional annotations in genetic risk prediction for human complex diseases. PLoS computational biology, 13(6):e1005589, 2017.

95. C. Marquez-Luna, et al. LDpred-funct: incorporating functional priors improves polygenic prediction accuracy in UK Biobank and 23andMe data sets. bioRxiv, 2020.

96. R.J. Kinsella et al. Ensembl BioMarts: a hub for data retrieval across taxonomic space. Database, 2011.

97. A. Kundaje et al. Integrative analysis of 111 reference human epigenomes. Nature, 518:317–330, 2015.

98. Y. Liu, A. Sarkar, and M. Kellis. Evidence of a recombination rate valley in human regulatory domains. Genome Biology, page 193, 2017.

99. B.J. Schmiedel et al. Impact of genetic polymorphisms on human immune cell gene expression. Cell, 175:1701–1715, 2018.

100. C.T. Ong and V.G. Corces. Enhancer function: new insights into the regulation of tissue-specific gene expression. Nature Reviews Genetics, 12:283–293, 2011.

101. J.Y. Ko, S. Oh, and K.H. Yoo. Functional enhancers as master regulators of tissue-specific gene regulation and cancer development. Molecules and cells, 40:169–177, 2017.

102. D. Szklarczyk et al. STRING v10: protein–protein interaction networks, integrated over the tree of life. Nucleic acids research, 43:D447–D452, 2014.

103. M.M. Hoffman, J. Ernst, S.P. Wilder, A. Kundaje, R.S. Harris, and M. Libbrecht. A method to predict the impact of regulatory variants from DNA sequence. Nucleic acids research, 41:827–841, 2012.

104. M.M. Hoffman, O.J. Buske, J. Wang, Z. Weng, J.A. Bilmes, and W.S. Noble. Unsupervised pattern discovery in human chromatin structure through genomic segmentation. Nature methods, 9:473–476, 2012.

105. W.J. Kent et al. The human genome browser at ucsc. Genome research, 12(6):996– 1006, 2002.

106. D. Karolchik et al. The ucsc table browser data retrieval tool. Nucleic acids research, 32 (Database Issue):D493–D496, 2004.

107. J. Ernst et al. Mapping and analysis of chromatin state dynamics in nine human cell types. Nature, 473:43–49, 2011.

108. B.M. Javierre et al. Lineage-specific genome architecture links enhancers and non-coding disease variants to target gene promoters. Cell, 167:1369–1384, 2016.

109. H.M. Amemiya, A. Kundaje, and A.P. Boyle. The ENCODE blacklist: identification of problematic regions of the genome. Scientific reports, 9(1):1–5, 2019.

110. ENCODE Project Consortium. An integrated encyclopedia of DNA elements in the human genome. Nature, 489:57–74, 2012.

111. Jan-Renier AJ Moonen et al. KLF4 Recruits SWI/SNF to Increase Chromatin Accessibility and Reprogram the Endothelial Enhancer Landscape under Laminar Shear Stress. bioRxiv, 2020.

112. GTEx Consortium. Genetic effects on gene expression across human tissues. Nature, 550(7675):204–213, 2017.

113. L. Jostins et al. Host-microbe interactions have shaped the genetic architecture of inflammatory bowel disease. Nature, 491:119–124, 2012.

114. Y. Okada et al. Genetics of rheumatoid arthritis contributes to biology and drug discovery. Nature, 506:376–381, 2014.

115. J. Bentham et al. Genetic association analyses implicate aberrant regulation of innate and adaptive immunity genes in the pathogenesis of systemic lupus erythematosus. Nature genetics, 47(12):1457–1464, 2015.

116. P.C. Dubois et al. Multiple common variants for celiac disease influencing immune gene expression. Nature genetics, 42(4):295–302, 2010.

117. A. Kamburov et al. ConsensusPathDB: toward a more complete picture of cell biology. Nucleic acids research, 39(suppl 1):D712–D717, 2010.

118. E.K. Speliotes et al. Association analyses of 249,796 individuals reveal 18 new loci associated with body mass index. Nature genetics, 42:937–948, 2010.

119. A. Okbay et al. Genome-wide association study identifies 74 loci associated with educational attainment. Nature, 533:539–542, 2016.

120. S. Ripke et al. Biological insights from 108 schizophrenia-associated genetic loci. Nature, 511(7510):421–427, 2014.

121. Tobacco and . Genetics Consortium. Genome-wide meta-analyses identify multiple loci associated with smoking behavior. Nature genetics, 42:441–447, 2010.

122. Psychiatric GWAS Consortium Bipolar Disorder Working Group. Large-scale genome-wide association analysis of bipolar disorder identifies a new susceptibility locus near odz4. Nature genetics, 43:977–983, 2011.

123. N. Barban et al. Genome-wide analysis identifies 12 loci influencing human reproductive behavior. Nature genetics, 48(12):1462–1472, 2016.

124. V. Boraska et al. A genome-wide association study of anorexia nervosa. Molecular psychiatry, 19(10):1085–1094, 2014.

125. 125. Cross-Disorder Group of the Psychiatric Genomics Consortium. Identification of risk loci with shared effects on five major psychiatric disorders: a genome-wide analysis. The Lancet, 381(9875):1371–1379, 2013.

126. H. Schunkert et al. Large-scale association analysis identifies 13 new susceptibility loci for coronary artery disease. Nature genetics, 43:333–338, 2011.

127. T.M. Teslovich et al. Biological, clinical and population relevance of 95 loci for blood lipids. Nature, 466:707–713, 2010.

128. H.L. Allen et al. Hundreds of variants clustered in genomic loci and biological pathways affect human height. Nature, 467:832–838, 2010.

129. A.P. Morris et al. Large-scale association analysis provides insights into the genetic architecture and pathophysiology of type 2 diabetes. Nature genetics, 44:981–990, 2012.

